# Exploring dimension-reduced embeddings with Sleepwalk

**DOI:** 10.1101/603589

**Authors:** Svetlana Ovchinnikova, Simon Anders

## Abstract

Dimension-reduction methods, such as t-SNE or UMAP, are widely used when exploring high-dimensional data describing many entities, e.g., RNA-seq data for many single cells. However, dimension reduction is commonly prone to introducing artefacts, and we hence need means to see where a dimension-reduced embedding is a faithful representation of the local neighbourhood and where it is not.

We present Sleepwalk, a simple but powerful tool that allows the user to interactively explore an embedding, using colour to depict original or any other distances from all points to the cell under the mouse cursor. We show how this approach not only highlights distortions, but also reveals otherwise hidden characteristics of the data, and how Sleepwalk’s comparative modes help integrate multi-sample data and understand differences between embedding and preprocessing methods. Sleepwalk is a versatile and intuitive tool that unlocks the full power of dimension reduction and will be of value not only in single-cell RNA-seq but also in any other area with matrix-shaped big data.

## Introduction

Whenever one is presented with large amounts of data, producing a suitable plot to get an overview is an important first step. So-called dimension reduction methods are commonly used. For example, in high-throughput transcriptomics projects using expression microarrays or RNA-seq, it is common practice, especially when working with many samples, to perform principal component analysis (PCA) on a suitably normalized and transformed expression matrix and then plot the samples’ first two principal components as a scatter plot. Of course, PCA has more uses than just providing such an overview plot (See Ringnér (2008) for a primer.), but nevertheless, the user’s expectation is often simply that samples with similar expression profile should appear close together (“cluster together”), while samples with strong differences should appear farther apart. PCA’s popularity in biology notwithstanding, the literature offers many methods designed specifically with this goal in mind, with the best-known classic example perhaps being classical multidimensional scaling (classical MDS, also known as principal coordinate analysis, PCoA), Kruskall’s non-metric multidimensional scaling (Kruskal 1964) and Kohonen’s self-organizing maps (SOM) (Kohonen 1982).

The recent rapid progress of single-cell RNA-seq methods, now enabling the measurement of expression profiles of thousands of individual cells in a sample, has renewed biologists’ interest in dimension reduction methods. Here, t-distributed stochastic neighbour embedding (t-SNE, van der Maaten and Hinton (2008), Figure 1) has become a de-facto standard, with Uniform Manifold Approximation and Projection (UMAP, McInnes et al. (2018)) currently gaining popularity as an alternative. Other dimension reduction methods, developed specifically for single-cell RNA-Seq include Destiny (Angerer et al. 2016) (an method based on diffusion maps (Coifman and Lafon 2006)), the Monocle methods (Trapnell et al. 2014; Qiu et al. 2017), DDRTree (Mao et al. 2015) and more. (See Nguyen and Holmes (2019) for a recent review.)

**Figure 1:**
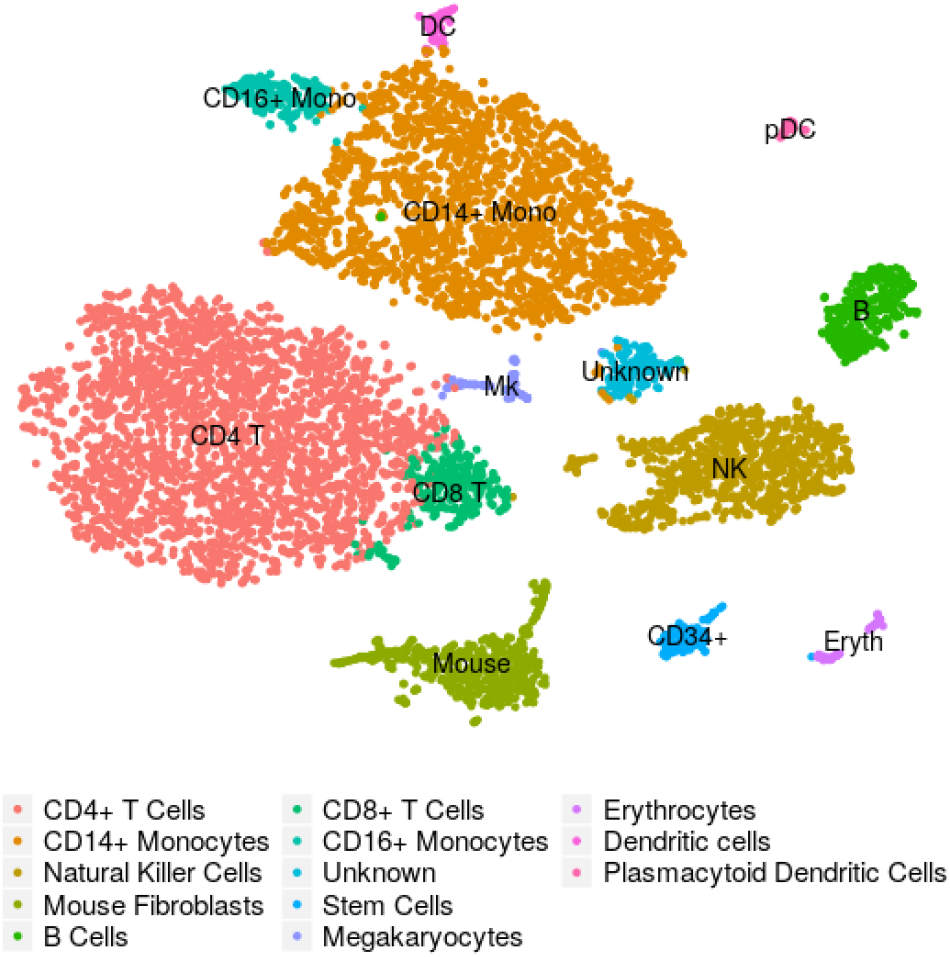
Example of a t-SNE plot: These are cord-blood mononuclear cells studied by Stoeckius et al. (2017). The embedding and the assignment of cell types have been taken from the Seurat (Butler et al. 2018) tutorial that uses this data set as example (Satija Lab 2018).

These varied methods have been developed with different design goals: for example, some methods strive to primarily preserve neighbourhood, others to represent the overall structure or larger-scale relations. Nevertheless, when using any of them in the field of single-cell transcriptomics, practitioner’s primary expectation is usually that cells depicted close to each other or within the same apparent structure or cluster have more or less similar expression profiles, while cells depicted in different regions of the plot or in different structures are more different. In other words, it is the preservation of neighbourhood relationships that is of importance. The term “neighbourhood” should here be understood as follows: We consider a high-dimensional space, the so-called feature space, in which each dimension corresponds to one gene and each cell is represented by a point, whose coordinates along the many dimensions are given by the expression strength of the corresponding gene. Two cells with similar expression profile will hence have similar coordinates and thus will be close to each other in feature space. Around each cell, we can imagine a hypersphere of nearby points, and consider all cells with the hypersphere as neighbours.

Any attempt to provide a two dimensional representation of the neighbourhood relations in this high-dimensional space will have to face what van der Maaten and Hinton (2008) called the “crowding problem”. The volume of a high-dimensional sphere is exponentially larger than the area of a two-dimensional circle, and therefore, a cell can easily have many more close neighbours in feature-space than cells can be drawn within a sufficiently small circle around the point representing the cell in two dimensional space.

This is, of course, not the only obstacle in achieving a faithful two dimensional representation of feature space, and the many possible kinds of distortions have been widely discussed in the literature. (See e.g. Aupetit (2007)) and Kaski et al. (2003).) In single-cell sequencing, it is, however, of particular relevance: given a dimension-reduced representation such as a t-SNE or UMAP embedding, how can we know for a specific cell of interest how far its neighbourhood reaches? Knowing this is of paramount importance to correctly interpret an embedding.

## Results

### The Sleepwalk app

Here, we present “Sleepwalk”, an interactive tool that provides an intuitive solution to the task just outlined.

It works as follows: The user provides an embedding, i.e., the two-dimensional coordinates output by a dimension-reducing method, as well as information on the distances between cells in feature space in some suitable metric or their coordinates in a appropriately transformed feature space. Whenever the user moves the mouse cursor over a cell, all cells are coloured according to their distance to this cell in feature space, thus indicating the cell’s closest neighbours with the strongest colour (Figure 2A-C). By moving the mouse over all the cells in the plot, the user can hence quickly obtain an intuitive overview over how neighbourhood may have been rendered differently in different regions of the plot. Buttons are provided to adjust the colour scale so that the user can chose which feature-space distance should be considered as “close neighbourhood” and hence given the strong (dark, green) colours.

**Figure 2:**
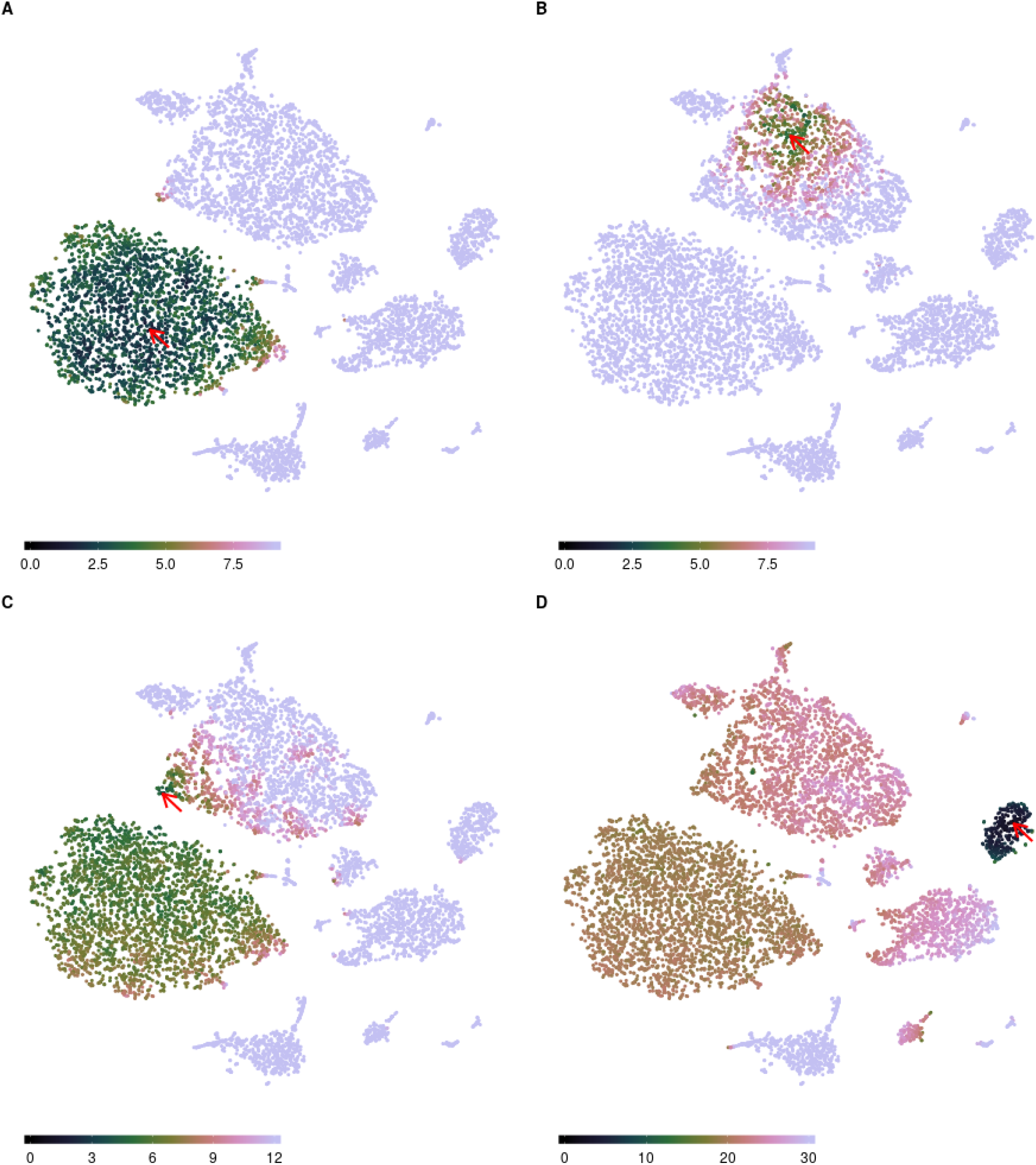
The “Sleepwalk” app, being used to explore the t-SNE rendition of the cord-blood data set from Figure 1. The plots here are snapshots of a running “Sleepwalk” app; a live version can be found at https://anders-biostat.github.io/sleepwalk/supplementary/. The red arrow shows the current mouse position. (A) By moving the mouse cursor through the embedding, we find, e.g., that the CD4+ T cell cluster is very tight and homogeneous, as can be seen from the fact that all cells show a colour indicating that they are all close to each other. (B) The monocyte cluster, in contrast shows much more heterogeneity, when comparing the colouring at the same colour scale: now only few monocytes are coloured green and are hence as similar to the cell under the mouse cursor as most of the T cells were in (B). (C) Placing the mouse on this small tip of the monocyte cluster reveals that the cells there are more similar to the T cells than to the other monocytes, indicating that the cluster boundary might be inaccurate in both the t-SNE rendition and the SNN clustering on which the Seurat workflow’s cell-type assignment is based. (D) With the colour scale set to a wider distance range, we can assess similarities *between* clusters: As expected, B cells are somewhat similar to T cells, less so to NK cells and monocytes, and distant to erythrocytes and the spiked-in mouse cells.

We invite the reader to pause here and try Sleepwalk himself or herself. At https://anders-biostat.github.io/sleepwalk/supplementary/ (and also in this paper’s supplemental HTML), live interactive Sleepwalk renditions of Figure 2 (and also of all the subsequent figures discussed below) are given. The Sleepwalk app runs in any Javascript-enabled web browser, i.e., it suffices to open the page or file in a browser, without need to install anything.

### Exploring an embedding

Sleepwalk makes aspects visible that are not apparent from a dimension reduction alone. For example, the two large clusters under the cursor in Figure 2A and Figure 2B have quite different characteristic. In the T cell cluster (Figure 2A), most of the cells are very close to each other: the cluster shows up as a large green cloud no matter where one points the mouse. The monocyte cluster (Figure 2B), however, spreads over larger distances: only a part shows up in green, which “follows” the mouse. In a static t-SNE plot (such as Figure 1), this cannot be seen.

We can also check the cluster borders and discover, for instance (Figure 2C), that some cells in the monocyte cluster are more similar to those in the T cell cluster than to those in their own cluster. They may have been assigned the wrong cell type, or might be doublets. Thus, the Sleepwalk exploration can alert the analyst to the need for further investigation of possibly misleading features of a dimension-reduced embedding.

In 2A-C, the colour scale was left at the automatically chosen range of only very small distances. When switching the colour scale to a wider distance range, we can also see here how relationships *between* clusters (Figure 2D) appear in the supplied distance values: we see which clusters are more and which are less similar to each other – an information that a static t-SNE does not show, due to the method’s design focus on faithful representation only of neighbourhoods. Care is needed here, however: once the considered distances exceed what one might consider as “close neighbourhood”, the choice of distance metric used will strongly influence interpretability of the visualization, as we discuss below.

### Feature-space distances

The colours in Sleepwalk are meant to indicate similarity or dissimilarity between the cells’ expression profiles, quantified as distances. There are multiple suggestions for useful distance measures in the literature, and the users can provide whichever they prefer. To produce the t-SNE embedding in Figure 1, we followed the Seurat workflow (Satija Lab 2018), which calculates distances in a specific manner (Methods), and these are then also used by the t-SNE routine. We have also used these same distances to colour the points in the Sleepwalk rendition (Figure 2); thus allowing us to see directly where t-SNE succeeded and where it failed in its design goal of preserving the neighbourhood relation in its input data.

t-SNE uses a flexible approach to define the distance scale over which cells are considered neighbours: it adjusts the distance scale for each cell such that all cells have approximately the same number of neighbours (the so-called perplexity). Sleepwalk, in contrast, uses a fixed distance scale. This is on purpose: it allows us to note where the neighbourhood has longer or shorter range (as shown in the comparison of Figures 2A and 2B). The app offers two buttons to increase or decrease the scale of the distance-to-colour mapping, allowing the user to manually chose what distance should be considered as close.

### Comparing embeddings

With the availability of choice in dimension reduction methods, the question arises which one to use. Benchmark comparisons may address this question in general; see for example Becht et al. (2019) for a comparison of UMAP with t-SNE and related methods. When working on a specific dataset, however, simply calculating multiple embeddings and comparing them side by side might be even more helpful. We demonstrate this here using data from a study of the development of the mouse cerebellum (Carter et al. 2018). In Figure 3, we show cells from development time point E13.5, first visualized with t-SNE (Figure 3A), then with UMAP (Figure 3B). The live apps in the supplemental HTML show, in addition to this, also the same comparison for the cord-blood data of Figures 1 and 2.

**Figure 3:**
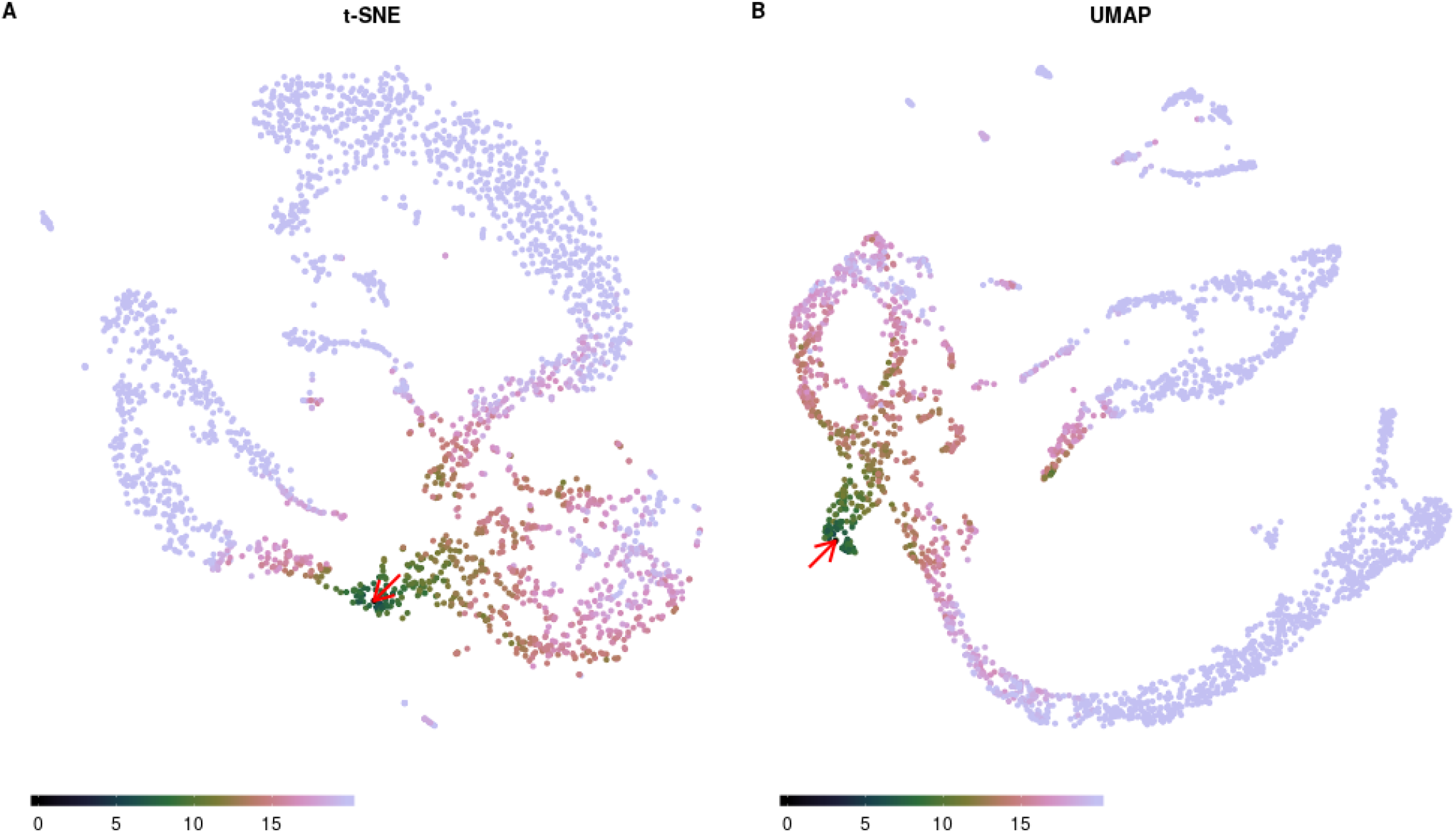
Sleepwalk being used to compare two embeddings of the same single-cell data of a developing murine cerebellum at embryonal time point E13.5 (Carter et al. 2018): t-SNE on the left and UMAP on the right. The user can explore one embedding in the same ways as in Figure 2, while all other embeddings that are displayed concurrently, are “slaved” to the one under the mouse cursor: each cell has the same colour in all embeddings. The red arrow shows the current mouse position. A live version (for this data and for the cord-blood data) can be found in the supplemental HTML file.

To compare the two embedding, we need, at minimum, a way to see which points in the two plots correspond to the same cells. A classical approach is “brushing” (Becker and Cleveland 1987): selecting with the mouse a group of adjacently depicted cells in one plot causes them to be highlighted in the other one, too. Sleepwalk adapts this idea, but instead of the usual brush, we simply use in all embeddings the same colour for points corresponding to the same cell. Moving the mouse over points in one plot then highlights the neighbourhood structure induced by the feature-space distance chosen for that embedding not only there but also in all displayed embeddings and so links them. This allows us to see for a structure in one embedding whether there are corresponding structures in the other embeddings.

In the example shown in Figure 3, we see a clear correspondence between the major structures generated by t-SNE and by UMAP. Even the arrangement of cells within these structures is the same, which one can follow in the life version of the app. There are, however, also differences: The cells at the mouse position in Figure 3 are part of the connecting “filament” in the t-SNE embedding, but lie in an external “protrusion” in the UMAP. Further exploration in the live version of Figure 3 can suggest that UMAP forced the two branches to intersect while still trying to repel cells of different lineages away from each other (note the gap between the highlighted branch in Figure 3B). This is another example of the dimensionality reduction artefacts that are hard to notice from a static image, but can be uncovered with “Sleepwalk”.

### Comparing samples

Until recently, most single-cell RNA-seq studies analysed only a single sample comprising many cells. Yet, the full value of the technique might become apparent only when it is used to compare between many samples. One currently popular approach to do so visually is to simply combine the data from the cells of all samples into one large expression matrix and perform t-SNE or UMAP on this. Often, global differences between samples, typically due to technical effects (Tung et al. 2017), will prevent similar cells from different samples to appear in the same cluster or structure in the dimension-reduced embedding. Methods to automatically remove such sample-to-sample differences (e.g., the CCA-based method in Butler et al. (2018) and the MNN method in Haghverdi et al. (2018)) address this issue, but will not work always and may risk also removing biological signal.

An visual alternative is to produce a dimension-reduced embedding separately for each sample, and then try to find correspondences between the features in these. In Figure 4, we show how Sleepwalk allows to perform such an exploration comparing UMAP renderings for the two E13.5 and one of the E14.5 samples of the mouse cerebellum data set. Exploring the data with the mouse shows the two replicas of E13.5 samples (Figures 4A and 4B) are almost identical. The two branches (GABAergic and glutamatergic neurons) can be easily followed from the early progenitor cells up to the most differentiated ones. Comparing between the two E13.5 replicates reveals which aspects of the peculiar two-pronged shape of the glutamatergic branch is simply due to random variation and what seems reproducible. In the later E14.5 sample, the branches have disconnected from the progenitor cells, but Sleepwalk allows us to still identify corresponding cells. Sleepwalk can show that the GABAergic lineage is differentiated further in the E14.5 than in E13.5 samples, as the endpoint of the branch in E14.5 corresponds to an intermediate point in E13.5. Sleepwalk allows one to discover such details immediately, with minimal effort. Of course, such a visual exploration cannot replace a tailored detailed analysis but it does provide a starting point and a first overview.

**Figure 4:**
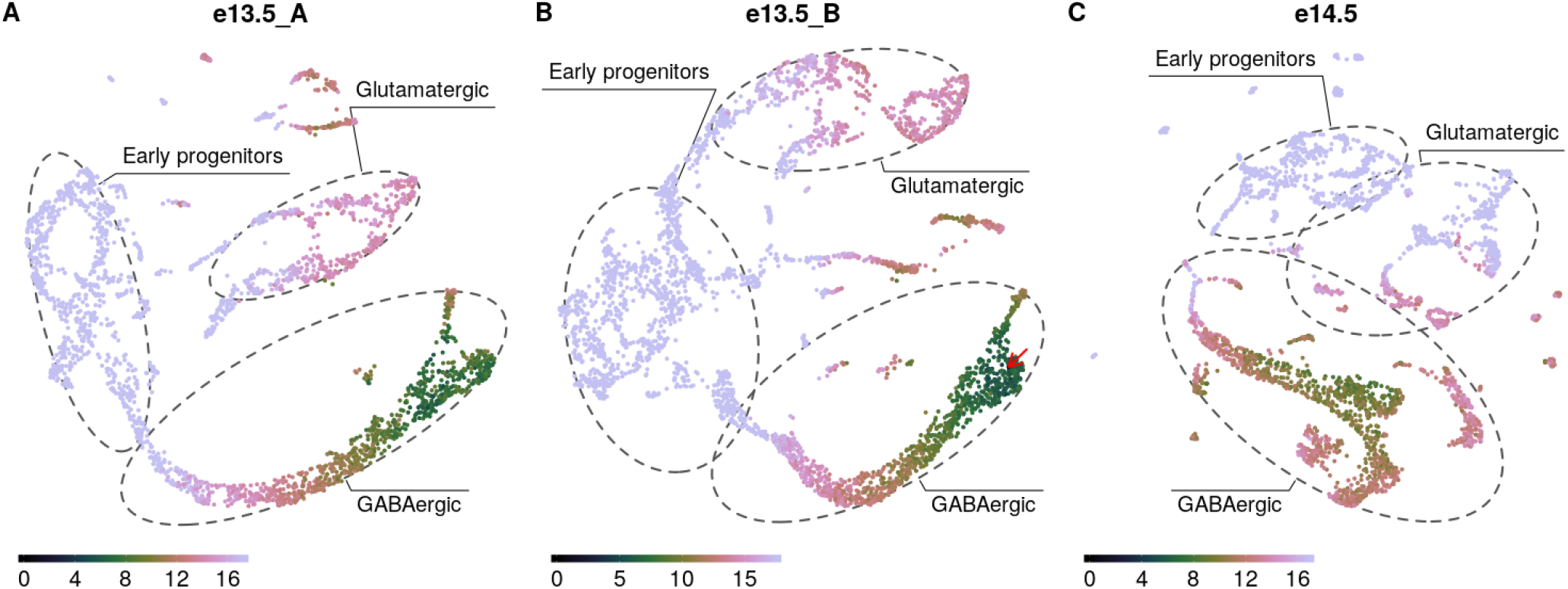
Sleepwalk in multi-sample mode, comparing three samples of a developing murine cerebellum: two samples of two different mice embryos at time point E13.5 (A, B) and the third (C) from E14.5. The red arrow shows the current mouse position. The dashed grey lines roughly indicate two different lineages and their common progenitor cells (expression of the marker genes that we used to draw the boundaries is shown in the Supplemental Figure S1). By following the GABAergic branch one can notice that its very tip in the E13.5 samples corresponds to cells in the middle of the branch in E14.5, indicating that the cells have differentiated further during the elapsed day.

Crucially, using Sleepwalk’s multi-sample comparison mode does not require any removal of global sample-to-sample differences with batch-effect correction methods. If the user selects a cell with the mouse in one sample, the cells that are similar to it will be highlighted, both in the same sample as well as in all other samples. This works even if the cells in the other samples seem more distant, due to the additional sampleto-sample distance; we only might need to increase the scale of the distance-to-colour mapping for the cross-sample comparisons.

### Comparing distance metrics

In the examples discussed so far, we have always coloured cells according to the default distance calculated by the Seurat workflow, namely the Euclidean distance in the space spanned by the first few principal components according to a PCA performed after certain preprocessing. The reason for this was not that this specific distance metric should be considered more correct or more “true” than any of the alternatives discussed in the literature, but simply because it is the distance metric that has been used as input to t-SNE and UMAP when calculating the discussed embeddings.

While this specific distance metric is popular due it appearing in standard workflows such as Seurat’s, this is of course no reason to consider as more correct or “true” than possible alternatives or modifications. For instance, we may either choose to use all genes in the distance calculation, or only some genes, which may either be chosen for having high expression or high signal-to-noise ratio, or perhaps chosen, via manual curation, as especially informative with respect to cell type or state. We may use the genes as they are or aggregate them before into meta- or eigen-genes, e.g., by a principal component analysis (as done in the Seurat workflow). The way how the expression data has been transformed, normalized or preprocessed can be understood as part of the choice of distance metric. The last, but not the least choice is, of course, the metric itself. There are numerous ways how to calculate distances from the selected and possibly preprocessed set of features. Besides Euclidean, one can calculate angular (“cosine”) or correlation distance or use kernel functions (Phillips and Venkatasubramanian 2011) as it has been done, for instance in Wang et al. (2017). Metrics can even be learned to suit the specific task researcher has in mind (Yang and Jin 2006). Dimensionality reduction techniques are typically based on the assumption that in feature space cells are located on the surface of a smooth manifold. The methods attempt to learn the manifold and then to replace the original distance with the geodesic one (i.e., distance within the manifold) (Cayton 2005; Moon et al. 2018). Diffusion distances (Coifman and Lafon 2006) are popular way of obtaining a manifold-following distance in a simple and efficient manner.

Some of these metrics may yield similar results, others can drastically change cell-to-cell distances. In order to impact of the choice of metric, Sleepwalk offers a variant to the mode for comparing embeddings described above, in which points that correspond to the same cell will get different colours in different panels of the app, each showing the same embedding but having a different distance matrix assigned to it. By hovering the mouse over a cell, the user can see how the cells neighbourhoods differ between the distance metrics.

Figure 5 shows Sleepwalk in the distance comparison mode. The live version of this figure can be found in the supplement to the paper. Once again, we use the murine cerebellum dataset (specifically, stage E13.5, visualized with UMAP) as an example. As before, we used the 2131 genes chosen by the Seurat workflow as “variable”, and then calculated distance matrices using four metrics: (i) Euclidean distance based directly on the normalized and logarithmised expressions of these genes, (ii) Euclidean distance in the space spanned by the first 50 principal components of a PCA performed using the variable genes, (iii) diffusion distance based on directly on the genes’ expression or (iv) on the first 50 principal components. As expected, Euclidean distance calculated on all variable genes (i) is almost useless when applied to so many dimensions. Most of the distances are condensed around some median value, making it almost impossible to distinguish any patterns in the data. However, all other distances are already good enough to see the two developmental branches, with perhaps the diffusion distance separating them most clearly.

**Figure 5:**
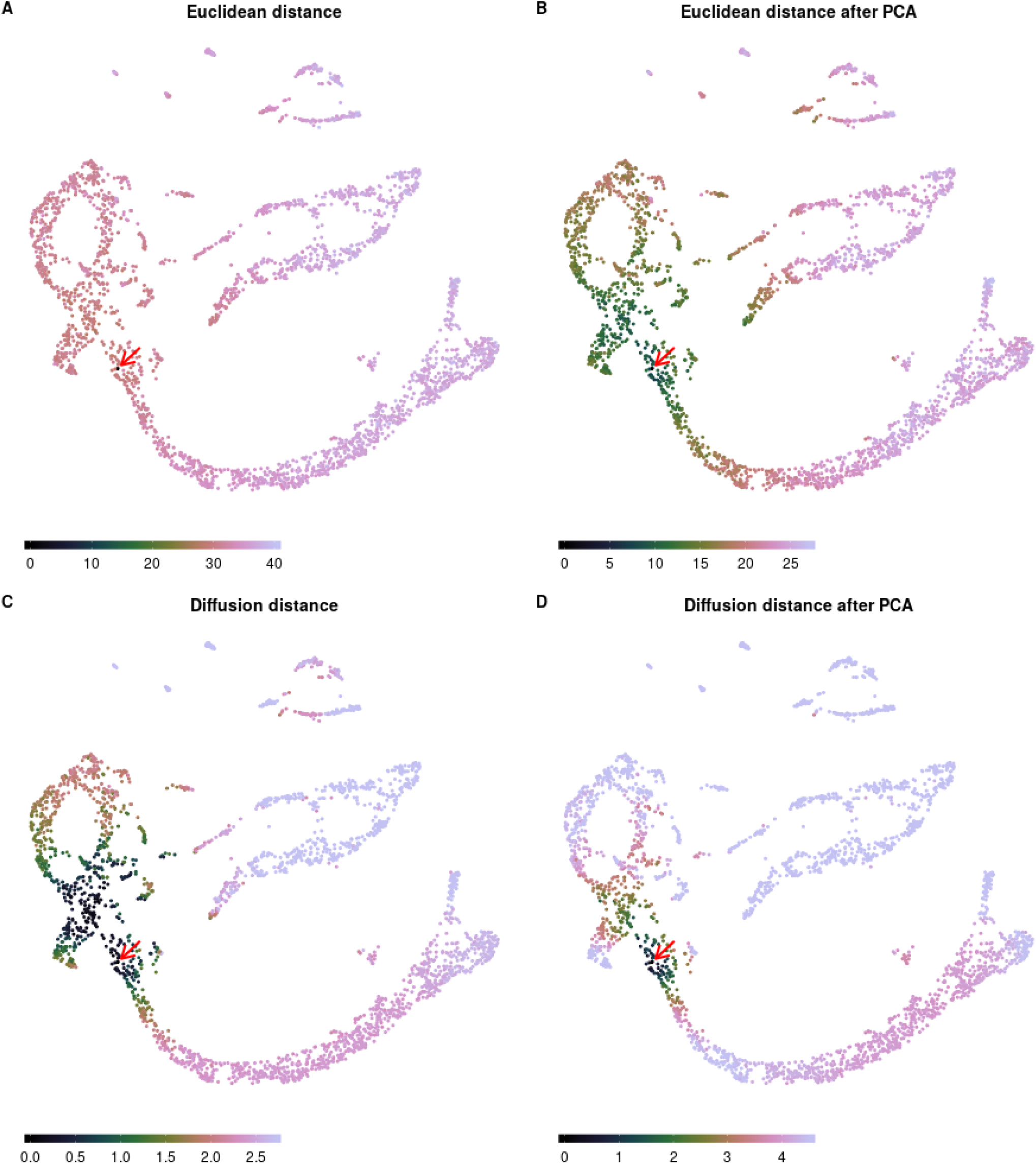
Metric comparison using Sleepwalk. All four panels now show the same embedding: A UMAP visualization of the E13.5 sample of murine cerebellum. The cells are now coloured based on four different metrics: Euclidean distance based directly on the normalized and logarithmised genes’ expressions (A); Euclidean distance in the space spanned by the first 50 principal components of a PCA performed using the variable genes (B); diffusion distance based on directly on the genes’ expression (C) or on the first 50 principal components (D). The red arrow shows the current mouse position, which is on the intersection of GABAergic and glutamatergic lineages. Colour scales were adjusted so that roughly stretch along the entire selected branch. The spread of colouring onto the other developmental branch shows how good is the metric in separating the two lineages.

In the live version in the supplemental HTML file, we also demonstrate a metric comparison using the cordblood dataset. There, the difference in the metrics does not lie in the details of its calculation but stems from using different input data, as each cell has been assessed in two modalities, and distances are calculated either from the single-cell RNA-Seq or from the single-cell proteomics (CITE-Seq epitome) aspect of the data.

One should keep in mind that, in the example of Figure 5, we are comparing the four metrics not just to each other, but also to the fifth one: The distance that was used to generate the embedding. UMAP is one of the manifold learning dimensionality reduction techniques, and as such it might be more similar to diffusion distances than to the Euclidean metric, even though the Euclidean distance in PCA space (distance (ii)) was used as input for the UMAP process. Taking this together, one might expect the combination of diffusion distance and PCA to correspond especially well with the embedding. Which of the four metrics, however, should be considered “best”, is a completely different question, as the suitability of metric will depend on the task at hand. While a metric can be effective in separating specific cell types, it might at the same time fail to arrange cells by their cell cycle stage (Buettner et al. 2015), and this can be considered a good or a bad thing, depending on whether differences due to cell cycle are considered a nuisance or a topic of interest in one’s experiment. We therefore do not wish to provide opinions or guidance on this question in the present paper. What Sleepwalk does offer here is a means to explore differences between metrics and embeddings and understand them, not necessarily to perform benchmarks. Once a researcher is aware of such differences, it is up to him or her to decide if they affect data interpretation.

### Beyond single-cell transcriptomics

In all the examples discussed so far, the points correspond to individual cells in samples assessed with single-cell transcriptomics. Another important use case for dimension-reduction methods are large-scale studies comprising dozens or even hundreds of samples. Brawand et al. (2011), for example, describe a collection of 131 bulk RNA-seq data sets comparing organ samples from several species and provide an overview PCA plot as their Figure 1, Dietrich et al. (2018) use a t-SNE plot to illustrate similarities and differences between their 246 blood cancer samples. Clearly, Sleepwalk can also be useful to explore dimension-reduced embeddings arising in such applications.

Research on dimension reduction originated in the machine learning field, with the original applications being the study of training data for machine learning applications. Of course, in this area, as well as in other applications of dimension reduction, Sleepwalk should also prove useful.

### Usage

Sleepwalk is provided as a package for the statistical programming language R (R Core Team 2019). The main function of the package is also called “sleepwalk”. The user provides it with the 2D coordinates for each object (cell) in the embedding and a square matrix of cell-to-cell distances, or, alternative to the latter, a data matrix from which sleepwalk can calculate Euclidean or angular distances. For both these parameters, the user can also supply multiple matrices in order to display multiple embeddings concurrently for comparison. This can be done either such that each embedding represents the same objects (as in Figure 3), or that each embedding represents a different set of objects but distances are given also between objects in different embeddings (as in Figure 4).

Sleepwalk can easily be used in combination with other single-cell analysis frameworks. Visualizing, for example, a Seurat data object can be done with one line of code, as explained on the documentation web page.

By default, the sleepwalk function displays the visualization app in a web browser. Alternatively, it can also write it into an HTML file, which can then be opened with any web browser without the need of having R or the Sleepwalk package running or even installed. This is useful when an analyst wishes to share a Sleepwalk visualization with colleagues or provide it on a web page or in a paper supplement.

For a description of further options of the function, please see the documentation.

The app offers a “lasso” functionality: The user can encircle a group of points with the mouse, and the indices of these points are then reported back to the R session, where they can be queried with a callback function. This is helpful if the analyst spots an interesting set of cells while exploring an embedding and wishes to perform further analysis on them.

We also mention the “slw snapshot” function, which produces static plots, like the figures in this paper.

## Discussion and Conclusion

Dimension-reduced embedding such as those provided by t-SNE and UMAP have become a core tool in single-cell transcriptomics. They provide an overview of a study, help to check for expected and unexpected features in the data, allow researchers to form new hypotheses and to plan and organise the subsequent analysis. As they generally contain artefacts, a common concern is that these plots may be over-interpreted.

Dimension reduction is a research area with a rich history, long predating the use of these techniques for single-cell biology. The issue with distortions has been long discussed, with the possible distortions being classified (e.g., Aupetit (2007)) and quantified (e.g., Kaski et al. (2003)), and advice on careful interpretation derived from these (e.g., Wattenberg et al. (2016)). To visually alert the viewer to distortions, some authors have suggested to colour each point by its so-called stress, i.e., the deviation of the point’s on-screen distance to the other points from the distances in feature space (Seifert et al. 2010) and others to colour the area around the points according to the amount of compression or stretching that the manifold underwent locally due to projection (Lespinats and Aupetit 2011).

Such visualizations are useful tools for developing and improving dimension reducing methods. Our approach, however, offers a novel aspect that is crucial: rather than merely alerting the user to distortions, Sleepwalk allows the user to directly see the underlying “truth” for the selected cell. This is possible due to our use of interactivity: by allowing the user to rapidly move focus from cell to cell, and the app instantly following in redrawing the colours, we are effectively escaping the confines of a two-dimensional representation (or, three-dimensional, if we also count static colouring as a dimension).

We have shown how this novel approach gives insights into dimension-reduced embeddings that would otherwise stay hidden and thus solves a core problem in the practical use of dimension reduction methods. We envision that Sleepwalk will be used in two manners: first as a tool of exploratory data analysis, helping researchers to better understand their data, but also second as a reporting and communication tool, allowing researchers to present their results in a more transparent way. For this latter application, Sleepwalk’s ability to produce stand-alone HTML pages is crucial, as these pages can then be used, e.g., as supplements to publications, where they allow readers to check embeddings themselves, without the need to install any software.

We should be clear that a visual, interactive data exploration with Sleepwalk does not replace formal inference but complements or typically precedes it. Once one has formed a hypothesis about one’s data using Sleepwalk, one should employ suitable formal analysis methods, such as statistical hypotheses tests, to confirm them. That analysis will then typically be done on the full, high-dimensional data. Dimension reduction methods are data reduction methods: this sacrifice of data is done to allow for visual inspection, but is a hindrance for any numerical analysis.

The principle of Sleepwalk is useful not only for inspection of a single data set but also lends itself for generalization to comparative tasks. We have shown several possible modes of comparison: between different embeddings of the same data, between embeddings from several samples, and between different ways of preprocessing the data and obtaining distances. The comparison between samples will find direct application in any study working with multiple samples, the other two are useful in method selection and in method development, as they allow for comparison of data processing pipelines.

We therefore expect that Sleepwalk will find broad use not only in single-cell transcriptomics, but essentially all instances of big data where experimental units (cells, samples, or the like) are described in a highdimensional feature space.

## Methods

### Sleepwalk implementation

Sleepwalk is written in JavaScript, using the D3.js data visualization framework (Bostock et al. 2011). JavaScript was chosen because it is available on all common platforms, usually without need to install anything, thus enabling the standalone HTML feature, because writing the app with D3.js was convenient, and because the JavaScript engines of most web browsers offer very good performance, enabling smooth rendering of the colour changes and thus the instant interactive feed-back that is required to provide an intuitive user experience.

A thin wrapper of R code around the JavaScript code turns Sleepwalk into an R package, allowing for convenient integration into currently popular workflows for single-cell analysis like Seurat. We use the httpuv R package (Cheng et al. 2019) to bridge between the R session and the web browser. It sets up a simple local server to serve the app to the browser and then uses its implementation of the WebSocket protocol (Fette and Melnikov 2011) to keep open a communication channel between R session and web browser. This allows the app to report back to the R session when the user has selected points using the lasso feature or to make snapshots and change the state of the app from an R session. The colour scheme used to depict distances is the “cubehelix” palette, a colour map originally developed for astronomy and optimized for good visual separation between levels throughout its dynamic range (Green 2011).

### Example data

#### Cord-blood data set

The cord-blood data from Stoeckius et al. (2017) are available at Gene Expression Omnibus (GEO) via accession GSE100866. The raw UMI counts were processed following the Seurat workflow proposed for exactly this data set (Satija Lab 2018). Data were normalized and log transformed. 976 variable genes were detected with y.cutoff = 0.5. These genes were scaled and used for principle components analysis. For further analysis, the first 13 principal components were used, which explain around 23% of the total variance. The t-SNE (Figures 1, 2, supplemental HTML file) and UMAP (Figure 2, supplemental HTML) embeddings were calculated using the default functions from the Seurat package. The assignments of cell types to clusters was taken, too, from the Seurat tutorial workflow (Satija Lab 2018). Normalized and log-transformed epitome data were used to calculate Euclidean distances for the distance comparison (all the scripts are available on GitHub). The resulting Seurat object can be downloaded from Figshare (doi:10.6084/m9.figshare.7908059).

#### Murine cerebellum data set

The raw sequence data from Carter et al. (2018) were downloaded from the European Nucleotide Archive under the accession number PRJEB23051. The reads were aligned and counted using the Cell Ranger (10x Genomics 2019) software (output files are accessible from Figshare, doi:10.6084/m9.figshare.7910483). Some genes and droplets were filtered out following the Methods section of Carter et al. (2018). We removed all the cells with more than 10% of all UMIs coming from mitochondrial genes. We then removed all ribosomal and mitochondrial genes. Finally, only cells that contain from 3500 to 15000 UMIs were kept. Lastly, we omitted all genes with zero expression in all the cells. The filtered raw data were then used to create Seurat objects that can be found at Figshare, doi:10.6084/m9.figshare.7910483), or on GitHub https://github.com/anders-biostat/sleepwalk/tree/paper in the “data” folder. We used Seurat to normalise and log-transform raw counts and find variable genes. The irlba R package (Baglama et al. 2019; Baglama and Reichel 2005) was used to generate a PCA embedding of the data (each sample separately, only variable genes). The first 50 principal components were used for further analysis. To render a t-SNE embedding we used the Rtsne package by Krijthe (2015), a wrapper around the code by van der Maaten (2014). The uwot package by Melville (2019) was used for UMAP embeddings. To calculate distances between cells from different samples (Figure 4), we used the variable genes shared between all the samples and produces a PCA embedding for all the cells. Euclidean distances in the space defined by the first 50 principal components are used to colour the points. To distinguish early progenitors form further differentiated cells of glutamatergic and GABAergic we used the following marker genes: Msx3 for early progenitors, Meis2 for the glutamatergic lineage, adn Lhx5 for the GABAergic lineage (Supplemental Figure S1). Countours in Figure 4 are drawn to include around 90% of cells that express each of the markers above a certain threshold using the “geom mark ellipse” function of the ggforce package (Pedersen 2019).

Calculation of diffusion distance in Figure 5 is based on the destiny package by Angerer et al. (2016). We used internal functions of the package to find nearest neighbours, to calculate local diffusion scale parameters “sigma” and to get initial transition probabilities. Then we manually propagated the diffusion with 16 time steps and calculated resulting distances.

## Supporting information

Supplemental HTML page

## Data and Software Availability

All data sets used in this paper where taken from published works and can be downloaded from the cited references (Stoeckius et al. 2017; Carter et al. 2018). While processing the data we also stored intermediate steps that are now available at doi:10.6084/m9.figshare.7908059.v1 (CiteSeq data), doi:10.6084/m9.figshare.7910483.v1 (three single-cell samples of developing murine cerebellum) and doi:10.6084/m9.figshare.7910504.v2 (some of the 2D embeddings calculated for the Sleepwalk examples). Scripts to generate these files, all the figures in the paper, and interactive apps are available on GitHub at https://github.com/anders-biostat/sleepwalk/tree/paper. Demonstration HTML file is available in the supplement and at https://anders-biostat.github.io/sleepwalk/supplementary/. The “sleepwalk” R package is available on the Comprehensive R Archive Network (CRAN) at https://cran.r-project.org/package=sleepwalk, released as opensource software under the GNU General Public License v3 (or later). Documentation, installation instructions and examples for Sleepwalk can be found on the project web page at https://anders-biostat.github.io/sleepwalk/. The source code is available on GitHub (https://github.com/anders-biostat/sleepwalk).

## Acknowledgments

We thank Kevin Leiss for helpful discussions and assistance in preprocessing the cerebellum data with CellRanger. This work was funded by the Deutsche Forschungsgemeinschaft (DFG) via the collaborative research centres SFB 1036 (project no. 201348542) and SFB 1366 (project no. 394046768).

Notes:

The symbol “ C? “ marks clickable DOI hyperlinks.
For cites R packages, see http://www.cran.org.

## Supplemental Figure

**Supplemental Figure S1.**
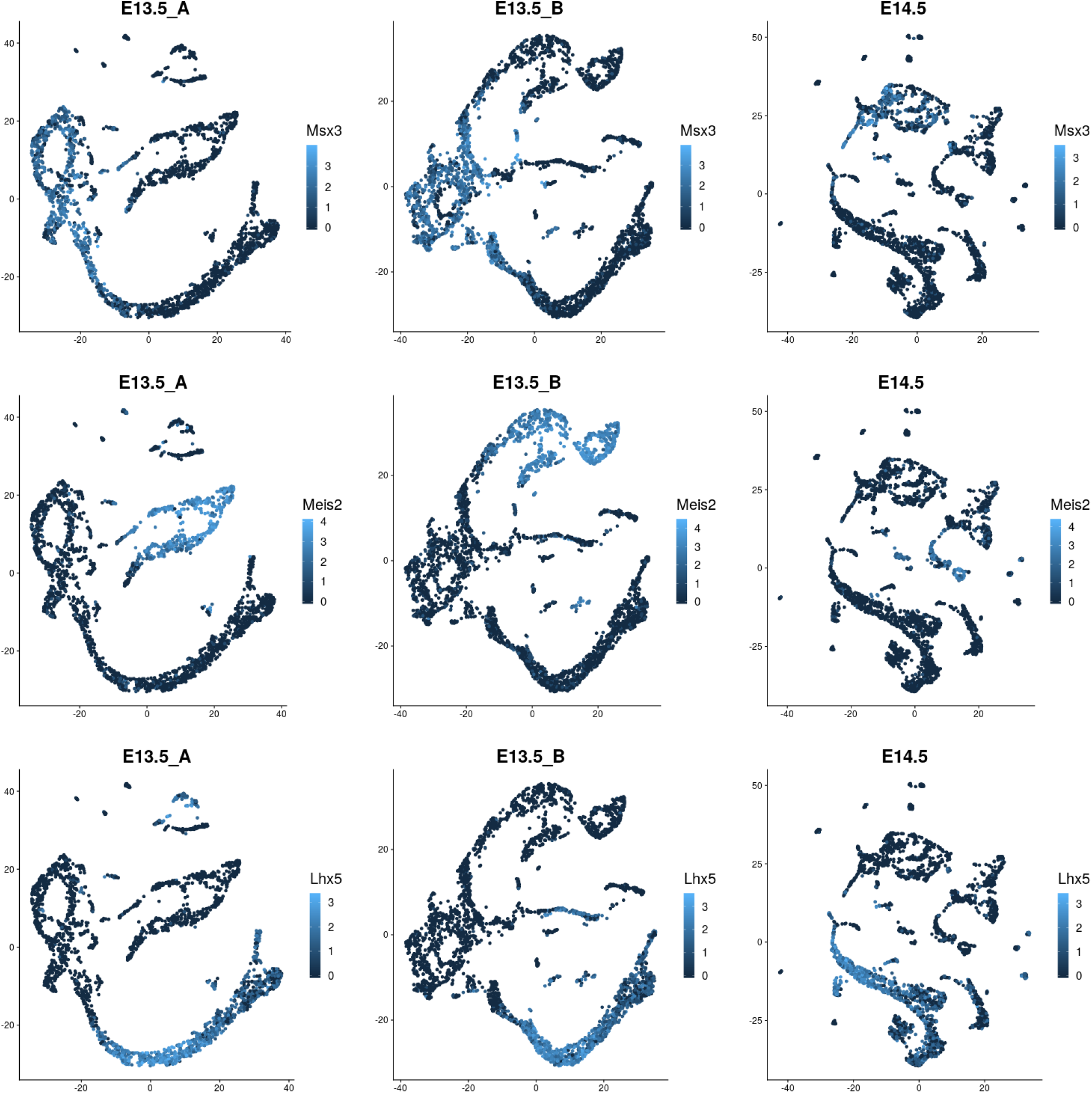
Markers used to identify lineages in developing murine cerebellum. Expression of three marker genes was used to identify GABAergic (Lhx5, bottom row) and glutamatergic (Meis2, middle row) lineages as well as early progenitor cells (Msx3, top row).

## References

10x Genomics (2019). What is Cell Ranger? https://support.10xgenomics.com/single-cell-gene-expression/software/pipelines/latest/what-is-cell-ranger

Angerer P, et al. (2016). destiny: diffusion maps for large-scale single-cell data in R. Bioinformatics, 32:1241.

Aupetit M (2007). Visualizing distortions and recovering topology in continuous projection techniques. Neurocomputing, 70:1304.

Baglama J and Reichel L (2005). Augmented implicitly restarted Lanczos bidiagonalization methods. SIAM Journal on Scientific Computing, 27:19.

Baglama J, Reichel L, and Lewis BW (2019). irlba: Fast Truncated Singular Value Decomposition and Principal Components Analysis for Large Dense and Sparse Matrices. R package version 2.3.3

Becht E, et al. (2019). Dimensionality reduction for visualizing single-cell data using UMAP. Nature Biotechnology, 37:38.

Becker RA and Cleveland WS (1987). Brushing scatterplots. Technometrics, 29:127.

Bostock M, Ogievetsky V, and Heer J (2011). *D*3 *data-driven documents*. IEEE Transactions on Visualization and Computer Graphics, 17:2301.

Brawand D, et al. (2011). The evolution of gene expression levels in mammalian organs. Nature, 478:343.

Buettner F, et al. (2015). Computational analysis of cell-to-cell heterogeneity in single-cell RNA-sequencing data reveals hidden subpopulations of cells. Nature Biotechnology, 33:155.

Butler A, et al. (2018). Integrating single-cell transcriptomic data across different conditions, technologies, and species. Nature Biotechnology, 36:411.

Carter RA, et al. (2018). A single-cell transcriptional atlas of the developing murine cerebellum. Current Biology, 28:2910.

Cayton L (2005). Algorithms for manifold learning. Technical Report CS2008-0923, University of California at San Diego. http://www.lcayton.com/resexam.pdf

Cheng J, et al. (2019). httpuv: HTTP and WebSocket Server Library. R package version 1.5.0

Coifman RR and Lafon S (2006). Diffusion maps. Applied and Computational Harmonic Analysis, 21:5.

Dietrich S, et al. (2018). Drug-perturbation-based stratification of blood cancer. Journal of Clinical Investigation, 128:427.

Fette I and Melnikov A (2011). The WebSocket protocol. RFC 6455, Internet Engineering Task Force. https://tools.ietf.org/html/rfc6455

Green DA (2011). A colour scheme for the display of astronomical intensity images. Bulletin of the Astromical Society of India, 39:289.

Haghverdi L, et al. (2018). Batch effects in single-cell RNA-sequencing data are corrected by matching mutual nearest neighbors. Nature Biotechnology, 36:421.

Kaski S, et al. (2003). Trustworthiness and metrics in visualizing similarity of gene expression. BMC Bioinformatics, 4:48.

Kohonen T (1982). Self-organized formation of topologically correct feature maps. Biological Cybernetics, 43:59.

Krijthe JH (2015). Rtsne: T-Distributed Stochastic Neighbor Embedding using Barnes-Hut Implementation. R package version 0.15

Kruskal JB (1964). Multidimensional scaling by optimizing goodness of fit to a nonmetric hypothesis. Psychometrika, 29:1.

Lespinats S and Aupetit M (2011). CheckViz: Sanity check and topological clues for linear and non-linear mappings. Computer Graphics Forum, 30:113.

Mao Q, et al. (2015). Dimensionality reduction via graph structure learning. In Proceedings of the 21th ACM SIGKDD International Conference on Knowledge Discovery and Data Mining, KDD ‘15, pp. 765–774. ACM, New York, NY, USA.

McInnes L, Healy J, and Melville J (2018). UMAP: Uniform manifold approximation and projection for dimension reduction. arXiv:1802.03426 [cs, stat]

Melville J (2019). uwot: The Uniform Manifold Approximation and Projection (UMAP) Method for Dimensionality Reduction. R package version 0.0.0.9010

Moon KR, et al. (2018). Manifold learning-based methods for analyzing single-cell rna-sequencing data. Current Opinion in Systems Biology, 7:36.

Nguyen LH and Holmes S (2019). Ten quick tips for effective dimensionality reduction. PLOS Computational Biology, 15:1.

Pedersen TL (2019). ggforce: Accelerating ggplot2. R package version 0.2.0

Phillips JM and Venkatasubramanian S (2011). A gentle introduction to the kernel distance. arXiv:1103.1625

Qiu X, et al. (2017). Reversed graph embedding resolves complex single-cell trajectories. Nature Methods, 14:979.

R Core Team (2019). R: A Language and Environment for Statistical Computing. R Foundation for Statistical Computing, Vienna, Austria

Ringnér M (2008). What is principal component analysis? Nature Biotechnology, 26:303.

Satija Lab (2018). Using Seurat with multi-modal data. https://satijalab.org/seurat/multimodalvignette.html

Seifert C, Sabol V, and Kienreich W (2010). Stress Maps: Analysing Local Phenomena in Dimensionality Reduction Based Visualisations. In J Kohlhammer and D Keim (Editors), EuroVAST 2010: International Symposium on Visual Analytics Science and Technology. The Eurographics Association.

Stoeckius M, et al. (2017). Simultaneous epitope and transcriptome measurement in single cells. Nature Methods, 14:865.

Trapnell C, et al. (2014). The dynamics and regulators of cell fate decisions are revealed by pseudotemporal ordering of single cells. Nature Biotechnology, 32:381.

Tung PY, et al. (2017). Batch effects and the effective design of single-cell gene expression studies. Scientific Reports, 7:39921.

van der Maaten L (2014). Accelerating t-SNE using tree-based algorithms. Journal of Machine Learning Research, 15:3221.

van der Maaten L and Hinton G (2008). Visualizing high-dimensional data using t-SNE. Journal of Machine Learning Research, 9:2579.

Wang B, et al. (2017). Visualization and analysis of single-cell RNA-seq data by kernel-based similarity learning. Nature Methods, 14:414.

Wattenberg M, Viégas F, and Johnson I (2016). How to use t-SNE effectively. Distill. http://doi.org/10.23915/distill.00002.

Yang L and Jin R (2006). Distance metric learning: A comprehensive survey. Technical report, Michigan State University. http://www.cs.cmu.edu/~liuy/framesurveyv2.pdf

