## Supplemental HTML page for "Exploring dimension-reduced embeddings with Sleepwalk": index.html

Sleepwalk supplement

### *Paper Supplement*

#### Svetlana Ovchinnikova and Simon Anders

This page contains interactive "live" versions of all the main figures in the Sleepwalk paper as well as few supplementary figures.

Sleepwalk displays a 2D embedding,i.e., a 2D representation of higher-dimensional data points,
and whenever the user hovers with the mouse over a data point, all points are coloured according to their distance to the focus point
under the mouse cursor. By moving the mouse over the plot, the user can explore how the faithfulness of the embedding varies between
regions of the plot. Buttons are provided to change the range of the colour scale.

For more information and a more comprehensive introduction to Sleepwalk, please refer to the paper or to the Sleepwalk web page

*Please select an example from the side bar*
