## Supplemental HTML page for "Exploring dimension-reduced embeddings with Sleepwalk": d3.min.js

```
// https://d3js.org v5.9.2 Copyright 2019 Mike Bostock
!function(t,n){"object"==typeof exports&&"undefined"!=typeof module?n(exports):"function"==typeof define&&define.amd?define(["exports"],n):n(t.d3=t.d3||{})}(this,function(t){"use strict";function n(t,n){return t<n?-1:t>n?1:t>=n?0:NaN}function e(t){var e;return 1===t.length&&(e=t,t=function(t,r){return n(e(t),r)}),{left:function(n,e,r,i){for(null==r&&(r=0),null==i&&(i=n.length);r<i;){var o=r+i>>>1;t(n[o],e)<0?r=o+1:i=o}return r},right:function(n,e,r,i){for(null==r&&(r=0),null==i&&(i=n.length);r<i;){var o=r+i>>>1;t(n[o],e)>0?i=o:r=o+1}return r}}}var r=e(n),i=r.right,o=r.left;function a(t,n){return[t,n]}function u(t){return null===t?NaN:+t}function c(t,n){var e,r,i=t.length,o=0,a=-1,c=0,f=0;if(null==n)for(;++a<i;)isNaN(e=u(t[a]))||(f+=(r=e-c)*(e-(c+=r/++o)));else for(;++a<i;)isNaN(e=u(n(t[a],a,t)))||(f+=(r=e-c)*(e-(c+=r/++o)));if(o>1)return f/(o-1)}function f(t,n){var e=c(t,n);return e?Math.sqrt(e):e}function s(t,n){var e,r,i,o=t.length,a=-1;if(null==n){for(;++a<o;)if(null!=(e=t[a])&&e>=e)for(r=i=e;++a<o;)null!=(e=t[a])&&(r>e&&(r=e),i<e&&(i=e))}else for(;++a<o;)if(null!=(e=n(t[a],a,t))&&e>=e)for(r=i=e;++a<o;)null!=(e=n(t[a],a,t))&&(r>e&&(r=e),i<e&&(i=e));return[r,i]}var l=Array.prototype,h=l.slice,d=l.map;function p(t){return function(){return t}}function v(t){return t}function g(t,n,e){t=+t,n=+n,e=(i=arguments.length)<2?(n=t,t=0,1):i<3?1:+e;for(var r=-1,i=0|Math.max(0,Math.ceil((n-t)/e)),o=new Array(i);++r<i;)o[r]=t+r*e;return o}var y=Math.sqrt(50),_=Math.sqrt(10),b=Math.sqrt(2);function m(t,n,e){var r,i,o,a,u=-1;if(e=+e,(t=+t)===(n=+n)&&e>0)return[t];if((r=n<t)&&(i=t,t=n,n=i),0===(a=x(t,n,e))||!isFinite(a))return[];if(a>0)for(t=Math.ceil(t/a),n=Math.floor(n/a),o=new Array(i=Math.ceil(n-t+1));++u<i;)o[u]=(t+u)*a;else for(t=Math.floor(t*a),n=Math.ceil(n*a),o=new Array(i=Math.ceil(t-n+1));++u<i;)o[u]=(t-u)/a;return r&&o.reverse(),o}function x(t,n,e){var r=(n-t)/Math.max(0,e),i=Math.floor(Math.log(r)/Math.LN10),o=r/Math.pow(10,i);return i>=0?(o>=y?10:o>=_?5:o>=b?2:1)*Math.pow(10,i):-Math.pow(10,-i)/(o>=y?10:o>=_?5:o>=b?2:1)}function w(t,n,e){var r=Math.abs(n-t)/Math.max(0,e),i=Math.pow(10,Math.floor(Math.log(r)/Math.LN10)),o=r/i;return o>=y?i*=10:o>=_?i*=5:o>=b&&(i*=2),n<t?-i:i}function M(t){return Math.ceil(Math.log(t.length)/Math.LN2)+1}function N(t,n,e){if(null==e&&(e=u),r=t.length){if((n=+n)<=0||r<2)return+e(t[0],0,t);if(n>=1)return+e(t[r-1],r-1,t);var r,i=(r-1)*n,o=Math.floor(i),a=+e(t[o],o,t);return a+(+e(t[o+1],o+1,t)-a)*(i-o)}}function A(t,n){var e,r,i=t.length,o=-1;if(null==n){for(;++o<i;)if(null!=(e=t[o])&&e>=e)for(r=e;++o<i;)null!=(e=t[o])&&e>r&&(r=e)}else for(;++o<i;)if(null!=(e=n(t[o],o,t))&&e>=e)for(r=e;++o<i;)null!=(e=n(t[o],o,t))&&e>r&&(r=e);return r}function T(t){for(var n,e,r,i=t.length,o=-1,a=0;++o<i;)a+=t[o].length;for(e=new Array(a);--i>=0;)for(n=(r=t[i]).length;--n>=0;)e[--a]=r[n];return e}function S(t,n){var e,r,i=t.length,o=-1;if(null==n){for(;++o<i;)if(null!=(e=t[o])&&e>=e)for(r=e;++o<i;)null!=(e=t[o])&&r>e&&(r=e)}else for(;++o<i;)if(null!=(e=n(t[o],o,t))&&e>=e)for(r=e;++o<i;)null!=(e=n(t[o],o,t))&&r>e&&(r=e);return r}function k(t){if(!(i=t.length))return[];for(var n=-1,e=S(t,E),r=new Array(e);++n<e;)for(var i,o=-1,a=r[n]=new Array(i);++o<i;)a[o]=t[o][n];return r}function E(t){return t.length}var C=Array.prototype.slice;function P(t){return t}var z=1,R=2,D=3,q=4,L=1e-6;function U(t){return"translate("+(t+.5)+",0)"}function O(t){return"translate(0,"+(t+.5)+")"}function B(){return!this.__axis}function Y(t,n){var e=[],r=null,i=null,o=6,a=6,u=3,c=t===z||t===q?-1:1,f=t===q||t===R?"x":"y",s=t===z||t===D?U:O;function l(l){var h=null==r?n.ticks?n.ticks.apply(n,e):n.domain():r,d=null==i?n.tickFormat?n.tickFormat.apply(n,e):P:i,p=Math.max(o,0)+u,v=n.range(),g=+v[0]+.5,y=+v[v.length-1]+.5,_=(n.bandwidth?function(t){var n=Math.max(0,t.bandwidth()-1)/2;return t.round()&&(n=Math.round(n)),function(e){return+t(e)+n}}:function(t){return function(n){return+t(n)}})(n.copy()),b=l.selection?l.selection():l,m=b.selectAll(".domain").data([null]),x=b.selectAll(".tick").data(h,n).order(),w=x.exit(),M=x.enter().append("g").attr("class","tick"),N=x.select("line"),A=x.select("text");m=m.merge(m.enter().insert("path",".tick").attr("class","domain").attr("stroke","currentColor")),x=x.merge(M),N=N.merge(M.append("line").attr("stroke","currentColor").attr(f+"2",c*o)),A=A.merge(M.append("text").attr("fill","currentColor").attr(f,c*p).attr("dy",t===z?"0em":t===D?"0.71em":"0.32em")),l!==b&&(m=m.transition(l),x=x.transition(l),N=N.transition(l),A=A.transition(l),w=w.transition(l).attr("opacity",L).attr("transform",function(t){return isFinite(t=_(t))?s(t):this.getAttribute("transform")}),M.attr("opacity",L).attr("transform",function(t){var n=this.parentNode.__axis;return s(n&&isFinite(n=n(t))?n:_(t))})),w.remove(),m.attr("d",t===q||t==R?a?"M"+c*a+","+g+"H0.5V"+y+"H"+c*a:"M0.5,"+g+"V"+y:a?"M"+g+","+c*a+"V0.5H"+y+"V"+c*a:"M"+g+",0.5H"+y),x.attr("opacity",1).attr("transform",function(t){return s(_(t))}),N.attr(f+"2",c*o),A.attr(f,c*p).text(d),b.filter(B).attr("fill","none").attr("font-size",10).attr("font-family","sans-serif").attr("text-anchor",t===R?"start":t===q?"end":"middle"),b.each(function(){this.__axis=_})}return l.scale=function(t){return arguments.length?(n=t,l):n},l.ticks=function(){return e=C.call(arguments),l},l.tickArguments=function(t){return arguments.length?(e=null==t?[]:C.call(t),l):e.slice()},l.tickValues=function(t){return arguments.length?(r=null==t?null:C.call(t),l):r&&r.slice()},l.tickFormat=function(t){return arguments.length?(i=t,l):i},l.tickSize=function(t){return arguments.length?(o=a=+t,l):o},l.tickSizeInner=function(t){return arguments.length?(o=+t,l):o},l.tickSizeOuter=function(t){return arguments.length?(a=+t,l):a},l.tickPadding=function(t){return arguments.length?(u=+t,l):u},l}var F={value:function(){}};function I(){for(var t,n=0,e=arguments.length,r={};n<e;++n){if(!(t=arguments[n]+"")||t in r)throw new Error("illegal type: "+t);r[t]=[]}return new j(r)}function j(t){this._=t}function H(t,n){for(var e,r=0,i=t.length;r<i;++r)if((e=t[r]).name===n)return e.value}function X(t,n,e){for(var r=0,i=t.length;r<i;++r)if(t[r].name===n){t[r]=F,t=t.slice(0,r).concat(t.slice(r+1));break}return null!=e&&t.push({name:n,value:e}),t}j.prototype=I.prototype={constructor:j,on:function(t,n){var e,r,i=this._,o=(r=i,(t+"").trim().split(/^|\s+/).map(function(t){var n="",e=t.indexOf(".");if(e>=0&&(n=t.slice(e+1),t=t.slice(0,e)),t&&!r.hasOwnProperty(t))throw new Error("unknown type: "+t);return{type:t,name:n}})),a=-1,u=o.length;if(!(arguments.length<2)){if(null!=n&&"function"!=typeof n)throw new Error("invalid callback: "+n);for(;++a<u;)if(e=(t=o[a]).type)i[e]=X(i[e],t.name,n);else if(null==n)for(e in i)i[e]=X(i[e],t.name,null);return this}for(;++a<u;)if((e=(t=o[a]).type)&&(e=H(i[e],t.name)))return e},copy:function(){var t={},n=this._;for(var e in n)t[e]=n[e].slice();return new j(t)},call:function(t,n){if((e=arguments.length-2)>0)for(var e,r,i=new Array(e),o=0;o<e;++o)i[o]=arguments[o+2];if(!this._.hasOwnProperty(t))throw new Error("unknown type: "+t);for(o=0,e=(r=this._[t]).length;o<e;++o)r[o].value.apply(n,i)},apply:function(t,n,e){if(!this._.hasOwnProperty(t))throw new Error("unknown type: "+t);for(var r=this._[t],i=0,o=r.length;i<o;++i)r[i].value.apply(n,e)}};var G="http://www.w3.org/1999/xhtml",V={svg:"http://www.w3.org/2000/svg",xhtml:G,xlink:"http://www.w3.org/1999/xlink",xml:"http://www.w3.org/XML/1998/namespace",xmlns:"http://www.w3.org/2000/xmlns/"};function $(t){var n=t+="",e=n.indexOf(":");return e>=0&&"xmlns"!==(n=t.slice(0,e))&&(t=t.slice(e+1)),V.hasOwnProperty(n)?{space:V[n],local:t}:t}function W(t){var n=$(t);return(n.local?function(t){return function(){return this.ownerDocument.createElementNS(t.space,t.local)}}:function(t){return function(){var n=this.ownerDocument,e=this.namespaceURI;return e===G&&n.documentElement.namespaceURI===G?n.createElement(t):n.createElementNS(e,t)}})(n)}function Z(){}function Q(t){return null==t?Z:function(){return this.querySelector(t)}}function J(){return[]}function K(t){return null==t?J:function(){return this.querySelectorAll(t)}}function tt(t){return function(){return this.matches(t)}}function nt(t){return new Array(t.length)}function et(t,n){this.ownerDocument=t.ownerDocument,this.namespaceURI=t.namespaceURI,this._next=null,this._parent=t,this.__data__=n}et.prototype={constructor:et,appendChild:function(t){return this._parent.insertBefore(t,this._next)},insertBefore:function(t,n){return this._parent.insertBefore(t,n)},querySelector:function(t){return this._parent.querySelector(t)},querySelectorAll:function(t){return this._parent.querySelectorAll(t)}};var rt="$";function it(t,n,e,r,i,o){for(var a,u=0,c=n.length,f=o.length;u<f;++u)(a=n[u])?(a.__data__=o[u],r[u]=a):e[u]=new et(t,o[u]);for(;u<c;++u)(a=n[u])&&(i[u]=a)}function ot(t,n,e,r,i,o,a){var u,c,f,s={},l=n.length,h=o.length,d=new Array(l);for(u=0;u<l;++u)(c=n[u])&&(d[u]=f=rt+a.call(c,c.__data__,u,n),f in s?i[u]=c:s[f]=c);for(u=0;u<h;++u)(c=s[f=rt+a.call(t,o[u],u,o)])?(r[u]=c,c.__data__=o[u],s[f]=null):e[u]=new et(t,o[u]);for(u=0;u<l;++u)(c=n[u])&&s[d[u]]===c&&(i[u]=c)}function at(t,n){return t<n?-1:t>n?1:t>=n?0:NaN}function ut(t){return t.ownerDocument&&t.ownerDocument.defaultView||t.document&&t||t.defaultView}function ct(t,n){return t.style.getPropertyValue(n)||ut(t).getComputedStyle(t,null).getPropertyValue(n)}function ft(t){return t.trim().split(/^|\s+/)}function st(t){return t.classList||new lt(t)}function lt(t){this._node=t,this._names=ft(t.getAttribute("class")||"")}function ht(t,n){for(var e=st(t),r=-1,i=n.length;++r<i;)e.add(n[r])}function dt(t,n){for(var e=st(t),r=-1,i=n.length;++r<i;)e.remove(n[r])}function pt(){this.textContent=""}function vt(){this.innerHTML=""}function gt(){this.nextSibling&&this.parentNode.appendChild(this)}function yt(){this.previousSibling&&this.parentNode.insertBefore(this,this.parentNode.firstChild)}function _t(){return null}function bt(){var t=this.parentNode;t&&t.removeChild(this)}function mt(){return this.parentNode.insertBefore(this.cloneNode(!1),this.nextSibling)}function xt(){return this.parentNode.insertBefore(this.cloneNode(!0),this.nextSibling)}lt.prototype={add:function(t){this._names.indexOf(t)<0&&(this._names.push(t),this._node.setAttribute("class",this._names.join(" ")))},remove:function(t){var n=this._names.indexOf(t);n>=0&&(this._names.splice(n,1),this._node.setAttribute("class",this._names.join(" ")))},contains:function(t){return this._names.indexOf(t)>=0}};var wt={};(t.event=null,"undefined"!=typeof document)&&("onmouseenter"in document.documentElement||(wt={mouseenter:"mouseover",mouseleave:"mouseout"}));function Mt(t,n,e){return t=Nt(t,n,e),function(n){var e=n.relatedTarget;e&&(e===this||8&e.compareDocumentPosition(this))||t.call(this,n)}}function Nt(n,e,r){return function(i){var o=t.event;t.event=i;try{n.call(this,this.__data__,e,r)}finally{t.event=o}}}function At(t){return function(){var n=this.__on;if(n){for(var e,r=0,i=-1,o=n.length;r<o;++r)e=n[r],t.type&&e.type!==t.type||e.name!==t.name?n[++i]=e:this.removeEventListener(e.type,e.listener,e.capture);++i?n.length=i:delete this.__on}}}function Tt(t,n,e){var r=wt.hasOwnProperty(t.type)?Mt:Nt;return function(i,o,a){var u,c=this.__on,f=r(n,o,a);if(c)for(var s=0,l=c.length;s<l;++s)if((u=c[s]).type===t.type&&u.name===t.name)return this.removeEventListener(u.type,u.listener,u.capture),this.addEventListener(u.type,u.listener=f,u.capture=e),void(u.value=n);this.addEventListener(t.type,f,e),u={type:t.type,name:t.name,value:n,listener:f,capture:e},c?c.push(u):this.__on=[u]}}function St(n,e,r,i){var o=t.event;n.sourceEvent=t.event,t.event=n;try{return e.apply(r,i)}finally{t.event=o}}function kt(t,n,e){var r=ut(t),i=r.CustomEvent;"function"==typeof i?i=new i(n,e):(i=r.document.createEvent("Event"),e?(i.initEvent(n,e.bubbles,e.cancelable),i.detail=e.detail):i.initEvent(n,!1,!1)),t.dispatchEvent(i)}var Et=[null];function Ct(t,n){this._groups=t,this._parents=n}function Pt(){return new Ct([[document.documentElement]],Et)}function zt(t){return"string"==typeof t?new Ct([[document.querySelector(t)]],[document.documentElement]):new Ct([[t]],Et)}Ct.prototype=Pt.prototype={constructor:Ct,select:function(t){"function"!=typeof t&&(t=Q(t));for(var n=this._groups,e=n.length,r=new Array(e),i=0;i<e;++i)for(var o,a,u=n[i],c=u.length,f=r[i]=new Array(c),s=0;s<c;++s)(o=u[s])&&(a=t.call(o,o.__data__,s,u))&&("__data__"in o&&(a.__data__=o.__data__),f[s]=a);return new Ct(r,this._parents)},selectAll:function(t){"function"!=typeof t&&(t=K(t));for(var n=this._groups,e=n.length,r=[],i=[],o=0;o<e;++o)for(var a,u=n[o],c=u.length,f=0;f<c;++f)(a=u[f])&&(r.push(t.call(a,a.__data__,f,u)),i.push(a));return new Ct(r,i)},filter:function(t){"function"!=typeof t&&(t=tt(t));for(var n=this._groups,e=n.length,r=new Array(e),i=0;i<e;++i)for(var o,a=n[i],u=a.length,c=r[i]=[],f=0;f<u;++f)(o=a[f])&&t.call(o,o.__data__,f,a)&&c.push(o);return new Ct(r,this._parents)},data:function(t,n){if(!t)return p=new Array(this.size()),s=-1,this.each(function(t){p[++s]=t}),p;var e,r=n?ot:it,i=this._parents,o=this._groups;"function"!=typeof t&&(e=t,t=function(){return e});for(var a=o.length,u=new Array(a),c=new Array(a),f=new Array(a),s=0;s<a;++s){var l=i[s],h=o[s],d=h.length,p=t.call(l,l&&l.__data__,s,i),v=p.length,g=c[s]=new Array(v),y=u[s]=new Array(v);r(l,h,g,y,f[s]=new Array(d),p,n);for(var _,b,m=0,x=0;m<v;++m)if(_=g[m]){for(m>=x&&(x=m+1);!(b=y[x])&&++x<v;);_._next=b||null}}return(u=new Ct(u,i))._enter=c,u._exit=f,u},enter:function(){return new Ct(this._enter||this._groups.map(nt),this._parents)},exit:function(){return new Ct(this._exit||this._groups.map(nt),this._parents)},join:function(t,n,e){var r=this.enter(),i=this,o=this.exit();return r="function"==typeof t?t(r):r.append(t+""),null!=n&&(i=n(i)),null==e?o.remove():e(o),r&&i?r.merge(i).order():i},merge:function(t){for(var n=this._groups,e=t._groups,r=n.length,i=e.length,o=Math.min(r,i),a=new Array(r),u=0;u<o;++u)for(var c,f=n[u],s=e[u],l=f.length,h=a[u]=new Array(l),d=0;d<l;++d)(c=f[d]||s[d])&&(h[d]=c);for(;u<r;++u)a[u]=n[u];return new Ct(a,this._parents)},order:function(){for(var t=this._groups,n=-1,e=t.length;++n<e;)for(var r,i=t[n],o=i.length-1,a=i[o];--o>=0;)(r=i[o])&&(a&&4^r.compareDocumentPosition(a)&&a.parentNode.insertBefore(r,a),a=r);return this},sort:function(t){function n(n,e){return n&&e?t(n.__data__,e.__data__):!n-!e}t||(t=at);for(var e=this._groups,r=e.length,i=new Array(r),o=0;o<r;++o){for(var a,u=e[o],c=u.length,f=i[o]=new Array(c),s=0;s<c;++s)(a=u[s])&&(f[s]=a);f.sort(n)}return new Ct(i,this._parents).order()},call:function(){var t=arguments[0];return arguments[0]=this,t.apply(null,arguments),this},nodes:function(){var t=new Array(this.size()),n=-1;return this.each(function(){t[++n]=this}),t},node:function(){for(var t=this._groups,n=0,e=t.length;n<e;++n)for(var r=t[n],i=0,o=r.length;i<o;++i){var a=r[i];if(a)return a}return null},size:function(){var t=0;return this.each(function(){++t}),t},empty:function(){return!this.node()},each:function(t){for(var n=this._groups,e=0,r=n.length;e<r;++e)for(var i,o=n[e],a=0,u=o.length;a<u;++a)(i=o[a])&&t.call(i,i.__data__,a,o);return this},attr:function(t,n){var e=$(t);if(arguments.length<2){var r=this.node();return e.local?r.getAttributeNS(e.space,e.local):r.getAttribute(e)}return this.each((null==n?e.local?function(t){return function(){this.removeAttributeNS(t.space,t.local)}}:function(t){return function(){this.removeAttribute(t)}}:"function"==typeof n?e.local?function(t,n){return function(){var e=n.apply(this,arguments);null==e?this.removeAttributeNS(t.space,t.local):this.setAttributeNS(t.space,t.local,e)}}:function(t,n){return function(){var e=n.apply(this,arguments);null==e?this.removeAttribute(t):this.setAttribute(t,e)}}:e.local?function(t,n){return function(){this.setAttributeNS(t.space,t.local,n)}}:function(t,n){return function(){this.setAttribute(t,n)}})(e,n))},style:function(t,n,e){return arguments.length>1?this.each((null==n?function(t){return function(){this.style.removeProperty(t)}}:"function"==typeof n?function(t,n,e){return function(){var r=n.apply(this,arguments);null==r?this.style.removeProperty(t):this.style.setProperty(t,r,e)}}:function(t,n,e){return function(){this.style.setProperty(t,n,e)}})(t,n,null==e?"":e)):ct(this.node(),t)},property:function(t,n){return arguments.length>1?this.each((null==n?function(t){return function(){delete this[t]}}:"function"==typeof n?function(t,n){return function(){var e=n.apply(this,arguments);null==e?delete this[t]:this[t]=e}}:function(t,n){return function(){this[t]=n}})(t,n)):this.node()[t]},classed:function(t,n){var e=ft(t+"");if(arguments.length<2){for(var r=st(this.node()),i=-1,o=e.length;++i<o;)if(!r.contains(e[i]))return!1;return!0}return this.each(("function"==typeof n?function(t,n){return function(){(n.apply(this,arguments)?ht:dt)(this,t)}}:n?function(t){return function(){ht(this,t)}}:function(t){return function(){dt(this,t)}})(e,n))},text:function(t){return arguments.length?this.each(null==t?pt:("function"==typeof t?function(t){return function(){var n=t.apply(this,arguments);this.textContent=null==n?"":n}}:function(t){return function(){this.textContent=t}})(t)):this.node().textContent},html:function(t){return arguments.length?this.each(null==t?vt:("function"==typeof t?function(t){return function(){var n=t.apply(this,arguments);this.innerHTML=null==n?"":n}}:function(t){return function(){this.innerHTML=t}})(t)):this.node().innerHTML},raise:function(){return this.each(gt)},lower:function(){return this.each(yt)},append:function(t){var n="function"==typeof t?t:W(t);return this.select(function(){return this.appendChild(n.apply(this,arguments))})},insert:function(t,n){var e="function"==typeof t?t:W(t),r=null==n?_t:"function"==typeof n?n:Q(n);return this.select(function(){return this.insertBefore(e.apply(this,arguments),r.apply(this,arguments)||null)})},remove:function(){return this.each(bt)},clone:function(t){return this.select(t?xt:mt)},datum:function(t){return arguments.length?this.property("__data__",t):this.node().__data__},on:function(t,n,e){var r,i,o=function(t){return t.trim().split(/^|\s+/).map(function(t){var n="",e=t.indexOf(".");return e>=0&&(n=t.slice(e+1),t=t.slice(0,e)),{type:t,name:n}})}(t+""),a=o.length;if(!(arguments.length<2)){for(u=n?Tt:At,null==e&&(e=!1),r=0;r<a;++r)this.each(u(o[r],n,e));return this}var u=this.node().__on;if(u)for(var c,f=0,s=u.length;f<s;++f)for(r=0,c=u[f];r<a;++r)if((i=o[r]).type===c.type&&i.name===c.name)return c.value},dispatch:function(t,n){return this.each(("function"==typeof n?function(t,n){return function(){return kt(this,t,n.apply(this,arguments))}}:function(t,n){return function(){return kt(this,t,n)}})(t,n))}};var Rt=0;function Dt(){return new qt}function qt(){this._="@"+(++Rt).toString(36)}function Lt(){for(var n,e=t.event;n=e.sourceEvent;)e=n;return e}function Ut(t,n){var e=t.ownerSVGElement||t;if(e.createSVGPoint){var r=e.createSVGPoint();return r.x=n.clientX,r.y=n.clientY,[(r=r.matrixTransform(t.getScreenCTM().inverse())).x,r.y]}var i=t.getBoundingClientRect();return[n.clientX-i.left-t.clientLeft,n.clientY-i.top-t.clientTop]}function Ot(t){var n=Lt();return n.changedTouches&&(n=n.changedTouches[0]),Ut(t,n)}function Bt(t,n,e){arguments.length<3&&(e=n,n=Lt().changedTouches);for(var r,i=0,o=n?n.length:0;i<o;++i)if((r=n[i]).identifier===e)return Ut(t,r);return null}function Yt(){t.event.stopImmediatePropagation()}function Ft(){t.event.preventDefault(),t.event.stopImmediatePropagation()}function It(t){var n=t.document.documentElement,e=zt(t).on("dragstart.drag",Ft,!0);"onselectstart"in n?e.on("selectstart.drag",Ft,!0):(n.__noselect=n.style.MozUserSelect,n.style.MozUserSelect="none")}function jt(t,n){var e=t.document.documentElement,r=zt(t).on("dragstart.drag",null);n&&(r.on("click.drag",Ft,!0),setTimeout(function(){r.on("click.drag",null)},0)),"onselectstart"in e?r.on("selectstart.drag",null):(e.style.MozUserSelect=e.__noselect,delete e.__noselect)}function Ht(t){return function(){return t}}function Xt(t,n,e,r,i,o,a,u,c,f){this.target=t,this.type=n,this.subject=e,this.identifier=r,this.active=i,this.x=o,this.y=a,this.dx=u,this.dy=c,this._=f}function Gt(){return!t.event.button}function Vt(){return this.parentNode}function $t(n){return null==n?{x:t.event.x,y:t.event.y}:n}function Wt(){return"ontouchstart"in this}function Zt(t,n,e){t.prototype=n.prototype=e,e.constructor=t}function Qt(t,n){var e=Object.create(t.prototype);for(var r in n)e[r]=n[r];return e}function Jt(){}qt.prototype=Dt.prototype={constructor:qt,get:function(t){for(var n=this._;!(n in t);)if(!(t=t.parentNode))return;return t[n]},set:function(t,n){return t[this._]=n},remove:function(t){return this._ in t&&delete t[this._]},toString:function(){return this._}},Xt.prototype.on=function(){var t=this._.on.apply(this._,arguments);return t===this._?this:t};var Kt="\\s*([+-]?\\d+)\\s*",tn="\\s*([+-]?\\d*\\.?\\d+(?:[eE][+-]?\\d+)?)\\s*",nn="\\s*([+-]?\\d*\\.?\\d+(?:[eE][+-]?\\d+)?)%\\s*",en=/^#([0-9a-f]{3})$/,rn=/^#([0-9a-f]{6})$/,on=new RegExp("^rgb\\("+[Kt,Kt,Kt]+"\\)$"),an=new RegExp("^rgb\\("+[nn,nn,nn]+"\\)$"),un=new RegExp("^rgba\\("+[Kt,Kt,Kt,tn]+"\\)$"),cn=new RegExp("^rgba\\("+[nn,nn,nn,tn]+"\\)$"),fn=new RegExp("^hsl\\("+[tn,nn,nn]+"\\)$"),sn=new RegExp("^hsla\\("+[tn,nn,nn,tn]+"\\)$"),ln={aliceblue:15792383,antiquewhite:16444375,aqua:65535,aquamarine:8388564,azure:15794175,beige:16119260,bisque:16770244,black:0,blanchedalmond:16772045,blue:255,blueviolet:9055202,brown:10824234,burlywood:14596231,cadetblue:6266528,chartreuse:8388352,chocolate:13789470,coral:16744272,cornflowerblue:6591981,cornsilk:16775388,crimson:14423100,cyan:65535,darkblue:139,darkcyan:35723,darkgoldenrod:12092939,darkgray:11119017,darkgreen:25600,darkgrey:11119017,darkkhaki:12433259,darkmagenta:9109643,darkolivegreen:5597999,darkorange:16747520,darkorchid:10040012,darkred:9109504,darksalmon:15308410,darkseagreen:9419919,darkslateblue:4734347,darkslategray:3100495,darkslategrey:3100495,darkturquoise:52945,darkviolet:9699539,deeppink:16716947,deepskyblue:49151,dimgray:6908265,dimgrey:6908265,dodgerblue:2003199,firebrick:11674146,floralwhite:16775920,forestgreen:2263842,fuchsia:16711935,gainsboro:14474460,ghostwhite:16316671,gold:16766720,goldenrod:14329120,gray:8421504,green:32768,greenyellow:11403055,grey:8421504,honeydew:15794160,hotpink:16738740,indianred:13458524,indigo:4915330,ivory:16777200,khaki:15787660,lavender:15132410,lavenderblush:16773365,lawngreen:8190976,lemonchiffon:16775885,lightblue:11393254,lightcoral:15761536,lightcyan:14745599,lightgoldenrodyellow:16448210,lightgray:13882323,lightgreen:9498256,lightgrey:13882323,lightpink:16758465,lightsalmon:16752762,lightseagreen:2142890,lightskyblue:8900346,lightslategray:7833753,lightslategrey:7833753,lightsteelblue:11584734,lightyellow:16777184,lime:65280,limegreen:3329330,linen:16445670,magenta:16711935,maroon:8388608,mediumaquamarine:6737322,mediumblue:205,mediumorchid:12211667,mediumpurple:9662683,mediumseagreen:3978097,mediumslateblue:8087790,mediumspringgreen:64154,mediumturquoise:4772300,mediumvioletred:13047173,midnightblue:1644912,mintcream:16121850,mistyrose:16770273,moccasin:16770229,navajowhite:16768685,navy:128,oldlace:16643558,olive:8421376,olivedrab:7048739,orange:16753920,orangered:16729344,orchid:14315734,palegoldenrod:15657130,palegreen:10025880,paleturquoise:11529966,palevioletred:14381203,papayawhip:16773077,peachpuff:16767673,peru:13468991,pink:16761035,plum:14524637,powderblue:11591910,purple:8388736,rebeccapurple:6697881,red:16711680,rosybrown:12357519,royalblue:4286945,saddlebrown:9127187,salmon:16416882,sandybrown:16032864,seagreen:3050327,seashell:16774638,sienna:10506797,silver:12632256,skyblue:8900331,slateblue:6970061,slategray:7372944,slategrey:7372944,snow:16775930,springgreen:65407,steelblue:4620980,tan:13808780,teal:32896,thistle:14204888,tomato:16737095,turquoise:4251856,violet:15631086,wheat:16113331,white:16777215,whitesmoke:16119285,yellow:16776960,yellowgreen:10145074};function hn(t){var n;return t=(t+"").trim().toLowerCase(),(n=en.exec(t))?new yn((n=parseInt(n[1],16))>>8&15|n>>4&240,n>>4&15|240&n,(15&n)<<4|15&n,1):(n=rn.exec(t))?dn(parseInt(n[1],16)):(n=on.exec(t))?new yn(n[1],n[2],n[3],1):(n=an.exec(t))?new yn(255*n[1]/100,255*n[2]/100,255*n[3]/100,1):(n=un.exec(t))?pn(n[1],n[2],n[3],n[4]):(n=cn.exec(t))?pn(255*n[1]/100,255*n[2]/100,255*n[3]/100,n[4]):(n=fn.exec(t))?bn(n[1],n[2]/100,n[3]/100,1):(n=sn.exec(t))?bn(n[1],n[2]/100,n[3]/100,n[4]):ln.hasOwnProperty(t)?dn(ln[t]):"transparent"===t?new yn(NaN,NaN,NaN,0):null}function dn(t){return new yn(t>>16&255,t>>8&255,255&t,1)}function pn(t,n,e,r){return r<=0&&(t=n=e=NaN),new yn(t,n,e,r)}function vn(t){return t instanceof Jt||(t=hn(t)),t?new yn((t=t.rgb()).r,t.g,t.b,t.opacity):new yn}function gn(t,n,e,r){return 1===arguments.length?vn(t):new yn(t,n,e,null==r?1:r)}function yn(t,n,e,r){this.r=+t,this.g=+n,this.b=+e,this.opacity=+r}function _n(t){return((t=Math.max(0,Math.min(255,Math.round(t)||0)))<16?"0":"")+t.toString(16)}function bn(t,n,e,r){return r<=0?t=n=e=NaN:e<=0||e>=1?t=n=NaN:n<=0&&(t=NaN),new xn(t,n,e,r)}function mn(t,n,e,r){return 1===arguments.length?function(t){if(t instanceof xn)return new xn(t.h,t.s,t.l,t.opacity);if(t instanceof Jt||(t=hn(t)),!t)return new xn;if(t instanceof xn)return t;var n=(t=t.rgb()).r/255,e=t.g/255,r=t.b/255,i=Math.min(n,e,r),o=Math.max(n,e,r),a=NaN,u=o-i,c=(o+i)/2;return u?(a=n===o?(e-r)/u+6*(e<r):e===o?(r-n)/u+2:(n-e)/u+4,u/=c<.5?o+i:2-o-i,a*=60):u=c>0&&c<1?0:a,new xn(a,u,c,t.opacity)}(t):new xn(t,n,e,null==r?1:r)}function xn(t,n,e,r){this.h=+t,this.s=+n,this.l=+e,this.opacity=+r}function wn(t,n,e){return 255*(t<60?n+(e-n)*t/60:t<180?e:t<240?n+(e-n)*(240-t)/60:n)}Zt(Jt,hn,{displayable:function(){return this.rgb().displayable()},hex:function(){return this.rgb().hex()},toString:function(){return this.rgb()+""}}),Zt(yn,gn,Qt(Jt,{brighter:function(t){return t=null==t?1/.7:Math.pow(1/.7,t),new yn(this.r*t,this.g*t,this.b*t,this.opacity)},darker:function(t){return t=null==t?.7:Math.pow(.7,t),new yn(this.r*t,this.g*t,this.b*t,this.opacity)},rgb:function(){return this},displayable:function(){return 0<=this.r&&this.r<=255&&0<=this.g&&this.g<=255&&0<=this.b&&this.b<=255&&0<=this.opacity&&this.opacity<=1},hex:function(){return"#"+_n(this.r)+_n(this.g)+_n(this.b)},toString:function(){var t=this.opacity;return(1===(t=isNaN(t)?1:Math.max(0,Math.min(1,t)))?"rgb(":"rgba(")+Math.max(0,Math.min(255,Math.round(this.r)||0))+", "+Math.max(0,Math.min(255,Math.round(this.g)||0))+", "+Math.max(0,Math.min(255,Math.round(this.b)||0))+(1===t?")":", "+t+")")}})),Zt(xn,mn,Qt(Jt,{brighter:function(t){return t=null==t?1/.7:Math.pow(1/.7,t),new xn(this.h,this.s,this.l*t,this.opacity)},darker:function(t){return t=null==t?.7:Math.pow(.7,t),new xn(this.h,this.s,this.l*t,this.opacity)},rgb:function(){var t=this.h%360+360*(this.h<0),n=isNaN(t)||isNaN(this.s)?0:this.s,e=this.l,r=e+(e<.5?e:1-e)*n,i=2*e-r;return new yn(wn(t>=240?t-240:t+120,i,r),wn(t,i,r),wn(t<120?t+240:t-120,i,r),this.opacity)},displayable:function(){return(0<=this.s&&this.s<=1||isNaN(this.s))&&0<=this.l&&this.l<=1&&0<=this.opacity&&this.opacity<=1}}));var Mn=Math.PI/180,Nn=180/Math.PI,An=.96422,Tn=1,Sn=.82521,kn=4/29,En=6/29,Cn=3*En*En,Pn=En*En*En;function zn(t){if(t instanceof Dn)return new Dn(t.l,t.a,t.b,t.opacity);if(t instanceof Fn){if(isNaN(t.h))return new Dn(t.l,0,0,t.opacity);var n=t.h*Mn;return new Dn(t.l,Math.cos(n)*t.c,Math.sin(n)*t.c,t.opacity)}t instanceof yn||(t=vn(t));var e,r,i=On(t.r),o=On(t.g),a=On(t.b),u=qn((.2225045*i+.7168786*o+.0606169*a)/Tn);return i===o&&o===a?e=r=u:(e=qn((.4360747*i+.3850649*o+.1430804*a)/An),r=qn((.0139322*i+.0971045*o+.7141733*a)/Sn)),new Dn(116*u-16,500*(e-u),200*(u-r),t.opacity)}function Rn(t,n,e,r){return 1===arguments.length?zn(t):new Dn(t,n,e,null==r?1:r)}function Dn(t,n,e,r){this.l=+t,this.a=+n,this.b=+e,this.opacity=+r}function qn(t){return t>Pn?Math.pow(t,1/3):t/Cn+kn}function Ln(t){return t>En?t*t*t:Cn*(t-kn)}function Un(t){return 255*(t<=.0031308?12.92*t:1.055*Math.pow(t,1/2.4)-.055)}function On(t){return(t/=255)<=.04045?t/12.92:Math.pow((t+.055)/1.055,2.4)}function Bn(t){if(t instanceof Fn)return new Fn(t.h,t.c,t.l,t.opacity);if(t instanceof Dn||(t=zn(t)),0===t.a&&0===t.b)return new Fn(NaN,0,t.l,t.opacity);var n=Math.atan2(t.b,t.a)*Nn;return new Fn(n<0?n+360:n,Math.sqrt(t.a*t.a+t.b*t.b),t.l,t.opacity)}function Yn(t,n,e,r){return 1===arguments.length?Bn(t):new Fn(t,n,e,null==r?1:r)}function Fn(t,n,e,r){this.h=+t,this.c=+n,this.l=+e,this.opacity=+r}Zt(Dn,Rn,Qt(Jt,{brighter:function(t){return new Dn(this.l+18*(null==t?1:t),this.a,this.b,this.opacity)},darker:function(t){return new Dn(this.l-18*(null==t?1:t),this.a,this.b,this.opacity)},rgb:function(){var t=(this.l+16)/116,n=isNaN(this.a)?t:t+this.a/500,e=isNaN(this.b)?t:t-this.b/200;return new yn(Un(3.1338561*(n=An*Ln(n))-1.6168667*(t=Tn*Ln(t))-.4906146*(e=Sn*Ln(e))),Un(-.9787684*n+1.9161415*t+.033454*e),Un(.0719453*n-.2289914*t+1.4052427*e),this.opacity)}})),Zt(Fn,Yn,Qt(Jt,{brighter:function(t){return new Fn(this.h,this.c,this.l+18*(null==t?1:t),this.opacity)},darker:function(t){return new Fn(this.h,this.c,this.l-18*(null==t?1:t),this.opacity)},rgb:function(){return zn(this).rgb()}}));var In=-.14861,jn=1.78277,Hn=-.29227,Xn=-.90649,Gn=1.97294,Vn=Gn*Xn,$n=Gn*jn,Wn=jn*Hn-Xn*In;function Zn(t,n,e,r){return 1===arguments.length?function(t){if(t instanceof Qn)return new Qn(t.h,t.s,t.l,t.opacity);t instanceof yn||(t=vn(t));var n=t.r/255,e=t.g/255,r=t.b/255,i=(Wn*r+Vn*n-$n*e)/(Wn+Vn-$n),o=r-i,a=(Gn*(e-i)-Hn*o)/Xn,u=Math.sqrt(a*a+o*o)/(Gn*i*(1-i)),c=u?Math.atan2(a,o)*Nn-120:NaN;return new Qn(c<0?c+360:c,u,i,t.opacity)}(t):new Qn(t,n,e,null==r?1:r)}function Qn(t,n,e,r){this.h=+t,this.s=+n,this.l=+e,this.opacity=+r}function Jn(t,n,e,r,i){var o=t*t,a=o*t;return((1-3*t+3*o-a)*n+(4-6*o+3*a)*e+(1+3*t+3*o-3*a)*r+a*i)/6}function Kn(t){var n=t.length-1;return function(e){var r=e<=0?e=0:e>=1?(e=1,n-1):Math.floor(e*n),i=t[r],o=t[r+1],a=r>0?t[r-1]:2*i-o,u=r<n-1?t[r+2]:2*o-i;return Jn((e-r/n)*n,a,i,o,u)}}function te(t){var n=t.length;return function(e){var r=Math.floor(((e%=1)<0?++e:e)*n),i=t[(r+n-1)%n],o=t[r%n],a=t[(r+1)%n],u=t[(r+2)%n];return Jn((e-r/n)*n,i,o,a,u)}}function ne(t){return function(){return t}}function ee(t,n){return function(e){return t+e*n}}function re(t,n){var e=n-t;return e?ee(t,e>180||e<-180?e-360*Math.round(e/360):e):ne(isNaN(t)?n:t)}function ie(t){return 1==(t=+t)?oe:function(n,e){return e-n?function(t,n,e){return t=Math.pow(t,e),n=Math.pow(n,e)-t,e=1/e,function(r){return Math.pow(t+r*n,e)}}(n,e,t):ne(isNaN(n)?e:n)}}function oe(t,n){var e=n-t;return e?ee(t,e):ne(isNaN(t)?n:t)}Zt(Qn,Zn,Qt(Jt,{brighter:function(t){return t=null==t?1/.7:Math.pow(1/.7,t),new Qn(this.h,this.s,this.l*t,this.opacity)},darker:function(t){return t=null==t?.7:Math.pow(.7,t),new Qn(this.h,this.s,this.l*t,this.opacity)},rgb:function(){var t=isNaN(this.h)?0:(this.h+120)*Mn,n=+this.l,e=isNaN(this.s)?0:this.s*n*(1-n),r=Math.cos(t),i=Math.sin(t);return new yn(255*(n+e*(In*r+jn*i)),255*(n+e*(Hn*r+Xn*i)),255*(n+e*(Gn*r)),this.opacity)}}));var ae=function t(n){var e=ie(n);function r(t,n){var r=e((t=gn(t)).r,(n=gn(n)).r),i=e(t.g,n.g),o=e(t.b,n.b),a=oe(t.opacity,n.opacity);return function(n){return t.r=r(n),t.g=i(n),t.b=o(n),t.opacity=a(n),t+""}}return r.gamma=t,r}(1);function ue(t){return function(n){var e,r,i=n.length,o=new Array(i),a=new Array(i),u=new Array(i);for(e=0;e<i;++e)r=gn(n[e]),o[e]=r.r||0,a[e]=r.g||0,u[e]=r.b||0;return o=t(o),a=t(a),u=t(u),r.opacity=1,function(t){return r.r=o(t),r.g=a(t),r.b=u(t),r+""}}}var ce=ue(Kn),fe=ue(te);function se(t,n){var e,r=n?n.length:0,i=t?Math.min(r,t.length):0,o=new Array(i),a=new Array(r);for(e=0;e<i;++e)o[e]=ye(t[e],n[e]);for(;e<r;++e)a[e]=n[e];return function(t){for(e=0;e<i;++e)a[e]=o[e](t);return a}}function le(t,n){var e=new Date;return n-=t=+t,function(r){return e.setTime(t+n*r),e}}function he(t,n){return n-=t=+t,function(e){return t+n*e}}function de(t,n){var e,r={},i={};for(e in null!==t&&"object"==typeof t||(t={}),null!==n&&"object"==typeof n||(n={}),n)e in t?r[e]=ye(t[e],n[e]):i[e]=n[e];return function(t){for(e in r)i[e]=r[e](t);return i}}var pe=/[-+]?(?:\d+\.?\d*|\.?\d+)(?:[eE][-+]?\d+)?/g,ve=new RegExp(pe.source,"g");function ge(t,n){var e,r,i,o=pe.lastIndex=ve.lastIndex=0,a=-1,u=[],c=[];for(t+="",n+="";(e=pe.exec(t))&&(r=ve.exec(n));)(i=r.index)>o&&(i=n.slice(o,i),u[a]?u[a]+=i:u[++a]=i),(e=e[0])===(r=r[0])?u[a]?u[a]+=r:u[++a]=r:(u[++a]=null,c.push({i:a,x:he(e,r)})),o=ve.lastIndex;return o<n.length&&(i=n.slice(o),u[a]?u[a]+=i:u[++a]=i),u.length<2?c[0]?function(t){return function(n){return t(n)+""}}(c[0].x):function(t){return function(){return t}}(n):(n=c.length,function(t){for(var e,r=0;r<n;++r)u[(e=c[r]).i]=e.x(t);return u.join("")})}function ye(t,n){var e,r=typeof n;return null==n||"boolean"===r?ne(n):("number"===r?he:"string"===r?(e=hn(n))?(n=e,ae):ge:n instanceof hn?ae:n instanceof Date?le:Array.isArray(n)?se:"function"!=typeof n.valueOf&&"function"!=typeof n.toString||isNaN(n)?de:he)(t,n)}function _e(t,n){return n-=t=+t,function(e){return Math.round(t+n*e)}}var be,me,xe,we,Me=180/Math.PI,Ne={translateX:0,translateY:0,rotate:0,skewX:0,scaleX:1,scaleY:1};function Ae(t,n,e,r,i,o){var a,u,c;return(a=Math.sqrt(t*t+n*n))&&(t/=a,n/=a),(c=t*e+n*r)&&(e-=t*c,r-=n*c),(u=Math.sqrt(e*e+r*r))&&(e/=u,r/=u,c/=u),t*r<n*e&&(t=-t,n=-n,c=-c,a=-a),{translateX:i,translateY:o,rotate:Math.atan2(n,t)*Me,skewX:Math.atan(c)*Me,scaleX:a,scaleY:u}}function Te(t,n,e,r){function i(t){return t.length?t.pop()+" ":""}return function(o,a){var u=[],c=[];return o=t(o),a=t(a),function(t,r,i,o,a,u){if(t!==i||r!==o){var c=a.push("translate(",null,n,null,e);u.push({i:c-4,x:he(t,i)},{i:c-2,x:he(r,o)})}else(i||o)&&a.push("translate("+i+n+o+e)}(o.translateX,o.translateY,a.translateX,a.translateY,u,c),function(t,n,e,o){t!==n?(t-n>180?n+=360:n-t>180&&(t+=360),o.push({i:e.push(i(e)+"rotate(",null,r)-2,x:he(t,n)})):n&&e.push(i(e)+"rotate("+n+r)}(o.rotate,a.rotate,u,c),function(t,n,e,o){t!==n?o.push({i:e.push(i(e)+"skewX(",null,r)-2,x:he(t,n)}):n&&e.push(i(e)+"skewX("+n+r)}(o.skewX,a.skewX,u,c),function(t,n,e,r,o,a){if(t!==e||n!==r){var u=o.push(i(o)+"scale(",null,",",null,")");a.push({i:u-4,x:he(t,e)},{i:u-2,x:he(n,r)})}else 1===e&&1===r||o.push(i(o)+"scale("+e+","+r+")")}(o.scaleX,o.scaleY,a.scaleX,a.scaleY,u,c),o=a=null,function(t){for(var n,e=-1,r=c.length;++e<r;)u[(n=c[e]).i]=n.x(t);return u.join("")}}}var Se=Te(function(t){return"none"===t?Ne:(be||(be=document.createElement("DIV"),me=document.documentElement,xe=document.defaultView),be.style.transform=t,t=xe.getComputedStyle(me.appendChild(be),null).getPropertyValue("transform"),me.removeChild(be),Ae(+(t=t.slice(7,-1).split(","))[0],+t[1],+t[2],+t[3],+t[4],+t[5]))},"px, ","px)","deg)"),ke=Te(function(t){return null==t?Ne:(we||(we=document.createElementNS("http://www.w3.org/2000/svg","g")),we.setAttribute("transform",t),(t=we.transform.baseVal.consolidate())?Ae((t=t.matrix).a,t.b,t.c,t.d,t.e,t.f):Ne)},", ",")",")"),Ee=Math.SQRT2,Ce=2,Pe=4,ze=1e-12;function Re(t){return((t=Math.exp(t))+1/t)/2}function De(t,n){var e,r,i=t[0],o=t[1],a=t[2],u=n[0],c=n[1],f=n[2],s=u-i,l=c-o,h=s*s+l*l;if(h<ze)r=Math.log(f/a)/Ee,e=function(t){return[i+t*s,o+t*l,a*Math.exp(Ee*t*r)]};else{var d=Math.sqrt(h),p=(f*f-a*a+Pe*h)/(2*a*Ce*d),v=(f*f-a*a-Pe*h)/(2*f*Ce*d),g=Math.log(Math.sqrt(p*p+1)-p),y=Math.log(Math.sqrt(v*v+1)-v);r=(y-g)/Ee,e=function(t){var n,e=t*r,u=Re(g),c=a/(Ce*d)*(u*(n=Ee*e+g,((n=Math.exp(2*n))-1)/(n+1))-function(t){return((t=Math.exp(t))-1/t)/2}(g));return[i+c*s,o+c*l,a*u/Re(Ee*e+g)]}}return e.duration=1e3*r,e}function qe(t){return function(n,e){var r=t((n=mn(n)).h,(e=mn(e)).h),i=oe(n.s,e.s),o=oe(n.l,e.l),a=oe(n.opacity,e.opacity);return function(t){return n.h=r(t),n.s=i(t),n.l=o(t),n.opacity=a(t),n+""}}}var Le=qe(re),Ue=qe(oe);function Oe(t){return function(n,e){var r=t((n=Yn(n)).h,(e=Yn(e)).h),i=oe(n.c,e.c),o=oe(n.l,e.l),a=oe(n.opacity,e.opacity);return function(t){return n.h=r(t),n.c=i(t),n.l=o(t),n.opacity=a(t),n+""}}}var Be=Oe(re),Ye=Oe(oe);function Fe(t){return function n(e){function r(n,r){var i=t((n=Zn(n)).h,(r=Zn(r)).h),o=oe(n.s,r.s),a=oe(n.l,r.l),u=oe(n.opacity,r.opacity);return function(t){return n.h=i(t),n.s=o(t),n.l=a(Math.pow(t,e)),n.opacity=u(t),n+""}}return e=+e,r.gamma=n,r}(1)}var Ie=Fe(re),je=Fe(oe);var He,Xe,Ge=0,Ve=0,$e=0,We=1e3,Ze=0,Qe=0,Je=0,Ke="object"==typeof performance&&performance.now?performance:Date,tr="object"==typeof window&&window.requestAnimationFrame?window.requestAnimationFrame.bind(window):function(t){setTimeout(t,17)};function nr(){return Qe||(tr(er),Qe=Ke.now()+Je)}function er(){Qe=0}function rr(){this._call=this._time=this._next=null}function ir(t,n,e){var r=new rr;return r.restart(t,n,e),r}function or(){nr(),++Ge;for(var t,n=He;n;)(t=Qe-n._time)>=0&&n._call.call(null,t),n=n._next;--Ge}function ar(){Qe=(Ze=Ke.now())+Je,Ge=Ve=0;try{or()}finally{Ge=0,function(){var t,n,e=He,r=1/0;for(;e;)e._call?(r>e._time&&(r=e._time),t=e,e=e._next):(n=e._next,e._next=null,e=t?t._next=n:He=n);Xe=t,cr(r)}(),Qe=0}}function ur(){var t=Ke.now(),n=t-Ze;n>We&&(Je-=n,Ze=t)}function cr(t){Ge||(Ve&&(Ve=clearTimeout(Ve)),t-Qe>24?(t<1/0&&(Ve=setTimeout(ar,t-Ke.now()-Je)),$e&&($e=clearInterval($e))):($e||(Ze=Ke.now(),$e=setInterval(ur,We)),Ge=1,tr(ar)))}function fr(t,n,e){var r=new rr;return n=null==n?0:+n,r.restart(function(e){r.stop(),t(e+n)},n,e),r}rr.prototype=ir.prototype={constructor:rr,restart:function(t,n,e){if("function"!=typeof t)throw new TypeError("callback is not a function");e=(null==e?nr():+e)+(null==n?0:+n),this._next||Xe===this||(Xe?Xe._next=this:He=this,Xe=this),this._call=t,this._time=e,cr()},stop:function(){this._call&&(this._call=null,this._time=1/0,cr())}};var sr=I("start","end","cancel","interrupt"),lr=[],hr=0,dr=1,pr=2,vr=3,gr=4,yr=5,_r=6;function br(t,n,e,r,i,o){var a=t.__transition;if(a){if(e in a)return}else t.__transition={};!function(t,n,e){var r,i=t.__transition;function o(c){var f,s,l,h;if(e.state!==dr)return u();for(f in i)if((h=i[f]).name===e.name){if(h.state===vr)return fr(o);h.state===gr?(h.state=_r,h.timer.stop(),h.on.call("interrupt",t,t.__data__,h.index,h.group),delete i[f]):+f<n&&(h.state=_r,h.timer.stop(),h.on.call("cancel",t,t.__data__,h.index,h.group),delete i[f])}if(fr(function(){e.state===vr&&(e.state=gr,e.timer.restart(a,e.delay,e.time),a(c))}),e.state=pr,e.on.call("start",t,t.__data__,e.index,e.group),e.state===pr){for(e.state=vr,r=new Array(l=e.tween.length),f=0,s=-1;f<l;++f)(h=e.tween[f].value.call(t,t.__data__,e.index,e.group))&&(r[++s]=h);r.length=s+1}}function a(n){for(var i=n<e.duration?e.ease.call(null,n/e.duration):(e.timer.restart(u),e.state=yr,1),o=-1,a=r.length;++o<a;)r[o].call(t,i);e.state===yr&&(e.on.call("end",t,t.__data__,e.index,e.group),u())}function u(){for(var r in e.state=_r,e.timer.stop(),delete i[n],i)return;delete t.__transition}i[n]=e,e.timer=ir(function(t){e.state=dr,e.timer.restart(o,e.delay,e.time),e.delay<=t&&o(t-e.delay)},0,e.time)}(t,e,{name:n,index:r,group:i,on:sr,tween:lr,time:o.time,delay:o.delay,duration:o.duration,ease:o.ease,timer:null,state:hr})}function mr(t,n){var e=wr(t,n);if(e.state>hr)throw new Error("too late; already scheduled");return e}function xr(t,n){var e=wr(t,n);if(e.state>vr)throw new Error("too late; already running");return e}function wr(t,n){var e=t.__transition;if(!e||!(e=e[n]))throw new Error("transition not found");return e}function Mr(t,n){var e,r,i,o=t.__transition,a=!0;if(o){for(i in n=null==n?null:n+"",o)(e=o[i]).name===n?(r=e.state>pr&&e.state<yr,e.state=_r,e.timer.stop(),e.on.call(r?"interrupt":"cancel",t,t.__data__,e.index,e.group),delete o[i]):a=!1;a&&delete t.__transition}}function Nr(t,n,e){var r=t._id;return t.each(function(){var t=xr(this,r);(t.value||(t.value={}))[n]=e.apply(this,arguments)}),function(t){return wr(t,r).value[n]}}function Ar(t,n){var e;return("number"==typeof n?he:n instanceof hn?ae:(e=hn(n))?(n=e,ae):ge)(t,n)}var Tr=Pt.prototype.constructor;function Sr(t){return function(){this.style.removeProperty(t)}}var kr=0;function Er(t,n,e,r){this._groups=t,this._parents=n,this._name=e,this._id=r}function Cr(t){return Pt().transition(t)}function Pr(){return++kr}var zr=Pt.prototype;function Rr(t){return((t*=2)<=1?t*t:--t*(2-t)+1)/2}function Dr(t){return((t*=2)<=1?t*t*t:(t-=2)*t*t+2)/2}Er.prototype=Cr.prototype={constructor:Er,select:function(t){var n=this._name,e=this._id;"function"!=typeof t&&(t=Q(t));for(var r=this._groups,i=r.length,o=new Array(i),a=0;a<i;++a)for(var u,c,f=r[a],s=f.length,l=o[a]=new Array(s),h=0;h<s;++h)(u=f[h])&&(c=t.call(u,u.__data__,h,f))&&("__data__"in u&&(c.__data__=u.__data__),l[h]=c,br(l[h],n,e,h,l,wr(u,e)));return new Er(o,this._parents,n,e)},selectAll:function(t){var n=this._name,e=this._id;"function"!=typeof t&&(t=K(t));for(var r=this._groups,i=r.length,o=[],a=[],u=0;u<i;++u)for(var c,f=r[u],s=f.length,l=0;l<s;++l)if(c=f[l]){for(var h,d=t.call(c,c.__data__,l,f),p=wr(c,e),v=0,g=d.length;v<g;++v)(h=d[v])&&br(h,n,e,v,d,p);o.push(d),a.push(c)}return new Er(o,a,n,e)},filter:function(t){"function"!=typeof t&&(t=tt(t));for(var n=this._groups,e=n.length,r=new Array(e),i=0;i<e;++i)for(var o,a=n[i],u=a.length,c=r[i]=[],f=0;f<u;++f)(o=a[f])&&t.call(o,o.__data__,f,a)&&c.push(o);return new Er(r,this._parents,this._name,this._id)},merge:function(t){if(t._id!==this._id)throw new Error;for(var n=this._groups,e=t._groups,r=n.length,i=e.length,o=Math.min(r,i),a=new Array(r),u=0;u<o;++u)for(var c,f=n[u],s=e[u],l=f.length,h=a[u]=new Array(l),d=0;d<l;++d)(c=f[d]||s[d])&&(h[d]=c);for(;u<r;++u)a[u]=n[u];return new Er(a,this._parents,this._name,this._id)},selection:function(){return new Tr(this._groups,this._parents)},transition:function(){for(var t=this._name,n=this._id,e=Pr(),r=this._groups,i=r.length,o=0;o<i;++o)for(var a,u=r[o],c=u.length,f=0;f<c;++f)if(a=u[f]){var s=wr(a,n);br(a,t,e,f,u,{time:s.time+s.delay+s.duration,delay:0,duration:s.duration,ease:s.ease})}return new Er(r,this._parents,t,e)},call:zr.call,nodes:zr.nodes,node:zr.node,size:zr.size,empty:zr.empty,each:zr.each,on:function(t,n){var e=this._id;return arguments.length<2?wr(this.node(),e).on.on(t):this.each(function(t,n,e){var r,i,o=function(t){return(t+"").trim().split(/^|\s+/).every(function(t){var n=t.indexOf(".");return n>=0&&(t=t.slice(0,n)),!t||"start"===t})}(n)?mr:xr;return function(){var a=o(this,t),u=a.on;u!==r&&(i=(r=u).copy()).on(n,e),a.on=i}}(e,t,n))},attr:function(t,n){var e=$(t),r="transform"===e?ke:Ar;return this.attrTween(t,"function"==typeof n?(e.local?function(t,n,e){var r,i,o;return function(){var a,u,c=e(this);if(null!=c)return(a=this.getAttributeNS(t.space,t.local))===(u=c+"")?null:a===r&&u===i?o:(i=u,o=n(r=a,c));this.removeAttributeNS(t.space,t.local)}}:function(t,n,e){var r,i,o;return function(){var a,u,c=e(this);if(null!=c)return(a=this.getAttribute(t))===(u=c+"")?null:a===r&&u===i?o:(i=u,o=n(r=a,c));this.removeAttribute(t)}})(e,r,Nr(this,"attr."+t,n)):null==n?(e.local?function(t){return function(){this.removeAttributeNS(t.space,t.local)}}:function(t){return function(){this.removeAttribute(t)}})(e):(e.local?function(t,n,e){var r,i,o=e+"";return function(){var a=this.getAttributeNS(t.space,t.local);return a===o?null:a===r?i:i=n(r=a,e)}}:function(t,n,e){var r,i,o=e+"";return function(){var a=this.getAttribute(t);return a===o?null:a===r?i:i=n(r=a,e)}})(e,r,n))},attrTween:function(t,n){var e="attr."+t;if(arguments.length<2)return(e=this.tween(e))&&e._value;if(null==n)return this.tween(e,null);if("function"!=typeof n)throw new Error;var r=$(t);return this.tween(e,(r.local?function(t,n){var e,r;function i(){var i=n.apply(this,arguments);return i!==r&&(e=(r=i)&&function(t,n){return function(e){this.setAttributeNS(t.space,t.local,n(e))}}(t,i)),e}return i._value=n,i}:function(t,n){var e,r;function i(){var i=n.apply(this,arguments);return i!==r&&(e=(r=i)&&function(t,n){return function(e){this.setAttribute(t,n(e))}}(t,i)),e}return i._value=n,i})(r,n))},style:function(t,n,e){var r="transform"==(t+="")?Se:Ar;return null==n?this.styleTween(t,function(t,n){var e,r,i;return function(){var o=ct(this,t),a=(this.style.removeProperty(t),ct(this,t));return o===a?null:o===e&&a===r?i:i=n(e=o,r=a)}}(t,r)).on("end.style."+t,Sr(t)):"function"==typeof n?this.styleTween(t,function(t,n,e){var r,i,o;return function(){var a=ct(this,t),u=e(this),c=u+"";return null==u&&(this.style.removeProperty(t),c=u=ct(this,t)),a===c?null:a===r&&c===i?o:(i=c,o=n(r=a,u))}}(t,r,Nr(this,"style."+t,n))).each(function(t,n){var e,r,i,o,a="style."+n,u="end."+a;return function(){var c=xr(this,t),f=c.on,s=null==c.value[a]?o||(o=Sr(n)):void 0;f===e&&i===s||(r=(e=f).copy()).on(u,i=s),c.on=r}}(this._id,t)):this.styleTween(t,function(t,n,e){var r,i,o=e+"";return function(){var a=ct(this,t);return a===o?null:a===r?i:i=n(r=a,e)}}(t,r,n),e).on("end.style."+t,null)},styleTween:function(t,n,e){var r="style."+(t+="");if(arguments.length<2)return(r=this.tween(r))&&r._value;if(null==n)return this.tween(r,null);if("function"!=typeof n)throw new Error;return this.tween(r,function(t,n,e){var r,i;function o(){var o=n.apply(this,arguments);return o!==i&&(r=(i=o)&&function(t,n,e){return function(r){this.style.setProperty(t,n(r),e)}}(t,o,e)),r}return o._value=n,o}(t,n,null==e?"":e))},text:function(t){return this.tween("text","function"==typeof t?function(t){return function(){var n=t(this);this.textContent=null==n?"":n}}(Nr(this,"text",t)):function(t){return function(){this.textContent=t}}(null==t?"":t+""))},remove:function(){return this.on("end.remove",(t=this._id,function(){var n=this.parentNode;for(var e in this.__transition)if(+e!==t)return;n&&n.removeChild(this)}));var t},tween:function(t,n){var e=this._id;if(t+="",arguments.length<2){for(var r,i=wr(this.node(),e).tween,o=0,a=i.length;o<a;++o)if((r=i[o]).name===t)return r.value;return null}return this.each((null==n?function(t,n){var e,r;return function(){var i=xr(this,t),o=i.tween;if(o!==e)for(var a=0,u=(r=e=o).length;a<u;++a)if(r[a].name===n){(r=r.slice()).splice(a,1);break}i.tween=r}}:function(t,n,e){var r,i;if("function"!=typeof e)throw new Error;return function(){var o=xr(this,t),a=o.tween;if(a!==r){i=(r=a).slice();for(var u={name:n,value:e},c=0,f=i.length;c<f;++c)if(i[c].name===n){i[c]=u;break}c===f&&i.push(u)}o.tween=i}})(e,t,n))},delay:function(t){var n=this._id;return arguments.length?this.each(("function"==typeof t?function(t,n){return function(){mr(this,t).delay=+n.apply(this,arguments)}}:function(t,n){return n=+n,function(){mr(this,t).delay=n}})(n,t)):wr(this.node(),n).delay},duration:function(t){var n=this._id;return arguments.length?this.each(("function"==typeof t?function(t,n){return function(){xr(this,t).duration=+n.apply(this,arguments)}}:function(t,n){return n=+n,function(){xr(this,t).duration=n}})(n,t)):wr(this.node(),n).duration},ease:function(t){var n=this._id;return arguments.length?this.each(function(t,n){if("function"!=typeof n)throw new Error;return function(){xr(this,t).ease=n}}(n,t)):wr(this.node(),n).ease},end:function(){var t,n,e=this,r=e._id,i=e.size();return new Promise(function(o,a){var u={value:a},c={value:function(){0==--i&&o()}};e.each(function(){var e=xr(this,r),i=e.on;i!==t&&((n=(t=i).copy())._.cancel.push(u),n._.interrupt.push(u),n._.end.push(c)),e.on=n})})}};var qr=function t(n){function e(t){return Math.pow(t,n)}return n=+n,e.exponent=t,e}(3),Lr=function t(n){function e(t){return 1-Math.pow(1-t,n)}return n=+n,e.exponent=t,e}(3),Ur=function t(n){function e(t){return((t*=2)<=1?Math.pow(t,n):2-Math.pow(2-t,n))/2}return n=+n,e.exponent=t,e}(3),Or=Math.PI,Br=Or/2;function Yr(t){return(1-Math.cos(Or*t))/2}function Fr(t){return((t*=2)<=1?Math.pow(2,10*t-10):2-Math.pow(2,10-10*t))/2}function Ir(t){return((t*=2)<=1?1-Math.sqrt(1-t*t):Math.sqrt(1-(t-=2)*t)+1)/2}var jr=4/11,Hr=6/11,Xr=8/11,Gr=.75,Vr=9/11,$r=10/11,Wr=.9375,Zr=21/22,Qr=63/64,Jr=1/jr/jr;function Kr(t){return(t=+t)<jr?Jr*t*t:t<Xr?Jr*(t-=Hr)*t+Gr:t<$r?Jr*(t-=Vr)*t+Wr:Jr*(t-=Zr)*t+Qr}var ti=function t(n){function e(t){return t*t*((n+1)*t-n)}return n=+n,e.overshoot=t,e}(1.70158),ni=function t(n){function e(t){return--t*t*((n+1)*t+n)+1}return n=+n,e.overshoot=t,e}(1.70158),ei=function t(n){function e(t){return((t*=2)<1?t*t*((n+1)*t-n):(t-=2)*t*((n+1)*t+n)+2)/2}return n=+n,e.overshoot=t,e}(1.70158),ri=2*Math.PI,ii=function t(n,e){var r=Math.asin(1/(n=Math.max(1,n)))*(e/=ri);function i(t){return n*Math.pow(2,10*--t)*Math.sin((r-t)/e)}return i.amplitude=function(n){return t(n,e*ri)},i.period=function(e){return t(n,e)},i}(1,.3),oi=function t(n,e){var r=Math.asin(1/(n=Math.max(1,n)))*(e/=ri);function i(t){return 1-n*Math.pow(2,-10*(t=+t))*Math.sin((t+r)/e)}return i.amplitude=function(n){return t(n,e*ri)},i.period=function(e){return t(n,e)},i}(1,.3),ai=function t(n,e){var r=Math.asin(1/(n=Math.max(1,n)))*(e/=ri);function i(t){return((t=2*t-1)<0?n*Math.pow(2,10*t)*Math.sin((r-t)/e):2-n*Math.pow(2,-10*t)*Math.sin((r+t)/e))/2}return i.amplitude=function(n){return t(n,e*ri)},i.period=function(e){return t(n,e)},i}(1,.3),ui={time:null,delay:0,duration:250,ease:Dr};function ci(t,n){for(var e;!(e=t.__transition)||!(e=e[n]);)if(!(t=t.parentNode))return ui.time=nr(),ui;return e}Pt.prototype.interrupt=function(t){return this.each(function(){Mr(this,t)})},Pt.prototype.transition=function(t){var n,e;t instanceof Er?(n=t._id,t=t._name):(n=Pr(),(e=ui).time=nr(),t=null==t?null:t+"");for(var r=this._groups,i=r.length,o=0;o<i;++o)for(var a,u=r[o],c=u.length,f=0;f<c;++f)(a=u[f])&&br(a,t,n,f,u,e||ci(a,n));return new Er(r,this._parents,t,n)};var fi=[null];function si(t){return function(){return t}}function li(t,n,e){this.target=t,this.type=n,this.selection=e}function hi(){t.event.stopImmediatePropagation()}function di(){t.event.preventDefault(),t.event.stopImmediatePropagation()}var pi={name:"drag"},vi={name:"space"},gi={name:"handle"},yi={name:"center"},_i={name:"x",handles:["e","w"].map(Ti),input:function(t,n){return t&&[[t[0],n[0][1]],[t[1],n[1][1]]]},output:function(t){return t&&[t[0][0],t[1][0]]}},bi={name:"y",handles:["n","s"].map(Ti),input:function(t,n){return t&&[[n[0][0],t[0]],[n[1][0],t[1]]]},output:function(t){return t&&[t[0][1],t[1][1]]}},mi={name:"xy",handles:["n","e","s","w","nw","ne","se","sw"].map(Ti),input:function(t){return t},output:function(t){return t}},xi={overlay:"crosshair",selection:"move",n:"ns-resize",e:"ew-resize",s:"ns-resize",w:"ew-resize",nw:"nwse-resize",ne:"nesw-resize",se:"nwse-resize",sw:"nesw-resize"},wi={e:"w",w:"e",nw:"ne",ne:"nw",se:"sw",sw:"se"},Mi={n:"s",s:"n",nw:"sw",ne:"se",se:"ne",sw:"nw"},Ni={overlay:1,selection:1,n:null,e:1,s:null,w:-1,nw:-1,ne:1,se:1,sw:-1},Ai={overlay:1,selection:1,n:-1,e:null,s:1,w:null,nw:-1,ne:-1,se:1,sw:1};function Ti(t){return{type:t}}function Si(){return!t.event.button}function ki(){var t=this.ownerSVGElement||this;return[[0,0],[t.width.baseVal.value,t.height.baseVal.value]]}function Ei(t){for(;!t.__brush;)if(!(t=t.parentNode))return;return t.__brush}function Ci(t){return t[0][0]===t[1][0]||t[0][1]===t[1][1]}function Pi(n){var e,r=ki,i=Si,o=I(u,"start","brush","end"),a=6;function u(t){var e=t.property("__brush",h).selectAll(".overlay").data([Ti("overlay")]);e.enter().append("rect").attr("class","overlay").attr("pointer-events","all").attr("cursor",xi.overlay).merge(e).each(function(){var t=Ei(this).extent;zt(this).attr("x",t[0][0]).attr("y",t[0][1]).attr("width",t[1][0]-t[0][0]).attr("height",t[1][1]-t[0][1])}),t.selectAll(".selection").data([Ti("selection")]).enter().append("rect").attr("class","selection").attr("cursor",xi.selection).attr("fill","#777").attr("fill-opacity",.3).attr("stroke","#fff").attr("shape-rendering","crispEdges");var r=t.selectAll(".handle").data(n.handles,function(t){return t.type});r.exit().remove(),r.enter().append("rect").attr("class",function(t){return"handle handle--"+t.type}).attr("cursor",function(t){return xi[t.type]}),t.each(c).attr("fill","none").attr("pointer-events","all").style("-webkit-tap-highlight-color","rgba(0,0,0,0)").on("mousedown.brush touchstart.brush",l)}function c(){var t=zt(this),n=Ei(this).selection;n?(t.selectAll(".selection").style("display",null).attr("x",n[0][0]).attr("y",n[0][1]).attr("width",n[1][0]-n[0][0]).attr("height",n[1][1]-n[0][1]),t.selectAll(".handle").style("display",null).attr("x",function(t){return"e"===t.type[t.type.length-1]?n[1][0]-a/2:n[0][0]-a/2}).attr("y",function(t){return"s"===t.type[0]?n[1][1]-a/2:n[0][1]-a/2}).attr("width",function(t){return"n"===t.type||"s"===t.type?n[1][0]-n[0][0]+a:a}).attr("height",function(t){return"e"===t.type||"w"===t.type?n[1][1]-n[0][1]+a:a})):t.selectAll(".selection,.handle").style("display","none").attr("x",null).attr("y",null).attr("width",null).attr("height",null)}function f(t,n){return t.__brush.emitter||new s(t,n)}function s(t,n){this.that=t,this.args=n,this.state=t.__brush,this.active=0}function l(){if(t.event.touches){if(t.event.changedTouches.length<t.event.touches.length)return di()}else if(e)return;if(i.apply(this,arguments)){var r,o,a,u,s,l,h,d,p,v,g,y,_,b=this,m=t.event.target.__data__.type,x="selection"===(t.event.metaKey?m="overlay":m)?pi:t.event.altKey?yi:gi,w=n===bi?null:Ni[m],M=n===_i?null:Ai[m],N=Ei(b),A=N.extent,T=N.selection,S=A[0][0],k=A[0][1],E=A[1][0],C=A[1][1],P=w&&M&&t.event.shiftKey,z=Ot(b),R=z,D=f(b,arguments).beforestart();"overlay"===m?N.selection=T=[[r=n===bi?S:z[0],a=n===_i?k:z[1]],[s=n===bi?E:r,h=n===_i?C:a]]:(r=T[0][0],a=T[0][1],s=T[1][0],h=T[1][1]),o=r,u=a,l=s,d=h;var q=zt(b).attr("pointer-events","none"),L=q.selectAll(".overlay").attr("cursor",xi[m]);if(t.event.touches)q.on("touchmove.brush",O,!0).on("touchend.brush touchcancel.brush",Y,!0);else{var U=zt(t.event.view).on("keydown.brush",function(){switch(t.event.keyCode){case 16:P=w&&M;break;case 18:x===gi&&(w&&(s=l-p*w,r=o+p*w),M&&(h=d-v*M,a=u+v*M),x=yi,B());break;case 32:x!==gi&&x!==yi||(w<0?s=l-p:w>0&&(r=o-p),M<0?h=d-v:M>0&&(a=u-v),x=vi,L.attr("cursor",xi.selection),B());break;default:return}di()},!0).on("keyup.brush",function(){switch(t.event.keyCode){case 16:P&&(y=_=P=!1,B());break;case 18:x===yi&&(w<0?s=l:w>0&&(r=o),M<0?h=d:M>0&&(a=u),x=gi,B());break;case 32:x===vi&&(t.event.altKey?(w&&(s=l-p*w,r=o+p*w),M&&(h=d-v*M,a=u+v*M),x=yi):(w<0?s=l:w>0&&(r=o),M<0?h=d:M>0&&(a=u),x=gi),L.attr("cursor",xi[m]),B());break;default:return}di()},!0).on("mousemove.brush",O,!0).on("mouseup.brush",Y,!0);It(t.event.view)}hi(),Mr(b),c.call(b),D.start()}function O(){var t=Ot(b);!P||y||_||(Math.abs(t[0]-R[0])>Math.abs(t[1]-R[1])?_=!0:y=!0),R=t,g=!0,di(),B()}function B(){var t;switch(p=R[0]-z[0],v=R[1]-z[1],x){case vi:case pi:w&&(p=Math.max(S-r,Math.min(E-s,p)),o=r+p,l=s+p),M&&(v=Math.max(k-a,Math.min(C-h,v)),u=a+v,d=h+v);break;case gi:w<0?(p=Math.max(S-r,Math.min(E-r,p)),o=r+p,l=s):w>0&&(p=Math.max(S-s,Math.min(E-s,p)),o=r,l=s+p),M<0?(v=Math.max(k-a,Math.min(C-a,v)),u=a+v,d=h):M>0&&(v=Math.max(k-h,Math.min(C-h,v)),u=a,d=h+v);break;case yi:w&&(o=Math.max(S,Math.min(E,r-p*w)),l=Math.max(S,Math.min(E,s+p*w))),M&&(u=Math.max(k,Math.min(C,a-v*M)),d=Math.max(k,Math.min(C,h+v*M)))}l<o&&(w*=-1,t=r,r=s,s=t,t=o,o=l,l=t,m in wi&&L.attr("cursor",xi[m=wi[m]])),d<u&&(M*=-1,t=a,a=h,h=t,t=u,u=d,d=t,m in Mi&&L.attr("cursor",xi[m=Mi[m]])),N.selection&&(T=N.selection),y&&(o=T[0][0],l=T[1][0]),_&&(u=T[0][1],d=T[1][1]),T[0][0]===o&&T[0][1]===u&&T[1][0]===l&&T[1][1]===d||(N.selection=[[o,u],[l,d]],c.call(b),D.brush())}function Y(){if(hi(),t.event.touches){if(t.event.touches.length)return;e&&clearTimeout(e),e=setTimeout(function(){e=null},500),q.on("touchmove.brush touchend.brush touchcancel.brush",null)}else jt(t.event.view,g),U.on("keydown.brush keyup.brush mousemove.brush mouseup.brush",null);q.attr("pointer-events","all"),L.attr("cursor",xi.overlay),N.selection&&(T=N.selection),Ci(T)&&(N.selection=null,c.call(b)),D.end()}}function h(){var t=this.__brush||{selection:null};return t.extent=r.apply(this,arguments),t.dim=n,t}return u.move=function(t,e){t.selection?t.on("start.brush",function(){f(this,arguments).beforestart().start()}).on("interrupt.brush end.brush",function(){f(this,arguments).end()}).tween("brush",function(){var t=this,r=t.__brush,i=f(t,arguments),o=r.selection,a=n.input("function"==typeof e?e.apply(this,arguments):e,r.extent),u=ye(o,a);function s(n){r.selection=1===n&&Ci(a)?null:u(n),c.call(t),i.brush()}return o&&a?s:s(1)}):t.each(function(){var t=arguments,r=this.__brush,i=n.input("function"==typeof e?e.apply(this,t):e,r.extent),o=f(this,t).beforestart();Mr(this),r.selection=null==i||Ci(i)?null:i,c.call(this),o.start().brush().end()})},s.prototype={beforestart:function(){return 1==++this.active&&(this.state.emitter=this,this.starting=!0),this},start:function(){return this.starting&&(this.starting=!1,this.emit("start")),this},brush:function(){return this.emit("brush"),this},end:function(){return 0==--this.active&&(delete this.state.emitter,this.emit("end")),this},emit:function(t){St(new li(u,t,n.output(this.state.selection)),o.apply,o,[t,this.that,this.args])}},u.extent=function(t){return arguments.length?(r="function"==typeof t?t:si([[+t[0][0],+t[0][1]],[+t[1][0],+t[1][1]]]),u):r},u.filter=function(t){return arguments.length?(i="function"==typeof t?t:si(!!t),u):i},u.handleSize=function(t){return arguments.length?(a=+t,u):a},u.on=function(){var t=o.on.apply(o,arguments);return t===o?u:t},u}var zi=Math.cos,Ri=Math.sin,Di=Math.PI,qi=Di/2,Li=2*Di,Ui=Math.max;var Oi=Array.prototype.slice;function Bi(t){return function(){return t}}var Yi=Math.PI,Fi=2*Yi,Ii=Fi-1e-6;function ji(){this._x0=this._y0=this._x1=this._y1=null,this._=""}function Hi(){return new ji}function Xi(t){return t.source}function Gi(t){return t.target}function Vi(t){return t.radius}function $i(t){return t.startAngle}function Wi(t){return t.endAngle}ji.prototype=Hi.prototype={constructor:ji,moveTo:function(t,n){this._+="M"+(this._x0=this._x1=+t)+","+(this._y0=this._y1=+n)},closePath:function(){null!==this._x1&&(this._x1=this._x0,this._y1=this._y0,this._+="Z")},lineTo:function(t,n){this._+="L"+(this._x1=+t)+","+(this._y1=+n)},quadraticCurveTo:function(t,n,e,r){this._+="Q"+ +t+","+ +n+","+(this._x1=+e)+","+(this._y1=+r)},bezierCurveTo:function(t,n,e,r,i,o){this._+="C"+ +t+","+ +n+","+ +e+","+ +r+","+(this._x1=+i)+","+(this._y1=+o)},arcTo:function(t,n,e,r,i){t=+t,n=+n,e=+e,r=+r,i=+i;var o=this._x1,a=this._y1,u=e-t,c=r-n,f=o-t,s=a-n,l=f*f+s*s;if(i<0)throw new Error("negative radius: "+i);if(null===this._x1)this._+="M"+(this._x1=t)+","+(this._y1=n);else if(l>1e-6)if(Math.abs(s*u-c*f)>1e-6&&i){var h=e-o,d=r-a,p=u*u+c*c,v=h*h+d*d,g=Math.sqrt(p),y=Math.sqrt(l),_=i*Math.tan((Yi-Math.acos((p+l-v)/(2*g*y)))/2),b=_/y,m=_/g;Math.abs(b-1)>1e-6&&(this._+="L"+(t+b*f)+","+(n+b*s)),this._+="A"+i+","+i+",0,0,"+ +(s*h>f*d)+","+(this._x1=t+m*u)+","+(this._y1=n+m*c)}else this._+="L"+(this._x1=t)+","+(this._y1=n);else;},arc:function(t,n,e,r,i,o){t=+t,n=+n;var a=(e=+e)*Math.cos(r),u=e*Math.sin(r),c=t+a,f=n+u,s=1^o,l=o?r-i:i-r;if(e<0)throw new Error("negative radius: "+e);null===this._x1?this._+="M"+c+","+f:(Math.abs(this._x1-c)>1e-6||Math.abs(this._y1-f)>1e-6)&&(this._+="L"+c+","+f),e&&(l<0&&(l=l%Fi+Fi),l>Ii?this._+="A"+e+","+e+",0,1,"+s+","+(t-a)+","+(n-u)+"A"+e+","+e+",0,1,"+s+","+(this._x1=c)+","+(this._y1=f):l>1e-6&&(this._+="A"+e+","+e+",0,"+ +(l>=Yi)+","+s+","+(this._x1=t+e*Math.cos(i))+","+(this._y1=n+e*Math.sin(i))))},rect:function(t,n,e,r){this._+="M"+(this._x0=this._x1=+t)+","+(this._y0=this._y1=+n)+"h"+ +e+"v"+ +r+"h"+-e+"Z"},toString:function(){return this._}};function Zi(){}function Qi(t,n){var e=new Zi;if(t instanceof Zi)t.each(function(t,n){e.set(n,t)});else if(Array.isArray(t)){var r,i=-1,o=t.length;if(null==n)for(;++i<o;)e.set(i,t[i]);else for(;++i<o;)e.set(n(r=t[i],i,t),r)}else if(t)for(var a in t)e.set(a,t[a]);return e}function Ji(){return{}}function Ki(t,n,e){t[n]=e}function to(){return Qi()}function no(t,n,e){t.set(n,e)}function eo(){}Zi.prototype=Qi.prototype={constructor:Zi,has:function(t){return"$"+t in this},get:function(t){return this["$"+t]},set:function(t,n){return this["$"+t]=n,this},remove:function(t){var n="$"+t;return n in this&&delete this[n]},clear:function(){for(var t in this)"$"===t[0]&&delete this[t]},keys:function(){var t=[];for(var n in this)"$"===n[0]&&t.push(n.slice(1));return t},values:function(){var t=[];for(var n in this)"$"===n[0]&&t.push(this[n]);return t},entries:function(){var t=[];for(var n in this)"$"===n[0]&&t.push({key:n.slice(1),value:this[n]});return t},size:function(){var t=0;for(var n in this)"$"===n[0]&&++t;return t},empty:function(){for(var t in this)if("$"===t[0])return!1;return!0},each:function(t){for(var n in this)"$"===n[0]&&t(this[n],n.slice(1),this)}};var ro=Qi.prototype;function io(t,n){var e=new eo;if(t instanceof eo)t.each(function(t){e.add(t)});else if(t){var r=-1,i=t.length;if(null==n)for(;++r<i;)e.add(t[r]);else for(;++r<i;)e.add(n(t[r],r,t))}return e}eo.prototype=io.prototype={constructor:eo,has:ro.has,add:function(t){return this["$"+(t+="")]=t,this},remove:ro.remove,clear:ro.clear,values:ro.keys,size:ro.size,empty:ro.empty,each:ro.each};var oo=Array.prototype.slice;function ao(t,n){return t-n}function uo(t){return function(){return t}}function co(t,n){for(var e,r=-1,i=n.length;++r<i;)if(e=fo(t,n[r]))return e;return 0}function fo(t,n){for(var e=n[0],r=n[1],i=-1,o=0,a=t.length,u=a-1;o<a;u=o++){var c=t[o],f=c[0],s=c[1],l=t[u],h=l[0],d=l[1];if(so(c,l,n))return 0;s>r!=d>r&&e<(h-f)*(r-s)/(d-s)+f&&(i=-i)}return i}function so(t,n,e){var r,i,o,a;return function(t,n,e){return(n[0]-t[0])*(e[1]-t[1])==(e[0]-t[0])*(n[1]-t[1])}(t,n,e)&&(i=t[r=+(t[0]===n[0])],o=e[r],a=n[r],i<=o&&o<=a||a<=o&&o<=i)}function lo(){}var ho=[[],[[[1,1.5],[.5,1]]],[[[1.5,1],[1,1.5]]],[[[1.5,1],[.5,1]]],[[[1,.5],[1.5,1]]],[[[1,1.5],[.5,1]],[[1,.5],[1.5,1]]],[[[1,.5],[1,1.5]]],[[[1,.5],[.5,1]]],[[[.5,1],[1,.5]]],[[[1,1.5],[1,.5]]],[[[.5,1],[1,.5]],[[1.5,1],[1,1.5]]],[[[1.5,1],[1,.5]]],[[[.5,1],[1.5,1]]],[[[1,1.5],[1.5,1]]],[[[.5,1],[1,1.5]]],[]];function po(){var t=1,n=1,e=M,r=u;function i(t){var n=e(t);if(Array.isArray(n))n=n.slice().sort(ao);else{var r=s(t),i=r[0],a=r[1];n=w(i,a,n),n=g(Math.floor(i/n)*n,Math.floor(a/n)*n,n)}return n.map(function(n){return o(t,n)})}function o(e,i){var o=[],u=[];return function(e,r,i){var o,u,c,f,s,l,h=new Array,d=new Array;o=u=-1,f=e[0]>=r,ho[f<<1].forEach(p);for(;++o<t-1;)c=f,f=e[o+1]>=r,ho[c|f<<1].forEach(p);ho[f<<0].forEach(p);for(;++u<n-1;){for(o=-1,f=e[u*t+t]>=r,s=e[u*t]>=r,ho[f<<1|s<<2].forEach(p);++o<t-1;)c=f,f=e[u*t+t+o+1]>=r,l=s,s=e[u*t+o+1]>=r,ho[c|f<<1|s<<2|l<<3].forEach(p);ho[f|s<<3].forEach(p)}o=-1,s=e[u*t]>=r,ho[s<<2].forEach(p);for(;++o<t-1;)l=s,s=e[u*t+o+1]>=r,ho[s<<2|l<<3].forEach(p);function p(t){var n,e,r=[t[0][0]+o,t[0][1]+u],c=[t[1][0]+o,t[1][1]+u],f=a(r),s=a(c);(n=d[f])?(e=h[s])?(delete d[n.end],delete h[e.start],n===e?(n.ring.push(c),i(n.ring)):h[n.start]=d[e.end]={start:n.start,end:e.end,ring:n.ring.concat(e.ring)}):(delete d[n.end],n.ring.push(c),d[n.end=s]=n):(n=h[s])?(e=d[f])?(delete h[n.start],delete d[e.end],n===e?(n.ring.push(c),i(n.ring)):h[e.start]=d[n.end]={start:e.start,end:n.end,ring:e.ring.concat(n.ring)}):(delete h[n.start],n.ring.unshift(r),h[n.start=f]=n):h[f]=d[s]={start:f,end:s,ring:[r,c]}}ho[s<<3].forEach(p)}(e,i,function(t){r(t,e,i),function(t){for(var n=0,e=t.length,r=t[e-1][1]*t[0][0]-t[e-1][0]*t[0][1];++n<e;)r+=t[n-1][1]*t[n][0]-t[n-1][0]*t[n][1];return r}(t)>0?o.push([t]):u.push(t)}),u.forEach(function(t){for(var n,e=0,r=o.length;e<r;++e)if(-1!==co((n=o[e])[0],t))return void n.push(t)}),{type:"MultiPolygon",value:i,coordinates:o}}function a(n){return 2*n[0]+n[1]*(t+1)*4}function u(e,r,i){e.forEach(function(e){var o,a=e[0],u=e[1],c=0|a,f=0|u,s=r[f*t+c];a>0&&a<t&&c===a&&(o=r[f*t+c-1],e[0]=a+(i-o)/(s-o)-.5),u>0&&u<n&&f===u&&(o=r[(f-1)*t+c],e[1]=u+(i-o)/(s-o)-.5)})}return i.contour=o,i.size=function(e){if(!arguments.length)return[t,n];var r=Math.ceil(e[0]),o=Math.ceil(e[1]);if(!(r>0&&o>0))throw new Error("invalid size");return t=r,n=o,i},i.thresholds=function(t){return arguments.length?(e="function"==typeof t?t:Array.isArray(t)?uo(oo.call(t)):uo(t),i):e},i.smooth=function(t){return arguments.length?(r=t?u:lo,i):r===u},i}function vo(t,n,e){for(var r=t.width,i=t.height,o=1+(e<<1),a=0;a<i;++a)for(var u=0,c=0;u<r+e;++u)u<r&&(c+=t.data[u+a*r]),u>=e&&(u>=o&&(c-=t.data[u-o+a*r]),n.data[u-e+a*r]=c/Math.min(u+1,r-1+o-u,o))}function go(t,n,e){for(var r=t.width,i=t.height,o=1+(e<<1),a=0;a<r;++a)for(var u=0,c=0;u<i+e;++u)u<i&&(c+=t.data[a+u*r]),u>=e&&(u>=o&&(c-=t.data[a+(u-o)*r]),n.data[a+(u-e)*r]=c/Math.min(u+1,i-1+o-u,o))}function yo(t){return t[0]}function _o(t){return t[1]}function bo(){return 1}var mo={},xo={},wo=34,Mo=10,No=13;function Ao(t){return new Function("d","return {"+t.map(function(t,n){return JSON.stringify(t)+": d["+n+"]"}).join(",")+"}")}function To(t){var n=Object.create(null),e=[];return t.forEach(function(t){for(var r in t)r in n||e.push(n[r]=r)}),e}function So(t,n){var e=t+"",r=e.length;return r<n?new Array(n-r+1).join(0)+e:e}function ko(t){var n,e=t.getUTCHours(),r=t.getUTCMinutes(),i=t.getUTCSeconds(),o=t.getUTCMilliseconds();return isNaN(t)?"Invalid Date":((n=t.getUTCFullYear())<0?"-"+So(-n,6):n>9999?"+"+So(n,6):So(n,4))+"-"+So(t.getUTCMonth()+1,2)+"-"+So(t.getUTCDate(),2)+(o?"T"+So(e,2)+":"+So(r,2)+":"+So(i,2)+"."+So(o,3)+"Z":i?"T"+So(e,2)+":"+So(r,2)+":"+So(i,2)+"Z":r||e?"T"+So(e,2)+":"+So(r,2)+"Z":"")}function Eo(t){var n=new RegExp('["'+t+"\n\r]"),e=t.charCodeAt(0);function r(t,n){var r,i=[],o=t.length,a=0,u=0,c=o<=0,f=!1;function s(){if(c)return xo;if(f)return f=!1,mo;var n,r,i=a;if(t.charCodeAt(i)===wo){for(;a++<o&&t.charCodeAt(a)!==wo||t.charCodeAt(++a)===wo;);return(n=a)>=o?c=!0:(r=t.charCodeAt(a++))===Mo?f=!0:r===No&&(f=!0,t.charCodeAt(a)===Mo&&++a),t.slice(i+1,n-1).replace(/""/g,'"')}for(;a<o;){if((r=t.charCodeAt(n=a++))===Mo)f=!0;else if(r===No)f=!0,t.charCodeAt(a)===Mo&&++a;else if(r!==e)continue;return t.slice(i,n)}return c=!0,t.slice(i,o)}for(t.charCodeAt(o-1)===Mo&&--o,t.charCodeAt(o-1)===No&&--o;(r=s())!==xo;){for(var l=[];r!==mo&&r!==xo;)l.push(r),r=s();n&&null==(l=n(l,u++))||i.push(l)}return i}function i(n,e){return n.map(function(n){return e.map(function(t){return a(n[t])}).join(t)})}function o(n){return n.map(a).join(t)}function a(t){return null==t?"":t instanceof Date?ko(t):n.test(t+="")?'"'+t.replace(/"/g,'""')+'"':t}return{parse:function(t,n){var e,i,o=r(t,function(t,r){if(e)return e(t,r-1);i=t,e=n?function(t,n){var e=Ao(t);return function(r,i){return n(e(r),i,t)}}(t,n):Ao(t)});return o.columns=i||[],o},parseRows:r,format:function(n,e){return null==e&&(e=To(n)),[e.map(a).join(t)].concat(i(n,e)).join("\n")},formatBody:function(t,n){return null==n&&(n=To(t)),i(t,n).join("\n")},formatRows:function(t){return t.map(o).join("\n")}}}var Co=Eo(","),Po=Co.parse,zo=Co.parseRows,Ro=Co.format,Do=Co.formatBody,qo=Co.formatRows,Lo=Eo("\t"),Uo=Lo.parse,Oo=Lo.parseRows,Bo=Lo.format,Yo=Lo.formatBody,Fo=Lo.formatRows;function Io(t){if(!t.ok)throw new Error(t.status+" "+t.statusText);return t.blob()}function jo(t){if(!t.ok)throw new Error(t.status+" "+t.statusText);return t.arrayBuffer()}function Ho(t){if(!t.ok)throw new Error(t.status+" "+t.statusText);return t.text()}function Xo(t,n){return fetch(t,n).then(Ho)}function Go(t){return function(n,e,r){return 2===arguments.length&&"function"==typeof e&&(r=e,e=void 0),Xo(n,e).then(function(n){return t(n,r)})}}var Vo=Go(Po),$o=Go(Uo);function Wo(t){if(!t.ok)throw new Error(t.status+" "+t.statusText);return t.json()}function Zo(t){return function(n,e){return Xo(n,e).then(function(n){return(new DOMParser).parseFromString(n,t)})}}var Qo=Zo("application/xml"),Jo=Zo("text/html"),Ko=Zo("image/svg+xml");function ta(t){return function(){return t}}function na(){return 1e-6*(Math.random()-.5)}function ea(t,n,e,r){if(isNaN(n)||isNaN(e))return t;var i,o,a,u,c,f,s,l,h,d=t._root,p={data:r},v=t._x0,g=t._y0,y=t._x1,_=t._y1;if(!d)return t._root=p,t;for(;d.length;)if((f=n>=(o=(v+y)/2))?v=o:y=o,(s=e>=(a=(g+_)/2))?g=a:_=a,i=d,!(d=d[l=s<<1|f]))return i[l]=p,t;if(u=+t._x.call(null,d.data),c=+t._y.call(null,d.data),n===u&&e===c)return p.next=d,i?i[l]=p:t._root=p,t;do{i=i?i[l]=new Array(4):t._root=new Array(4),(f=n>=(o=(v+y)/2))?v=o:y=o,(s=e>=(a=(g+_)/2))?g=a:_=a}while((l=s<<1|f)==(h=(c>=a)<<1|u>=o));return i[h]=d,i[l]=p,t}function ra(t,n,e,r,i){this.node=t,this.x0=n,this.y0=e,this.x1=r,this.y1=i}function ia(t){return t[0]}function oa(t){return t[1]}function aa(t,n,e){var r=new ua(null==n?ia:n,null==e?oa:e,NaN,NaN,NaN,NaN);return null==t?r:r.addAll(t)}function ua(t,n,e,r,i,o){this._x=t,this._y=n,this._x0=e,this._y0=r,this._x1=i,this._y1=o,this._root=void 0}function ca(t){for(var n={data:t.data},e=n;t=t.next;)e=e.next={data:t.data};return n}var fa=aa.prototype=ua.prototype;function sa(t){return t.x+t.vx}function la(t){return t.y+t.vy}function ha(t){return t.index}function da(t,n){var e=t.get(n);if(!e)throw new Error("missing: "+n);return e}function pa(t){return t.x}function va(t){return t.y}fa.copy=function(){var t,n,e=new ua(this._x,this._y,this._x0,this._y0,this._x1,this._y1),r=this._root;if(!r)return e;if(!r.length)return e._root=ca(r),e;for(t=[{source:r,target:e._root=new Array(4)}];r=t.pop();)for(var i=0;i<4;++i)(n=r.source[i])&&(n.length?t.push({source:n,target:r.target[i]=new Array(4)}):r.target[i]=ca(n));return e},fa.add=function(t){var n=+this._x.call(null,t),e=+this._y.call(null,t);return ea(this.cover(n,e),n,e,t)},fa.addAll=function(t){var n,e,r,i,o=t.length,a=new Array(o),u=new Array(o),c=1/0,f=1/0,s=-1/0,l=-1/0;for(e=0;e<o;++e)isNaN(r=+this._x.call(null,n=t[e]))||isNaN(i=+this._y.call(null,n))||(a[e]=r,u[e]=i,r<c&&(c=r),r>s&&(s=r),i<f&&(f=i),i>l&&(l=i));if(c>s||f>l)return this;for(this.cover(c,f).cover(s,l),e=0;e<o;++e)ea(this,a[e],u[e],t[e]);return this},fa.cover=function(t,n){if(isNaN(t=+t)||isNaN(n=+n))return this;var e=this._x0,r=this._y0,i=this._x1,o=this._y1;if(isNaN(e))i=(e=Math.floor(t))+1,o=(r=Math.floor(n))+1;else{for(var a,u,c=i-e,f=this._root;e>t||t>=i||r>n||n>=o;)switch(u=(n<r)<<1|t<e,(a=new Array(4))[u]=f,f=a,c*=2,u){case 0:i=e+c,o=r+c;break;case 1:e=i-c,o=r+c;break;case 2:i=e+c,r=o-c;break;case 3:e=i-c,r=o-c}this._root&&this._root.length&&(this._root=f)}return this._x0=e,this._y0=r,this._x1=i,this._y1=o,this},fa.data=function(){var t=[];return this.visit(function(n){if(!n.length)do{t.push(n.data)}while(n=n.next)}),t},fa.extent=function(t){return arguments.length?this.cover(+t[0][0],+t[0][1]).cover(+t[1][0],+t[1][1]):isNaN(this._x0)?void 0:[[this._x0,this._y0],[this._x1,this._y1]]},fa.find=function(t,n,e){var r,i,o,a,u,c,f,s=this._x0,l=this._y0,h=this._x1,d=this._y1,p=[],v=this._root;for(v&&p.push(new ra(v,s,l,h,d)),null==e?e=1/0:(s=t-e,l=n-e,h=t+e,d=n+e,e*=e);c=p.pop();)if(!(!(v=c.node)||(i=c.x0)>h||(o=c.y0)>d||(a=c.x1)<s||(u=c.y1)<l))if(v.length){var g=(i+a)/2,y=(o+u)/2;p.push(new ra(v[3],g,y,a,u),new ra(v[2],i,y,g,u),new ra(v[1],g,o,a,y),new ra(v[0],i,o,g,y)),(f=(n>=y)<<1|t>=g)&&(c=p[p.length-1],p[p.length-1]=p[p.length-1-f],p[p.length-1-f]=c)}else{var _=t-+this._x.call(null,v.data),b=n-+this._y.call(null,v.data),m=_*_+b*b;if(m<e){var x=Math.sqrt(e=m);s=t-x,l=n-x,h=t+x,d=n+x,r=v.data}}return r},fa.remove=function(t){if(isNaN(o=+this._x.call(null,t))||isNaN(a=+this._y.call(null,t)))return this;var n,e,r,i,o,a,u,c,f,s,l,h,d=this._root,p=this._x0,v=this._y0,g=this._x1,y=this._y1;if(!d)return this;if(d.length)for(;;){if((f=o>=(u=(p+g)/2))?p=u:g=u,(s=a>=(c=(v+y)/2))?v=c:y=c,n=d,!(d=d[l=s<<1|f]))return this;if(!d.length)break;(n[l+1&3]||n[l+2&3]||n[l+3&3])&&(e=n,h=l)}for(;d.data!==t;)if(r=d,!(d=d.next))return this;return(i=d.next)&&delete d.next,r?(i?r.next=i:delete r.next,this):n?(i?n[l]=i:delete n[l],(d=n[0]||n[1]||n[2]||n[3])&&d===(n[3]||n[2]||n[1]||n[0])&&!d.length&&(e?e[h]=d:this._root=d),this):(this._root=i,this)},fa.removeAll=function(t){for(var n=0,e=t.length;n<e;++n)this.remove(t[n]);return this},fa.root=function(){return this._root},fa.size=function(){var t=0;return this.visit(function(n){if(!n.length)do{++t}while(n=n.next)}),t},fa.visit=function(t){var n,e,r,i,o,a,u=[],c=this._root;for(c&&u.push(new ra(c,this._x0,this._y0,this._x1,this._y1));n=u.pop();)if(!t(c=n.node,r=n.x0,i=n.y0,o=n.x1,a=n.y1)&&c.length){var f=(r+o)/2,s=(i+a)/2;(e=c[3])&&u.push(new ra(e,f,s,o,a)),(e=c[2])&&u.push(new ra(e,r,s,f,a)),(e=c[1])&&u.push(new ra(e,f,i,o,s)),(e=c[0])&&u.push(new ra(e,r,i,f,s))}return this},fa.visitAfter=function(t){var n,e=[],r=[];for(this._root&&e.push(new ra(this._root,this._x0,this._y0,this._x1,this._y1));n=e.pop();){var i=n.node;if(i.length){var o,a=n.x0,u=n.y0,c=n.x1,f=n.y1,s=(a+c)/2,l=(u+f)/2;(o=i[0])&&e.push(new ra(o,a,u,s,l)),(o=i[1])&&e.push(new ra(o,s,u,c,l)),(o=i[2])&&e.push(new ra(o,a,l,s,f)),(o=i[3])&&e.push(new ra(o,s,l,c,f))}r.push(n)}for(;n=r.pop();)t(n.node,n.x0,n.y0,n.x1,n.y1);return this},fa.x=function(t){return arguments.length?(this._x=t,this):this._x},fa.y=function(t){return arguments.length?(this._y=t,this):this._y};var ga=10,ya=Math.PI*(3-Math.sqrt(5));function _a(t,n){if((e=(t=n?t.toExponential(n-1):t.toExponential()).indexOf("e"))<0)return null;var e,r=t.slice(0,e);return[r.length>1?r[0]+r.slice(2):r,+t.slice(e+1)]}function ba(t){return(t=_a(Math.abs(t)))?t[1]:NaN}var ma,xa=/^(?:(.)?([<>=^]))?([+\-( ])?([$#])?(0)?(\d+)?(,)?(\.\d+)?(~)?([a-z%])?$/i;function wa(t){return new Ma(t)}function Ma(t){if(!(n=xa.exec(t)))throw new Error("invalid format: "+t);var n;this.fill=n[1]||" ",this.align=n[2]||">",this.sign=n[3]||"-",this.symbol=n[4]||"",this.zero=!!n[5],this.width=n[6]&&+n[6],this.comma=!!n[7],this.precision=n[8]&&+n[8].slice(1),this.trim=!!n[9],this.type=n[10]||""}function Na(t,n){var e=_a(t,n);if(!e)return t+"";var r=e[0],i=e[1];return i<0?"0."+new Array(-i).join("0")+r:r.length>i+1?r.slice(0,i+1)+"."+r.slice(i+1):r+new Array(i-r.length+2).join("0")}wa.prototype=Ma.prototype,Ma.prototype.toString=function(){return this.fill+this.align+this.sign+this.symbol+(this.zero?"0":"")+(null==this.width?"":Math.max(1,0|this.width))+(this.comma?",":"")+(null==this.precision?"":"."+Math.max(0,0|this.precision))+(this.trim?"~":"")+this.type};var Aa={"%":function(t,n){return(100*t).toFixed(n)},b:function(t){return Math.round(t).toString(2)},c:function(t){return t+""},d:function(t){return Math.round(t).toString(10)},e:function(t,n){return t.toExponential(n)},f:function(t,n){return t.toFixed(n)},g:function(t,n){return t.toPrecision(n)},o:function(t){return Math.round(t).toString(8)},p:function(t,n){return Na(100*t,n)},r:Na,s:function(t,n){var e=_a(t,n);if(!e)return t+"";var r=e[0],i=e[1],o=i-(ma=3*Math.max(-8,Math.min(8,Math.floor(i/3))))+1,a=r.length;return o===a?r:o>a?r+new Array(o-a+1).join("0"):o>0?r.slice(0,o)+"."+r.slice(o):"0."+new Array(1-o).join("0")+_a(t,Math.max(0,n+o-1))[0]},X:function(t){return Math.round(t).toString(16).toUpperCase()},x:function(t){return Math.round(t).toString(16)}};function Ta(t){return t}var Sa,ka=["y","z","a","f","p","n","µ","m","","k","M","G","T","P","E","Z","Y"];function Ea(t){var n,e,r=t.grouping&&t.thousands?(n=t.grouping,e=t.thousands,function(t,r){for(var i=t.length,o=[],a=0,u=n[0],c=0;i>0&&u>0&&(c+u+1>r&&(u=Math.max(1,r-c)),o.push(t.substring(i-=u,i+u)),!((c+=u+1)>r));)u=n[a=(a+1)%n.length];return o.reverse().join(e)}):Ta,i=t.currency,o=t.decimal,a=t.numerals?function(t){return function(n){return n.replace(/[0-9]/g,function(n){return t[+n]})}}(t.numerals):Ta,u=t.percent||"%";function c(t){var n=(t=wa(t)).fill,e=t.align,c=t.sign,f=t.symbol,s=t.zero,l=t.width,h=t.comma,d=t.precision,p=t.trim,v=t.type;"n"===v?(h=!0,v="g"):Aa[v]||(null==d&&(d=12),p=!0,v="g"),(s||"0"===n&&"="===e)&&(s=!0,n="0",e="=");var g="$"===f?i[0]:"#"===f&&/[boxX]/.test(v)?"0"+v.toLowerCase():"",y="$"===f?i[1]:/[%p]/.test(v)?u:"",_=Aa[v],b=/[defgprs%]/.test(v);function m(t){var i,u,f,m=g,x=y;if("c"===v)x=_(t)+x,t="";else{var w=(t=+t)<0;if(t=_(Math.abs(t),d),p&&(t=function(t){t:for(var n,e=t.length,r=1,i=-1;r<e;++r)switch(t[r]){case".":i=n=r;break;case"0":0===i&&(i=r),n=r;break;default:if(i>0){if(!+t[r])break t;i=0}}return i>0?t.slice(0,i)+t.slice(n+1):t}(t)),w&&0==+t&&(w=!1),m=(w?"("===c?c:"-":"-"===c||"("===c?"":c)+m,x=("s"===v?ka[8+ma/3]:"")+x+(w&&"("===c?")":""),b)for(i=-1,u=t.length;++i<u;)if(48>(f=t.charCodeAt(i))||f>57){x=(46===f?o+t.slice(i+1):t.slice(i))+x,t=t.slice(0,i);break}}h&&!s&&(t=r(t,1/0));var M=m.length+t.length+x.length,N=M<l?new Array(l-M+1).join(n):"";switch(h&&s&&(t=r(N+t,N.length?l-x.length:1/0),N=""),e){case"<":t=m+t+x+N;break;case"=":t=m+N+t+x;break;case"^":t=N.slice(0,M=N.length>>1)+m+t+x+N.slice(M);break;default:t=N+m+t+x}return a(t)}return d=null==d?6:/[gprs]/.test(v)?Math.max(1,Math.min(21,d)):Math.max(0,Math.min(20,d)),m.toString=function(){return t+""},m}return{format:c,formatPrefix:function(t,n){var e=c(((t=wa(t)).type="f",t)),r=3*Math.max(-8,Math.min(8,Math.floor(ba(n)/3))),i=Math.pow(10,-r),o=ka[8+r/3];return function(t){return e(i*t)+o}}}}function Ca(n){return Sa=Ea(n),t.format=Sa.format,t.formatPrefix=Sa.formatPrefix,Sa}function Pa(t){return Math.max(0,-ba(Math.abs(t)))}function za(t,n){return Math.max(0,3*Math.max(-8,Math.min(8,Math.floor(ba(n)/3)))-ba(Math.abs(t)))}function Ra(t,n){return t=Math.abs(t),n=Math.abs(n)-t,Math.max(0,ba(n)-ba(t))+1}function Da(){return new qa}function qa(){this.reset()}Ca({decimal:".",thousands:",",grouping:[3],currency:["$",""]}),qa.prototype={constructor:qa,reset:function(){this.s=this.t=0},add:function(t){Ua(La,t,this.t),Ua(this,La.s,this.s),this.s?this.t+=La.t:this.s=La.t},valueOf:function(){return this.s}};var La=new qa;function Ua(t,n,e){var r=t.s=n+e,i=r-n,o=r-i;t.t=n-o+(e-i)}var Oa=1e-6,Ba=1e-12,Ya=Math.PI,Fa=Ya/2,Ia=Ya/4,ja=2*Ya,Ha=180/Ya,Xa=Ya/180,Ga=Math.abs,Va=Math.atan,$a=Math.atan2,Wa=Math.cos,Za=Math.ceil,Qa=Math.exp,Ja=Math.log,Ka=Math.pow,tu=Math.sin,nu=Math.sign||function(t){return t>0?1:t<0?-1:0},eu=Math.sqrt,ru=Math.tan;function iu(t){return t>1?0:t<-1?Ya:Math.acos(t)}function ou(t){return t>1?Fa:t<-1?-Fa:Math.asin(t)}function au(t){return(t=tu(t/2))*t}function uu(){}function cu(t,n){t&&su.hasOwnProperty(t.type)&&su[t.type](t,n)}var fu={Feature:function(t,n){cu(t.geometry,n)},FeatureCollection:function(t,n){for(var e=t.features,r=-1,i=e.length;++r<i;)cu(e[r].geometry,n)}},su={Sphere:function(t,n){n.sphere()},Point:function(t,n){t=t.coordinates,n.point(t[0],t[1],t[2])},MultiPoint:function(t,n){for(var e=t.coordinates,r=-1,i=e.length;++r<i;)t=e[r],n.point(t[0],t[1],t[2])},LineString:function(t,n){lu(t.coordinates,n,0)},MultiLineString:function(t,n){for(var e=t.coordinates,r=-1,i=e.length;++r<i;)lu(e[r],n,0)},Polygon:function(t,n){hu(t.coordinates,n)},MultiPolygon:function(t,n){for(var e=t.coordinates,r=-1,i=e.length;++r<i;)hu(e[r],n)},GeometryCollection:function(t,n){for(var e=t.geometries,r=-1,i=e.length;++r<i;)cu(e[r],n)}};function lu(t,n,e){var r,i=-1,o=t.length-e;for(n.lineStart();++i<o;)r=t[i],n.point(r[0],r[1],r[2]);n.lineEnd()}function hu(t,n){var e=-1,r=t.length;for(n.polygonStart();++e<r;)lu(t[e],n,1);n.polygonEnd()}function du(t,n){t&&fu.hasOwnProperty(t.type)?fu[t.type](t,n):cu(t,n)}var pu,vu,gu,yu,_u,bu=Da(),mu=Da(),xu={point:uu,lineStart:uu,lineEnd:uu,polygonStart:function(){bu.reset(),xu.lineStart=wu,xu.lineEnd=Mu},polygonEnd:function(){var t=+bu;mu.add(t<0?ja+t:t),this.lineStart=this.lineEnd=this.point=uu},sphere:function(){mu.add(ja)}};function wu(){xu.point=Nu}function Mu(){Au(pu,vu)}function Nu(t,n){xu.point=Au,pu=t,vu=n,gu=t*=Xa,yu=Wa(n=(n*=Xa)/2+Ia),_u=tu(n)}function Au(t,n){var e=(t*=Xa)-gu,r=e>=0?1:-1,i=r*e,o=Wa(n=(n*=Xa)/2+Ia),a=tu(n),u=_u*a,c=yu*o+u*Wa(i),f=u*r*tu(i);bu.add($a(f,c)),gu=t,yu=o,_u=a}function Tu(t){return[$a(t[1],t[0]),ou(t[2])]}function Su(t){var n=t[0],e=t[1],r=Wa(e);return[r*Wa(n),r*tu(n),tu(e)]}function ku(t,n){return t[0]*n[0]+t[1]*n[1]+t[2]*n[2]}function Eu(t,n){return[t[1]*n[2]-t[2]*n[1],t[2]*n[0]-t[0]*n[2],t[0]*n[1]-t[1]*n[0]]}function Cu(t,n){t[0]+=n[0],t[1]+=n[1],t[2]+=n[2]}function Pu(t,n){return[t[0]*n,t[1]*n,t[2]*n]}function zu(t){var n=eu(t[0]*t[0]+t[1]*t[1]+t[2]*t[2]);t[0]/=n,t[1]/=n,t[2]/=n}var Ru,Du,qu,Lu,Uu,Ou,Bu,Yu,Fu,Iu,ju,Hu,Xu,Gu,Vu,$u,Wu,Zu,Qu,Ju,Ku,tc,nc,ec,rc,ic,oc=Da(),ac={point:uc,lineStart:fc,lineEnd:sc,polygonStart:function(){ac.point=lc,ac.lineStart=hc,ac.lineEnd=dc,oc.reset(),xu.polygonStart()},polygonEnd:function(){xu.polygonEnd(),ac.point=uc,ac.lineStart=fc,ac.lineEnd=sc,bu<0?(Ru=-(qu=180),Du=-(Lu=90)):oc>Oa?Lu=90:oc<-Oa&&(Du=-90),Iu[0]=Ru,Iu[1]=qu}};function uc(t,n){Fu.push(Iu=[Ru=t,qu=t]),n<Du&&(Du=n),n>Lu&&(Lu=n)}function cc(t,n){var e=Su([t*Xa,n*Xa]);if(Yu){var r=Eu(Yu,e),i=Eu([r[1],-r[0],0],r);zu(i),i=Tu(i);var o,a=t-Uu,u=a>0?1:-1,c=i[0]*Ha*u,f=Ga(a)>180;f^(u*Uu<c&&c<u*t)?(o=i[1]*Ha)>Lu&&(Lu=o):f^(u*Uu<(c=(c+360)%360-180)&&c<u*t)?(o=-i[1]*Ha)<Du&&(Du=o):(n<Du&&(Du=n),n>Lu&&(Lu=n)),f?t<Uu?pc(Ru,t)>pc(Ru,qu)&&(qu=t):pc(t,qu)>pc(Ru,qu)&&(Ru=t):qu>=Ru?(t<Ru&&(Ru=t),t>qu&&(qu=t)):t>Uu?pc(Ru,t)>pc(Ru,qu)&&(qu=t):pc(t,qu)>pc(Ru,qu)&&(Ru=t)}else Fu.push(Iu=[Ru=t,qu=t]);n<Du&&(Du=n),n>Lu&&(Lu=n),Yu=e,Uu=t}function fc(){ac.point=cc}function sc(){Iu[0]=Ru,Iu[1]=qu,ac.point=uc,Yu=null}function lc(t,n){if(Yu){var e=t-Uu;oc.add(Ga(e)>180?e+(e>0?360:-360):e)}else Ou=t,Bu=n;xu.point(t,n),cc(t,n)}function hc(){xu.lineStart()}function dc(){lc(Ou,Bu),xu.lineEnd(),Ga(oc)>Oa&&(Ru=-(qu=180)),Iu[0]=Ru,Iu[1]=qu,Yu=null}function pc(t,n){return(n-=t)<0?n+360:n}function vc(t,n){return t[0]-n[0]}function gc(t,n){return t[0]<=t[1]?t[0]<=n&&n<=t[1]:n<t[0]||t[1]<n}var yc={sphere:uu,point:_c,lineStart:mc,lineEnd:Mc,polygonStart:function(){yc.lineStart=Nc,yc.lineEnd=Ac},polygonEnd:function(){yc.lineStart=mc,yc.lineEnd=Mc}};function _c(t,n){t*=Xa;var e=Wa(n*=Xa);bc(e*Wa(t),e*tu(t),tu(n))}function bc(t,n,e){Xu+=(t-Xu)/++ju,Gu+=(n-Gu)/ju,Vu+=(e-Vu)/ju}function mc(){yc.point=xc}function xc(t,n){t*=Xa;var e=Wa(n*=Xa);ec=e*Wa(t),rc=e*tu(t),ic=tu(n),yc.point=wc,bc(ec,rc,ic)}function wc(t,n){t*=Xa;var e=Wa(n*=Xa),r=e*Wa(t),i=e*tu(t),o=tu(n),a=$a(eu((a=rc*o-ic*i)*a+(a=ic*r-ec*o)*a+(a=ec*i-rc*r)*a),ec*r+rc*i+ic*o);Hu+=a,$u+=a*(ec+(ec=r)),Wu+=a*(rc+(rc=i)),Zu+=a*(ic+(ic=o)),bc(ec,rc,ic)}function Mc(){yc.point=_c}function Nc(){yc.point=Tc}function Ac(){Sc(tc,nc),yc.point=_c}function Tc(t,n){tc=t,nc=n,t*=Xa,n*=Xa,yc.point=Sc;var e=Wa(n);ec=e*Wa(t),rc=e*tu(t),ic=tu(n),bc(ec,rc,ic)}function Sc(t,n){t*=Xa;var e=Wa(n*=Xa),r=e*Wa(t),i=e*tu(t),o=tu(n),a=rc*o-ic*i,u=ic*r-ec*o,c=ec*i-rc*r,f=eu(a*a+u*u+c*c),s=ou(f),l=f&&-s/f;Qu+=l*a,Ju+=l*u,Ku+=l*c,Hu+=s,$u+=s*(ec+(ec=r)),Wu+=s*(rc+(rc=i)),Zu+=s*(ic+(ic=o)),bc(ec,rc,ic)}function kc(t){return function(){return t}}function Ec(t,n){function e(e,r){return e=t(e,r),n(e[0],e[1])}return t.invert&&n.invert&&(e.invert=function(e,r){return(e=n.invert(e,r))&&t.invert(e[0],e[1])}),e}function Cc(t,n){return[Ga(t)>Ya?t+Math.round(-t/ja)*ja:t,n]}function Pc(t,n,e){return(t%=ja)?n||e?Ec(Rc(t),Dc(n,e)):Rc(t):n||e?Dc(n,e):Cc}function zc(t){return function(n,e){return[(n+=t)>Ya?n-ja:n<-Ya?n+ja:n,e]}}function Rc(t){var n=zc(t);return n.invert=zc(-t),n}function Dc(t,n){var e=Wa(t),r=tu(t),i=Wa(n),o=tu(n);function a(t,n){var a=Wa(n),u=Wa(t)*a,c=tu(t)*a,f=tu(n),s=f*e+u*r;return[$a(c*i-s*o,u*e-f*r),ou(s*i+c*o)]}return a.invert=function(t,n){var a=Wa(n),u=Wa(t)*a,c=tu(t)*a,f=tu(n),s=f*i-c*o;return[$a(c*i+f*o,u*e+s*r),ou(s*e-u*r)]},a}function qc(t){function n(n){return(n=t(n[0]*Xa,n[1]*Xa))[0]*=Ha,n[1]*=Ha,n}return t=Pc(t[0]*Xa,t[1]*Xa,t.length>2?t[2]*Xa:0),n.invert=function(n){return(n=t.invert(n[0]*Xa,n[1]*Xa))[0]*=Ha,n[1]*=Ha,n},n}function Lc(t,n,e,r,i,o){if(e){var a=Wa(n),u=tu(n),c=r*e;null==i?(i=n+r*ja,o=n-c/2):(i=Uc(a,i),o=Uc(a,o),(r>0?i<o:i>o)&&(i+=r*ja));for(var f,s=i;r>0?s>o:s<o;s-=c)f=Tu([a,-u*Wa(s),-u*tu(s)]),t.point(f[0],f[1])}}function Uc(t,n){(n=Su(n))[0]-=t,zu(n);var e=iu(-n[1]);return((-n[2]<0?-e:e)+ja-Oa)%ja}function Oc(){var t,n=[];return{point:function(n,e){t.push([n,e])},lineStart:function(){n.push(t=[])},lineEnd:uu,rejoin:function(){n.length>1&&n.push(n.pop().concat(n.shift()))},result:function(){var e=n;return n=[],t=null,e}}}function Bc(t,n){return Ga(t[0]-n[0])<Oa&&Ga(t[1]-n[1])<Oa}function Yc(t,n,e,r){this.x=t,this.z=n,this.o=e,this.e=r,this.v=!1,this.n=this.p=null}function Fc(t,n,e,r,i){var o,a,u=[],c=[];if(t.forEach(function(t){if(!((n=t.length-1)<=0)){var n,e,r=t[0],a=t[n];if(Bc(r,a)){for(i.lineStart(),o=0;o<n;++o)i.point((r=t[o])[0],r[1]);i.lineEnd()}else u.push(e=new Yc(r,t,null,!0)),c.push(e.o=new Yc(r,null,e,!1)),u.push(e=new Yc(a,t,null,!1)),c.push(e.o=new Yc(a,null,e,!0))}}),u.length){for(c.sort(n),Ic(u),Ic(c),o=0,a=c.length;o<a;++o)c[o].e=e=!e;for(var f,s,l=u[0];;){for(var h=l,d=!0;h.v;)if((h=h.n)===l)return;f=h.z,i.lineStart();do{if(h.v=h.o.v=!0,h.e){if(d)for(o=0,a=f.length;o<a;++o)i.point((s=f[o])[0],s[1]);else r(h.x,h.n.x,1,i);h=h.n}else{if(d)for(f=h.p.z,o=f.length-1;o>=0;--o)i.point((s=f[o])[0],s[1]);else r(h.x,h.p.x,-1,i);h=h.p}f=(h=h.o).z,d=!d}while(!h.v);i.lineEnd()}}}function Ic(t){if(n=t.length){for(var n,e,r=0,i=t[0];++r<n;)i.n=e=t[r],e.p=i,i=e;i.n=e=t[0],e.p=i}}Cc.invert=Cc;var jc=Da();function Hc(t,n){var e=n[0],r=n[1],i=tu(r),o=[tu(e),-Wa(e),0],a=0,u=0;jc.reset(),1===i?r=Fa+Oa:-1===i&&(r=-Fa-Oa);for(var c=0,f=t.length;c<f;++c)if(l=(s=t[c]).length)for(var s,l,h=s[l-1],d=h[0],p=h[1]/2+Ia,v=tu(p),g=Wa(p),y=0;y<l;++y,d=b,v=x,g=w,h=_){var _=s[y],b=_[0],m=_[1]/2+Ia,x=tu(m),w=Wa(m),M=b-d,N=M>=0?1:-1,A=N*M,T=A>Ya,S=v*x;if(jc.add($a(S*N*tu(A),g*w+S*Wa(A))),a+=T?M+N*ja:M,T^d>=e^b>=e){var k=Eu(Su(h),Su(_));zu(k);var E=Eu(o,k);zu(E);var C=(T^M>=0?-1:1)*ou(E[2]);(r>C||r===C&&(k[0]||k[1]))&&(u+=T^M>=0?1:-1)}}return(a<-Oa||a<Oa&&jc<-Oa)^1&u}function Xc(t,n,e,r){return function(i){var o,a,u,c=n(i),f=Oc(),s=n(f),l=!1,h={point:d,lineStart:v,lineEnd:g,polygonStart:function(){h.point=y,h.lineStart=_,h.lineEnd=b,a=[],o=[]},polygonEnd:function(){h.point=d,h.lineStart=v,h.lineEnd=g,a=T(a);var t=Hc(o,r);a.length?(l||(i.polygonStart(),l=!0),Fc(a,Vc,t,e,i)):t&&(l||(i.polygonStart(),l=!0),i.lineStart(),e(null,null,1,i),i.lineEnd()),l&&(i.polygonEnd(),l=!1),a=o=null},sphere:function(){i.polygonStart(),i.lineStart(),e(null,null,1,i),i.lineEnd(),i.polygonEnd()}};function d(n,e){t(n,e)&&i.point(n,e)}function p(t,n){c.point(t,n)}function v(){h.point=p,c.lineStart()}function g(){h.point=d,c.lineEnd()}function y(t,n){u.push([t,n]),s.point(t,n)}function _(){s.lineStart(),u=[]}function b(){y(u[0][0],u[0][1]),s.lineEnd();var t,n,e,r,c=s.clean(),h=f.result(),d=h.length;if(u.pop(),o.push(u),u=null,d)if(1&c){if((n=(e=h[0]).length-1)>0){for(l||(i.polygonStart(),l=!0),i.lineStart(),t=0;t<n;++t)i.point((r=e[t])[0],r[1]);i.lineEnd()}}else d>1&&2&c&&h.push(h.pop().concat(h.shift())),a.push(h.filter(Gc))}return h}}function Gc(t){return t.length>1}function Vc(t,n){return((t=t.x)[0]<0?t[1]-Fa-Oa:Fa-t[1])-((n=n.x)[0]<0?n[1]-Fa-Oa:Fa-n[1])}var $c=Xc(function(){return!0},function(t){var n,e=NaN,r=NaN,i=NaN;return{lineStart:function(){t.lineStart(),n=1},point:function(o,a){var u=o>0?Ya:-Ya,c=Ga(o-e);Ga(c-Ya)<Oa?(t.point(e,r=(r+a)/2>0?Fa:-Fa),t.point(i,r),t.lineEnd(),t.lineStart(),t.point(u,r),t.point(o,r),n=0):i!==u&&c>=Ya&&(Ga(e-i)<Oa&&(e-=i*Oa),Ga(o-u)<Oa&&(o-=u*Oa),r=function(t,n,e,r){var i,o,a=tu(t-e);return Ga(a)>Oa?Va((tu(n)*(o=Wa(r))*tu(e)-tu(r)*(i=Wa(n))*tu(t))/(i*o*a)):(n+r)/2}(e,r,o,a),t.point(i,r),t.lineEnd(),t.lineStart(),t.point(u,r),n=0),t.point(e=o,r=a),i=u},lineEnd:function(){t.lineEnd(),e=r=NaN},clean:function(){return 2-n}}},function(t,n,e,r){var i;if(null==t)i=e*Fa,r.point(-Ya,i),r.point(0,i),r.point(Ya,i),r.point(Ya,0),r.point(Ya,-i),r.point(0,-i),r.point(-Ya,-i),r.point(-Ya,0),r.point(-Ya,i);else if(Ga(t[0]-n[0])>Oa){var o=t[0]<n[0]?Ya:-Ya;i=e*o/2,r.point(-o,i),r.point(0,i),r.point(o,i)}else r.point(n[0],n[1])},[-Ya,-Fa]);function Wc(t){var n=Wa(t),e=6*Xa,r=n>0,i=Ga(n)>Oa;function o(t,e){return Wa(t)*Wa(e)>n}function a(t,e,r){var i=[1,0,0],o=Eu(Su(t),Su(e)),a=ku(o,o),u=o[0],c=a-u*u;if(!c)return!r&&t;var f=n*a/c,s=-n*u/c,l=Eu(i,o),h=Pu(i,f);Cu(h,Pu(o,s));var d=l,p=ku(h,d),v=ku(d,d),g=p*p-v*(ku(h,h)-1);if(!(g<0)){var y=eu(g),_=Pu(d,(-p-y)/v);if(Cu(_,h),_=Tu(_),!r)return _;var b,m=t[0],x=e[0],w=t[1],M=e[1];x<m&&(b=m,m=x,x=b);var N=x-m,A=Ga(N-Ya)<Oa;if(!A&&M<w&&(b=w,w=M,M=b),A||N<Oa?A?w+M>0^_[1]<(Ga(_[0]-m)<Oa?w:M):w<=_[1]&&_[1]<=M:N>Ya^(m<=_[0]&&_[0]<=x)){var T=Pu(d,(-p+y)/v);return Cu(T,h),[_,Tu(T)]}}}function u(n,e){var i=r?t:Ya-t,o=0;return n<-i?o|=1:n>i&&(o|=2),e<-i?o|=4:e>i&&(o|=8),o}return Xc(o,function(t){var n,e,c,f,s;return{lineStart:function(){f=c=!1,s=1},point:function(l,h){var d,p=[l,h],v=o(l,h),g=r?v?0:u(l,h):v?u(l+(l<0?Ya:-Ya),h):0;if(!n&&(f=c=v)&&t.lineStart(),v!==c&&(!(d=a(n,p))||Bc(n,d)||Bc(p,d))&&(p[0]+=Oa,p[1]+=Oa,v=o(p[0],p[1])),v!==c)s=0,v?(t.lineStart(),d=a(p,n),t.point(d[0],d[1])):(d=a(n,p),t.point(d[0],d[1]),t.lineEnd()),n=d;else if(i&&n&&r^v){var y;g&e||!(y=a(p,n,!0))||(s=0,r?(t.lineStart(),t.point(y[0][0],y[0][1]),t.point(y[1][0],y[1][1]),t.lineEnd()):(t.point(y[1][0],y[1][1]),t.lineEnd(),t.lineStart(),t.point(y[0][0],y[0][1])))}!v||n&&Bc(n,p)||t.point(p[0],p[1]),n=p,c=v,e=g},lineEnd:function(){c&&t.lineEnd(),n=null},clean:function(){return s|(f&&c)<<1}}},function(n,r,i,o){Lc(o,t,e,i,n,r)},r?[0,-t]:[-Ya,t-Ya])}var Zc=1e9,Qc=-Zc;function Jc(t,n,e,r){function i(i,o){return t<=i&&i<=e&&n<=o&&o<=r}function o(i,o,u,f){var s=0,l=0;if(null==i||(s=a(i,u))!==(l=a(o,u))||c(i,o)<0^u>0)do{f.point(0===s||3===s?t:e,s>1?r:n)}while((s=(s+u+4)%4)!==l);else f.point(o[0],o[1])}function a(r,i){return Ga(r[0]-t)<Oa?i>0?0:3:Ga(r[0]-e)<Oa?i>0?2:1:Ga(r[1]-n)<Oa?i>0?1:0:i>0?3:2}function u(t,n){return c(t.x,n.x)}function c(t,n){var e=a(t,1),r=a(n,1);return e!==r?e-r:0===e?n[1]-t[1]:1===e?t[0]-n[0]:2===e?t[1]-n[1]:n[0]-t[0]}return function(a){var c,f,s,l,h,d,p,v,g,y,_,b=a,m=Oc(),x={point:w,lineStart:function(){x.point=M,f&&f.push(s=[]);y=!0,g=!1,p=v=NaN},lineEnd:function(){c&&(M(l,h),d&&g&&m.rejoin(),c.push(m.result()));x.point=w,g&&b.lineEnd()},polygonStart:function(){b=m,c=[],f=[],_=!0},polygonEnd:function(){var n=function(){for(var n=0,e=0,i=f.length;e<i;++e)for(var o,a,u=f[e],c=1,s=u.length,l=u[0],h=l[0],d=l[1];c<s;++c)o=h,a=d,l=u[c],h=l[0],d=l[1],a<=r?d>r&&(h-o)*(r-a)>(d-a)*(t-o)&&++n:d<=r&&(h-o)*(r-a)<(d-a)*(t-o)&&--n;return n}(),e=_&&n,i=(c=T(c)).length;(e||i)&&(a.polygonStart(),e&&(a.lineStart(),o(null,null,1,a),a.lineEnd()),i&&Fc(c,u,n,o,a),a.polygonEnd());b=a,c=f=s=null}};function w(t,n){i(t,n)&&b.point(t,n)}function M(o,a){var u=i(o,a);if(f&&s.push([o,a]),y)l=o,h=a,d=u,y=!1,u&&(b.lineStart(),b.point(o,a));else if(u&&g)b.point(o,a);else{var c=[p=Math.max(Qc,Math.min(Zc,p)),v=Math.max(Qc,Math.min(Zc,v))],m=[o=Math.max(Qc,Math.min(Zc,o)),a=Math.max(Qc,Math.min(Zc,a))];!function(t,n,e,r,i,o){var a,u=t[0],c=t[1],f=0,s=1,l=n[0]-u,h=n[1]-c;if(a=e-u,l||!(a>0)){if(a/=l,l<0){if(a<f)return;a<s&&(s=a)}else if(l>0){if(a>s)return;a>f&&(f=a)}if(a=i-u,l||!(a<0)){if(a/=l,l<0){if(a>s)return;a>f&&(f=a)}else if(l>0){if(a<f)return;a<s&&(s=a)}if(a=r-c,h||!(a>0)){if(a/=h,h<0){if(a<f)return;a<s&&(s=a)}else if(h>0){if(a>s)return;a>f&&(f=a)}if(a=o-c,h||!(a<0)){if(a/=h,h<0){if(a>s)return;a>f&&(f=a)}else if(h>0){if(a<f)return;a<s&&(s=a)}return f>0&&(t[0]=u+f*l,t[1]=c+f*h),s<1&&(n[0]=u+s*l,n[1]=c+s*h),!0}}}}}(c,m,t,n,e,r)?u&&(b.lineStart(),b.point(o,a),_=!1):(g||(b.lineStart(),b.point(c[0],c[1])),b.point(m[0],m[1]),u||b.lineEnd(),_=!1)}p=o,v=a,g=u}return x}}var Kc,tf,nf,ef=Da(),rf={sphere:uu,point:uu,lineStart:function(){rf.point=af,rf.lineEnd=of},lineEnd:uu,polygonStart:uu,polygonEnd:uu};function of(){rf.point=rf.lineEnd=uu}function af(t,n){Kc=t*=Xa,tf=tu(n*=Xa),nf=Wa(n),rf.point=uf}function uf(t,n){t*=Xa;var e=tu(n*=Xa),r=Wa(n),i=Ga(t-Kc),o=Wa(i),a=r*tu(i),u=nf*e-tf*r*o,c=tf*e+nf*r*o;ef.add($a(eu(a*a+u*u),c)),Kc=t,tf=e,nf=r}function cf(t){return ef.reset(),du(t,rf),+ef}var ff=[null,null],sf={type:"LineString",coordinates:ff};function lf(t,n){return ff[0]=t,ff[1]=n,cf(sf)}var hf={Feature:function(t,n){return pf(t.geometry,n)},FeatureCollection:function(t,n){for(var e=t.features,r=-1,i=e.length;++r<i;)if(pf(e[r].geometry,n))return!0;return!1}},df={Sphere:function(){return!0},Point:function(t,n){return vf(t.coordinates,n)},MultiPoint:function(t,n){for(var e=t.coordinates,r=-1,i=e.length;++r<i;)if(vf(e[r],n))return!0;return!1},LineString:function(t,n){return gf(t.coordinates,n)},MultiLineString:function(t,n){for(var e=t.coordinates,r=-1,i=e.length;++r<i;)if(gf(e[r],n))return!0;return!1},Polygon:function(t,n){return yf(t.coordinates,n)},MultiPolygon:function(t,n){for(var e=t.coordinates,r=-1,i=e.length;++r<i;)if(yf(e[r],n))return!0;return!1},GeometryCollection:function(t,n){for(var e=t.geometries,r=-1,i=e.length;++r<i;)if(pf(e[r],n))return!0;return!1}};function pf(t,n){return!(!t||!df.hasOwnProperty(t.type))&&df[t.type](t,n)}function vf(t,n){return 0===lf(t,n)}function gf(t,n){var e=lf(t[0],t[1]);return lf(t[0],n)+lf(n,t[1])<=e+Oa}function yf(t,n){return!!Hc(t.map(_f),bf(n))}function _f(t){return(t=t.map(bf)).pop(),t}function bf(t){return[t[0]*Xa,t[1]*Xa]}function mf(t,n,e){var r=g(t,n-Oa,e).concat(n);return function(t){return r.map(function(n){return[t,n]})}}function xf(t,n,e){var r=g(t,n-Oa,e).concat(n);return function(t){return r.map(function(n){return[n,t]})}}function wf(){var t,n,e,r,i,o,a,u,c,f,s,l,h=10,d=h,p=90,v=360,y=2.5;function _(){return{type:"MultiLineString",coordinates:b()}}function b(){return g(Za(r/p)*p,e,p).map(s).concat(g(Za(u/v)*v,a,v).map(l)).concat(g(Za(n/h)*h,t,h).filter(function(t){return Ga(t%p)>Oa}).map(c)).concat(g(Za(o/d)*d,i,d).filter(function(t){return Ga(t%v)>Oa}).map(f))}return _.lines=function(){return b().map(function(t){return{type:"LineString",coordinates:t}})},_.outline=function(){return{type:"Polygon",coordinates:[s(r).concat(l(a).slice(1),s(e).reverse().slice(1),l(u).reverse().slice(1))]}},_.extent=function(t){return arguments.length?_.extentMajor(t).extentMinor(t):_.extentMinor()},_.extentMajor=function(t){return arguments.length?(r=+t[0][0],e=+t[1][0],u=+t[0][1],a=+t[1][1],r>e&&(t=r,r=e,e=t),u>a&&(t=u,u=a,a=t),_.precision(y)):[[r,u],[e,a]]},_.extentMinor=function(e){return arguments.length?(n=+e[0][0],t=+e[1][0],o=+e[0][1],i=+e[1][1],n>t&&(e=n,n=t,t=e),o>i&&(e=o,o=i,i=e),_.precision(y)):[[n,o],[t,i]]},_.step=function(t){return arguments.length?_.stepMajor(t).stepMinor(t):_.stepMinor()},_.stepMajor=function(t){return arguments.length?(p=+t[0],v=+t[1],_):[p,v]},_.stepMinor=function(t){return arguments.length?(h=+t[0],d=+t[1],_):[h,d]},_.precision=function(h){return arguments.length?(y=+h,c=mf(o,i,90),f=xf(n,t,y),s=mf(u,a,90),l=xf(r,e,y),_):y},_.extentMajor([[-180,-90+Oa],[180,90-Oa]]).extentMinor([[-180,-80-Oa],[180,80+Oa]])}function Mf(t){return t}var Nf,Af,Tf,Sf,kf=Da(),Ef=Da(),Cf={point:uu,lineStart:uu,lineEnd:uu,polygonStart:function(){Cf.lineStart=Pf,Cf.lineEnd=Df},polygonEnd:function(){Cf.lineStart=Cf.lineEnd=Cf.point=uu,kf.add(Ga(Ef)),Ef.reset()},result:function(){var t=kf/2;return kf.reset(),t}};function Pf(){Cf.point=zf}function zf(t,n){Cf.point=Rf,Nf=Tf=t,Af=Sf=n}function Rf(t,n){Ef.add(Sf*t-Tf*n),Tf=t,Sf=n}function Df(){Rf(Nf,Af)}var qf=1/0,Lf=qf,Uf=-qf,Of=Uf,Bf={point:function(t,n){t<qf&&(qf=t);t>Uf&&(Uf=t);n<Lf&&(Lf=n);n>Of&&(Of=n)},lineStart:uu,lineEnd:uu,polygonStart:uu,polygonEnd:uu,result:function(){var t=[[qf,Lf],[Uf,Of]];return Uf=Of=-(Lf=qf=1/0),t}};var Yf,Ff,If,jf,Hf=0,Xf=0,Gf=0,Vf=0,$f=0,Wf=0,Zf=0,Qf=0,Jf=0,Kf={point:ts,lineStart:ns,lineEnd:is,polygonStart:function(){Kf.lineStart=os,Kf.lineEnd=as},polygonEnd:function(){Kf.point=ts,Kf.lineStart=ns,Kf.lineEnd=is},result:function(){var t=Jf?[Zf/Jf,Qf/Jf]:Wf?[Vf/Wf,$f/Wf]:Gf?[Hf/Gf,Xf/Gf]:[NaN,NaN];return Hf=Xf=Gf=Vf=$f=Wf=Zf=Qf=Jf=0,t}};function ts(t,n){Hf+=t,Xf+=n,++Gf}function ns(){Kf.point=es}function es(t,n){Kf.point=rs,ts(If=t,jf=n)}function rs(t,n){var e=t-If,r=n-jf,i=eu(e*e+r*r);Vf+=i*(If+t)/2,$f+=i*(jf+n)/2,Wf+=i,ts(If=t,jf=n)}function is(){Kf.point=ts}function os(){Kf.point=us}function as(){cs(Yf,Ff)}function us(t,n){Kf.point=cs,ts(Yf=If=t,Ff=jf=n)}function cs(t,n){var e=t-If,r=n-jf,i=eu(e*e+r*r);Vf+=i*(If+t)/2,$f+=i*(jf+n)/2,Wf+=i,Zf+=(i=jf*t-If*n)*(If+t),Qf+=i*(jf+n),Jf+=3*i,ts(If=t,jf=n)}function fs(t){this._context=t}fs.prototype={_radius:4.5,pointRadius:function(t){return this._radius=t,this},polygonStart:function(){this._line=0},polygonEnd:function(){this._line=NaN},lineStart:function(){this._point=0},lineEnd:function(){0===this._line&&this._context.closePath(),this._point=NaN},point:function(t,n){switch(this._point){case 0:this._context.moveTo(t,n),this._point=1;break;case 1:this._context.lineTo(t,n);break;default:this._context.moveTo(t+this._radius,n),this._context.arc(t,n,this._radius,0,ja)}},result:uu};var ss,ls,hs,ds,ps,vs=Da(),gs={point:uu,lineStart:function(){gs.point=ys},lineEnd:function(){ss&&_s(ls,hs),gs.point=uu},polygonStart:function(){ss=!0},polygonEnd:function(){ss=null},result:function(){var t=+vs;return vs.reset(),t}};function ys(t,n){gs.point=_s,ls=ds=t,hs=ps=n}function _s(t,n){ds-=t,ps-=n,vs.add(eu(ds*ds+ps*ps)),ds=t,ps=n}function bs(){this._string=[]}function ms(t){return"m0,"+t+"a"+t+","+t+" 0 1,1 0,"+-2*t+"a"+t+","+t+" 0 1,1 0,"+2*t+"z"}function xs(t){return function(n){var e=new ws;for(var r in t)e[r]=t[r];return e.stream=n,e}}function ws(){}function Ms(t,n,e){var r=t.clipExtent&&t.clipExtent();return t.scale(150).translate([0,0]),null!=r&&t.clipExtent(null),du(e,t.stream(Bf)),n(Bf.result()),null!=r&&t.clipExtent(r),t}function Ns(t,n,e){return Ms(t,function(e){var r=n[1][0]-n[0][0],i=n[1][1]-n[0][1],o=Math.min(r/(e[1][0]-e[0][0]),i/(e[1][1]-e[0][1])),a=+n[0][0]+(r-o*(e[1][0]+e[0][0]))/2,u=+n[0][1]+(i-o*(e[1][1]+e[0][1]))/2;t.scale(150*o).translate([a,u])},e)}function As(t,n,e){return Ns(t,[[0,0],n],e)}function Ts(t,n,e){return Ms(t,function(e){var r=+n,i=r/(e[1][0]-e[0][0]),o=(r-i*(e[1][0]+e[0][0]))/2,a=-i*e[0][1];t.scale(150*i).translate([o,a])},e)}function Ss(t,n,e){return Ms(t,function(e){var r=+n,i=r/(e[1][1]-e[0][1]),o=-i*e[0][0],a=(r-i*(e[1][1]+e[0][1]))/2;t.scale(150*i).translate([o,a])},e)}bs.prototype={_radius:4.5,_circle:ms(4.5),pointRadius:function(t){return(t=+t)!==this._radius&&(this._radius=t,this._circle=null),this},polygonStart:function(){this._line=0},polygonEnd:function(){this._line=NaN},lineStart:function(){this._point=0},lineEnd:function(){0===this._line&&this._string.push("Z"),this._point=NaN},point:function(t,n){switch(this._point){case 0:this._string.push("M",t,",",n),this._point=1;break;case 1:this._string.push("L",t,",",n);break;default:null==this._circle&&(this._circle=ms(this._radius)),this._string.push("M",t,",",n,this._circle)}},result:function(){if(this._string.length){var t=this._string.join("");return this._string=[],t}return null}},ws.prototype={constructor:ws,point:function(t,n){this.stream.point(t,n)},sphere:function(){this.stream.sphere()},lineStart:function(){this.stream.lineStart()},lineEnd:function(){this.stream.lineEnd()},polygonStart:function(){this.stream.polygonStart()},polygonEnd:function(){this.stream.polygonEnd()}};var ks=16,Es=Wa(30*Xa);function Cs(t,n){return+n?function(t,n){function e(r,i,o,a,u,c,f,s,l,h,d,p,v,g){var y=f-r,_=s-i,b=y*y+_*_;if(b>4*n&&v--){var m=a+h,x=u+d,w=c+p,M=eu(m*m+x*x+w*w),N=ou(w/=M),A=Ga(Ga(w)-1)<Oa||Ga(o-l)<Oa?(o+l)/2:$a(x,m),T=t(A,N),S=T[0],k=T[1],E=S-r,C=k-i,P=_*E-y*C;(P*P/b>n||Ga((y*E+_*C)/b-.5)>.3||a*h+u*d+c*p<Es)&&(e(r,i,o,a,u,c,S,k,A,m/=M,x/=M,w,v,g),g.point(S,k),e(S,k,A,m,x,w,f,s,l,h,d,p,v,g))}}return function(n){var r,i,o,a,u,c,f,s,l,h,d,p,v={point:g,lineStart:y,lineEnd:b,polygonStart:function(){n.polygonStart(),v.lineStart=m},polygonEnd:function(){n.polygonEnd(),v.lineStart=y}};function g(e,r){e=t(e,r),n.point(e[0],e[1])}function y(){s=NaN,v.point=_,n.lineStart()}function _(r,i){var o=Su([r,i]),a=t(r,i);e(s,l,f,h,d,p,s=a[0],l=a[1],f=r,h=o[0],d=o[1],p=o[2],ks,n),n.point(s,l)}function b(){v.point=g,n.lineEnd()}function m(){y(),v.point=x,v.lineEnd=w}function x(t,n){_(r=t,n),i=s,o=l,a=h,u=d,c=p,v.point=_}function w(){e(s,l,f,h,d,p,i,o,r,a,u,c,ks,n),v.lineEnd=b,b()}return v}}(t,n):function(t){return xs({point:function(n,e){n=t(n,e),this.stream.point(n[0],n[1])}})}(t)}var Ps=xs({point:function(t,n){this.stream.point(t*Xa,n*Xa)}});function zs(t,n,e,r){var i=Wa(r),o=tu(r),a=i*t,u=o*t,c=i/t,f=o/t,s=(o*e-i*n)/t,l=(o*n+i*e)/t;function h(t,r){return[a*t-u*r+n,e-u*t-a*r]}return h.invert=function(t,n){return[c*t-f*n+s,l-f*t-c*n]},h}function Rs(t){return Ds(function(){return t})()}function Ds(t){var n,e,r,i,o,a,u,c,f,s,l=150,h=480,d=250,p=0,v=0,g=0,y=0,_=0,b=0,m=null,x=$c,w=null,M=Mf,N=.5;function A(t){return c(t[0]*Xa,t[1]*Xa)}function T(t){return(t=c.invert(t[0],t[1]))&&[t[0]*Ha,t[1]*Ha]}function S(){var t=zs(l,0,0,b).apply(null,n(p,v)),r=(b?zs:function(t,n,e){function r(r,i){return[n+t*r,e-t*i]}return r.invert=function(r,i){return[(r-n)/t,(e-i)/t]},r})(l,h-t[0],d-t[1],b);return e=Pc(g,y,_),u=Ec(n,r),c=Ec(e,u),a=Cs(u,N),k()}function k(){return f=s=null,A}return A.stream=function(t){return f&&s===t?f:f=Ps(function(t){return xs({point:function(n,e){var r=t(n,e);return this.stream.point(r[0],r[1])}})}(e)(x(a(M(s=t)))))},A.preclip=function(t){return arguments.length?(x=t,m=void 0,k()):x},A.postclip=function(t){return arguments.length?(M=t,w=r=i=o=null,k()):M},A.clipAngle=function(t){return arguments.length?(x=+t?Wc(m=t*Xa):(m=null,$c),k()):m*Ha},A.clipExtent=function(t){return arguments.length?(M=null==t?(w=r=i=o=null,Mf):Jc(w=+t[0][0],r=+t[0][1],i=+t[1][0],o=+t[1][1]),k()):null==w?null:[[w,r],[i,o]]},A.scale=function(t){return arguments.length?(l=+t,S()):l},A.translate=function(t){return arguments.length?(h=+t[0],d=+t[1],S()):[h,d]},A.center=function(t){return arguments.length?(p=t[0]%360*Xa,v=t[1]%360*Xa,S()):[p*Ha,v*Ha]},A.rotate=function(t){return arguments.length?(g=t[0]%360*Xa,y=t[1]%360*Xa,_=t.length>2?t[2]%360*Xa:0,S()):[g*Ha,y*Ha,_*Ha]},A.angle=function(t){return arguments.length?(b=t%360*Xa,S()):b*Ha},A.precision=function(t){return arguments.length?(a=Cs(u,N=t*t),k()):eu(N)},A.fitExtent=function(t,n){return Ns(A,t,n)},A.fitSize=function(t,n){return As(A,t,n)},A.fitWidth=function(t,n){return Ts(A,t,n)},A.fitHeight=function(t,n){return Ss(A,t,n)},function(){return n=t.apply(this,arguments),A.invert=n.invert&&T,S()}}function qs(t){var n=0,e=Ya/3,r=Ds(t),i=r(n,e);return i.parallels=function(t){return arguments.length?r(n=t[0]*Xa,e=t[1]*Xa):[n*Ha,e*Ha]},i}function Ls(t,n){var e=tu(t),r=(e+tu(n))/2;if(Ga(r)<Oa)return function(t){var n=Wa(t);function e(t,e){return[t*n,tu(e)/n]}return e.invert=function(t,e){return[t/n,ou(e*n)]},e}(t);var i=1+e*(2*r-e),o=eu(i)/r;function a(t,n){var e=eu(i-2*r*tu(n))/r;return[e*tu(t*=r),o-e*Wa(t)]}return a.invert=function(t,n){var e=o-n;return[$a(t,Ga(e))/r*nu(e),ou((i-(t*t+e*e)*r*r)/(2*r))]},a}function Us(){return qs(Ls).scale(155.424).center([0,33.6442])}function Os(){return Us().parallels([29.5,45.5]).scale(1070).translate([480,250]).rotate([96,0]).center([-.6,38.7])}function Bs(t){return function(n,e){var r=Wa(n),i=Wa(e),o=t(r*i);return[o*i*tu(n),o*tu(e)]}}function Ys(t){return function(n,e){var r=eu(n*n+e*e),i=t(r),o=tu(i),a=Wa(i);return[$a(n*o,r*a),ou(r&&e*o/r)]}}var Fs=Bs(function(t){return eu(2/(1+t))});Fs.invert=Ys(function(t){return 2*ou(t/2)});var Is=Bs(function(t){return(t=iu(t))&&t/tu(t)});function js(t,n){return[t,Ja(ru((Fa+n)/2))]}function Hs(t){var n,e,r,i=Rs(t),o=i.center,a=i.scale,u=i.translate,c=i.clipExtent,f=null;function s(){var o=Ya*a(),u=i(qc(i.rotate()).invert([0,0]));return c(null==f?[[u[0]-o,u[1]-o],[u[0]+o,u[1]+o]]:t===js?[[Math.max(u[0]-o,f),n],[Math.min(u[0]+o,e),r]]:[[f,Math.max(u[1]-o,n)],[e,Math.min(u[1]+o,r)]])}return i.scale=function(t){return arguments.length?(a(t),s()):a()},i.translate=function(t){return arguments.length?(u(t),s()):u()},i.center=function(t){return arguments.length?(o(t),s()):o()},i.clipExtent=function(t){return arguments.length?(null==t?f=n=e=r=null:(f=+t[0][0],n=+t[0][1],e=+t[1][0],r=+t[1][1]),s()):null==f?null:[[f,n],[e,r]]},s()}function Xs(t){return ru((Fa+t)/2)}function Gs(t,n){var e=Wa(t),r=t===n?tu(t):Ja(e/Wa(n))/Ja(Xs(n)/Xs(t)),i=e*Ka(Xs(t),r)/r;if(!r)return js;function o(t,n){i>0?n<-Fa+Oa&&(n=-Fa+Oa):n>Fa-Oa&&(n=Fa-Oa);var e=i/Ka(Xs(n),r);return[e*tu(r*t),i-e*Wa(r*t)]}return o.invert=function(t,n){var e=i-n,o=nu(r)*eu(t*t+e*e);return[$a(t,Ga(e))/r*nu(e),2*Va(Ka(i/o,1/r))-Fa]},o}function Vs(t,n){return[t,n]}function $s(t,n){var e=Wa(t),r=t===n?tu(t):(e-Wa(n))/(n-t),i=e/r+t;if(Ga(r)<Oa)return Vs;function o(t,n){var e=i-n,o=r*t;return[e*tu(o),i-e*Wa(o)]}return o.invert=function(t,n){var e=i-n;return[$a(t,Ga(e))/r*nu(e),i-nu(r)*eu(t*t+e*e)]},o}Is.invert=Ys(function(t){return t}),js.invert=function(t,n){return[t,2*Va(Qa(n))-Fa]},Vs.invert=Vs;var Ws=1.340264,Zs=-.081106,Qs=893e-6,Js=.003796,Ks=eu(3)/2;function tl(t,n){var e=ou(Ks*tu(n)),r=e*e,i=r*r*r;return[t*Wa(e)/(Ks*(Ws+3*Zs*r+i*(7*Qs+9*Js*r))),e*(Ws+Zs*r+i*(Qs+Js*r))]}function nl(t,n){var e=Wa(n),r=Wa(t)*e;return[e*tu(t)/r,tu(n)/r]}function el(t,n,e,r){return 1===t&&1===n&&0===e&&0===r?Mf:xs({point:function(i,o){this.stream.point(i*t+e,o*n+r)}})}function rl(t,n){var e=n*n,r=e*e;return[t*(.8707-.131979*e+r*(r*(.003971*e-.001529*r)-.013791)),n*(1.007226+e*(.015085+r*(.028874*e-.044475-.005916*r)))]}function il(t,n){return[Wa(n)*tu(t),tu(n)]}function ol(t,n){var e=Wa(n),r=1+Wa(t)*e;return[e*tu(t)/r,tu(n)/r]}function al(t,n){return[Ja(ru((Fa+n)/2)),-t]}function ul(t,n){return t.parent===n.parent?1:2}function cl(t,n){return t+n.x}function fl(t,n){return Math.max(t,n.y)}function sl(t){var n=0,e=t.children,r=e&&e.length;if(r)for(;--r>=0;)n+=e[r].value;else n=1;t.value=n}function ll(t,n){var e,r,i,o,a,u=new vl(t),c=+t.value&&(u.value=t.value),f=[u];for(null==n&&(n=hl);e=f.pop();)if(c&&(e.value=+e.data.value),(i=n(e.data))&&(a=i.length))for(e.children=new Array(a),o=a-1;o>=0;--o)f.push(r=e.children[o]=new vl(i[o])),r.parent=e,r.depth=e.depth+1;return u.eachBefore(pl)}function hl(t){return t.children}function dl(t){t.data=t.data.data}function pl(t){var n=0;do{t.height=n}while((t=t.parent)&&t.height<++n)}function vl(t){this.data=t,this.depth=this.height=0,this.parent=null}tl.invert=function(t,n){for(var e,r=n,i=r*r,o=i*i*i,a=0;a<12&&(o=(i=(r-=e=(r*(Ws+Zs*i+o*(Qs+Js*i))-n)/(Ws+3*Zs*i+o*(7*Qs+9*Js*i)))*r)*i*i,!(Ga(e)<Ba));++a);return[Ks*t*(Ws+3*Zs*i+o*(7*Qs+9*Js*i))/Wa(r),ou(tu(r)/Ks)]},nl.invert=Ys(Va),rl.invert=function(t,n){var e,r=n,i=25;do{var o=r*r,a=o*o;r-=e=(r*(1.007226+o*(.015085+a*(.028874*o-.044475-.005916*a)))-n)/(1.007226+o*(.045255+a*(.259866*o-.311325-.005916*11*a)))}while(Ga(e)>Oa&&--i>0);return[t/(.8707+(o=r*r)*(o*(o*o*o*(.003971-.001529*o)-.013791)-.131979)),r]},il.invert=Ys(ou),ol.invert=Ys(function(t){return 2*Va(t)}),al.invert=function(t,n){return[-n,2*Va(Qa(t))-Fa]},vl.prototype=ll.prototype={constructor:vl,count:function(){return this.eachAfter(sl)},each:function(t){var n,e,r,i,o=this,a=[o];do{for(n=a.reverse(),a=[];o=n.pop();)if(t(o),e=o.children)for(r=0,i=e.length;r<i;++r)a.push(e[r])}while(a.length);return this},eachAfter:function(t){for(var n,e,r,i=this,o=[i],a=[];i=o.pop();)if(a.push(i),n=i.children)for(e=0,r=n.length;e<r;++e)o.push(n[e]);for(;i=a.pop();)t(i);return this},eachBefore:function(t){for(var n,e,r=this,i=[r];r=i.pop();)if(t(r),n=r.children)for(e=n.length-1;e>=0;--e)i.push(n[e]);return this},sum:function(t){return this.eachAfter(function(n){for(var e=+t(n.data)||0,r=n.children,i=r&&r.length;--i>=0;)e+=r[i].value;n.value=e})},sort:function(t){return this.eachBefore(function(n){n.children&&n.children.sort(t)})},path:function(t){for(var n=this,e=function(t,n){if(t===n)return t;var e=t.ancestors(),r=n.ancestors(),i=null;for(t=e.pop(),n=r.pop();t===n;)i=t,t=e.pop(),n=r.pop();return i}(n,t),r=[n];n!==e;)n=n.parent,r.push(n);for(var i=r.length;t!==e;)r.splice(i,0,t),t=t.parent;return r},ancestors:function(){for(var t=this,n=[t];t=t.parent;)n.push(t);return n},descendants:function(){var t=[];return this.each(function(n){t.push(n)}),t},leaves:function(){var t=[];return this.eachBefore(function(n){n.children||t.push(n)}),t},links:function(){var t=this,n=[];return t.each(function(e){e!==t&&n.push({source:e.parent,target:e})}),n},copy:function(){return ll(this).eachBefore(dl)}};var gl=Array.prototype.slice;function yl(t){for(var n,e,r=0,i=(t=function(t){for(var n,e,r=t.length;r;)e=Math.random()*r--|0,n=t[r],t[r]=t[e],t[e]=n;return t}(gl.call(t))).length,o=[];r<i;)n=t[r],e&&ml(e,n)?++r:(e=wl(o=_l(o,n)),r=0);return e}function _l(t,n){var e,r;if(xl(n,t))return[n];for(e=0;e<t.length;++e)if(bl(n,t[e])&&xl(Ml(t[e],n),t))return[t[e],n];for(e=0;e<t.length-1;++e)for(r=e+1;r<t.length;++r)if(bl(Ml(t[e],t[r]),n)&&bl(Ml(t[e],n),t[r])&&bl(Ml(t[r],n),t[e])&&xl(Nl(t[e],t[r],n),t))return[t[e],t[r],n];throw new Error}function bl(t,n){var e=t.r-n.r,r=n.x-t.x,i=n.y-t.y;return e<0||e*e<r*r+i*i}function ml(t,n){var e=t.r-n.r+1e-6,r=n.x-t.x,i=n.y-t.y;return e>0&&e*e>r*r+i*i}function xl(t,n){for(var e=0;e<n.length;++e)if(!ml(t,n[e]))return!1;return!0}function wl(t){switch(t.length){case 1:return{x:(n=t[0]).x,y:n.y,r:n.r};case 2:return Ml(t[0],t[1]);case 3:return Nl(t[0],t[1],t[2])}var n}function Ml(t,n){var e=t.x,r=t.y,i=t.r,o=n.x,a=n.y,u=n.r,c=o-e,f=a-r,s=u-i,l=Math.sqrt(c*c+f*f);return{x:(e+o+c/l*s)/2,y:(r+a+f/l*s)/2,r:(l+i+u)/2}}function Nl(t,n,e){var r=t.x,i=t.y,o=t.r,a=n.x,u=n.y,c=n.r,f=e.x,s=e.y,l=e.r,h=r-a,d=r-f,p=i-u,v=i-s,g=c-o,y=l-o,_=r*r+i*i-o*o,b=_-a*a-u*u+c*c,m=_-f*f-s*s+l*l,x=d*p-h*v,w=(p*m-v*b)/(2*x)-r,M=(v*g-p*y)/x,N=(d*b-h*m)/(2*x)-i,A=(h*y-d*g)/x,T=M*M+A*A-1,S=2*(o+w*M+N*A),k=w*w+N*N-o*o,E=-(T?(S+Math.sqrt(S*S-4*T*k))/(2*T):k/S);return{x:r+w+M*E,y:i+N+A*E,r:E}}function Al(t,n,e){var r,i,o,a,u=t.x-n.x,c=t.y-n.y,f=u*u+c*c;f?(i=n.r+e.r,i*=i,a=t.r+e.r,i>(a*=a)?(r=(f+a-i)/(2*f),o=Math.sqrt(Math.max(0,a/f-r*r)),e.x=t.x-r*u-o*c,e.y=t.y-r*c+o*u):(r=(f+i-a)/(2*f),o=Math.sqrt(Math.max(0,i/f-r*r)),e.x=n.x+r*u-o*c,e.y=n.y+r*c+o*u)):(e.x=n.x+e.r,e.y=n.y)}function Tl(t,n){var e=t.r+n.r-1e-6,r=n.x-t.x,i=n.y-t.y;return e>0&&e*e>r*r+i*i}function Sl(t){var n=t._,e=t.next._,r=n.r+e.r,i=(n.x*e.r+e.x*n.r)/r,o=(n.y*e.r+e.y*n.r)/r;return i*i+o*o}function kl(t){this._=t,this.next=null,this.previous=null}function El(t){if(!(i=t.length))return 0;var n,e,r,i,o,a,u,c,f,s,l;if((n=t[0]).x=0,n.y=0,!(i>1))return n.r;if(e=t[1],n.x=-e.r,e.x=n.r,e.y=0,!(i>2))return n.r+e.r;Al(e,n,r=t[2]),n=new kl(n),e=new kl(e),r=new kl(r),n.next=r.previous=e,e.next=n.previous=r,r.next=e.previous=n;t:for(u=3;u<i;++u){Al(n._,e._,r=t[u]),r=new kl(r),c=e.next,f=n.previous,s=e._.r,l=n._.r;do{if(s<=l){if(Tl(c._,r._)){e=c,n.next=e,e.previous=n,--u;continue t}s+=c._.r,c=c.next}else{if(Tl(f._,r._)){(n=f).next=e,e.previous=n,--u;continue t}l+=f._.r,f=f.previous}}while(c!==f.next);for(r.previous=n,r.next=e,n.next=e.previous=e=r,o=Sl(n);(r=r.next)!==e;)(a=Sl(r))<o&&(n=r,o=a);e=n.next}for(n=[e._],r=e;(r=r.next)!==e;)n.push(r._);for(r=yl(n),u=0;u<i;++u)(n=t[u]).x-=r.x,n.y-=r.y;return r.r}function Cl(t){if("function"!=typeof t)throw new Error;return t}function Pl(){return 0}function zl(t){return function(){return t}}function Rl(t){return Math.sqrt(t.value)}function Dl(t){return function(n){n.children||(n.r=Math.max(0,+t(n)||0))}}function ql(t,n){return function(e){if(r=e.children){var r,i,o,a=r.length,u=t(e)*n||0;if(u)for(i=0;i<a;++i)r[i].r+=u;if(o=El(r),u)for(i=0;i<a;++i)r[i].r-=u;e.r=o+u}}}function Ll(t){return function(n){var e=n.parent;n.r*=t,e&&(n.x=e.x+t*n.x,n.y=e.y+t*n.y)}}function Ul(t){t.x0=Math.round(t.x0),t.y0=Math.round(t.y0),t.x1=Math.round(t.x1),t.y1=Math.round(t.y1)}function Ol(t,n,e,r,i){for(var o,a=t.children,u=-1,c=a.length,f=t.value&&(r-n)/t.value;++u<c;)(o=a[u]).y0=e,o.y1=i,o.x0=n,o.x1=n+=o.value*f}var Bl="$",Yl={depth:-1},Fl={};function Il(t){return t.id}function jl(t){return t.parentId}function Hl(t,n){return t.parent===n.parent?1:2}function Xl(t){var n=t.children;return n?n[0]:t.t}function Gl(t){var n=t.children;return n?n[n.length-1]:t.t}function Vl(t,n,e){var r=e/(n.i-t.i);n.c-=r,n.s+=e,t.c+=r,n.z+=e,n.m+=e}function $l(t,n,e){return t.a.parent===n.parent?t.a:e}function Wl(t,n){this._=t,this.parent=null,this.children=null,this.A=null,this.a=this,this.z=0,this.m=0,this.c=0,this.s=0,this.t=null,this.i=n}function Zl(t,n,e,r,i){for(var o,a=t.children,u=-1,c=a.length,f=t.value&&(i-e)/t.value;++u<c;)(o=a[u]).x0=n,o.x1=r,o.y0=e,o.y1=e+=o.value*f}Wl.prototype=Object.create(vl.prototype);var Ql=(1+Math.sqrt(5))/2;function Jl(t,n,e,r,i,o){for(var a,u,c,f,s,l,h,d,p,v,g,y=[],_=n.children,b=0,m=0,x=_.length,w=n.value;b<x;){c=i-e,f=o-r;do{s=_[m++].value}while(!s&&m<x);for(l=h=s,g=s*s*(v=Math.max(f/c,c/f)/(w*t)),p=Math.max(h/g,g/l);m<x;++m){if(s+=u=_[m].value,u<l&&(l=u),u>h&&(h=u),g=s*s*v,(d=Math.max(h/g,g/l))>p){s-=u;break}p=d}y.push(a={value:s,dice:c<f,children:_.slice(b,m)}),a.dice?Ol(a,e,r,i,w?r+=f*s/w:o):Zl(a,e,r,w?e+=c*s/w:i,o),w-=s,b=m}return y}var Kl=function t(n){function e(t,e,r,i,o){Jl(n,t,e,r,i,o)}return e.ratio=function(n){return t((n=+n)>1?n:1)},e}(Ql);var th=function t(n){function e(t,e,r,i,o){if((a=t._squarify)&&a.ratio===n)for(var a,u,c,f,s,l=-1,h=a.length,d=t.value;++l<h;){for(c=(u=a[l]).children,f=u.value=0,s=c.length;f<s;++f)u.value+=c[f].value;u.dice?Ol(u,e,r,i,r+=(o-r)*u.value/d):Zl(u,e,r,e+=(i-e)*u.value/d,o),d-=u.value}else t._squarify=a=Jl(n,t,e,r,i,o),a.ratio=n}return e.ratio=function(n){return t((n=+n)>1?n:1)},e}(Ql);function nh(t,n){return t[0]-n[0]||t[1]-n[1]}function eh(t){for(var n,e,r,i=t.length,o=[0,1],a=2,u=2;u<i;++u){for(;a>1&&(n=t[o[a-2]],e=t[o[a-1]],r=t[u],(e[0]-n[0])*(r[1]-n[1])-(e[1]-n[1])*(r[0]-n[0])<=0);)--a;o[a++]=u}return o.slice(0,a)}function rh(){return Math.random()}var ih=function t(n){function e(t,e){return t=null==t?0:+t,e=null==e?1:+e,1===arguments.length?(e=t,t=0):e-=t,function(){return n()*e+t}}return e.source=t,e}(rh),oh=function t(n){function e(t,e){var r,i;return t=null==t?0:+t,e=null==e?1:+e,function(){var o;if(null!=r)o=r,r=null;else do{r=2*n()-1,o=2*n()-1,i=r*r+o*o}while(!i||i>1);return t+e*o*Math.sqrt(-2*Math.log(i)/i)}}return e.source=t,e}(rh),ah=function t(n){function e(){var t=oh.source(n).apply(this,arguments);return function(){return Math.exp(t())}}return e.source=t,e}(rh),uh=function t(n){function e(t){return function(){for(var e=0,r=0;r<t;++r)e+=n();return e}}return e.source=t,e}(rh),ch=function t(n){function e(t){var e=uh.source(n)(t);return function(){return e()/t}}return e.source=t,e}(rh),fh=function t(n){function e(t){return function(){return-Math.log(1-n())/t}}return e.source=t,e}(rh);function sh(t,n){switch(arguments.length){case 0:break;case 1:this.range(t);break;default:this.range(n).domain(t)}return this}function lh(t,n){switch(arguments.length){case 0:break;case 1:this.interpolator(t);break;default:this.interpolator(n).domain(t)}return this}var hh=Array.prototype,dh=hh.map,ph=hh.slice,vh={name:"implicit"};function gh(){var t=Qi(),n=[],e=[],r=vh;function i(i){var o=i+"",a=t.get(o);if(!a){if(r!==vh)return r;t.set(o,a=n.push(i))}return e[(a-1)%e.length]}return i.domain=function(e){if(!arguments.length)return n.slice();n=[],t=Qi();for(var r,o,a=-1,u=e.length;++a<u;)t.has(o=(r=e[a])+"")||t.set(o,n.push(r));return i},i.range=function(t){return arguments.length?(e=ph.call(t),i):e.slice()},i.unknown=function(t){return arguments.length?(r=t,i):r},i.copy=function(){return gh(n,e).unknown(r)},sh.apply(i,arguments),i}function yh(){var t,n,e=gh().unknown(void 0),r=e.domain,i=e.range,o=[0,1],a=!1,u=0,c=0,f=.5;function s(){var e=r().length,s=o[1]<o[0],l=o[s-0],h=o[1-s];t=(h-l)/Math.max(1,e-u+2*c),a&&(t=Math.floor(t)),l+=(h-l-t*(e-u))*f,n=t*(1-u),a&&(l=Math.round(l),n=Math.round(n));var d=g(e).map(function(n){return l+t*n});return i(s?d.reverse():d)}return delete e.unknown,e.domain=function(t){return arguments.length?(r(t),s()):r()},e.range=function(t){return arguments.length?(o=[+t[0],+t[1]],s()):o.slice()},e.rangeRound=function(t){return o=[+t[0],+t[1]],a=!0,s()},e.bandwidth=function(){return n},e.step=function(){return t},e.round=function(t){return arguments.length?(a=!!t,s()):a},e.padding=function(t){return arguments.length?(u=Math.min(1,c=+t),s()):u},e.paddingInner=function(t){return arguments.length?(u=Math.min(1,t),s()):u},e.paddingOuter=function(t){return arguments.length?(c=+t,s()):c},e.align=function(t){return arguments.length?(f=Math.max(0,Math.min(1,t)),s()):f},e.copy=function(){return yh(r(),o).round(a).paddingInner(u).paddingOuter(c).align(f)},sh.apply(s(),arguments)}function _h(t){return+t}var bh=[0,1];function mh(t){return t}function xh(t,n){return(n-=t=+t)?function(e){return(e-t)/n}:(e=isNaN(n)?NaN:.5,function(){return e});var e}function wh(t){var n,e=t[0],r=t[t.length-1];return e>r&&(n=e,e=r,r=n),function(t){return Math.max(e,Math.min(r,t))}}function Mh(t,n,e){var r=t[0],i=t[1],o=n[0],a=n[1];return i<r?(r=xh(i,r),o=e(a,o)):(r=xh(r,i),o=e(o,a)),function(t){return o(r(t))}}function Nh(t,n,e){var r=Math.min(t.length,n.length)-1,o=new Array(r),a=new Array(r),u=-1;for(t[r]<t[0]&&(t=t.slice().reverse(),n=n.slice().reverse());++u<r;)o[u]=xh(t[u],t[u+1]),a[u]=e(n[u],n[u+1]);return function(n){var e=i(t,n,1,r)-1;return a[e](o[e](n))}}function Ah(t,n){return n.domain(t.domain()).range(t.range()).interpolate(t.interpolate()).clamp(t.clamp()).unknown(t.unknown())}function Th(){var t,n,e,r,i,o,a=bh,u=bh,c=ye,f=mh;function s(){return r=Math.min(a.length,u.length)>2?Nh:Mh,i=o=null,l}function l(n){return isNaN(n=+n)?e:(i||(i=r(a.map(t),u,c)))(t(f(n)))}return l.invert=function(e){return f(n((o||(o=r(u,a.map(t),he)))(e)))},l.domain=function(t){return arguments.length?(a=dh.call(t,_h),f===mh||(f=wh(a)),s()):a.slice()},l.range=function(t){return arguments.length?(u=ph.call(t),s()):u.slice()},l.rangeRound=function(t){return u=ph.call(t),c=_e,s()},l.clamp=function(t){return arguments.length?(f=t?wh(a):mh,l):f!==mh},l.interpolate=function(t){return arguments.length?(c=t,s()):c},l.unknown=function(t){return arguments.length?(e=t,l):e},function(e,r){return t=e,n=r,s()}}function Sh(t,n){return Th()(t,n)}function kh(n,e,r,i){var o,a=w(n,e,r);switch((i=wa(null==i?",f":i)).type){case"s":var u=Math.max(Math.abs(n),Math.abs(e));return null!=i.precision||isNaN(o=za(a,u))||(i.precision=o),t.formatPrefix(i,u);case"":case"e":case"g":case"p":case"r":null!=i.precision||isNaN(o=Ra(a,Math.max(Math.abs(n),Math.abs(e))))||(i.precision=o-("e"===i.type));break;case"f":case"%":null!=i.precision||isNaN(o=Pa(a))||(i.precision=o-2*("%"===i.type))}return t.format(i)}function Eh(t){var n=t.domain;return t.ticks=function(t){var e=n();return m(e[0],e[e.length-1],null==t?10:t)},t.tickFormat=function(t,e){var r=n();return kh(r[0],r[r.length-1],null==t?10:t,e)},t.nice=function(e){null==e&&(e=10);var r,i=n(),o=0,a=i.length-1,u=i[o],c=i[a];return c<u&&(r=u,u=c,c=r,r=o,o=a,a=r),(r=x(u,c,e))>0?r=x(u=Math.floor(u/r)*r,c=Math.ceil(c/r)*r,e):r<0&&(r=x(u=Math.ceil(u*r)/r,c=Math.floor(c*r)/r,e)),r>0?(i[o]=Math.floor(u/r)*r,i[a]=Math.ceil(c/r)*r,n(i)):r<0&&(i[o]=Math.ceil(u*r)/r,i[a]=Math.floor(c*r)/r,n(i)),t},t}function Ch(t,n){var e,r=0,i=(t=t.slice()).length-1,o=t[r],a=t[i];return a<o&&(e=r,r=i,i=e,e=o,o=a,a=e),t[r]=n.floor(o),t[i]=n.ceil(a),t}function Ph(t){return Math.log(t)}function zh(t){return Math.exp(t)}function Rh(t){return-Math.log(-t)}function Dh(t){return-Math.exp(-t)}function qh(t){return isFinite(t)?+("1e"+t):t<0?0:t}function Lh(t){return function(n){return-t(-n)}}function Uh(n){var e,r,i=n(Ph,zh),o=i.domain,a=10;function u(){return e=function(t){return t===Math.E?Math.log:10===t&&Math.log10||2===t&&Math.log2||(t=Math.log(t),function(n){return Math.log(n)/t})}(a),r=function(t){return 10===t?qh:t===Math.E?Math.exp:function(n){return Math.pow(t,n)}}(a),o()[0]<0?(e=Lh(e),r=Lh(r),n(Rh,Dh)):n(Ph,zh),i}return i.base=function(t){return arguments.length?(a=+t,u()):a},i.domain=function(t){return arguments.length?(o(t),u()):o()},i.ticks=function(t){var n,i=o(),u=i[0],c=i[i.length-1];(n=c<u)&&(h=u,u=c,c=h);var f,s,l,h=e(u),d=e(c),p=null==t?10:+t,v=[];if(!(a%1)&&d-h<p){if(h=Math.round(h)-1,d=Math.round(d)+1,u>0){for(;h<d;++h)for(s=1,f=r(h);s<a;++s)if(!((l=f*s)<u)){if(l>c)break;v.push(l)}}else for(;h<d;++h)for(s=a-1,f=r(h);s>=1;--s)if(!((l=f*s)<u)){if(l>c)break;v.push(l)}}else v=m(h,d,Math.min(d-h,p)).map(r);return n?v.reverse():v},i.tickFormat=function(n,o){if(null==o&&(o=10===a?".0e":","),"function"!=typeof o&&(o=t.format(o)),n===1/0)return o;null==n&&(n=10);var u=Math.max(1,a*n/i.ticks().length);return function(t){var n=t/r(Math.round(e(t)));return n*a<a-.5&&(n*=a),n<=u?o(t):""}},i.nice=function(){return o(Ch(o(),{floor:function(t){return r(Math.floor(e(t)))},ceil:function(t){return r(Math.ceil(e(t)))}}))},i}function Oh(t){return function(n){return Math.sign(n)*Math.log1p(Math.abs(n/t))}}function Bh(t){return function(n){return Math.sign(n)*Math.expm1(Math.abs(n))*t}}function Yh(t){var n=1,e=t(Oh(n),Bh(n));return e.constant=function(e){return arguments.length?t(Oh(n=+e),Bh(n)):n},Eh(e)}function Fh(t){return function(n){return n<0?-Math.pow(-n,t):Math.pow(n,t)}}function Ih(t){return t<0?-Math.sqrt(-t):Math.sqrt(t)}function jh(t){return t<0?-t*t:t*t}function Hh(t){var n=t(mh,mh),e=1;return n.exponent=function(n){return arguments.length?1===(e=+n)?t(mh,mh):.5===e?t(Ih,jh):t(Fh(e),Fh(1/e)):e},Eh(n)}function Xh(){var t=Hh(Th());return t.copy=function(){return Ah(t,Xh()).exponent(t.exponent())},sh.apply(t,arguments),t}var Gh=new Date,Vh=new Date;function $h(t,n,e,r){function i(n){return t(n=new Date(+n)),n}return i.floor=i,i.ceil=function(e){return t(e=new Date(e-1)),n(e,1),t(e),e},i.round=function(t){var n=i(t),e=i.ceil(t);return t-n<e-t?n:e},i.offset=function(t,e){return n(t=new Date(+t),null==e?1:Math.floor(e)),t},i.range=function(e,r,o){var a,u=[];if(e=i.ceil(e),o=null==o?1:Math.floor(o),!(e<r&&o>0))return u;do{u.push(a=new Date(+e)),n(e,o),t(e)}while(a<e&&e<r);return u},i.filter=function(e){return $h(function(n){if(n>=n)for(;t(n),!e(n);)n.setTime(n-1)},function(t,r){if(t>=t)if(r<0)for(;++r<=0;)for(;n(t,-1),!e(t););else for(;--r>=0;)for(;n(t,1),!e(t););})},e&&(i.count=function(n,r){return Gh.setTime(+n),Vh.setTime(+r),t(Gh),t(Vh),Math.floor(e(Gh,Vh))},i.every=function(t){return t=Math.floor(t),isFinite(t)&&t>0?t>1?i.filter(r?function(n){return r(n)%t==0}:function(n){return i.count(0,n)%t==0}):i:null}),i}var Wh=$h(function(){},function(t,n){t.setTime(+t+n)},function(t,n){return n-t});Wh.every=function(t){return t=Math.floor(t),isFinite(t)&&t>0?t>1?$h(function(n){n.setTime(Math.floor(n/t)*t)},function(n,e){n.setTime(+n+e*t)},function(n,e){return(e-n)/t}):Wh:null};var Zh=Wh.range,Qh=6e4,Jh=6048e5,Kh=$h(function(t){t.setTime(t-t.getMilliseconds())},function(t,n){t.setTime(+t+1e3*n)},function(t,n){return(n-t)/1e3},function(t){return t.getUTCSeconds()}),td=Kh.range,nd=$h(function(t){t.setTime(t-t.getMilliseconds()-1e3*t.getSeconds())},function(t,n){t.setTime(+t+n*Qh)},function(t,n){return(n-t)/Qh},function(t){return t.getMinutes()}),ed=nd.range,rd=$h(function(t){t.setTime(t-t.getMilliseconds()-1e3*t.getSeconds()-t.getMinutes()*Qh)},function(t,n){t.setTime(+t+36e5*n)},function(t,n){return(n-t)/36e5},function(t){return t.getHours()}),id=rd.range,od=$h(function(t){t.setHours(0,0,0,0)},function(t,n){t.setDate(t.getDate()+n)},function(t,n){return(n-t-(n.getTimezoneOffset()-t.getTimezoneOffset())*Qh)/864e5},function(t){return t.getDate()-1}),ad=od.range;function ud(t){return $h(function(n){n.setDate(n.getDate()-(n.getDay()+7-t)%7),n.setHours(0,0,0,0)},function(t,n){t.setDate(t.getDate()+7*n)},function(t,n){return(n-t-(n.getTimezoneOffset()-t.getTimezoneOffset())*Qh)/Jh})}var cd=ud(0),fd=ud(1),sd=ud(2),ld=ud(3),hd=ud(4),dd=ud(5),pd=ud(6),vd=cd.range,gd=fd.range,yd=sd.range,_d=ld.range,bd=hd.range,md=dd.range,xd=pd.range,wd=$h(function(t){t.setDate(1),t.setHours(0,0,0,0)},function(t,n){t.setMonth(t.getMonth()+n)},function(t,n){return n.getMonth()-t.getMonth()+12*(n.getFullYear()-t.getFullYear())},function(t){return t.getMonth()}),Md=wd.range,Nd=$h(function(t){t.setMonth(0,1),t.setHours(0,0,0,0)},function(t,n){t.setFullYear(t.getFullYear()+n)},function(t,n){return n.getFullYear()-t.getFullYear()},function(t){return t.getFullYear()});Nd.every=function(t){return isFinite(t=Math.floor(t))&&t>0?$h(function(n){n.setFullYear(Math.floor(n.getFullYear()/t)*t),n.setMonth(0,1),n.setHours(0,0,0,0)},function(n,e){n.setFullYear(n.getFullYear()+e*t)}):null};var Ad=Nd.range,Td=$h(function(t){t.setUTCSeconds(0,0)},function(t,n){t.setTime(+t+n*Qh)},function(t,n){return(n-t)/Qh},function(t){return t.getUTCMinutes()}),Sd=Td.range,kd=$h(function(t){t.setUTCMinutes(0,0,0)},function(t,n){t.setTime(+t+36e5*n)},function(t,n){return(n-t)/36e5},function(t){return t.getUTCHours()}),Ed=kd.range,Cd=$h(function(t){t.setUTCHours(0,0,0,0)},function(t,n){t.setUTCDate(t.getUTCDate()+n)},function(t,n){return(n-t)/864e5},function(t){return t.getUTCDate()-1}),Pd=Cd.range;function zd(t){return $h(function(n){n.setUTCDate(n.getUTCDate()-(n.getUTCDay()+7-t)%7),n.setUTCHours(0,0,0,0)},function(t,n){t.setUTCDate(t.getUTCDate()+7*n)},function(t,n){return(n-t)/Jh})}var Rd=zd(0),Dd=zd(1),qd=zd(2),Ld=zd(3),Ud=zd(4),Od=zd(5),Bd=zd(6),Yd=Rd.range,Fd=Dd.range,Id=qd.range,jd=Ld.range,Hd=Ud.range,Xd=Od.range,Gd=Bd.range,Vd=$h(function(t){t.setUTCDate(1),t.setUTCHours(0,0,0,0)},function(t,n){t.setUTCMonth(t.getUTCMonth()+n)},function(t,n){return n.getUTCMonth()-t.getUTCMonth()+12*(n.getUTCFullYear()-t.getUTCFullYear())},function(t){return t.getUTCMonth()}),$d=Vd.range,Wd=$h(function(t){t.setUTCMonth(0,1),t.setUTCHours(0,0,0,0)},function(t,n){t.setUTCFullYear(t.getUTCFullYear()+n)},function(t,n){return n.getUTCFullYear()-t.getUTCFullYear()},function(t){return t.getUTCFullYear()});Wd.every=function(t){return isFinite(t=Math.floor(t))&&t>0?$h(function(n){n.setUTCFullYear(Math.floor(n.getUTCFullYear()/t)*t),n.setUTCMonth(0,1),n.setUTCHours(0,0,0,0)},function(n,e){n.setUTCFullYear(n.getUTCFullYear()+e*t)}):null};var Zd=Wd.range;function Qd(t){if(0<=t.y&&t.y<100){var n=new Date(-1,t.m,t.d,t.H,t.M,t.S,t.L);return n.setFullYear(t.y),n}return new Date(t.y,t.m,t.d,t.H,t.M,t.S,t.L)}function Jd(t){if(0<=t.y&&t.y<100){var n=new Date(Date.UTC(-1,t.m,t.d,t.H,t.M,t.S,t.L));return n.setUTCFullYear(t.y),n}return new Date(Date.UTC(t.y,t.m,t.d,t.H,t.M,t.S,t.L))}function Kd(t){return{y:t,m:0,d:1,H:0,M:0,S:0,L:0}}function tp(t){var n=t.dateTime,e=t.date,r=t.time,i=t.periods,o=t.days,a=t.shortDays,u=t.months,c=t.shortMonths,f=cp(i),s=fp(i),l=cp(o),h=fp(o),d=cp(a),p=fp(a),v=cp(u),g=fp(u),y=cp(c),_=fp(c),b={a:function(t){return a[t.getDay()]},A:function(t){return o[t.getDay()]},b:function(t){return c[t.getMonth()]},B:function(t){return u[t.getMonth()]},c:null,d:Ep,e:Ep,f:Dp,H:Cp,I:Pp,j:zp,L:Rp,m:qp,M:Lp,p:function(t){return i[+(t.getHours()>=12)]},Q:sv,s:lv,S:Up,u:Op,U:Bp,V:Yp,w:Fp,W:Ip,x:null,X:null,y:jp,Y:Hp,Z:Xp,"%":fv},m={a:function(t){return a[t.getUTCDay()]},A:function(t){return o[t.getUTCDay()]},b:function(t){return c[t.getUTCMonth()]},B:function(t){return u[t.getUTCMonth()]},c:null,d:Gp,e:Gp,f:Qp,H:Vp,I:$p,j:Wp,L:Zp,m:Jp,M:Kp,p:function(t){return i[+(t.getUTCHours()>=12)]},Q:sv,s:lv,S:tv,u:nv,U:ev,V:rv,w:iv,W:ov,x:null,X:null,y:av,Y:uv,Z:cv,"%":fv},x={a:function(t,n,e){var r=d.exec(n.slice(e));return r?(t.w=p[r[0].toLowerCase()],e+r[0].length):-1},A:function(t,n,e){var r=l.exec(n.slice(e));return r?(t.w=h[r[0].toLowerCase()],e+r[0].length):-1},b:function(t,n,e){var r=y.exec(n.slice(e));return r?(t.m=_[r[0].toLowerCase()],e+r[0].length):-1},B:function(t,n,e){var r=v.exec(n.slice(e));return r?(t.m=g[r[0].toLowerCase()],e+r[0].length):-1},c:function(t,e,r){return N(t,n,e,r)},d:bp,e:bp,f:Ap,H:xp,I:xp,j:mp,L:Np,m:_p,M:wp,p:function(t,n,e){var r=f.exec(n.slice(e));return r?(t.p=s[r[0].toLowerCase()],e+r[0].length):-1},Q:Sp,s:kp,S:Mp,u:lp,U:hp,V:dp,w:sp,W:pp,x:function(t,n,r){return N(t,e,n,r)},X:function(t,n,e){return N(t,r,n,e)},y:gp,Y:vp,Z:yp,"%":Tp};function w(t,n){return function(e){var r,i,o,a=[],u=-1,c=0,f=t.length;for(e instanceof Date||(e=new Date(+e));++u<f;)37===t.charCodeAt(u)&&(a.push(t.slice(c,u)),null!=(i=ep[r=t.charAt(++u)])?r=t.charAt(++u):i="e"===r?" ":"0",(o=n[r])&&(r=o(e,i)),a.push(r),c=u+1);return a.push(t.slice(c,u)),a.join("")}}function M(t,n){return function(e){var r,i,o=Kd(1900);if(N(o,t,e+="",0)!=e.length)return null;if("Q"in o)return new Date(o.Q);if("p"in o&&(o.H=o.H%12+12*o.p),"V"in o){if(o.V<1||o.V>53)return null;"w"in o||(o.w=1),"Z"in o?(i=(r=Jd(Kd(o.y))).getUTCDay(),r=i>4||0===i?Dd.ceil(r):Dd(r),r=Cd.offset(r,7*(o.V-1)),o.y=r.getUTCFullYear(),o.m=r.getUTCMonth(),o.d=r.getUTCDate()+(o.w+6)%7):(i=(r=n(Kd(o.y))).getDay(),r=i>4||0===i?fd.ceil(r):fd(r),r=od.offset(r,7*(o.V-1)),o.y=r.getFullYear(),o.m=r.getMonth(),o.d=r.getDate()+(o.w+6)%7)}else("W"in o||"U"in o)&&("w"in o||(o.w="u"in o?o.u%7:"W"in o?1:0),i="Z"in o?Jd(Kd(o.y)).getUTCDay():n(Kd(o.y)).getDay(),o.m=0,o.d="W"in o?(o.w+6)%7+7*o.W-(i+5)%7:o.w+7*o.U-(i+6)%7);return"Z"in o?(o.H+=o.Z/100|0,o.M+=o.Z%100,Jd(o)):n(o)}}function N(t,n,e,r){for(var i,o,a=0,u=n.length,c=e.length;a<u;){if(r>=c)return-1;if(37===(i=n.charCodeAt(a++))){if(i=n.charAt(a++),!(o=x[i in ep?n.charAt(a++):i])||(r=o(t,e,r))<0)return-1}else if(i!=e.charCodeAt(r++))return-1}return r}return b.x=w(e,b),b.X=w(r,b),b.c=w(n,b),m.x=w(e,m),m.X=w(r,m),m.c=w(n,m),{format:function(t){var n=w(t+="",b);return n.toString=function(){return t},n},parse:function(t){var n=M(t+="",Qd);return n.toString=function(){return t},n},utcFormat:function(t){var n=w(t+="",m);return n.toString=function(){return t},n},utcParse:function(t){var n=M(t,Jd);return n.toString=function(){return t},n}}}var np,ep={"-":"",_:" ",0:"0"},rp=/^\s*\d+/,ip=/^%/,op=/[\\^$*+?|[\]().{}]/g;function ap(t,n,e){var r=t<0?"-":"",i=(r?-t:t)+"",o=i.length;return r+(o<e?new Array(e-o+1).join(n)+i:i)}function up(t){return t.replace(op,"\\$&")}function cp(t){return new RegExp("^(?:"+t.map(up).join("|")+")","i")}function fp(t){for(var n={},e=-1,r=t.length;++e<r;)n[t[e].toLowerCase()]=e;return n}function sp(t,n,e){var r=rp.exec(n.slice(e,e+1));return r?(t.w=+r[0],e+r[0].length):-1}function lp(t,n,e){var r=rp.exec(n.slice(e,e+1));return r?(t.u=+r[0],e+r[0].length):-1}function hp(t,n,e){var r=rp.exec(n.slice(e,e+2));return r?(t.U=+r[0],e+r[0].length):-1}function dp(t,n,e){var r=rp.exec(n.slice(e,e+2));return r?(t.V=+r[0],e+r[0].length):-1}function pp(t,n,e){var r=rp.exec(n.slice(e,e+2));return r?(t.W=+r[0],e+r[0].length):-1}function vp(t,n,e){var r=rp.exec(n.slice(e,e+4));return r?(t.y=+r[0],e+r[0].length):-1}function gp(t,n,e){var r=rp.exec(n.slice(e,e+2));return r?(t.y=+r[0]+(+r[0]>68?1900:2e3),e+r[0].length):-1}function yp(t,n,e){var r=/^(Z)|([+-]\d\d)(?::?(\d\d))?/.exec(n.slice(e,e+6));return r?(t.Z=r[1]?0:-(r[2]+(r[3]||"00")),e+r[0].length):-1}function _p(t,n,e){var r=rp.exec(n.slice(e,e+2));return r?(t.m=r[0]-1,e+r[0].length):-1}function bp(t,n,e){var r=rp.exec(n.slice(e,e+2));return r?(t.d=+r[0],e+r[0].length):-1}function mp(t,n,e){var r=rp.exec(n.slice(e,e+3));return r?(t.m=0,t.d=+r[0],e+r[0].length):-1}function xp(t,n,e){var r=rp.exec(n.slice(e,e+2));return r?(t.H=+r[0],e+r[0].length):-1}function wp(t,n,e){var r=rp.exec(n.slice(e,e+2));return r?(t.M=+r[0],e+r[0].length):-1}function Mp(t,n,e){var r=rp.exec(n.slice(e,e+2));return r?(t.S=+r[0],e+r[0].length):-1}function Np(t,n,e){var r=rp.exec(n.slice(e,e+3));return r?(t.L=+r[0],e+r[0].length):-1}function Ap(t,n,e){var r=rp.exec(n.slice(e,e+6));return r?(t.L=Math.floor(r[0]/1e3),e+r[0].length):-1}function Tp(t,n,e){var r=ip.exec(n.slice(e,e+1));return r?e+r[0].length:-1}function Sp(t,n,e){var r=rp.exec(n.slice(e));return r?(t.Q=+r[0],e+r[0].length):-1}function kp(t,n,e){var r=rp.exec(n.slice(e));return r?(t.Q=1e3*+r[0],e+r[0].length):-1}function Ep(t,n){return ap(t.getDate(),n,2)}function Cp(t,n){return ap(t.getHours(),n,2)}function Pp(t,n){return ap(t.getHours()%12||12,n,2)}function zp(t,n){return ap(1+od.count(Nd(t),t),n,3)}function Rp(t,n){return ap(t.getMilliseconds(),n,3)}function Dp(t,n){return Rp(t,n)+"000"}function qp(t,n){return ap(t.getMonth()+1,n,2)}function Lp(t,n){return ap(t.getMinutes(),n,2)}function Up(t,n){return ap(t.getSeconds(),n,2)}function Op(t){var n=t.getDay();return 0===n?7:n}function Bp(t,n){return ap(cd.count(Nd(t),t),n,2)}function Yp(t,n){var e=t.getDay();return t=e>=4||0===e?hd(t):hd.ceil(t),ap(hd.count(Nd(t),t)+(4===Nd(t).getDay()),n,2)}function Fp(t){return t.getDay()}function Ip(t,n){return ap(fd.count(Nd(t),t),n,2)}function jp(t,n){return ap(t.getFullYear()%100,n,2)}function Hp(t,n){return ap(t.getFullYear()%1e4,n,4)}function Xp(t){var n=t.getTimezoneOffset();return(n>0?"-":(n*=-1,"+"))+ap(n/60|0,"0",2)+ap(n%60,"0",2)}function Gp(t,n){return ap(t.getUTCDate(),n,2)}function Vp(t,n){return ap(t.getUTCHours(),n,2)}function $p(t,n){return ap(t.getUTCHours()%12||12,n,2)}function Wp(t,n){return ap(1+Cd.count(Wd(t),t),n,3)}function Zp(t,n){return ap(t.getUTCMilliseconds(),n,3)}function Qp(t,n){return Zp(t,n)+"000"}function Jp(t,n){return ap(t.getUTCMonth()+1,n,2)}function Kp(t,n){return ap(t.getUTCMinutes(),n,2)}function tv(t,n){return ap(t.getUTCSeconds(),n,2)}function nv(t){var n=t.getUTCDay();return 0===n?7:n}function ev(t,n){return ap(Rd.count(Wd(t),t),n,2)}function rv(t,n){var e=t.getUTCDay();return t=e>=4||0===e?Ud(t):Ud.ceil(t),ap(Ud.count(Wd(t),t)+(4===Wd(t).getUTCDay()),n,2)}function iv(t){return t.getUTCDay()}function ov(t,n){return ap(Dd.count(Wd(t),t),n,2)}function av(t,n){return ap(t.getUTCFullYear()%100,n,2)}function uv(t,n){return ap(t.getUTCFullYear()%1e4,n,4)}function cv(){return"+0000"}function fv(){return"%"}function sv(t){return+t}function lv(t){return Math.floor(+t/1e3)}function hv(n){return np=tp(n),t.timeFormat=np.format,t.timeParse=np.parse,t.utcFormat=np.utcFormat,t.utcParse=np.utcParse,np}hv({dateTime:"%x, %X",date:"%-m/%-d/%Y",time:"%-I:%M:%S %p",periods:["AM","PM"],days:["Sunday","Monday","Tuesday","Wednesday","Thursday","Friday","Saturday"],shortDays:["Sun","Mon","Tue","Wed","Thu","Fri","Sat"],months:["January","February","March","April","May","June","July","August","September","October","November","December"],shortMonths:["Jan","Feb","Mar","Apr","May","Jun","Jul","Aug","Sep","Oct","Nov","Dec"]});var dv=Date.prototype.toISOString?function(t){return t.toISOString()}:t.utcFormat("%Y-%m-%dT%H:%M:%S.%LZ");var pv=+new Date("2000-01-01T00:00:00.000Z")?function(t){var n=new Date(t);return isNaN(n)?null:n}:t.utcParse("%Y-%m-%dT%H:%M:%S.%LZ"),vv=1e3,gv=60*vv,yv=60*gv,_v=24*yv,bv=7*_v,mv=30*_v,xv=365*_v;function wv(t){return new Date(t)}function Mv(t){return t instanceof Date?+t:+new Date(+t)}function Nv(t,n,r,i,o,a,u,c,f){var s=Sh(mh,mh),l=s.invert,h=s.domain,d=f(".%L"),p=f(":%S"),v=f("%I:%M"),g=f("%I %p"),y=f("%a %d"),_=f("%b %d"),b=f("%B"),m=f("%Y"),x=[[u,1,vv],[u,5,5*vv],[u,15,15*vv],[u,30,30*vv],[a,1,gv],[a,5,5*gv],[a,15,15*gv],[a,30,30*gv],[o,1,yv],[o,3,3*yv],[o,6,6*yv],[o,12,12*yv],[i,1,_v],[i,2,2*_v],[r,1,bv],[n,1,mv],[n,3,3*mv],[t,1,xv]];function M(e){return(u(e)<e?d:a(e)<e?p:o(e)<e?v:i(e)<e?g:n(e)<e?r(e)<e?y:_:t(e)<e?b:m)(e)}function N(n,r,i,o){if(null==n&&(n=10),"number"==typeof n){var a=Math.abs(i-r)/n,u=e(function(t){return t[2]}).right(x,a);u===x.length?(o=w(r/xv,i/xv,n),n=t):u?(o=(u=x[a/x[u-1][2]<x[u][2]/a?u-1:u])[1],n=u[0]):(o=Math.max(w(r,i,n),1),n=c)}return null==o?n:n.every(o)}return s.invert=function(t){return new Date(l(t))},s.domain=function(t){return arguments.length?h(dh.call(t,Mv)):h().map(wv)},s.ticks=function(t,n){var e,r=h(),i=r[0],o=r[r.length-1],a=o<i;return a&&(e=i,i=o,o=e),e=(e=N(t,i,o,n))?e.range(i,o+1):[],a?e.reverse():e},s.tickFormat=function(t,n){return null==n?M:f(n)},s.nice=function(t,n){var e=h();return(t=N(t,e[0],e[e.length-1],n))?h(Ch(e,t)):s},s.copy=function(){return Ah(s,Nv(t,n,r,i,o,a,u,c,f))},s}function Av(){var t,n,e,r,i,o=0,a=1,u=mh,c=!1;function f(n){return isNaN(n=+n)?i:u(0===e?.5:(n=(r(n)-t)*e,c?Math.max(0,Math.min(1,n)):n))}return f.domain=function(i){return arguments.length?(t=r(o=+i[0]),n=r(a=+i[1]),e=t===n?0:1/(n-t),f):[o,a]},f.clamp=function(t){return arguments.length?(c=!!t,f):c},f.interpolator=function(t){return arguments.length?(u=t,f):u},f.unknown=function(t){return arguments.length?(i=t,f):i},function(i){return r=i,t=i(o),n=i(a),e=t===n?0:1/(n-t),f}}function Tv(t,n){return n.domain(t.domain()).interpolator(t.interpolator()).clamp(t.clamp()).unknown(t.unknown())}function Sv(){var t=Hh(Av());return t.copy=function(){return Tv(t,Sv()).exponent(t.exponent())},lh.apply(t,arguments)}function kv(){var t,n,e,r,i,o,a,u=0,c=.5,f=1,s=mh,l=!1;function h(t){return isNaN(t=+t)?a:(t=.5+((t=+o(t))-n)*(t<n?r:i),s(l?Math.max(0,Math.min(1,t)):t))}return h.domain=function(a){return arguments.length?(t=o(u=+a[0]),n=o(c=+a[1]),e=o(f=+a[2]),r=t===n?0:.5/(n-t),i=n===e?0:.5/(e-n),h):[u,c,f]},h.clamp=function(t){return arguments.length?(l=!!t,h):l},h.interpolator=function(t){return arguments.length?(s=t,h):s},h.unknown=function(t){return arguments.length?(a=t,h):a},function(a){return o=a,t=a(u),n=a(c),e=a(f),r=t===n?0:.5/(n-t),i=n===e?0:.5/(e-n),h}}function Ev(){var t=Hh(kv());return t.copy=function(){return Tv(t,Ev()).exponent(t.exponent())},lh.apply(t,arguments)}function Cv(t){for(var n=t.length/6|0,e=new Array(n),r=0;r<n;)e[r]="#"+t.slice(6*r,6*++r);return e}var Pv=Cv("1f77b4ff7f0e2ca02cd627289467bd8c564be377c27f7f7fbcbd2217becf"),zv=Cv("7fc97fbeaed4fdc086ffff99386cb0f0027fbf5b17666666"),Rv=Cv("1b9e77d95f027570b3e7298a66a61ee6ab02a6761d666666"),Dv=Cv("a6cee31f78b4b2df8a33a02cfb9a99e31a1cfdbf6fff7f00cab2d66a3d9affff99b15928"),qv=Cv("fbb4aeb3cde3ccebc5decbe4fed9a6ffffcce5d8bdfddaecf2f2f2"),Lv=Cv("b3e2cdfdcdaccbd5e8f4cae4e6f5c9fff2aef1e2cccccccc"),Uv=Cv("e41a1c377eb84daf4a984ea3ff7f00ffff33a65628f781bf999999"),Ov=Cv("66c2a5fc8d628da0cbe78ac3a6d854ffd92fe5c494b3b3b3"),Bv=Cv("8dd3c7ffffb3bebadafb807280b1d3fdb462b3de69fccde5d9d9d9bc80bdccebc5ffed6f");function Yv(t){return ce(t[t.length-1])}var Fv=new Array(3).concat("d8b365f5f5f55ab4ac","a6611adfc27d80cdc1018571","a6611adfc27df5f5f580cdc1018571","8c510ad8b365f6e8c3c7eae55ab4ac01665e","8c510ad8b365f6e8c3f5f5f5c7eae55ab4ac01665e","8c510abf812ddfc27df6e8c3c7eae580cdc135978f01665e","8c510abf812ddfc27df6e8c3f5f5f5c7eae580cdc135978f01665e","5430058c510abf812ddfc27df6e8c3c7eae580cdc135978f01665e003c30","5430058c510abf812ddfc27df6e8c3f5f5f5c7eae580cdc135978f01665e003c30").map(Cv),Iv=Yv(Fv),jv=new Array(3).concat("af8dc3f7f7f77fbf7b","7b3294c2a5cfa6dba0008837","7b3294c2a5cff7f7f7a6dba0008837","762a83af8dc3e7d4e8d9f0d37fbf7b1b7837","762a83af8dc3e7d4e8f7f7f7d9f0d37fbf7b1b7837","762a839970abc2a5cfe7d4e8d9f0d3a6dba05aae611b7837","762a839970abc2a5cfe7d4e8f7f7f7d9f0d3a6dba05aae611b7837","40004b762a839970abc2a5cfe7d4e8d9f0d3a6dba05aae611b783700441b","40004b762a839970abc2a5cfe7d4e8f7f7f7d9f0d3a6dba05aae611b783700441b").map(Cv),Hv=Yv(jv),Xv=new Array(3).concat("e9a3c9f7f7f7a1d76a","d01c8bf1b6dab8e1864dac26","d01c8bf1b6daf7f7f7b8e1864dac26","c51b7de9a3c9fde0efe6f5d0a1d76a4d9221","c51b7de9a3c9fde0eff7f7f7e6f5d0a1d76a4d9221","c51b7dde77aef1b6dafde0efe6f5d0b8e1867fbc414d9221","c51b7dde77aef1b6dafde0eff7f7f7e6f5d0b8e1867fbc414d9221","8e0152c51b7dde77aef1b6dafde0efe6f5d0b8e1867fbc414d9221276419","8e0152c51b7dde77aef1b6dafde0eff7f7f7e6f5d0b8e1867fbc414d9221276419").map(Cv),Gv=Yv(Xv),Vv=new Array(3).concat("998ec3f7f7f7f1a340","5e3c99b2abd2fdb863e66101","5e3c99b2abd2f7f7f7fdb863e66101","542788998ec3d8daebfee0b6f1a340b35806","542788998ec3d8daebf7f7f7fee0b6f1a340b35806","5427888073acb2abd2d8daebfee0b6fdb863e08214b35806","5427888073acb2abd2d8daebf7f7f7fee0b6fdb863e08214b35806","2d004b5427888073acb2abd2d8daebfee0b6fdb863e08214b358067f3b08","2d004b5427888073acb2abd2d8daebf7f7f7fee0b6fdb863e08214b358067f3b08").map(Cv),$v=Yv(Vv),Wv=new Array(3).concat("ef8a62f7f7f767a9cf","ca0020f4a58292c5de0571b0","ca0020f4a582f7f7f792c5de0571b0","b2182bef8a62fddbc7d1e5f067a9cf2166ac","b2182bef8a62fddbc7f7f7f7d1e5f067a9cf2166ac","b2182bd6604df4a582fddbc7d1e5f092c5de4393c32166ac","b2182bd6604df4a582fddbc7f7f7f7d1e5f092c5de4393c32166ac","67001fb2182bd6604df4a582fddbc7d1e5f092c5de4393c32166ac053061","67001fb2182bd6604df4a582fddbc7f7f7f7d1e5f092c5de4393c32166ac053061").map(Cv),Zv=Yv(Wv),Qv=new Array(3).concat("ef8a62ffffff999999","ca0020f4a582bababa404040","ca0020f4a582ffffffbababa404040","b2182bef8a62fddbc7e0e0e09999994d4d4d","b2182bef8a62fddbc7ffffffe0e0e09999994d4d4d","b2182bd6604df4a582fddbc7e0e0e0bababa8787874d4d4d","b2182bd6604df4a582fddbc7ffffffe0e0e0bababa8787874d4d4d","67001fb2182bd6604df4a582fddbc7e0e0e0bababa8787874d4d4d1a1a1a","67001fb2182bd6604df4a582fddbc7ffffffe0e0e0bababa8787874d4d4d1a1a1a").map(Cv),Jv=Yv(Qv),Kv=new Array(3).concat("fc8d59ffffbf91bfdb","d7191cfdae61abd9e92c7bb6","d7191cfdae61ffffbfabd9e92c7bb6","d73027fc8d59fee090e0f3f891bfdb4575b4","d73027fc8d59fee090ffffbfe0f3f891bfdb4575b4","d73027f46d43fdae61fee090e0f3f8abd9e974add14575b4","d73027f46d43fdae61fee090ffffbfe0f3f8abd9e974add14575b4","a50026d73027f46d43fdae61fee090e0f3f8abd9e974add14575b4313695","a50026d73027f46d43fdae61fee090ffffbfe0f3f8abd9e974add14575b4313695").map(Cv),tg=Yv(Kv),ng=new Array(3).concat("fc8d59ffffbf91cf60","d7191cfdae61a6d96a1a9641","d7191cfdae61ffffbfa6d96a1a9641","d73027fc8d59fee08bd9ef8b91cf601a9850","d73027fc8d59fee08bffffbfd9ef8b91cf601a9850","d73027f46d43fdae61fee08bd9ef8ba6d96a66bd631a9850","d73027f46d43fdae61fee08bffffbfd9ef8ba6d96a66bd631a9850","a50026d73027f46d43fdae61fee08bd9ef8ba6d96a66bd631a9850006837","a50026d73027f46d43fdae61fee08bffffbfd9ef8ba6d96a66bd631a9850006837").map(Cv),eg=Yv(ng),rg=new Array(3).concat("fc8d59ffffbf99d594","d7191cfdae61abdda42b83ba","d7191cfdae61ffffbfabdda42b83ba","d53e4ffc8d59fee08be6f59899d5943288bd","d53e4ffc8d59fee08bffffbfe6f59899d5943288bd","d53e4ff46d43fdae61fee08be6f598abdda466c2a53288bd","d53e4ff46d43fdae61fee08bffffbfe6f598abdda466c2a53288bd","9e0142d53e4ff46d43fdae61fee08be6f598abdda466c2a53288bd5e4fa2","9e0142d53e4ff46d43fdae61fee08bffffbfe6f598abdda466c2a53288bd5e4fa2").map(Cv),ig=Yv(rg),og=new Array(3).concat("e5f5f999d8c92ca25f","edf8fbb2e2e266c2a4238b45","edf8fbb2e2e266c2a42ca25f006d2c","edf8fbccece699d8c966c2a42ca25f006d2c","edf8fbccece699d8c966c2a441ae76238b45005824","f7fcfde5f5f9ccece699d8c966c2a441ae76238b45005824","f7fcfde5f5f9ccece699d8c966c2a441ae76238b45006d2c00441b").map(Cv),ag=Yv(og),ug=new Array(3).concat("e0ecf49ebcda8856a7","edf8fbb3cde38c96c688419d","edf8fbb3cde38c96c68856a7810f7c","edf8fbbfd3e69ebcda8c96c68856a7810f7c","edf8fbbfd3e69ebcda8c96c68c6bb188419d6e016b","f7fcfde0ecf4bfd3e69ebcda8c96c68c6bb188419d6e016b","f7fcfde0ecf4bfd3e69ebcda8c96c68c6bb188419d810f7c4d004b").map(Cv),cg=Yv(ug),fg=new Array(3).concat("e0f3dba8ddb543a2ca","f0f9e8bae4bc7bccc42b8cbe","f0f9e8bae4bc7bccc443a2ca0868ac","f0f9e8ccebc5a8ddb57bccc443a2ca0868ac","f0f9e8ccebc5a8ddb57bccc44eb3d32b8cbe08589e","f7fcf0e0f3dbccebc5a8ddb57bccc44eb3d32b8cbe08589e","f7fcf0e0f3dbccebc5a8ddb57bccc44eb3d32b8cbe0868ac084081").map(Cv),sg=Yv(fg),lg=new Array(3).concat("fee8c8fdbb84e34a33","fef0d9fdcc8afc8d59d7301f","fef0d9fdcc8afc8d59e34a33b30000","fef0d9fdd49efdbb84fc8d59e34a33b30000","fef0d9fdd49efdbb84fc8d59ef6548d7301f990000","fff7ecfee8c8fdd49efdbb84fc8d59ef6548d7301f990000","fff7ecfee8c8fdd49efdbb84fc8d59ef6548d7301fb300007f0000").map(Cv),hg=Yv(lg),dg=new Array(3).concat("ece2f0a6bddb1c9099","f6eff7bdc9e167a9cf02818a","f6eff7bdc9e167a9cf1c9099016c59","f6eff7d0d1e6a6bddb67a9cf1c9099016c59","f6eff7d0d1e6a6bddb67a9cf3690c002818a016450","fff7fbece2f0d0d1e6a6bddb67a9cf3690c002818a016450","fff7fbece2f0d0d1e6a6bddb67a9cf3690c002818a016c59014636").map(Cv),pg=Yv(dg),vg=new Array(3).concat("ece7f2a6bddb2b8cbe","f1eef6bdc9e174a9cf0570b0","f1eef6bdc9e174a9cf2b8cbe045a8d","f1eef6d0d1e6a6bddb74a9cf2b8cbe045a8d","f1eef6d0d1e6a6bddb74a9cf3690c00570b0034e7b","fff7fbece7f2d0d1e6a6bddb74a9cf3690c00570b0034e7b","fff7fbece7f2d0d1e6a6bddb74a9cf3690c00570b0045a8d023858").map(Cv),gg=Yv(vg),yg=new Array(3).concat("e7e1efc994c7dd1c77","f1eef6d7b5d8df65b0ce1256","f1eef6d7b5d8df65b0dd1c77980043","f1eef6d4b9dac994c7df65b0dd1c77980043","f1eef6d4b9dac994c7df65b0e7298ace125691003f","f7f4f9e7e1efd4b9dac994c7df65b0e7298ace125691003f","f7f4f9e7e1efd4b9dac994c7df65b0e7298ace125698004367001f").map(Cv),_g=Yv(yg),bg=new Array(3).concat("fde0ddfa9fb5c51b8a","feebe2fbb4b9f768a1ae017e","feebe2fbb4b9f768a1c51b8a7a0177","feebe2fcc5c0fa9fb5f768a1c51b8a7a0177","feebe2fcc5c0fa9fb5f768a1dd3497ae017e7a0177","fff7f3fde0ddfcc5c0fa9fb5f768a1dd3497ae017e7a0177","fff7f3fde0ddfcc5c0fa9fb5f768a1dd3497ae017e7a017749006a").map(Cv),mg=Yv(bg),xg=new Array(3).concat("edf8b17fcdbb2c7fb8","ffffcca1dab441b6c4225ea8","ffffcca1dab441b6c42c7fb8253494","ffffccc7e9b47fcdbb41b6c42c7fb8253494","ffffccc7e9b47fcdbb41b6c41d91c0225ea80c2c84","ffffd9edf8b1c7e9b47fcdbb41b6c41d91c0225ea80c2c84","ffffd9edf8b1c7e9b47fcdbb41b6c41d91c0225ea8253494081d58").map(Cv),wg=Yv(xg),Mg=new Array(3).concat("f7fcb9addd8e31a354","ffffccc2e69978c679238443","ffffccc2e69978c67931a354006837","ffffccd9f0a3addd8e78c67931a354006837","ffffccd9f0a3addd8e78c67941ab5d238443005a32","ffffe5f7fcb9d9f0a3addd8e78c67941ab5d238443005a32","ffffe5f7fcb9d9f0a3addd8e78c67941ab5d238443006837004529").map(Cv),Ng=Yv(Mg),Ag=new Array(3).concat("fff7bcfec44fd95f0e","ffffd4fed98efe9929cc4c02","ffffd4fed98efe9929d95f0e993404","ffffd4fee391fec44ffe9929d95f0e993404","ffffd4fee391fec44ffe9929ec7014cc4c028c2d04","ffffe5fff7bcfee391fec44ffe9929ec7014cc4c028c2d04","ffffe5fff7bcfee391fec44ffe9929ec7014cc4c02993404662506").map(Cv),Tg=Yv(Ag),Sg=new Array(3).concat("ffeda0feb24cf03b20","ffffb2fecc5cfd8d3ce31a1c","ffffb2fecc5cfd8d3cf03b20bd0026","ffffb2fed976feb24cfd8d3cf03b20bd0026","ffffb2fed976feb24cfd8d3cfc4e2ae31a1cb10026","ffffccffeda0fed976feb24cfd8d3cfc4e2ae31a1cb10026","ffffccffeda0fed976feb24cfd8d3cfc4e2ae31a1cbd0026800026").map(Cv),kg=Yv(Sg),Eg=new Array(3).concat("deebf79ecae13182bd","eff3ffbdd7e76baed62171b5","eff3ffbdd7e76baed63182bd08519c","eff3ffc6dbef9ecae16baed63182bd08519c","eff3ffc6dbef9ecae16baed64292c62171b5084594","f7fbffdeebf7c6dbef9ecae16baed64292c62171b5084594","f7fbffdeebf7c6dbef9ecae16baed64292c62171b508519c08306b").map(Cv),Cg=Yv(Eg),Pg=new Array(3).concat("e5f5e0a1d99b31a354","edf8e9bae4b374c476238b45","edf8e9bae4b374c47631a354006d2c","edf8e9c7e9c0a1d99b74c47631a354006d2c","edf8e9c7e9c0a1d99b74c47641ab5d238b45005a32","f7fcf5e5f5e0c7e9c0a1d99b74c47641ab5d238b45005a32","f7fcf5e5f5e0c7e9c0a1d99b74c47641ab5d238b45006d2c00441b").map(Cv),zg=Yv(Pg),Rg=new Array(3).concat("f0f0f0bdbdbd636363","f7f7f7cccccc969696525252","f7f7f7cccccc969696636363252525","f7f7f7d9d9d9bdbdbd969696636363252525","f7f7f7d9d9d9bdbdbd969696737373525252252525","fffffff0f0f0d9d9d9bdbdbd969696737373525252252525","fffffff0f0f0d9d9d9bdbdbd969696737373525252252525000000").map(Cv),Dg=Yv(Rg),qg=new Array(3).concat("efedf5bcbddc756bb1","f2f0f7cbc9e29e9ac86a51a3","f2f0f7cbc9e29e9ac8756bb154278f","f2f0f7dadaebbcbddc9e9ac8756bb154278f","f2f0f7dadaebbcbddc9e9ac8807dba6a51a34a1486","fcfbfdefedf5dadaebbcbddc9e9ac8807dba6a51a34a1486","fcfbfdefedf5dadaebbcbddc9e9ac8807dba6a51a354278f3f007d").map(Cv),Lg=Yv(qg),Ug=new Array(3).concat("fee0d2fc9272de2d26","fee5d9fcae91fb6a4acb181d","fee5d9fcae91fb6a4ade2d26a50f15","fee5d9fcbba1fc9272fb6a4ade2d26a50f15","fee5d9fcbba1fc9272fb6a4aef3b2ccb181d99000d","fff5f0fee0d2fcbba1fc9272fb6a4aef3b2ccb181d99000d","fff5f0fee0d2fcbba1fc9272fb6a4aef3b2ccb181da50f1567000d").map(Cv),Og=Yv(Ug),Bg=new Array(3).concat("fee6cefdae6be6550d","feeddefdbe85fd8d3cd94701","feeddefdbe85fd8d3ce6550da63603","feeddefdd0a2fdae6bfd8d3ce6550da63603","feeddefdd0a2fdae6bfd8d3cf16913d948018c2d04","fff5ebfee6cefdd0a2fdae6bfd8d3cf16913d948018c2d04","fff5ebfee6cefdd0a2fdae6bfd8d3cf16913d94801a636037f2704").map(Cv),Yg=Yv(Bg),Fg=je(Zn(300,.5,0),Zn(-240,.5,1)),Ig=je(Zn(-100,.75,.35),Zn(80,1.5,.8)),jg=je(Zn(260,.75,.35),Zn(80,1.5,.8)),Hg=Zn();var Xg=gn(),Gg=Math.PI/3,Vg=2*Math.PI/3;function $g(t){var n=t.length;return function(e){return t[Math.max(0,Math.min(n-1,Math.floor(e*n)))]}}var Wg=$g(Cv("44015444025645045745055946075a46085c460a5d460b5e470d60470e6147106347116447136548146748166848176948186a481a6c481b6d481c6e481d6f481f70482071482173482374482475482576482677482878482979472a7a472c7a472d7b472e7c472f7d46307e46327e46337f463480453581453781453882443983443a83443b84433d84433e85423f854240864241864142874144874045884046883f47883f48893e49893e4a893e4c8a3d4d8a3d4e8a3c4f8a3c508b3b518b3b528b3a538b3a548c39558c39568c38588c38598c375a8c375b8d365c8d365d8d355e8d355f8d34608d34618d33628d33638d32648e32658e31668e31678e31688e30698e306a8e2f6b8e2f6c8e2e6d8e2e6e8e2e6f8e2d708e2d718e2c718e2c728e2c738e2b748e2b758e2a768e2a778e2a788e29798e297a8e297b8e287c8e287d8e277e8e277f8e27808e26818e26828e26828e25838e25848e25858e24868e24878e23888e23898e238a8d228b8d228c8d228d8d218e8d218f8d21908d21918c20928c20928c20938c1f948c1f958b1f968b1f978b1f988b1f998a1f9a8a1e9b8a1e9c891e9d891f9e891f9f881fa0881fa1881fa1871fa28720a38620a48621a58521a68522a78522a88423a98324aa8325ab8225ac8226ad8127ad8128ae8029af7f2ab07f2cb17e2db27d2eb37c2fb47c31b57b32b67a34b67935b77937b87838b9773aba763bbb753dbc743fbc7340bd7242be7144bf7046c06f48c16e4ac16d4cc26c4ec36b50c46a52c56954c56856c66758c7655ac8645cc8635ec96260ca6063cb5f65cb5e67cc5c69cd5b6ccd5a6ece5870cf5773d05675d05477d1537ad1517cd2507fd34e81d34d84d44b86d54989d5488bd6468ed64590d74393d74195d84098d83e9bd93c9dd93ba0da39a2da37a5db36a8db34aadc32addc30b0dd2fb2dd2db5de2bb8de29bade28bddf26c0df25c2df23c5e021c8e020cae11fcde11dd0e11cd2e21bd5e21ad8e219dae319dde318dfe318e2e418e5e419e7e419eae51aece51befe51cf1e51df4e61ef6e620f8e621fbe723fde725")),Zg=$g(Cv("00000401000501010601010802010902020b02020d03030f03031204041405041606051806051a07061c08071e0907200a08220b09240c09260d0a290e0b2b100b2d110c2f120d31130d34140e36150e38160f3b180f3d19103f1a10421c10441d11471e114920114b21114e22115024125325125527125829115a2a115c2c115f2d11612f116331116533106734106936106b38106c390f6e3b0f703d0f713f0f72400f74420f75440f764510774710784910784a10794c117a4e117b4f127b51127c52137c54137d56147d57157e59157e5a167e5c167f5d177f5f187f601880621980641a80651a80671b80681c816a1c816b1d816d1d816e1e81701f81721f817320817521817621817822817922827b23827c23827e24828025828125818326818426818627818827818928818b29818c29818e2a81902a81912b81932b80942c80962c80982d80992d809b2e7f9c2e7f9e2f7fa02f7fa1307ea3307ea5317ea6317da8327daa337dab337cad347cae347bb0357bb2357bb3367ab5367ab73779b83779ba3878bc3978bd3977bf3a77c03a76c23b75c43c75c53c74c73d73c83e73ca3e72cc3f71cd4071cf4070d0416fd2426fd3436ed5446dd6456cd8456cd9466bdb476adc4869de4968df4a68e04c67e24d66e34e65e44f64e55064e75263e85362e95462ea5661eb5760ec5860ed5a5fee5b5eef5d5ef05f5ef1605df2625df2645cf3655cf4675cf4695cf56b5cf66c5cf66e5cf7705cf7725cf8745cf8765cf9785df9795df97b5dfa7d5efa7f5efa815ffb835ffb8560fb8761fc8961fc8a62fc8c63fc8e64fc9065fd9266fd9467fd9668fd9869fd9a6afd9b6bfe9d6cfe9f6dfea16efea36ffea571fea772fea973feaa74feac76feae77feb078feb27afeb47bfeb67cfeb77efeb97ffebb81febd82febf84fec185fec287fec488fec68afec88cfeca8dfecc8ffecd90fecf92fed194fed395fed597fed799fed89afdda9cfddc9efddea0fde0a1fde2a3fde3a5fde5a7fde7a9fde9aafdebacfcecaefceeb0fcf0b2fcf2b4fcf4b6fcf6b8fcf7b9fcf9bbfcfbbdfcfdbf")),Qg=$g(Cv("00000401000501010601010802010a02020c02020e03021004031204031405041706041907051b08051d09061f0a07220b07240c08260d08290e092b10092d110a30120a32140b34150b37160b39180c3c190c3e1b0c411c0c431e0c451f0c48210c4a230c4c240c4f260c51280b53290b552b0b572d0b592f0a5b310a5c320a5e340a5f3609613809623909633b09643d09653e0966400a67420a68440a68450a69470b6a490b6a4a0c6b4c0c6b4d0d6c4f0d6c510e6c520e6d540f6d550f6d57106e59106e5a116e5c126e5d126e5f136e61136e62146e64156e65156e67166e69166e6a176e6c186e6d186e6f196e71196e721a6e741a6e751b6e771c6d781c6d7a1d6d7c1d6d7d1e6d7f1e6c801f6c82206c84206b85216b87216b88226a8a226a8c23698d23698f24699025689225689326679526679727669827669a28659b29649d29649f2a63a02a63a22b62a32c61a52c60a62d60a82e5fa92e5eab2f5ead305dae305cb0315bb1325ab3325ab43359b63458b73557b93556ba3655bc3754bd3853bf3952c03a51c13a50c33b4fc43c4ec63d4dc73e4cc83f4bca404acb4149cc4248ce4347cf4446d04545d24644d34743d44842d54a41d74b3fd84c3ed94d3dda4e3cdb503bdd513ade5238df5337e05536e15635e25734e35933e45a31e55c30e65d2fe75e2ee8602de9612bea632aeb6429eb6628ec6726ed6925ee6a24ef6c23ef6e21f06f20f1711ff1731df2741cf3761bf37819f47918f57b17f57d15f67e14f68013f78212f78410f8850ff8870ef8890cf98b0bf98c0af98e09fa9008fa9207fa9407fb9606fb9706fb9906fb9b06fb9d07fc9f07fca108fca309fca50afca60cfca80dfcaa0ffcac11fcae12fcb014fcb216fcb418fbb61afbb81dfbba1ffbbc21fbbe23fac026fac228fac42afac62df9c72ff9c932f9cb35f8cd37f8cf3af7d13df7d340f6d543f6d746f5d949f5db4cf4dd4ff4df53f4e156f3e35af3e55df2e661f2e865f2ea69f1ec6df1ed71f1ef75f1f179f2f27df2f482f3f586f3f68af4f88ef5f992f6fa96f8fb9af9fc9dfafda1fcffa4")),Jg=$g(Cv("0d088710078813078916078a19068c1b068d1d068e20068f2206902406912605912805922a05932c05942e05952f059631059733059735049837049938049a3a049a3c049b3e049c3f049c41049d43039e44039e46039f48039f4903a04b03a14c02a14e02a25002a25102a35302a35502a45601a45801a45901a55b01a55c01a65e01a66001a66100a76300a76400a76600a76700a86900a86a00a86c00a86e00a86f00a87100a87201a87401a87501a87701a87801a87a02a87b02a87d03a87e03a88004a88104a78305a78405a78606a68707a68808a68a09a58b0aa58d0ba58e0ca48f0da4910ea3920fa39410a29511a19613a19814a099159f9a169f9c179e9d189d9e199da01a9ca11b9ba21d9aa31e9aa51f99a62098a72197a82296aa2395ab2494ac2694ad2793ae2892b02991b12a90b22b8fb32c8eb42e8db52f8cb6308bb7318ab83289ba3388bb3488bc3587bd3786be3885bf3984c03a83c13b82c23c81c33d80c43e7fc5407ec6417dc7427cc8437bc9447aca457acb4679cc4778cc4977cd4a76ce4b75cf4c74d04d73d14e72d24f71d35171d45270d5536fd5546ed6556dd7566cd8576bd9586ada5a6ada5b69db5c68dc5d67dd5e66de5f65de6164df6263e06363e16462e26561e26660e3685fe4695ee56a5de56b5de66c5ce76e5be76f5ae87059e97158e97257ea7457eb7556eb7655ec7754ed7953ed7a52ee7b51ef7c51ef7e50f07f4ff0804ef1814df1834cf2844bf3854bf3874af48849f48948f58b47f58c46f68d45f68f44f79044f79143f79342f89441f89540f9973ff9983ef99a3efa9b3dfa9c3cfa9e3bfb9f3afba139fba238fca338fca537fca636fca835fca934fdab33fdac33fdae32fdaf31fdb130fdb22ffdb42ffdb52efeb72dfeb82cfeba2cfebb2bfebd2afebe2afec029fdc229fdc328fdc527fdc627fdc827fdca26fdcb26fccd25fcce25fcd025fcd225fbd324fbd524fbd724fad824fada24f9dc24f9dd25f8df25f8e125f7e225f7e425f6e626f6e826f5e926f5eb27f4ed27f3ee27f3f027f2f227f1f426f1f525f0f724f0f921"));function Kg(t){return function(){return t}}var ty=Math.abs,ny=Math.atan2,ey=Math.cos,ry=Math.max,iy=Math.min,oy=Math.sin,ay=Math.sqrt,uy=1e-12,cy=Math.PI,fy=cy/2,sy=2*cy;function ly(t){return t>=1?fy:t<=-1?-fy:Math.asin(t)}function hy(t){return t.innerRadius}function dy(t){return t.outerRadius}function py(t){return t.startAngle}function vy(t){return t.endAngle}function gy(t){return t&&t.padAngle}function yy(t,n,e,r,i,o,a){var u=t-e,c=n-r,f=(a?o:-o)/ay(u*u+c*c),s=f*c,l=-f*u,h=t+s,d=n+l,p=e+s,v=r+l,g=(h+p)/2,y=(d+v)/2,_=p-h,b=v-d,m=_*_+b*b,x=i-o,w=h*v-p*d,M=(b<0?-1:1)*ay(ry(0,x*x*m-w*w)),N=(w*b-_*M)/m,A=(-w*_-b*M)/m,T=(w*b+_*M)/m,S=(-w*_+b*M)/m,k=N-g,E=A-y,C=T-g,P=S-y;return k*k+E*E>C*C+P*P&&(N=T,A=S),{cx:N,cy:A,x01:-s,y01:-l,x11:N*(i/x-1),y11:A*(i/x-1)}}function _y(t){this._context=t}function by(t){return new _y(t)}function my(t){return t[0]}function xy(t){return t[1]}function wy(){var t=my,n=xy,e=Kg(!0),r=null,i=by,o=null;function a(a){var u,c,f,s=a.length,l=!1;for(null==r&&(o=i(f=Hi())),u=0;u<=s;++u)!(u<s&&e(c=a[u],u,a))===l&&((l=!l)?o.lineStart():o.lineEnd()),l&&o.point(+t(c,u,a),+n(c,u,a));if(f)return o=null,f+""||null}return a.x=function(n){return arguments.length?(t="function"==typeof n?n:Kg(+n),a):t},a.y=function(t){return arguments.length?(n="function"==typeof t?t:Kg(+t),a):n},a.defined=function(t){return arguments.length?(e="function"==typeof t?t:Kg(!!t),a):e},a.curve=function(t){return arguments.length?(i=t,null!=r&&(o=i(r)),a):i},a.context=function(t){return arguments.length?(null==t?r=o=null:o=i(r=t),a):r},a}function My(){var t=my,n=null,e=Kg(0),r=xy,i=Kg(!0),o=null,a=by,u=null;function c(c){var f,s,l,h,d,p=c.length,v=!1,g=new Array(p),y=new Array(p);for(null==o&&(u=a(d=Hi())),f=0;f<=p;++f){if(!(f<p&&i(h=c[f],f,c))===v)if(v=!v)s=f,u.areaStart(),u.lineStart();else{for(u.lineEnd(),u.lineStart(),l=f-1;l>=s;--l)u.point(g[l],y[l]);u.lineEnd(),u.areaEnd()}v&&(g[f]=+t(h,f,c),y[f]=+e(h,f,c),u.point(n?+n(h,f,c):g[f],r?+r(h,f,c):y[f]))}if(d)return u=null,d+""||null}function f(){return wy().defined(i).curve(a).context(o)}return c.x=function(e){return arguments.length?(t="function"==typeof e?e:Kg(+e),n=null,c):t},c.x0=function(n){return arguments.length?(t="function"==typeof n?n:Kg(+n),c):t},c.x1=function(t){return arguments.length?(n=null==t?null:"function"==typeof t?t:Kg(+t),c):n},c.y=function(t){return arguments.length?(e="function"==typeof t?t:Kg(+t),r=null,c):e},c.y0=function(t){return arguments.length?(e="function"==typeof t?t:Kg(+t),c):e},c.y1=function(t){return arguments.length?(r=null==t?null:"function"==typeof t?t:Kg(+t),c):r},c.lineX0=c.lineY0=function(){return f().x(t).y(e)},c.lineY1=function(){return f().x(t).y(r)},c.lineX1=function(){return f().x(n).y(e)},c.defined=function(t){return arguments.length?(i="function"==typeof t?t:Kg(!!t),c):i},c.curve=function(t){return arguments.length?(a=t,null!=o&&(u=a(o)),c):a},c.context=function(t){return arguments.length?(null==t?o=u=null:u=a(o=t),c):o},c}function Ny(t,n){return n<t?-1:n>t?1:n>=t?0:NaN}function Ay(t){return t}_y.prototype={areaStart:function(){this._line=0},areaEnd:function(){this._line=NaN},lineStart:function(){this._point=0},lineEnd:function(){(this._line||0!==this._line&&1===this._point)&&this._context.closePath(),this._line=1-this._line},point:function(t,n){switch(t=+t,n=+n,this._point){case 0:this._point=1,this._line?this._context.lineTo(t,n):this._context.moveTo(t,n);break;case 1:this._point=2;default:this._context.lineTo(t,n)}}};var Ty=ky(by);function Sy(t){this._curve=t}function ky(t){function n(n){return new Sy(t(n))}return n._curve=t,n}function Ey(t){var n=t.curve;return t.angle=t.x,delete t.x,t.radius=t.y,delete t.y,t.curve=function(t){return arguments.length?n(ky(t)):n()._curve},t}function Cy(){return Ey(wy().curve(Ty))}function Py(){var t=My().curve(Ty),n=t.curve,e=t.lineX0,r=t.lineX1,i=t.lineY0,o=t.lineY1;return t.angle=t.x,delete t.x,t.startAngle=t.x0,delete t.x0,t.endAngle=t.x1,delete t.x1,t.radius=t.y,delete t.y,t.innerRadius=t.y0,delete t.y0,t.outerRadius=t.y1,delete t.y1,t.lineStartAngle=function(){return Ey(e())},delete t.lineX0,t.lineEndAngle=function(){return Ey(r())},delete t.lineX1,t.lineInnerRadius=function(){return Ey(i())},delete t.lineY0,t.lineOuterRadius=function(){return Ey(o())},delete t.lineY1,t.curve=function(t){return arguments.length?n(ky(t)):n()._curve},t}function zy(t,n){return[(n=+n)*Math.cos(t-=Math.PI/2),n*Math.sin(t)]}Sy.prototype={areaStart:function(){this._curve.areaStart()},areaEnd:function(){this._curve.areaEnd()},lineStart:function(){this._curve.lineStart()},lineEnd:function(){this._curve.lineEnd()},point:function(t,n){this._curve.point(n*Math.sin(t),n*-Math.cos(t))}};var Ry=Array.prototype.slice;function Dy(t){return t.source}function qy(t){return t.target}function Ly(t){var n=Dy,e=qy,r=my,i=xy,o=null;function a(){var a,u=Ry.call(arguments),c=n.apply(this,u),f=e.apply(this,u);if(o||(o=a=Hi()),t(o,+r.apply(this,(u[0]=c,u)),+i.apply(this,u),+r.apply(this,(u[0]=f,u)),+i.apply(this,u)),a)return o=null,a+""||null}return a.source=function(t){return arguments.length?(n=t,a):n},a.target=function(t){return arguments.length?(e=t,a):e},a.x=function(t){return arguments.length?(r="function"==typeof t?t:Kg(+t),a):r},a.y=function(t){return arguments.length?(i="function"==typeof t?t:Kg(+t),a):i},a.context=function(t){return arguments.length?(o=null==t?null:t,a):o},a}function Uy(t,n,e,r,i){t.moveTo(n,e),t.bezierCurveTo(n=(n+r)/2,e,n,i,r,i)}function Oy(t,n,e,r,i){t.moveTo(n,e),t.bezierCurveTo(n,e=(e+i)/2,r,e,r,i)}function By(t,n,e,r,i){var o=zy(n,e),a=zy(n,e=(e+i)/2),u=zy(r,e),c=zy(r,i);t.moveTo(o[0],o[1]),t.bezierCurveTo(a[0],a[1],u[0],u[1],c[0],c[1])}var Yy={draw:function(t,n){var e=Math.sqrt(n/cy);t.moveTo(e,0),t.arc(0,0,e,0,sy)}},Fy={draw:function(t,n){var e=Math.sqrt(n/5)/2;t.moveTo(-3*e,-e),t.lineTo(-e,-e),t.lineTo(-e,-3*e),t.lineTo(e,-3*e),t.lineTo(e,-e),t.lineTo(3*e,-e),t.lineTo(3*e,e),t.lineTo(e,e),t.lineTo(e,3*e),t.lineTo(-e,3*e),t.lineTo(-e,e),t.lineTo(-3*e,e),t.closePath()}},Iy=Math.sqrt(1/3),jy=2*Iy,Hy={draw:function(t,n){var e=Math.sqrt(n/jy),r=e*Iy;t.moveTo(0,-e),t.lineTo(r,0),t.lineTo(0,e),t.lineTo(-r,0),t.closePath()}},Xy=Math.sin(cy/10)/Math.sin(7*cy/10),Gy=Math.sin(sy/10)*Xy,Vy=-Math.cos(sy/10)*Xy,$y={draw:function(t,n){var e=Math.sqrt(.8908130915292852*n),r=Gy*e,i=Vy*e;t.moveTo(0,-e),t.lineTo(r,i);for(var o=1;o<5;++o){var a=sy*o/5,u=Math.cos(a),c=Math.sin(a);t.lineTo(c*e,-u*e),t.lineTo(u*r-c*i,c*r+u*i)}t.closePath()}},Wy={draw:function(t,n){var e=Math.sqrt(n),r=-e/2;t.rect(r,r,e,e)}},Zy=Math.sqrt(3),Qy={draw:function(t,n){var e=-Math.sqrt(n/(3*Zy));t.moveTo(0,2*e),t.lineTo(-Zy*e,-e),t.lineTo(Zy*e,-e),t.closePath()}},Jy=Math.sqrt(3)/2,Ky=1/Math.sqrt(12),t_=3*(Ky/2+1),n_={draw:function(t,n){var e=Math.sqrt(n/t_),r=e/2,i=e*Ky,o=r,a=e*Ky+e,u=-o,c=a;t.moveTo(r,i),t.lineTo(o,a),t.lineTo(u,c),t.lineTo(-.5*r-Jy*i,Jy*r+-.5*i),t.lineTo(-.5*o-Jy*a,Jy*o+-.5*a),t.lineTo(-.5*u-Jy*c,Jy*u+-.5*c),t.lineTo(-.5*r+Jy*i,-.5*i-Jy*r),t.lineTo(-.5*o+Jy*a,-.5*a-Jy*o),t.lineTo(-.5*u+Jy*c,-.5*c-Jy*u),t.closePath()}},e_=[Yy,Fy,Hy,Wy,$y,Qy,n_];function r_(){}function i_(t,n,e){t._context.bezierCurveTo((2*t._x0+t._x1)/3,(2*t._y0+t._y1)/3,(t._x0+2*t._x1)/3,(t._y0+2*t._y1)/3,(t._x0+4*t._x1+n)/6,(t._y0+4*t._y1+e)/6)}function o_(t){this._context=t}function a_(t){this._context=t}function u_(t){this._context=t}function c_(t,n){this._basis=new o_(t),this._beta=n}o_.prototype={areaStart:function(){this._line=0},areaEnd:function(){this._line=NaN},lineStart:function(){this._x0=this._x1=this._y0=this._y1=NaN,this._point=0},lineEnd:function(){switch(this._point){case 3:i_(this,this._x1,this._y1);case 2:this._context.lineTo(this._x1,this._y1)}(this._line||0!==this._line&&1===this._point)&&this._context.closePath(),this._line=1-this._line},point:function(t,n){switch(t=+t,n=+n,this._point){case 0:this._point=1,this._line?this._context.lineTo(t,n):this._context.moveTo(t,n);break;case 1:this._point=2;break;case 2:this._point=3,this._context.lineTo((5*this._x0+this._x1)/6,(5*this._y0+this._y1)/6);default:i_(this,t,n)}this._x0=this._x1,this._x1=t,this._y0=this._y1,this._y1=n}},a_.prototype={areaStart:r_,areaEnd:r_,lineStart:function(){this._x0=this._x1=this._x2=this._x3=this._x4=this._y0=this._y1=this._y2=this._y3=this._y4=NaN,this._point=0},lineEnd:function(){switch(this._point){case 1:this._context.moveTo(this._x2,this._y2),this._context.closePath();break;case 2:this._context.moveTo((this._x2+2*this._x3)/3,(this._y2+2*this._y3)/3),this._context.lineTo((this._x3+2*this._x2)/3,(this._y3+2*this._y2)/3),this._context.closePath();break;case 3:this.point(this._x2,this._y2),this.point(this._x3,this._y3),this.point(this._x4,this._y4)}},point:function(t,n){switch(t=+t,n=+n,this._point){case 0:this._point=1,this._x2=t,this._y2=n;break;case 1:this._point=2,this._x3=t,this._y3=n;break;case 2:this._point=3,this._x4=t,this._y4=n,this._context.moveTo((this._x0+4*this._x1+t)/6,(this._y0+4*this._y1+n)/6);break;default:i_(this,t,n)}this._x0=this._x1,this._x1=t,this._y0=this._y1,this._y1=n}},u_.prototype={areaStart:function(){this._line=0},areaEnd:function(){this._line=NaN},lineStart:function(){this._x0=this._x1=this._y0=this._y1=NaN,this._point=0},lineEnd:function(){(this._line||0!==this._line&&3===this._point)&&this._context.closePath(),this._line=1-this._line},point:function(t,n){switch(t=+t,n=+n,this._point){case 0:this._point=1;break;case 1:this._point=2;break;case 2:this._point=3;var e=(this._x0+4*this._x1+t)/6,r=(this._y0+4*this._y1+n)/6;this._line?this._context.lineTo(e,r):this._context.moveTo(e,r);break;case 3:this._point=4;default:i_(this,t,n)}this._x0=this._x1,this._x1=t,this._y0=this._y1,this._y1=n}},c_.prototype={lineStart:function(){this._x=[],this._y=[],this._basis.lineStart()},lineEnd:function(){var t=this._x,n=this._y,e=t.length-1;if(e>0)for(var r,i=t[0],o=n[0],a=t[e]-i,u=n[e]-o,c=-1;++c<=e;)r=c/e,this._basis.point(this._beta*t[c]+(1-this._beta)*(i+r*a),this._beta*n[c]+(1-this._beta)*(o+r*u));this._x=this._y=null,this._basis.lineEnd()},point:function(t,n){this._x.push(+t),this._y.push(+n)}};var f_=function t(n){function e(t){return 1===n?new o_(t):new c_(t,n)}return e.beta=function(n){return t(+n)},e}(.85);function s_(t,n,e){t._context.bezierCurveTo(t._x1+t._k*(t._x2-t._x0),t._y1+t._k*(t._y2-t._y0),t._x2+t._k*(t._x1-n),t._y2+t._k*(t._y1-e),t._x2,t._y2)}function l_(t,n){this._context=t,this._k=(1-n)/6}l_.prototype={areaStart:function(){this._line=0},areaEnd:function(){this._line=NaN},lineStart:function(){this._x0=this._x1=this._x2=this._y0=this._y1=this._y2=NaN,this._point=0},lineEnd:function(){switch(this._point){case 2:this._context.lineTo(this._x2,this._y2);break;case 3:s_(this,this._x1,this._y1)}(this._line||0!==this._line&&1===this._point)&&this._context.closePath(),this._line=1-this._line},point:function(t,n){switch(t=+t,n=+n,this._point){case 0:this._point=1,this._line?this._context.lineTo(t,n):this._context.moveTo(t,n);break;case 1:this._point=2,this._x1=t,this._y1=n;break;case 2:this._point=3;default:s_(this,t,n)}this._x0=this._x1,this._x1=this._x2,this._x2=t,this._y0=this._y1,this._y1=this._y2,this._y2=n}};var h_=function t(n){function e(t){return new l_(t,n)}return e.tension=function(n){return t(+n)},e}(0);function d_(t,n){this._context=t,this._k=(1-n)/6}d_.prototype={areaStart:r_,areaEnd:r_,lineStart:function(){this._x0=this._x1=this._x2=this._x3=this._x4=this._x5=this._y0=this._y1=this._y2=this._y3=this._y4=this._y5=NaN,this._point=0},lineEnd:function(){switch(this._point){case 1:this._context.moveTo(this._x3,this._y3),this._context.closePath();break;case 2:this._context.lineTo(this._x3,this._y3),this._context.closePath();break;case 3:this.point(this._x3,this._y3),this.point(this._x4,this._y4),this.point(this._x5,this._y5)}},point:function(t,n){switch(t=+t,n=+n,this._point){case 0:this._point=1,this._x3=t,this._y3=n;break;case 1:this._point=2,this._context.moveTo(this._x4=t,this._y4=n);break;case 2:this._point=3,this._x5=t,this._y5=n;break;default:s_(this,t,n)}this._x0=this._x1,this._x1=this._x2,this._x2=t,this._y0=this._y1,this._y1=this._y2,this._y2=n}};var p_=function t(n){function e(t){return new d_(t,n)}return e.tension=function(n){return t(+n)},e}(0);function v_(t,n){this._context=t,this._k=(1-n)/6}v_.prototype={areaStart:function(){this._line=0},areaEnd:function(){this._line=NaN},lineStart:function(){this._x0=this._x1=this._x2=this._y0=this._y1=this._y2=NaN,this._point=0},lineEnd:function(){(this._line||0!==this._line&&3===this._point)&&this._context.closePath(),this._line=1-this._line},point:function(t,n){switch(t=+t,n=+n,this._point){case 0:this._point=1;break;case 1:this._point=2;break;case 2:this._point=3,this._line?this._context.lineTo(this._x2,this._y2):this._context.moveTo(this._x2,this._y2);break;case 3:this._point=4;default:s_(this,t,n)}this._x0=this._x1,this._x1=this._x2,this._x2=t,this._y0=this._y1,this._y1=this._y2,this._y2=n}};var g_=function t(n){function e(t){return new v_(t,n)}return e.tension=function(n){return t(+n)},e}(0);function y_(t,n,e){var r=t._x1,i=t._y1,o=t._x2,a=t._y2;if(t._l01_a>uy){var u=2*t._l01_2a+3*t._l01_a*t._l12_a+t._l12_2a,c=3*t._l01_a*(t._l01_a+t._l12_a);r=(r*u-t._x0*t._l12_2a+t._x2*t._l01_2a)/c,i=(i*u-t._y0*t._l12_2a+t._y2*t._l01_2a)/c}if(t._l23_a>uy){var f=2*t._l23_2a+3*t._l23_a*t._l12_a+t._l12_2a,s=3*t._l23_a*(t._l23_a+t._l12_a);o=(o*f+t._x1*t._l23_2a-n*t._l12_2a)/s,a=(a*f+t._y1*t._l23_2a-e*t._l12_2a)/s}t._context.bezierCurveTo(r,i,o,a,t._x2,t._y2)}function __(t,n){this._context=t,this._alpha=n}__.prototype={areaStart:function(){this._line=0},areaEnd:function(){this._line=NaN},lineStart:function(){this._x0=this._x1=this._x2=this._y0=this._y1=this._y2=NaN,this._l01_a=this._l12_a=this._l23_a=this._l01_2a=this._l12_2a=this._l23_2a=this._point=0},lineEnd:function(){switch(this._point){case 2:this._context.lineTo(this._x2,this._y2);break;case 3:this.point(this._x2,this._y2)}(this._line||0!==this._line&&1===this._point)&&this._context.closePath(),this._line=1-this._line},point:function(t,n){if(t=+t,n=+n,this._point){var e=this._x2-t,r=this._y2-n;this._l23_a=Math.sqrt(this._l23_2a=Math.pow(e*e+r*r,this._alpha))}switch(this._point){case 0:this._point=1,this._line?this._context.lineTo(t,n):this._context.moveTo(t,n);break;case 1:this._point=2;break;case 2:this._point=3;default:y_(this,t,n)}this._l01_a=this._l12_a,this._l12_a=this._l23_a,this._l01_2a=this._l12_2a,this._l12_2a=this._l23_2a,this._x0=this._x1,this._x1=this._x2,this._x2=t,this._y0=this._y1,this._y1=this._y2,this._y2=n}};var b_=function t(n){function e(t){return n?new __(t,n):new l_(t,0)}return e.alpha=function(n){return t(+n)},e}(.5);function m_(t,n){this._context=t,this._alpha=n}m_.prototype={areaStart:r_,areaEnd:r_,lineStart:function(){this._x0=this._x1=this._x2=this._x3=this._x4=this._x5=this._y0=this._y1=this._y2=this._y3=this._y4=this._y5=NaN,this._l01_a=this._l12_a=this._l23_a=this._l01_2a=this._l12_2a=this._l23_2a=this._point=0},lineEnd:function(){switch(this._point){case 1:this._context.moveTo(this._x3,this._y3),this._context.closePath();break;case 2:this._context.lineTo(this._x3,this._y3),this._context.closePath();break;case 3:this.point(this._x3,this._y3),this.point(this._x4,this._y4),this.point(this._x5,this._y5)}},point:function(t,n){if(t=+t,n=+n,this._point){var e=this._x2-t,r=this._y2-n;this._l23_a=Math.sqrt(this._l23_2a=Math.pow(e*e+r*r,this._alpha))}switch(this._point){case 0:this._point=1,this._x3=t,this._y3=n;break;case 1:this._point=2,this._context.moveTo(this._x4=t,this._y4=n);break;case 2:this._point=3,this._x5=t,this._y5=n;break;default:y_(this,t,n)}this._l01_a=this._l12_a,this._l12_a=this._l23_a,this._l01_2a=this._l12_2a,this._l12_2a=this._l23_2a,this._x0=this._x1,this._x1=this._x2,this._x2=t,this._y0=this._y1,this._y1=this._y2,this._y2=n}};var x_=function t(n){function e(t){return n?new m_(t,n):new d_(t,0)}return e.alpha=function(n){return t(+n)},e}(.5);function w_(t,n){this._context=t,this._alpha=n}w_.prototype={areaStart:function(){this._line=0},areaEnd:function(){this._line=NaN},lineStart:function(){this._x0=this._x1=this._x2=this._y0=this._y1=this._y2=NaN,this._l01_a=this._l12_a=this._l23_a=this._l01_2a=this._l12_2a=this._l23_2a=this._point=0},lineEnd:function(){(this._line||0!==this._line&&3===this._point)&&this._context.closePath(),this._line=1-this._line},point:function(t,n){if(t=+t,n=+n,this._point){var e=this._x2-t,r=this._y2-n;this._l23_a=Math.sqrt(this._l23_2a=Math.pow(e*e+r*r,this._alpha))}switch(this._point){case 0:this._point=1;break;case 1:this._point=2;break;case 2:this._point=3,this._line?this._context.lineTo(this._x2,this._y2):this._context.moveTo(this._x2,this._y2);break;case 3:this._point=4;default:y_(this,t,n)}this._l01_a=this._l12_a,this._l12_a=this._l23_a,this._l01_2a=this._l12_2a,this._l12_2a=this._l23_2a,this._x0=this._x1,this._x1=this._x2,this._x2=t,this._y0=this._y1,this._y1=this._y2,this._y2=n}};var M_=function t(n){function e(t){return n?new w_(t,n):new v_(t,0)}return e.alpha=function(n){return t(+n)},e}(.5);function N_(t){this._context=t}function A_(t){return t<0?-1:1}function T_(t,n,e){var r=t._x1-t._x0,i=n-t._x1,o=(t._y1-t._y0)/(r||i<0&&-0),a=(e-t._y1)/(i||r<0&&-0),u=(o*i+a*r)/(r+i);return(A_(o)+A_(a))*Math.min(Math.abs(o),Math.abs(a),.5*Math.abs(u))||0}function S_(t,n){var e=t._x1-t._x0;return e?(3*(t._y1-t._y0)/e-n)/2:n}function k_(t,n,e){var r=t._x0,i=t._y0,o=t._x1,a=t._y1,u=(o-r)/3;t._context.bezierCurveTo(r+u,i+u*n,o-u,a-u*e,o,a)}function E_(t){this._context=t}function C_(t){this._context=new P_(t)}function P_(t){this._context=t}function z_(t){this._context=t}function R_(t){var n,e,r=t.length-1,i=new Array(r),o=new Array(r),a=new Array(r);for(i[0]=0,o[0]=2,a[0]=t[0]+2*t[1],n=1;n<r-1;++n)i[n]=1,o[n]=4,a[n]=4*t[n]+2*t[n+1];for(i[r-1]=2,o[r-1]=7,a[r-1]=8*t[r-1]+t[r],n=1;n<r;++n)e=i[n]/o[n-1],o[n]-=e,a[n]-=e*a[n-1];for(i[r-1]=a[r-1]/o[r-1],n=r-2;n>=0;--n)i[n]=(a[n]-i[n+1])/o[n];for(o[r-1]=(t[r]+i[r-1])/2,n=0;n<r-1;++n)o[n]=2*t[n+1]-i[n+1];return[i,o]}function D_(t,n){this._context=t,this._t=n}function q_(t,n){if((i=t.length)>1)for(var e,r,i,o=1,a=t[n[0]],u=a.length;o<i;++o)for(r=a,a=t[n[o]],e=0;e<u;++e)a[e][1]+=a[e][0]=isNaN(r[e][1])?r[e][0]:r[e][1]}function L_(t){for(var n=t.length,e=new Array(n);--n>=0;)e[n]=n;return e}function U_(t,n){return t[n]}function O_(t){var n=t.map(B_);return L_(t).sort(function(t,e){return n[t]-n[e]})}function B_(t){for(var n,e=-1,r=0,i=t.length,o=-1/0;++e<i;)(n=+t[e][1])>o&&(o=n,r=e);return r}function Y_(t){var n=t.map(F_);return L_(t).sort(function(t,e){return n[t]-n[e]})}function F_(t){for(var n,e=0,r=-1,i=t.length;++r<i;)(n=+t[r][1])&&(e+=n);return e}function I_(t){return function(){return t}}function j_(t){return t[0]}function H_(t){return t[1]}function X_(){this._=null}function G_(t){t.U=t.C=t.L=t.R=t.P=t.N=null}function V_(t,n){var e=n,r=n.R,i=e.U;i?i.L===e?i.L=r:i.R=r:t._=r,r.U=i,e.U=r,e.R=r.L,e.R&&(e.R.U=e),r.L=e}function $_(t,n){var e=n,r=n.L,i=e.U;i?i.L===e?i.L=r:i.R=r:t._=r,r.U=i,e.U=r,e.L=r.R,e.L&&(e.L.U=e),r.R=e}function W_(t){for(;t.L;)t=t.L;return t}function Z_(t,n,e,r){var i=[null,null],o=mb.push(i)-1;return i.left=t,i.right=n,e&&J_(i,t,n,e),r&&J_(i,n,t,r),_b[t.index].halfedges.push(o),_b[n.index].halfedges.push(o),i}function Q_(t,n,e){var r=[n,e];return r.left=t,r}function J_(t,n,e,r){t[0]||t[1]?t.left===e?t[1]=r:t[0]=r:(t[0]=r,t.left=n,t.right=e)}function K_(t,n,e,r,i){var o,a=t[0],u=t[1],c=a[0],f=a[1],s=0,l=1,h=u[0]-c,d=u[1]-f;if(o=n-c,h||!(o>0)){if(o/=h,h<0){if(o<s)return;o<l&&(l=o)}else if(h>0){if(o>l)return;o>s&&(s=o)}if(o=r-c,h||!(o<0)){if(o/=h,h<0){if(o>l)return;o>s&&(s=o)}else if(h>0){if(o<s)return;o<l&&(l=o)}if(o=e-f,d||!(o>0)){if(o/=d,d<0){if(o<s)return;o<l&&(l=o)}else if(d>0){if(o>l)return;o>s&&(s=o)}if(o=i-f,d||!(o<0)){if(o/=d,d<0){if(o>l)return;o>s&&(s=o)}else if(d>0){if(o<s)return;o<l&&(l=o)}return!(s>0||l<1)||(s>0&&(t[0]=[c+s*h,f+s*d]),l<1&&(t[1]=[c+l*h,f+l*d]),!0)}}}}}function tb(t,n,e,r,i){var o=t[1];if(o)return!0;var a,u,c=t[0],f=t.left,s=t.right,l=f[0],h=f[1],d=s[0],p=s[1],v=(l+d)/2,g=(h+p)/2;if(p===h){if(v<n||v>=r)return;if(l>d){if(c){if(c[1]>=i)return}else c=[v,e];o=[v,i]}else{if(c){if(c[1]<e)return}else c=[v,i];o=[v,e]}}else if(u=g-(a=(l-d)/(p-h))*v,a<-1||a>1)if(l>d){if(c){if(c[1]>=i)return}else c=[(e-u)/a,e];o=[(i-u)/a,i]}else{if(c){if(c[1]<e)return}else c=[(i-u)/a,i];o=[(e-u)/a,e]}else if(h<p){if(c){if(c[0]>=r)return}else c=[n,a*n+u];o=[r,a*r+u]}else{if(c){if(c[0]<n)return}else c=[r,a*r+u];o=[n,a*n+u]}return t[0]=c,t[1]=o,!0}function nb(t,n){var e=t.site,r=n.left,i=n.right;return e===i&&(i=r,r=e),i?Math.atan2(i[1]-r[1],i[0]-r[0]):(e===r?(r=n[1],i=n[0]):(r=n[0],i=n[1]),Math.atan2(r[0]-i[0],i[1]-r[1]))}function eb(t,n){return n[+(n.left!==t.site)]}function rb(t,n){return n[+(n.left===t.site)]}N_.prototype={areaStart:r_,areaEnd:r_,lineStart:function(){this._point=0},lineEnd:function(){this._point&&this._context.closePath()},point:function(t,n){t=+t,n=+n,this._point?this._context.lineTo(t,n):(this._point=1,this._context.moveTo(t,n))}},E_.prototype={areaStart:function(){this._line=0},areaEnd:function(){this._line=NaN},lineStart:function(){this._x0=this._x1=this._y0=this._y1=this._t0=NaN,this._point=0},lineEnd:function(){switch(this._point){case 2:this._context.lineTo(this._x1,this._y1);break;case 3:k_(this,this._t0,S_(this,this._t0))}(this._line||0!==this._line&&1===this._point)&&this._context.closePath(),this._line=1-this._line},point:function(t,n){var e=NaN;if(n=+n,(t=+t)!==this._x1||n!==this._y1){switch(this._point){case 0:this._point=1,this._line?this._context.lineTo(t,n):this._context.moveTo(t,n);break;case 1:this._point=2;break;case 2:this._point=3,k_(this,S_(this,e=T_(this,t,n)),e);break;default:k_(this,this._t0,e=T_(this,t,n))}this._x0=this._x1,this._x1=t,this._y0=this._y1,this._y1=n,this._t0=e}}},(C_.prototype=Object.create(E_.prototype)).point=function(t,n){E_.prototype.point.call(this,n,t)},P_.prototype={moveTo:function(t,n){this._context.moveTo(n,t)},closePath:function(){this._context.closePath()},lineTo:function(t,n){this._context.lineTo(n,t)},bezierCurveTo:function(t,n,e,r,i,o){this._context.bezierCurveTo(n,t,r,e,o,i)}},z_.prototype={areaStart:function(){this._line=0},areaEnd:function(){this._line=NaN},lineStart:function(){this._x=[],this._y=[]},lineEnd:function(){var t=this._x,n=this._y,e=t.length;if(e)if(this._line?this._context.lineTo(t[0],n[0]):this._context.moveTo(t[0],n[0]),2===e)this._context.lineTo(t[1],n[1]);else for(var r=R_(t),i=R_(n),o=0,a=1;a<e;++o,++a)this._context.bezierCurveTo(r[0][o],i[0][o],r[1][o],i[1][o],t[a],n[a]);(this._line||0!==this._line&&1===e)&&this._context.closePath(),this._line=1-this._line,this._x=this._y=null},point:function(t,n){this._x.push(+t),this._y.push(+n)}},D_.prototype={areaStart:function(){this._line=0},areaEnd:function(){this._line=NaN},lineStart:function(){this._x=this._y=NaN,this._point=0},lineEnd:function(){0<this._t&&this._t<1&&2===this._point&&this._context.lineTo(this._x,this._y),(this._line||0!==this._line&&1===this._point)&&this._context.closePath(),this._line>=0&&(this._t=1-this._t,this._line=1-this._line)},point:function(t,n){switch(t=+t,n=+n,this._point){case 0:this._point=1,this._line?this._context.lineTo(t,n):this._context.moveTo(t,n);break;case 1:this._point=2;default:if(this._t<=0)this._context.lineTo(this._x,n),this._context.lineTo(t,n);else{var e=this._x*(1-this._t)+t*this._t;this._context.lineTo(e,this._y),this._context.lineTo(e,n)}}this._x=t,this._y=n}},X_.prototype={constructor:X_,insert:function(t,n){var e,r,i;if(t){if(n.P=t,n.N=t.N,t.N&&(t.N.P=n),t.N=n,t.R){for(t=t.R;t.L;)t=t.L;t.L=n}else t.R=n;e=t}else this._?(t=W_(this._),n.P=null,n.N=t,t.P=t.L=n,e=t):(n.P=n.N=null,this._=n,e=null);for(n.L=n.R=null,n.U=e,n.C=!0,t=n;e&&e.C;)e===(r=e.U).L?(i=r.R)&&i.C?(e.C=i.C=!1,r.C=!0,t=r):(t===e.R&&(V_(this,e),e=(t=e).U),e.C=!1,r.C=!0,$_(this,r)):(i=r.L)&&i.C?(e.C=i.C=!1,r.C=!0,t=r):(t===e.L&&($_(this,e),e=(t=e).U),e.C=!1,r.C=!0,V_(this,r)),e=t.U;this._.C=!1},remove:function(t){t.N&&(t.N.P=t.P),t.P&&(t.P.N=t.N),t.N=t.P=null;var n,e,r,i=t.U,o=t.L,a=t.R;if(e=o?a?W_(a):o:a,i?i.L===t?i.L=e:i.R=e:this._=e,o&&a?(r=e.C,e.C=t.C,e.L=o,o.U=e,e!==a?(i=e.U,e.U=t.U,t=e.R,i.L=t,e.R=a,a.U=e):(e.U=i,i=e,t=e.R)):(r=t.C,t=e),t&&(t.U=i),!r)if(t&&t.C)t.C=!1;else{do{if(t===this._)break;if(t===i.L){if((n=i.R).C&&(n.C=!1,i.C=!0,V_(this,i),n=i.R),n.L&&n.L.C||n.R&&n.R.C){n.R&&n.R.C||(n.L.C=!1,n.C=!0,$_(this,n),n=i.R),n.C=i.C,i.C=n.R.C=!1,V_(this,i),t=this._;break}}else if((n=i.L).C&&(n.C=!1,i.C=!0,$_(this,i),n=i.L),n.L&&n.L.C||n.R&&n.R.C){n.L&&n.L.C||(n.R.C=!1,n.C=!0,V_(this,n),n=i.L),n.C=i.C,i.C=n.L.C=!1,$_(this,i),t=this._;break}n.C=!0,t=i,i=i.U}while(!t.C);t&&(t.C=!1)}}};var ib,ob=[];function ab(){G_(this),this.x=this.y=this.arc=this.site=this.cy=null}function ub(t){var n=t.P,e=t.N;if(n&&e){var r=n.site,i=t.site,o=e.site;if(r!==o){var a=i[0],u=i[1],c=r[0]-a,f=r[1]-u,s=o[0]-a,l=o[1]-u,h=2*(c*l-f*s);if(!(h>=-wb)){var d=c*c+f*f,p=s*s+l*l,v=(l*d-f*p)/h,g=(c*p-s*d)/h,y=ob.pop()||new ab;y.arc=t,y.site=i,y.x=v+a,y.y=(y.cy=g+u)+Math.sqrt(v*v+g*g),t.circle=y;for(var _=null,b=bb._;b;)if(y.y<b.y||y.y===b.y&&y.x<=b.x){if(!b.L){_=b.P;break}b=b.L}else{if(!b.R){_=b;break}b=b.R}bb.insert(_,y),_||(ib=y)}}}}function cb(t){var n=t.circle;n&&(n.P||(ib=n.N),bb.remove(n),ob.push(n),G_(n),t.circle=null)}var fb=[];function sb(){G_(this),this.edge=this.site=this.circle=null}function lb(t){var n=fb.pop()||new sb;return n.site=t,n}function hb(t){cb(t),yb.remove(t),fb.push(t),G_(t)}function db(t){var n=t.circle,e=n.x,r=n.cy,i=[e,r],o=t.P,a=t.N,u=[t];hb(t);for(var c=o;c.circle&&Math.abs(e-c.circle.x)<xb&&Math.abs(r-c.circle.cy)<xb;)o=c.P,u.unshift(c),hb(c),c=o;u.unshift(c),cb(c);for(var f=a;f.circle&&Math.abs(e-f.circle.x)<xb&&Math.abs(r-f.circle.cy)<xb;)a=f.N,u.push(f),hb(f),f=a;u.push(f),cb(f);var s,l=u.length;for(s=1;s<l;++s)f=u[s],c=u[s-1],J_(f.edge,c.site,f.site,i);c=u[0],(f=u[l-1]).edge=Z_(c.site,f.site,null,i),ub(c),ub(f)}function pb(t){for(var n,e,r,i,o=t[0],a=t[1],u=yb._;u;)if((r=vb(u,a)-o)>xb)u=u.L;else{if(!((i=o-gb(u,a))>xb)){r>-xb?(n=u.P,e=u):i>-xb?(n=u,e=u.N):n=e=u;break}if(!u.R){n=u;break}u=u.R}!function(t){_b[t.index]={site:t,halfedges:[]}}(t);var c=lb(t);if(yb.insert(n,c),n||e){if(n===e)return cb(n),e=lb(n.site),yb.insert(c,e),c.edge=e.edge=Z_(n.site,c.site),ub(n),void ub(e);if(e){cb(n),cb(e);var f=n.site,s=f[0],l=f[1],h=t[0]-s,d=t[1]-l,p=e.site,v=p[0]-s,g=p[1]-l,y=2*(h*g-d*v),_=h*h+d*d,b=v*v+g*g,m=[(g*_-d*b)/y+s,(h*b-v*_)/y+l];J_(e.edge,f,p,m),c.edge=Z_(f,t,null,m),e.edge=Z_(t,p,null,m),ub(n),ub(e)}else c.edge=Z_(n.site,c.site)}}function vb(t,n){var e=t.site,r=e[0],i=e[1],o=i-n;if(!o)return r;var a=t.P;if(!a)return-1/0;var u=(e=a.site)[0],c=e[1],f=c-n;if(!f)return u;var s=u-r,l=1/o-1/f,h=s/f;return l?(-h+Math.sqrt(h*h-2*l*(s*s/(-2*f)-c+f/2+i-o/2)))/l+r:(r+u)/2}function gb(t,n){var e=t.N;if(e)return vb(e,n);var r=t.site;return r[1]===n?r[0]:1/0}var yb,_b,bb,mb,xb=1e-6,wb=1e-12;function Mb(t,n){return n[1]-t[1]||n[0]-t[0]}function Nb(t,n){var e,r,i,o=t.sort(Mb).pop();for(mb=[],_b=new Array(t.length),yb=new X_,bb=new X_;;)if(i=ib,o&&(!i||o[1]<i.y||o[1]===i.y&&o[0]<i.x))o[0]===e&&o[1]===r||(pb(o),e=o[0],r=o[1]),o=t.pop();else{if(!i)break;db(i.arc)}if(function(){for(var t,n,e,r,i=0,o=_b.length;i<o;++i)if((t=_b[i])&&(r=(n=t.halfedges).length)){var a=new Array(r),u=new Array(r);for(e=0;e<r;++e)a[e]=e,u[e]=nb(t,mb[n[e]]);for(a.sort(function(t,n){return u[n]-u[t]}),e=0;e<r;++e)u[e]=n[a[e]];for(e=0;e<r;++e)n[e]=u[e]}}(),n){var a=+n[0][0],u=+n[0][1],c=+n[1][0],f=+n[1][1];!function(t,n,e,r){for(var i,o=mb.length;o--;)tb(i=mb[o],t,n,e,r)&&K_(i,t,n,e,r)&&(Math.abs(i[0][0]-i[1][0])>xb||Math.abs(i[0][1]-i[1][1])>xb)||delete mb[o]}(a,u,c,f),function(t,n,e,r){var i,o,a,u,c,f,s,l,h,d,p,v,g=_b.length,y=!0;for(i=0;i<g;++i)if(o=_b[i]){for(a=o.site,u=(c=o.halfedges).length;u--;)mb[c[u]]||c.splice(u,1);for(u=0,f=c.length;u<f;)p=(d=rb(o,mb[c[u]]))[0],v=d[1],l=(s=eb(o,mb[c[++u%f]]))[0],h=s[1],(Math.abs(p-l)>xb||Math.abs(v-h)>xb)&&(c.splice(u,0,mb.push(Q_(a,d,Math.abs(p-t)<xb&&r-v>xb?[t,Math.abs(l-t)<xb?h:r]:Math.abs(v-r)<xb&&e-p>xb?[Math.abs(h-r)<xb?l:e,r]:Math.abs(p-e)<xb&&v-n>xb?[e,Math.abs(l-e)<xb?h:n]:Math.abs(v-n)<xb&&p-t>xb?[Math.abs(h-n)<xb?l:t,n]:null))-1),++f);f&&(y=!1)}if(y){var _,b,m,x=1/0;for(i=0,y=null;i<g;++i)(o=_b[i])&&(m=(_=(a=o.site)[0]-t)*_+(b=a[1]-n)*b)<x&&(x=m,y=o);if(y){var w=[t,n],M=[t,r],N=[e,r],A=[e,n];y.halfedges.push(mb.push(Q_(a=y.site,w,M))-1,mb.push(Q_(a,M,N))-1,mb.push(Q_(a,N,A))-1,mb.push(Q_(a,A,w))-1)}}for(i=0;i<g;++i)(o=_b[i])&&(o.halfedges.length||delete _b[i])}(a,u,c,f)}this.edges=mb,this.cells=_b,yb=bb=mb=_b=null}function Ab(t){return function(){return t}}function Tb(t,n,e){this.target=t,this.type=n,this.transform=e}function Sb(t,n,e){this.k=t,this.x=n,this.y=e}Nb.prototype={constructor:Nb,polygons:function(){var t=this.edges;return this.cells.map(function(n){var e=n.halfedges.map(function(e){return eb(n,t[e])});return e.data=n.site.data,e})},triangles:function(){var t=[],n=this.edges;return this.cells.forEach(function(e,r){if(o=(i=e.halfedges).length)for(var i,o,a,u,c,f,s=e.site,l=-1,h=n[i[o-1]],d=h.left===s?h.right:h.left;++l<o;)a=d,d=(h=n[i[l]]).left===s?h.right:h.left,a&&d&&r<a.index&&r<d.index&&(c=a,f=d,((u=s)[0]-f[0])*(c[1]-u[1])-(u[0]-c[0])*(f[1]-u[1])<0)&&t.push([s.data,a.data,d.data])}),t},links:function(){return this.edges.filter(function(t){return t.right}).map(function(t){return{source:t.left.data,target:t.right.data}})},find:function(t,n,e){for(var r,i,o=this,a=o._found||0,u=o.cells.length;!(i=o.cells[a]);)if(++a>=u)return null;var c=t-i.site[0],f=n-i.site[1],s=c*c+f*f;do{i=o.cells[r=a],a=null,i.halfedges.forEach(function(e){var r=o.edges[e],u=r.left;if(u!==i.site&&u||(u=r.right)){var c=t-u[0],f=n-u[1],l=c*c+f*f;l<s&&(s=l,a=u.index)}})}while(null!==a);return o._found=r,null==e||s<=e*e?i.site:null}},Sb.prototype={constructor:Sb,scale:function(t){return 1===t?this:new Sb(this.k*t,this.x,this.y)},translate:function(t,n){return 0===t&0===n?this:new Sb(this.k,this.x+this.k*t,this.y+this.k*n)},apply:function(t){return[t[0]*this.k+this.x,t[1]*this.k+this.y]},applyX:function(t){return t*this.k+this.x},applyY:function(t){return t*this.k+this.y},invert:function(t){return[(t[0]-this.x)/this.k,(t[1]-this.y)/this.k]},invertX:function(t){return(t-this.x)/this.k},invertY:function(t){return(t-this.y)/this.k},rescaleX:function(t){return t.copy().domain(t.range().map(this.invertX,this).map(t.invert,t))},rescaleY:function(t){return t.copy().domain(t.range().map(this.invertY,this).map(t.invert,t))},toString:function(){return"translate("+this.x+","+this.y+") scale("+this.k+")"}};var kb=new Sb(1,0,0);function Eb(t){return t.__zoom||kb}function Cb(){t.event.stopImmediatePropagation()}function Pb(){t.event.preventDefault(),t.event.stopImmediatePropagation()}function zb(){return!t.event.button}function Rb(){var t,n,e=this;return e instanceof SVGElement?(t=(e=e.ownerSVGElement||e).width.baseVal.value,n=e.height.baseVal.value):(t=e.clientWidth,n=e.clientHeight),[[0,0],[t,n]]}function Db(){return this.__zoom||kb}function qb(){return-t.event.deltaY*(t.event.deltaMode?120:1)/500}function Lb(){return"ontouchstart"in this}function Ub(t,n,e){var r=t.invertX(n[0][0])-e[0][0],i=t.invertX(n[1][0])-e[1][0],o=t.invertY(n[0][1])-e[0][1],a=t.invertY(n[1][1])-e[1][1];return t.translate(i>r?(r+i)/2:Math.min(0,r)||Math.max(0,i),a>o?(o+a)/2:Math.min(0,o)||Math.max(0,a))}Eb.prototype=Sb.prototype,t.version="5.9.2",t.bisect=i,t.bisectRight=i,t.bisectLeft=o,t.ascending=n,t.bisector=e,t.cross=function(t,n,e){var r,i,o,u,c=t.length,f=n.length,s=new Array(c*f);for(null==e&&(e=a),r=o=0;r<c;++r)for(u=t[r],i=0;i<f;++i,++o)s[o]=e(u,n[i]);return s},t.descending=function(t,n){return n<t?-1:n>t?1:n>=t?0:NaN},t.deviation=f,t.extent=s,t.histogram=function(){var t=v,n=s,e=M;function r(r){var o,a,u=r.length,c=new Array(u);for(o=0;o<u;++o)c[o]=t(r[o],o,r);var f=n(c),s=f[0],l=f[1],h=e(c,s,l);Array.isArray(h)||(h=w(s,l,h),h=g(Math.ceil(s/h)*h,l,h));for(var d=h.length;h[0]<=s;)h.shift(),--d;for(;h[d-1]>l;)h.pop(),--d;var p,v=new Array(d+1);for(o=0;o<=d;++o)(p=v[o]=[]).x0=o>0?h[o-1]:s,p.x1=o<d?h[o]:l;for(o=0;o<u;++o)s<=(a=c[o])&&a<=l&&v[i(h,a,0,d)].push(r[o]);return v}return r.value=function(n){return arguments.length?(t="function"==typeof n?n:p(n),r):t},r.domain=function(t){return arguments.length?(n="function"==typeof t?t:p([t[0],t[1]]),r):n},r.thresholds=function(t){return arguments.length?(e="function"==typeof t?t:Array.isArray(t)?p(h.call(t)):p(t),r):e},r},t.thresholdFreedmanDiaconis=function(t,e,r){return t=d.call(t,u).sort(n),Math.ceil((r-e)/(2*(N(t,.75)-N(t,.25))*Math.pow(t.length,-1/3)))},t.thresholdScott=function(t,n,e){return Math.ceil((e-n)/(3.5*f(t)*Math.pow(t.length,-1/3)))},t.thresholdSturges=M,t.max=A,t.mean=function(t,n){var e,r=t.length,i=r,o=-1,a=0;if(null==n)for(;++o<r;)isNaN(e=u(t[o]))?--i:a+=e;else for(;++o<r;)isNaN(e=u(n(t[o],o,t)))?--i:a+=e;if(i)return a/i},t.median=function(t,e){var r,i=t.length,o=-1,a=[];if(null==e)for(;++o<i;)isNaN(r=u(t[o]))||a.push(r);else for(;++o<i;)isNaN(r=u(e(t[o],o,t)))||a.push(r);return N(a.sort(n),.5)},t.merge=T,t.min=S,t.pairs=function(t,n){null==n&&(n=a);for(var e=0,r=t.length-1,i=t[0],o=new Array(r<0?0:r);e<r;)o[e]=n(i,i=t[++e]);return o},t.permute=function(t,n){for(var e=n.length,r=new Array(e);e--;)r[e]=t[n[e]];return r},t.quantile=N,t.range=g,t.scan=function(t,e){if(r=t.length){var r,i,o=0,a=0,u=t[a];for(null==e&&(e=n);++o<r;)(e(i=t[o],u)<0||0!==e(u,u))&&(u=i,a=o);return 0===e(u,u)?a:void 0}},t.shuffle=function(t,n,e){for(var r,i,o=(null==e?t.length:e)-(n=null==n?0:+n);o;)i=Math.random()*o--|0,r=t[o+n],t[o+n]=t[i+n],t[i+n]=r;return t},t.sum=function(t,n){var e,r=t.length,i=-1,o=0;if(null==n)for(;++i<r;)(e=+t[i])&&(o+=e);else for(;++i<r;)(e=+n(t[i],i,t))&&(o+=e);return o},t.ticks=m,t.tickIncrement=x,t.tickStep=w,t.transpose=k,t.variance=c,t.zip=function(){return k(arguments)},t.axisTop=function(t){return Y(z,t)},t.axisRight=function(t){return Y(R,t)},t.axisBottom=function(t){return Y(D,t)},t.axisLeft=function(t){return Y(q,t)},t.brush=function(){return Pi(mi)},t.brushX=function(){return Pi(_i)},t.brushY=function(){return Pi(bi)},t.brushSelection=function(t){var n=t.__brush;return n?n.dim.output(n.selection):null},t.chord=function(){var t=0,n=null,e=null,r=null;function i(i){var o,a,u,c,f,s,l=i.length,h=[],d=g(l),p=[],v=[],y=v.groups=new Array(l),_=new Array(l*l);for(o=0,f=-1;++f<l;){for(a=0,s=-1;++s<l;)a+=i[f][s];h.push(a),p.push(g(l)),o+=a}for(n&&d.sort(function(t,e){return n(h[t],h[e])}),e&&p.forEach(function(t,n){t.sort(function(t,r){return e(i[n][t],i[n][r])})}),c=(o=Ui(0,Li-t*l)/o)?t:Li/l,a=0,f=-1;++f<l;){for(u=a,s=-1;++s<l;){var b=d[f],m=p[b][s],x=i[b][m],w=a,M=a+=x*o;_[m*l+b]={index:b,subindex:m,startAngle:w,endAngle:M,value:x}}y[b]={index:b,startAngle:u,endAngle:a,value:h[b]},a+=c}for(f=-1;++f<l;)for(s=f-1;++s<l;){var N=_[s*l+f],A=_[f*l+s];(N.value||A.value)&&v.push(N.value<A.value?{source:A,target:N}:{source:N,target:A})}return r?v.sort(r):v}return i.padAngle=function(n){return arguments.length?(t=Ui(0,n),i):t},i.sortGroups=function(t){return arguments.length?(n=t,i):n},i.sortSubgroups=function(t){return arguments.length?(e=t,i):e},i.sortChords=function(t){return arguments.length?(null==t?r=null:(n=t,r=function(t,e){return n(t.source.value+t.target.value,e.source.value+e.target.value)})._=t,i):r&&r._;var n},i},t.ribbon=function(){var t=Xi,n=Gi,e=Vi,r=$i,i=Wi,o=null;function a(){var a,u=Oi.call(arguments),c=t.apply(this,u),f=n.apply(this,u),s=+e.apply(this,(u[0]=c,u)),l=r.apply(this,u)-qi,h=i.apply(this,u)-qi,d=s*zi(l),p=s*Ri(l),v=+e.apply(this,(u[0]=f,u)),g=r.apply(this,u)-qi,y=i.apply(this,u)-qi;if(o||(o=a=Hi()),o.moveTo(d,p),o.arc(0,0,s,l,h),l===g&&h===y||(o.quadraticCurveTo(0,0,v*zi(g),v*Ri(g)),o.arc(0,0,v,g,y)),o.quadraticCurveTo(0,0,d,p),o.closePath(),a)return o=null,a+""||null}return a.radius=function(t){return arguments.length?(e="function"==typeof t?t:Bi(+t),a):e},a.startAngle=function(t){return arguments.length?(r="function"==typeof t?t:Bi(+t),a):r},a.endAngle=function(t){return arguments.length?(i="function"==typeof t?t:Bi(+t),a):i},a.source=function(n){return arguments.length?(t=n,a):t},a.target=function(t){return arguments.length?(n=t,a):n},a.context=function(t){return arguments.length?(o=null==t?null:t,a):o},a},t.nest=function(){var t,n,e,r=[],i=[];function o(e,i,a,u){if(i>=r.length)return null!=t&&e.sort(t),null!=n?n(e):e;for(var c,f,s,l=-1,h=e.length,d=r[i++],p=Qi(),v=a();++l<h;)(s=p.get(c=d(f=e[l])+""))?s.push(f):p.set(c,[f]);return p.each(function(t,n){u(v,n,o(t,i,a,u))}),v}return e={object:function(t){return o(t,0,Ji,Ki)},map:function(t){return o(t,0,to,no)},entries:function(t){return function t(e,o){if(++o>r.length)return e;var a,u=i[o-1];return null!=n&&o>=r.length?a=e.entries():(a=[],e.each(function(n,e){a.push({key:e,values:t(n,o)})})),null!=u?a.sort(function(t,n){return u(t.key,n.key)}):a}(o(t,0,to,no),0)},key:function(t){return r.push(t),e},sortKeys:function(t){return i[r.length-1]=t,e},sortValues:function(n){return t=n,e},rollup:function(t){return n=t,e}}},t.set=io,t.map=Qi,t.keys=function(t){var n=[];for(var e in t)n.push(e);return n},t.values=function(t){var n=[];for(var e in t)n.push(t[e]);return n},t.entries=function(t){var n=[];for(var e in t)n.push({key:e,value:t[e]});return n},t.color=hn,t.rgb=gn,t.hsl=mn,t.lab=Rn,t.hcl=Yn,t.lch=function(t,n,e,r){return 1===arguments.length?Bn(t):new Fn(e,n,t,null==r?1:r)},t.gray=function(t,n){return new Dn(t,0,0,null==n?1:n)},t.cubehelix=Zn,t.contours=po,t.contourDensity=function(){var t=yo,n=_o,e=bo,r=960,i=500,o=20,a=2,u=3*o,c=r+2*u>>a,f=i+2*u>>a,s=uo(20);function l(r){var i=new Float32Array(c*f),l=new Float32Array(c*f);r.forEach(function(r,o,s){var l=+t(r,o,s)+u>>a,h=+n(r,o,s)+u>>a,d=+e(r,o,s);l>=0&&l<c&&h>=0&&h<f&&(i[l+h*c]+=d)}),vo({width:c,height:f,data:i},{width:c,height:f,data:l},o>>a),go({width:c,height:f,data:l},{width:c,height:f,data:i},o>>a),vo({width:c,height:f,data:i},{width:c,height:f,data:l},o>>a),go({width:c,height:f,data:l},{width:c,height:f,data:i},o>>a),vo({width:c,height:f,data:i},{width:c,height:f,data:l},o>>a),go({width:c,height:f,data:l},{width:c,height:f,data:i},o>>a);var d=s(i);if(!Array.isArray(d)){var p=A(i);d=w(0,p,d),(d=g(0,Math.floor(p/d)*d,d)).shift()}return po().thresholds(d).size([c,f])(i).map(h)}function h(t){return t.value*=Math.pow(2,-2*a),t.coordinates.forEach(d),t}function d(t){t.forEach(p)}function p(t){t.forEach(v)}function v(t){t[0]=t[0]*Math.pow(2,a)-u,t[1]=t[1]*Math.pow(2,a)-u}function y(){return c=r+2*(u=3*o)>>a,f=i+2*u>>a,l}return l.x=function(n){return arguments.length?(t="function"==typeof n?n:uo(+n),l):t},l.y=function(t){return arguments.length?(n="function"==typeof t?t:uo(+t),l):n},l.weight=function(t){return arguments.length?(e="function"==typeof t?t:uo(+t),l):e},l.size=function(t){if(!arguments.length)return[r,i];var n=Math.ceil(t[0]),e=Math.ceil(t[1]);if(!(n>=0||n>=0))throw new Error("invalid size");return r=n,i=e,y()},l.cellSize=function(t){if(!arguments.length)return 1<<a;if(!((t=+t)>=1))throw new Error("invalid cell size");return a=Math.floor(Math.log(t)/Math.LN2),y()},l.thresholds=function(t){return arguments.length?(s="function"==typeof t?t:Array.isArray(t)?uo(oo.call(t)):uo(t),l):s},l.bandwidth=function(t){if(!arguments.length)return Math.sqrt(o*(o+1));if(!((t=+t)>=0))throw new Error("invalid bandwidth");return o=Math.round((Math.sqrt(4*t*t+1)-1)/2),y()},l},t.dispatch=I,t.drag=function(){var n,e,r,i,o=Gt,a=Vt,u=$t,c=Wt,f={},s=I("start","drag","end"),l=0,h=0;function d(t){t.on("mousedown.drag",p).filter(c).on("touchstart.drag",y).on("touchmove.drag",_).on("touchend.drag touchcancel.drag",b).style("touch-action","none").style("-webkit-tap-highlight-color","rgba(0,0,0,0)")}function p(){if(!i&&o.apply(this,arguments)){var u=m("mouse",a.apply(this,arguments),Ot,this,arguments);u&&(zt(t.event.view).on("mousemove.drag",v,!0).on("mouseup.drag",g,!0),It(t.event.view),Yt(),r=!1,n=t.event.clientX,e=t.event.clientY,u("start"))}}function v(){if(Ft(),!r){var i=t.event.clientX-n,o=t.event.clientY-e;r=i*i+o*o>h}f.mouse("drag")}function g(){zt(t.event.view).on("mousemove.drag mouseup.drag",null),jt(t.event.view,r),Ft(),f.mouse("end")}function y(){if(o.apply(this,arguments)){var n,e,r=t.event.changedTouches,i=a.apply(this,arguments),u=r.length;for(n=0;n<u;++n)(e=m(r[n].identifier,i,Bt,this,arguments))&&(Yt(),e("start"))}}function _(){var n,e,r=t.event.changedTouches,i=r.length;for(n=0;n<i;++n)(e=f[r[n].identifier])&&(Ft(),e("drag"))}function b(){var n,e,r=t.event.changedTouches,o=r.length;for(i&&clearTimeout(i),i=setTimeout(function(){i=null},500),n=0;n<o;++n)(e=f[r[n].identifier])&&(Yt(),e("end"))}function m(n,e,r,i,o){var a,c,h,p=r(e,n),v=s.copy();if(St(new Xt(d,"beforestart",a,n,l,p[0],p[1],0,0,v),function(){return null!=(t.event.subject=a=u.apply(i,o))&&(c=a.x-p[0]||0,h=a.y-p[1]||0,!0)}))return function t(u){var s,g=p;switch(u){case"start":f[n]=t,s=l++;break;case"end":delete f[n],--l;case"drag":p=r(e,n),s=l}St(new Xt(d,u,a,n,s,p[0]+c,p[1]+h,p[0]-g[0],p[1]-g[1],v),v.apply,v,[u,i,o])}}return d.filter=function(t){return arguments.length?(o="function"==typeof t?t:Ht(!!t),d):o},d.container=function(t){return arguments.length?(a="function"==typeof t?t:Ht(t),d):a},d.subject=function(t){return arguments.length?(u="function"==typeof t?t:Ht(t),d):u},d.touchable=function(t){return arguments.length?(c="function"==typeof t?t:Ht(!!t),d):c},d.on=function(){var t=s.on.apply(s,arguments);return t===s?d:t},d.clickDistance=function(t){return arguments.length?(h=(t=+t)*t,d):Math.sqrt(h)},d},t.dragDisable=It,t.dragEnable=jt,t.dsvFormat=Eo,t.csvParse=Po,t.csvParseRows=zo,t.csvFormat=Ro,t.csvFormatBody=Do,t.csvFormatRows=qo,t.tsvParse=Uo,t.tsvParseRows=Oo,t.tsvFormat=Bo,t.tsvFormatBody=Yo,t.tsvFormatRows=Fo,t.autoType=function(t){for(var n in t){var e,r=t[n].trim();if(r)if("true"===r)r=!0;else if("false"===r)r=!1;else if("NaN"===r)r=NaN;else if(isNaN(e=+r)){if(!/^([-+]\d{2})?\d{4}(-\d{2}(-\d{2})?)?(T\d{2}:\d{2}(:\d{2}(\.\d{3})?)?(Z|[-+]\d{2}:\d{2})?)?$/.test(r))continue;r=new Date(r)}else r=e;else r=null;t[n]=r}return t},t.easeLinear=function(t){return+t},t.easeQuad=Rr,t.easeQuadIn=function(t){return t*t},t.easeQuadOut=function(t){return t*(2-t)},t.easeQuadInOut=Rr,t.easeCubic=Dr,t.easeCubicIn=function(t){return t*t*t},t.easeCubicOut=function(t){return--t*t*t+1},t.easeCubicInOut=Dr,t.easePoly=Ur,t.easePolyIn=qr,t.easePolyOut=Lr,t.easePolyInOut=Ur,t.easeSin=Yr,t.easeSinIn=function(t){return 1-Math.cos(t*Br)},t.easeSinOut=function(t){return Math.sin(t*Br)},t.easeSinInOut=Yr,t.easeExp=Fr,t.easeExpIn=function(t){return Math.pow(2,10*t-10)},t.easeExpOut=function(t){return 1-Math.pow(2,-10*t)},t.easeExpInOut=Fr,t.easeCircle=Ir,t.easeCircleIn=function(t){return 1-Math.sqrt(1-t*t)},t.easeCircleOut=function(t){return Math.sqrt(1- --t*t)},t.easeCircleInOut=Ir,t.easeBounce=Kr,t.easeBounceIn=function(t){return 1-Kr(1-t)},t.easeBounceOut=Kr,t.easeBounceInOut=function(t){return((t*=2)<=1?1-Kr(1-t):Kr(t-1)+1)/2},t.easeBack=ei,t.easeBackIn=ti,t.easeBackOut=ni,t.easeBackInOut=ei,t.easeElastic=oi,t.easeElasticIn=ii,t.easeElasticOut=oi,t.easeElasticInOut=ai,t.blob=function(t,n){return fetch(t,n).then(Io)},t.buffer=function(t,n){return fetch(t,n).then(jo)},t.dsv=function(t,n,e,r){3===arguments.length&&"function"==typeof e&&(r=e,e=void 0);var i=Eo(t);return Xo(n,e).then(function(t){return i.parse(t,r)})},t.csv=Vo,t.tsv=$o,t.image=function(t,n){return new Promise(function(e,r){var i=new Image;for(var o in n)i[o]=n[o];i.onerror=r,i.onload=function(){e(i)},i.src=t})},t.json=function(t,n){return fetch(t,n).then(Wo)},t.text=Xo,t.xml=Qo,t.html=Jo,t.svg=Ko,t.forceCenter=function(t,n){var e;function r(){var r,i,o=e.length,a=0,u=0;for(r=0;r<o;++r)a+=(i=e[r]).x,u+=i.y;for(a=a/o-t,u=u/o-n,r=0;r<o;++r)(i=e[r]).x-=a,i.y-=u}return null==t&&(t=0),null==n&&(n=0),r.initialize=function(t){e=t},r.x=function(n){return arguments.length?(t=+n,r):t},r.y=function(t){return arguments.length?(n=+t,r):n},r},t.forceCollide=function(t){var n,e,r=1,i=1;function o(){for(var t,o,u,c,f,s,l,h=n.length,d=0;d<i;++d)for(o=aa(n,sa,la).visitAfter(a),t=0;t<h;++t)u=n[t],s=e[u.index],l=s*s,c=u.x+u.vx,f=u.y+u.vy,o.visit(p);function p(t,n,e,i,o){var a=t.data,h=t.r,d=s+h;if(!a)return n>c+d||i<c-d||e>f+d||o<f-d;if(a.index>u.index){var p=c-a.x-a.vx,v=f-a.y-a.vy,g=p*p+v*v;g<d*d&&(0===p&&(g+=(p=na())*p),0===v&&(g+=(v=na())*v),g=(d-(g=Math.sqrt(g)))/g*r,u.vx+=(p*=g)*(d=(h*=h)/(l+h)),u.vy+=(v*=g)*d,a.vx-=p*(d=1-d),a.vy-=v*d)}}}function a(t){if(t.data)return t.r=e[t.data.index];for(var n=t.r=0;n<4;++n)t[n]&&t[n].r>t.r&&(t.r=t[n].r)}function u(){if(n){var r,i,o=n.length;for(e=new Array(o),r=0;r<o;++r)i=n[r],e[i.index]=+t(i,r,n)}}return"function"!=typeof t&&(t=ta(null==t?1:+t)),o.initialize=function(t){n=t,u()},o.iterations=function(t){return arguments.length?(i=+t,o):i},o.strength=function(t){return arguments.length?(r=+t,o):r},o.radius=function(n){return arguments.length?(t="function"==typeof n?n:ta(+n),u(),o):t},o},t.forceLink=function(t){var n,e,r,i,o,a=ha,u=function(t){return 1/Math.min(i[t.source.index],i[t.target.index])},c=ta(30),f=1;function s(r){for(var i=0,a=t.length;i<f;++i)for(var u,c,s,l,h,d,p,v=0;v<a;++v)c=(u=t[v]).source,l=(s=u.target).x+s.vx-c.x-c.vx||na(),h=s.y+s.vy-c.y-c.vy||na(),l*=d=((d=Math.sqrt(l*l+h*h))-e[v])/d*r*n[v],h*=d,s.vx-=l*(p=o[v]),s.vy-=h*p,c.vx+=l*(p=1-p),c.vy+=h*p}function l(){if(r){var u,c,f=r.length,s=t.length,l=Qi(r,a);for(u=0,i=new Array(f);u<s;++u)(c=t[u]).index=u,"object"!=typeof c.source&&(c.source=da(l,c.source)),"object"!=typeof c.target&&(c.target=da(l,c.target)),i[c.source.index]=(i[c.source.index]||0)+1,i[c.target.index]=(i[c.target.index]||0)+1;for(u=0,o=new Array(s);u<s;++u)c=t[u],o[u]=i[c.source.index]/(i[c.source.index]+i[c.target.index]);n=new Array(s),h(),e=new Array(s),d()}}function h(){if(r)for(var e=0,i=t.length;e<i;++e)n[e]=+u(t[e],e,t)}function d(){if(r)for(var n=0,i=t.length;n<i;++n)e[n]=+c(t[n],n,t)}return null==t&&(t=[]),s.initialize=function(t){r=t,l()},s.links=function(n){return arguments.length?(t=n,l(),s):t},s.id=function(t){return arguments.length?(a=t,s):a},s.iterations=function(t){return arguments.length?(f=+t,s):f},s.strength=function(t){return arguments.length?(u="function"==typeof t?t:ta(+t),h(),s):u},s.distance=function(t){return arguments.length?(c="function"==typeof t?t:ta(+t),d(),s):c},s},t.forceManyBody=function(){var t,n,e,r,i=ta(-30),o=1,a=1/0,u=.81;function c(r){var i,o=t.length,a=aa(t,pa,va).visitAfter(s);for(e=r,i=0;i<o;++i)n=t[i],a.visit(l)}function f(){if(t){var n,e,o=t.length;for(r=new Array(o),n=0;n<o;++n)e=t[n],r[e.index]=+i(e,n,t)}}function s(t){var n,e,i,o,a,u=0,c=0;if(t.length){for(i=o=a=0;a<4;++a)(n=t[a])&&(e=Math.abs(n.value))&&(u+=n.value,c+=e,i+=e*n.x,o+=e*n.y);t.x=i/c,t.y=o/c}else{(n=t).x=n.data.x,n.y=n.data.y;do{u+=r[n.data.index]}while(n=n.next)}t.value=u}function l(t,i,c,f){if(!t.value)return!0;var s=t.x-n.x,l=t.y-n.y,h=f-i,d=s*s+l*l;if(h*h/u<d)return d<a&&(0===s&&(d+=(s=na())*s),0===l&&(d+=(l=na())*l),d<o&&(d=Math.sqrt(o*d)),n.vx+=s*t.value*e/d,n.vy+=l*t.value*e/d),!0;if(!(t.length||d>=a)){(t.data!==n||t.next)&&(0===s&&(d+=(s=na())*s),0===l&&(d+=(l=na())*l),d<o&&(d=Math.sqrt(o*d)));do{t.data!==n&&(h=r[t.data.index]*e/d,n.vx+=s*h,n.vy+=l*h)}while(t=t.next)}}return c.initialize=function(n){t=n,f()},c.strength=function(t){return arguments.length?(i="function"==typeof t?t:ta(+t),f(),c):i},c.distanceMin=function(t){return arguments.length?(o=t*t,c):Math.sqrt(o)},c.distanceMax=function(t){return arguments.length?(a=t*t,c):Math.sqrt(a)},c.theta=function(t){return arguments.length?(u=t*t,c):Math.sqrt(u)},c},t.forceRadial=function(t,n,e){var r,i,o,a=ta(.1);function u(t){for(var a=0,u=r.length;a<u;++a){var c=r[a],f=c.x-n||1e-6,s=c.y-e||1e-6,l=Math.sqrt(f*f+s*s),h=(o[a]-l)*i[a]*t/l;c.vx+=f*h,c.vy+=s*h}}function c(){if(r){var n,e=r.length;for(i=new Array(e),o=new Array(e),n=0;n<e;++n)o[n]=+t(r[n],n,r),i[n]=isNaN(o[n])?0:+a(r[n],n,r)}}return"function"!=typeof t&&(t=ta(+t)),null==n&&(n=0),null==e&&(e=0),u.initialize=function(t){r=t,c()},u.strength=function(t){return arguments.length?(a="function"==typeof t?t:ta(+t),c(),u):a},u.radius=function(n){return arguments.length?(t="function"==typeof n?n:ta(+n),c(),u):t},u.x=function(t){return arguments.length?(n=+t,u):n},u.y=function(t){return arguments.length?(e=+t,u):e},u},t.forceSimulation=function(t){var n,e=1,r=.001,i=1-Math.pow(r,1/300),o=0,a=.6,u=Qi(),c=ir(s),f=I("tick","end");function s(){l(),f.call("tick",n),e<r&&(c.stop(),f.call("end",n))}function l(r){var c,f,s=t.length;void 0===r&&(r=1);for(var l=0;l<r;++l)for(e+=(o-e)*i,u.each(function(t){t(e)}),c=0;c<s;++c)null==(f=t[c]).fx?f.x+=f.vx*=a:(f.x=f.fx,f.vx=0),null==f.fy?f.y+=f.vy*=a:(f.y=f.fy,f.vy=0);return n}function h(){for(var n,e=0,r=t.length;e<r;++e){if((n=t[e]).index=e,null!=n.fx&&(n.x=n.fx),null!=n.fy&&(n.y=n.fy),isNaN(n.x)||isNaN(n.y)){var i=ga*Math.sqrt(e),o=e*ya;n.x=i*Math.cos(o),n.y=i*Math.sin(o)}(isNaN(n.vx)||isNaN(n.vy))&&(n.vx=n.vy=0)}}function d(n){return n.initialize&&n.initialize(t),n}return null==t&&(t=[]),h(),n={tick:l,restart:function(){return c.restart(s),n},stop:function(){return c.stop(),n},nodes:function(e){return arguments.length?(t=e,h(),u.each(d),n):t},alpha:function(t){return arguments.length?(e=+t,n):e},alphaMin:function(t){return arguments.length?(r=+t,n):r},alphaDecay:function(t){return arguments.length?(i=+t,n):+i},alphaTarget:function(t){return arguments.length?(o=+t,n):o},velocityDecay:function(t){return arguments.length?(a=1-t,n):1-a},force:function(t,e){return arguments.length>1?(null==e?u.remove(t):u.set(t,d(e)),n):u.get(t)},find:function(n,e,r){var i,o,a,u,c,f=0,s=t.length;for(null==r?r=1/0:r*=r,f=0;f<s;++f)(a=(i=n-(u=t[f]).x)*i+(o=e-u.y)*o)<r&&(c=u,r=a);return c},on:function(t,e){return arguments.length>1?(f.on(t,e),n):f.on(t)}}},t.forceX=function(t){var n,e,r,i=ta(.1);function o(t){for(var i,o=0,a=n.length;o<a;++o)(i=n[o]).vx+=(r[o]-i.x)*e[o]*t}function a(){if(n){var o,a=n.length;for(e=new Array(a),r=new Array(a),o=0;o<a;++o)e[o]=isNaN(r[o]=+t(n[o],o,n))?0:+i(n[o],o,n)}}return"function"!=typeof t&&(t=ta(null==t?0:+t)),o.initialize=function(t){n=t,a()},o.strength=function(t){return arguments.length?(i="function"==typeof t?t:ta(+t),a(),o):i},o.x=function(n){return arguments.length?(t="function"==typeof n?n:ta(+n),a(),o):t},o},t.forceY=function(t){var n,e,r,i=ta(.1);function o(t){for(var i,o=0,a=n.length;o<a;++o)(i=n[o]).vy+=(r[o]-i.y)*e[o]*t}function a(){if(n){var o,a=n.length;for(e=new Array(a),r=new Array(a),o=0;o<a;++o)e[o]=isNaN(r[o]=+t(n[o],o,n))?0:+i(n[o],o,n)}}return"function"!=typeof t&&(t=ta(null==t?0:+t)),o.initialize=function(t){n=t,a()},o.strength=function(t){return arguments.length?(i="function"==typeof t?t:ta(+t),a(),o):i},o.y=function(n){return arguments.length?(t="function"==typeof n?n:ta(+n),a(),o):t},o},t.formatDefaultLocale=Ca,t.formatLocale=Ea,t.formatSpecifier=wa,t.precisionFixed=Pa,t.precisionPrefix=za,t.precisionRound=Ra,t.geoArea=function(t){return mu.reset(),du(t,xu),2*mu},t.geoBounds=function(t){var n,e,r,i,o,a,u;if(Lu=qu=-(Ru=Du=1/0),Fu=[],du(t,ac),e=Fu.length){for(Fu.sort(vc),n=1,o=[r=Fu[0]];n<e;++n)gc(r,(i=Fu[n])[0])||gc(r,i[1])?(pc(r[0],i[1])>pc(r[0],r[1])&&(r[1]=i[1]),pc(i[0],r[1])>pc(r[0],r[1])&&(r[0]=i[0])):o.push(r=i);for(a=-1/0,n=0,r=o[e=o.length-1];n<=e;r=i,++n)i=o[n],(u=pc(r[1],i[0]))>a&&(a=u,Ru=i[0],qu=r[1])}return Fu=Iu=null,Ru===1/0||Du===1/0?[[NaN,NaN],[NaN,NaN]]:[[Ru,Du],[qu,Lu]]},t.geoCentroid=function(t){ju=Hu=Xu=Gu=Vu=$u=Wu=Zu=Qu=Ju=Ku=0,du(t,yc);var n=Qu,e=Ju,r=Ku,i=n*n+e*e+r*r;return i<Ba&&(n=$u,e=Wu,r=Zu,Hu<Oa&&(n=Xu,e=Gu,r=Vu),(i=n*n+e*e+r*r)<Ba)?[NaN,NaN]:[$a(e,n)*Ha,ou(r/eu(i))*Ha]},t.geoCircle=function(){var t,n,e=kc([0,0]),r=kc(90),i=kc(6),o={point:function(e,r){t.push(e=n(e,r)),e[0]*=Ha,e[1]*=Ha}};function a(){var a=e.apply(this,arguments),u=r.apply(this,arguments)*Xa,c=i.apply(this,arguments)*Xa;return t=[],n=Pc(-a[0]*Xa,-a[1]*Xa,0).invert,Lc(o,u,c,1),a={type:"Polygon",coordinates:[t]},t=n=null,a}return a.center=function(t){return arguments.length?(e="function"==typeof t?t:kc([+t[0],+t[1]]),a):e},a.radius=function(t){return arguments.length?(r="function"==typeof t?t:kc(+t),a):r},a.precision=function(t){return arguments.length?(i="function"==typeof t?t:kc(+t),a):i},a},t.geoClipAntimeridian=$c,t.geoClipCircle=Wc,t.geoClipExtent=function(){var t,n,e,r=0,i=0,o=960,a=500;return e={stream:function(e){return t&&n===e?t:t=Jc(r,i,o,a)(n=e)},extent:function(u){return arguments.length?(r=+u[0][0],i=+u[0][1],o=+u[1][0],a=+u[1][1],t=n=null,e):[[r,i],[o,a]]}}},t.geoClipRectangle=Jc,t.geoContains=function(t,n){return(t&&hf.hasOwnProperty(t.type)?hf[t.type]:pf)(t,n)},t.geoDistance=lf,t.geoGraticule=wf,t.geoGraticule10=function(){return wf()()},t.geoInterpolate=function(t,n){var e=t[0]*Xa,r=t[1]*Xa,i=n[0]*Xa,o=n[1]*Xa,a=Wa(r),u=tu(r),c=Wa(o),f=tu(o),s=a*Wa(e),l=a*tu(e),h=c*Wa(i),d=c*tu(i),p=2*ou(eu(au(o-r)+a*c*au(i-e))),v=tu(p),g=p?function(t){var n=tu(t*=p)/v,e=tu(p-t)/v,r=e*s+n*h,i=e*l+n*d,o=e*u+n*f;return[$a(i,r)*Ha,$a(o,eu(r*r+i*i))*Ha]}:function(){return[e*Ha,r*Ha]};return g.distance=p,g},t.geoLength=cf,t.geoPath=function(t,n){var e,r,i=4.5;function o(t){return t&&("function"==typeof i&&r.pointRadius(+i.apply(this,arguments)),du(t,e(r))),r.result()}return o.area=function(t){return du(t,e(Cf)),Cf.result()},o.measure=function(t){return du(t,e(gs)),gs.result()},o.bounds=function(t){return du(t,e(Bf)),Bf.result()},o.centroid=function(t){return du(t,e(Kf)),Kf.result()},o.projection=function(n){return arguments.length?(e=null==n?(t=null,Mf):(t=n).stream,o):t},o.context=function(t){return arguments.length?(r=null==t?(n=null,new bs):new fs(n=t),"function"!=typeof i&&r.pointRadius(i),o):n},o.pointRadius=function(t){return arguments.length?(i="function"==typeof t?t:(r.pointRadius(+t),+t),o):i},o.projection(t).context(n)},t.geoAlbers=Os,t.geoAlbersUsa=function(){var t,n,e,r,i,o,a=Os(),u=Us().rotate([154,0]).center([-2,58.5]).parallels([55,65]),c=Us().rotate([157,0]).center([-3,19.9]).parallels([8,18]),f={point:function(t,n){o=[t,n]}};function s(t){var n=t[0],a=t[1];return o=null,e.point(n,a),o||(r.point(n,a),o)||(i.point(n,a),o)}function l(){return t=n=null,s}return s.invert=function(t){var n=a.scale(),e=a.translate(),r=(t[0]-e[0])/n,i=(t[1]-e[1])/n;return(i>=.12&&i<.234&&r>=-.425&&r<-.214?u:i>=.166&&i<.234&&r>=-.214&&r<-.115?c:a).invert(t)},s.stream=function(e){return t&&n===e?t:(r=[a.stream(n=e),u.stream(e),c.stream(e)],i=r.length,t={point:function(t,n){for(var e=-1;++e<i;)r[e].point(t,n)},sphere:function(){for(var t=-1;++t<i;)r[t].sphere()},lineStart:function(){for(var t=-1;++t<i;)r[t].lineStart()},lineEnd:function(){for(var t=-1;++t<i;)r[t].lineEnd()},polygonStart:function(){for(var t=-1;++t<i;)r[t].polygonStart()},polygonEnd:function(){for(var t=-1;++t<i;)r[t].polygonEnd()}});var r,i},s.precision=function(t){return arguments.length?(a.precision(t),u.precision(t),c.precision(t),l()):a.precision()},s.scale=function(t){return arguments.length?(a.scale(t),u.scale(.35*t),c.scale(t),s.translate(a.translate())):a.scale()},s.translate=function(t){if(!arguments.length)return a.translate();var n=a.scale(),o=+t[0],s=+t[1];return e=a.translate(t).clipExtent([[o-.455*n,s-.238*n],[o+.455*n,s+.238*n]]).stream(f),r=u.translate([o-.307*n,s+.201*n]).clipExtent([[o-.425*n+Oa,s+.12*n+Oa],[o-.214*n-Oa,s+.234*n-Oa]]).stream(f),i=c.translate([o-.205*n,s+.212*n]).clipExtent([[o-.214*n+Oa,s+.166*n+Oa],[o-.115*n-Oa,s+.234*n-Oa]]).stream(f),l()},s.fitExtent=function(t,n){return Ns(s,t,n)},s.fitSize=function(t,n){return As(s,t,n)},s.fitWidth=function(t,n){return Ts(s,t,n)},s.fitHeight=function(t,n){return Ss(s,t,n)},s.scale(1070)},t.geoAzimuthalEqualArea=function(){return Rs(Fs).scale(124.75).clipAngle(179.999)},t.geoAzimuthalEqualAreaRaw=Fs,t.geoAzimuthalEquidistant=function(){return Rs(Is).scale(79.4188).clipAngle(179.999)},t.geoAzimuthalEquidistantRaw=Is,t.geoConicConformal=function(){return qs(Gs).scale(109.5).parallels([30,30])},t.geoConicConformalRaw=Gs,t.geoConicEqualArea=Us,t.geoConicEqualAreaRaw=Ls,t.geoConicEquidistant=function(){return qs($s).scale(131.154).center([0,13.9389])},t.geoConicEquidistantRaw=$s,t.geoEqualEarth=function(){return Rs(tl).scale(177.158)},t.geoEqualEarthRaw=tl,t.geoEquirectangular=function(){return Rs(Vs).scale(152.63)},t.geoEquirectangularRaw=Vs,t.geoGnomonic=function(){return Rs(nl).scale(144.049).clipAngle(60)},t.geoGnomonicRaw=nl,t.geoIdentity=function(){var t,n,e,r,i,o,a=1,u=0,c=0,f=1,s=1,l=Mf,h=null,d=Mf;function p(){return r=i=null,o}return o={stream:function(t){return r&&i===t?r:r=l(d(i=t))},postclip:function(r){return arguments.length?(d=r,h=t=n=e=null,p()):d},clipExtent:function(r){return arguments.length?(d=null==r?(h=t=n=e=null,Mf):Jc(h=+r[0][0],t=+r[0][1],n=+r[1][0],e=+r[1][1]),p()):null==h?null:[[h,t],[n,e]]},scale:function(t){return arguments.length?(l=el((a=+t)*f,a*s,u,c),p()):a},translate:function(t){return arguments.length?(l=el(a*f,a*s,u=+t[0],c=+t[1]),p()):[u,c]},reflectX:function(t){return arguments.length?(l=el(a*(f=t?-1:1),a*s,u,c),p()):f<0},reflectY:function(t){return arguments.length?(l=el(a*f,a*(s=t?-1:1),u,c),p()):s<0},fitExtent:function(t,n){return Ns(o,t,n)},fitSize:function(t,n){return As(o,t,n)},fitWidth:function(t,n){return Ts(o,t,n)},fitHeight:function(t,n){return Ss(o,t,n)}}},t.geoProjection=Rs,t.geoProjectionMutator=Ds,t.geoMercator=function(){return Hs(js).scale(961/ja)},t.geoMercatorRaw=js,t.geoNaturalEarth1=function(){return Rs(rl).scale(175.295)},t.geoNaturalEarth1Raw=rl,t.geoOrthographic=function(){return Rs(il).scale(249.5).clipAngle(90+Oa)},t.geoOrthographicRaw=il,t.geoStereographic=function(){return Rs(ol).scale(250).clipAngle(142)},t.geoStereographicRaw=ol,t.geoTransverseMercator=function(){var t=Hs(al),n=t.center,e=t.rotate;return t.center=function(t){return arguments.length?n([-t[1],t[0]]):[(t=n())[1],-t[0]]},t.rotate=function(t){return arguments.length?e([t[0],t[1],t.length>2?t[2]+90:90]):[(t=e())[0],t[1],t[2]-90]},e([0,0,90]).scale(159.155)},t.geoTransverseMercatorRaw=al,t.geoRotation=qc,t.geoStream=du,t.geoTransform=function(t){return{stream:xs(t)}},t.cluster=function(){var t=ul,n=1,e=1,r=!1;function i(i){var o,a=0;i.eachAfter(function(n){var e=n.children;e?(n.x=function(t){return t.reduce(cl,0)/t.length}(e),n.y=function(t){return 1+t.reduce(fl,0)}(e)):(n.x=o?a+=t(n,o):0,n.y=0,o=n)});var u=function(t){for(var n;n=t.children;)t=n[0];return t}(i),c=function(t){for(var n;n=t.children;)t=n[n.length-1];return t}(i),f=u.x-t(u,c)/2,s=c.x+t(c,u)/2;return i.eachAfter(r?function(t){t.x=(t.x-i.x)*n,t.y=(i.y-t.y)*e}:function(t){t.x=(t.x-f)/(s-f)*n,t.y=(1-(i.y?t.y/i.y:1))*e})}return i.separation=function(n){return arguments.length?(t=n,i):t},i.size=function(t){return arguments.length?(r=!1,n=+t[0],e=+t[1],i):r?null:[n,e]},i.nodeSize=function(t){return arguments.length?(r=!0,n=+t[0],e=+t[1],i):r?[n,e]:null},i},t.hierarchy=ll,t.pack=function(){var t=null,n=1,e=1,r=Pl;function i(i){return i.x=n/2,i.y=e/2,t?i.eachBefore(Dl(t)).eachAfter(ql(r,.5)).eachBefore(Ll(1)):i.eachBefore(Dl(Rl)).eachAfter(ql(Pl,1)).eachAfter(ql(r,i.r/Math.min(n,e))).eachBefore(Ll(Math.min(n,e)/(2*i.r))),i}return i.radius=function(n){return arguments.length?(t=null==(e=n)?null:Cl(e),i):t;var e},i.size=function(t){return arguments.length?(n=+t[0],e=+t[1],i):[n,e]},i.padding=function(t){return arguments.length?(r="function"==typeof t?t:zl(+t),i):r},i},t.packSiblings=function(t){return El(t),t},t.packEnclose=yl,t.partition=function(){var t=1,n=1,e=0,r=!1;function i(i){var o=i.height+1;return i.x0=i.y0=e,i.x1=t,i.y1=n/o,i.eachBefore(function(t,n){return function(r){r.children&&Ol(r,r.x0,t*(r.depth+1)/n,r.x1,t*(r.depth+2)/n);var i=r.x0,o=r.y0,a=r.x1-e,u=r.y1-e;a<i&&(i=a=(i+a)/2),u<o&&(o=u=(o+u)/2),r.x0=i,r.y0=o,r.x1=a,r.y1=u}}(n,o)),r&&i.eachBefore(Ul),i}return i.round=function(t){return arguments.length?(r=!!t,i):r},i.size=function(e){return arguments.length?(t=+e[0],n=+e[1],i):[t,n]},i.padding=function(t){return arguments.length?(e=+t,i):e},i},t.stratify=function(){var t=Il,n=jl;function e(e){var r,i,o,a,u,c,f,s=e.length,l=new Array(s),h={};for(i=0;i<s;++i)r=e[i],u=l[i]=new vl(r),null!=(c=t(r,i,e))&&(c+="")&&(h[f=Bl+(u.id=c)]=f in h?Fl:u);for(i=0;i<s;++i)if(u=l[i],null!=(c=n(e[i],i,e))&&(c+="")){if(!(a=h[Bl+c]))throw new Error("missing: "+c);if(a===Fl)throw new Error("ambiguous: "+c);a.children?a.children.push(u):a.children=[u],u.parent=a}else{if(o)throw new Error("multiple roots");o=u}if(!o)throw new Error("no root");if(o.parent=Yl,o.eachBefore(function(t){t.depth=t.parent.depth+1,--s}).eachBefore(pl),o.parent=null,s>0)throw new Error("cycle");return o}return e.id=function(n){return arguments.length?(t=Cl(n),e):t},e.parentId=function(t){return arguments.length?(n=Cl(t),e):n},e},t.tree=function(){var t=Hl,n=1,e=1,r=null;function i(i){var c=function(t){for(var n,e,r,i,o,a=new Wl(t,0),u=[a];n=u.pop();)if(r=n._.children)for(n.children=new Array(o=r.length),i=o-1;i>=0;--i)u.push(e=n.children[i]=new Wl(r[i],i)),e.parent=n;return(a.parent=new Wl(null,0)).children=[a],a}(i);if(c.eachAfter(o),c.parent.m=-c.z,c.eachBefore(a),r)i.eachBefore(u);else{var f=i,s=i,l=i;i.eachBefore(function(t){t.x<f.x&&(f=t),t.x>s.x&&(s=t),t.depth>l.depth&&(l=t)});var h=f===s?1:t(f,s)/2,d=h-f.x,p=n/(s.x+h+d),v=e/(l.depth||1);i.eachBefore(function(t){t.x=(t.x+d)*p,t.y=t.depth*v})}return i}function o(n){var e=n.children,r=n.parent.children,i=n.i?r[n.i-1]:null;if(e){!function(t){for(var n,e=0,r=0,i=t.children,o=i.length;--o>=0;)(n=i[o]).z+=e,n.m+=e,e+=n.s+(r+=n.c)}(n);var o=(e[0].z+e[e.length-1].z)/2;i?(n.z=i.z+t(n._,i._),n.m=n.z-o):n.z=o}else i&&(n.z=i.z+t(n._,i._));n.parent.A=function(n,e,r){if(e){for(var i,o=n,a=n,u=e,c=o.parent.children[0],f=o.m,s=a.m,l=u.m,h=c.m;u=Gl(u),o=Xl(o),u&&o;)c=Xl(c),(a=Gl(a)).a=n,(i=u.z+l-o.z-f+t(u._,o._))>0&&(Vl($l(u,n,r),n,i),f+=i,s+=i),l+=u.m,f+=o.m,h+=c.m,s+=a.m;u&&!Gl(a)&&(a.t=u,a.m+=l-s),o&&!Xl(c)&&(c.t=o,c.m+=f-h,r=n)}return r}(n,i,n.parent.A||r[0])}function a(t){t._.x=t.z+t.parent.m,t.m+=t.parent.m}function u(t){t.x*=n,t.y=t.depth*e}return i.separation=function(n){return arguments.length?(t=n,i):t},i.size=function(t){return arguments.length?(r=!1,n=+t[0],e=+t[1],i):r?null:[n,e]},i.nodeSize=function(t){return arguments.length?(r=!0,n=+t[0],e=+t[1],i):r?[n,e]:null},i},t.treemap=function(){var t=Kl,n=!1,e=1,r=1,i=[0],o=Pl,a=Pl,u=Pl,c=Pl,f=Pl;function s(t){return t.x0=t.y0=0,t.x1=e,t.y1=r,t.eachBefore(l),i=[0],n&&t.eachBefore(Ul),t}function l(n){var e=i[n.depth],r=n.x0+e,s=n.y0+e,l=n.x1-e,h=n.y1-e;l<r&&(r=l=(r+l)/2),h<s&&(s=h=(s+h)/2),n.x0=r,n.y0=s,n.x1=l,n.y1=h,n.children&&(e=i[n.depth+1]=o(n)/2,r+=f(n)-e,s+=a(n)-e,(l-=u(n)-e)<r&&(r=l=(r+l)/2),(h-=c(n)-e)<s&&(s=h=(s+h)/2),t(n,r,s,l,h))}return s.round=function(t){return arguments.length?(n=!!t,s):n},s.size=function(t){return arguments.length?(e=+t[0],r=+t[1],s):[e,r]},s.tile=function(n){return arguments.length?(t=Cl(n),s):t},s.padding=function(t){return arguments.length?s.paddingInner(t).paddingOuter(t):s.paddingInner()},s.paddingInner=function(t){return arguments.length?(o="function"==typeof t?t:zl(+t),s):o},s.paddingOuter=function(t){return arguments.length?s.paddingTop(t).paddingRight(t).paddingBottom(t).paddingLeft(t):s.paddingTop()},s.paddingTop=function(t){return arguments.length?(a="function"==typeof t?t:zl(+t),s):a},s.paddingRight=function(t){return arguments.length?(u="function"==typeof t?t:zl(+t),s):u},s.paddingBottom=function(t){return arguments.length?(c="function"==typeof t?t:zl(+t),s):c},s.paddingLeft=function(t){return arguments.length?(f="function"==typeof t?t:zl(+t),s):f},s},t.treemapBinary=function(t,n,e,r,i){var o,a,u=t.children,c=u.length,f=new Array(c+1);for(f[0]=a=o=0;o<c;++o)f[o+1]=a+=u[o].value;!function t(n,e,r,i,o,a,c){if(n>=e-1){var s=u[n];return s.x0=i,s.y0=o,s.x1=a,void(s.y1=c)}for(var l=f[n],h=r/2+l,d=n+1,p=e-1;d<p;){var v=d+p>>>1;f[v]<h?d=v+1:p=v}h-f[d-1]<f[d]-h&&n+1<d&&--d;var g=f[d]-l,y=r-g;if(a-i>c-o){var _=(i*y+a*g)/r;t(n,d,g,i,o,_,c),t(d,e,y,_,o,a,c)}else{var b=(o*y+c*g)/r;t(n,d,g,i,o,a,b),t(d,e,y,i,b,a,c)}}(0,c,t.value,n,e,r,i)},t.treemapDice=Ol,t.treemapSlice=Zl,t.treemapSliceDice=function(t,n,e,r,i){(1&t.depth?Zl:Ol)(t,n,e,r,i)},t.treemapSquarify=Kl,t.treemapResquarify=th,t.interpolate=ye,t.interpolateArray=se,t.interpolateBasis=Kn,t.interpolateBasisClosed=te,t.interpolateDate=le,t.interpolateDiscrete=function(t){var n=t.length;return function(e){return t[Math.max(0,Math.min(n-1,Math.floor(e*n)))]}},t.interpolateHue=function(t,n){var e=re(+t,+n);return function(t){var n=e(t);return n-360*Math.floor(n/360)}},t.interpolateNumber=he,t.interpolateObject=de,t.interpolateRound=_e,t.interpolateString=ge,t.interpolateTransformCss=Se,t.interpolateTransformSvg=ke,t.interpolateZoom=De,t.interpolateRgb=ae,t.interpolateRgbBasis=ce,t.interpolateRgbBasisClosed=fe,t.interpolateHsl=Le,t.interpolateHslLong=Ue,t.interpolateLab=function(t,n){var e=oe((t=Rn(t)).l,(n=Rn(n)).l),r=oe(t.a,n.a),i=oe(t.b,n.b),o=oe(t.opacity,n.opacity);return function(n){return t.l=e(n),t.a=r(n),t.b=i(n),t.opacity=o(n),t+""}},t.interpolateHcl=Be,t.interpolateHclLong=Ye,t.interpolateCubehelix=Ie,t.interpolateCubehelixLong=je,t.piecewise=function(t,n){for(var e=0,r=n.length-1,i=n[0],o=new Array(r<0?0:r);e<r;)o[e]=t(i,i=n[++e]);return function(t){var n=Math.max(0,Math.min(r-1,Math.floor(t*=r)));return o[n](t-n)}},t.quantize=function(t,n){for(var e=new Array(n),r=0;r<n;++r)e[r]=t(r/(n-1));return e},t.path=Hi,t.polygonArea=function(t){for(var n,e=-1,r=t.length,i=t[r-1],o=0;++e<r;)n=i,i=t[e],o+=n[1]*i[0]-n[0]*i[1];return o/2},t.polygonCentroid=function(t){for(var n,e,r=-1,i=t.length,o=0,a=0,u=t[i-1],c=0;++r<i;)n=u,u=t[r],c+=e=n[0]*u[1]-u[0]*n[1],o+=(n[0]+u[0])*e,a+=(n[1]+u[1])*e;return[o/(c*=3),a/c]},t.polygonHull=function(t){if((e=t.length)<3)return null;var n,e,r=new Array(e),i=new Array(e);for(n=0;n<e;++n)r[n]=[+t[n][0],+t[n][1],n];for(r.sort(nh),n=0;n<e;++n)i[n]=[r[n][0],-r[n][1]];var o=eh(r),a=eh(i),u=a[0]===o[0],c=a[a.length-1]===o[o.length-1],f=[];for(n=o.length-1;n>=0;--n)f.push(t[r[o[n]][2]]);for(n=+u;n<a.length-c;++n)f.push(t[r[a[n]][2]]);return f},t.polygonContains=function(t,n){for(var e,r,i=t.length,o=t[i-1],a=n[0],u=n[1],c=o[0],f=o[1],s=!1,l=0;l<i;++l)e=(o=t[l])[0],(r=o[1])>u!=f>u&&a<(c-e)*(u-r)/(f-r)+e&&(s=!s),c=e,f=r;return s},t.polygonLength=function(t){for(var n,e,r=-1,i=t.length,o=t[i-1],a=o[0],u=o[1],c=0;++r<i;)n=a,e=u,n-=a=(o=t[r])[0],e-=u=o[1],c+=Math.sqrt(n*n+e*e);return c},t.quadtree=aa,t.randomUniform=ih,t.randomNormal=oh,t.randomLogNormal=ah,t.randomBates=ch,t.randomIrwinHall=uh,t.randomExponential=fh,t.scaleBand=yh,t.scalePoint=function(){return function t(n){var e=n.copy;return n.padding=n.paddingOuter,delete n.paddingInner,delete n.paddingOuter,n.copy=function(){return t(e())},n}(yh.apply(null,arguments).paddingInner(1))},t.scaleIdentity=function t(n){var e;function r(t){return isNaN(t=+t)?e:t}return r.invert=r,r.domain=r.range=function(t){return arguments.length?(n=dh.call(t,_h),r):n.slice()},r.unknown=function(t){return arguments.length?(e=t,r):e},r.copy=function(){return t(n).unknown(e)},n=arguments.length?dh.call(n,_h):[0,1],Eh(r)},t.scaleLinear=function t(){var n=Sh(mh,mh);return n.copy=function(){return Ah(n,t())},sh.apply(n,arguments),Eh(n)},t.scaleLog=function t(){var n=Uh(Th()).domain([1,10]);return n.copy=function(){return Ah(n,t()).base(n.base())},sh.apply(n,arguments),n},t.scaleSymlog=function t(){var n=Yh(Th());return n.copy=function(){return Ah(n,t()).constant(n.constant())},sh.apply(n,arguments)},t.scaleOrdinal=gh,t.scaleImplicit=vh,t.scalePow=Xh,t.scaleSqrt=function(){return Xh.apply(null,arguments).exponent(.5)},t.scaleQuantile=function t(){var e,r=[],o=[],a=[];function u(){var t=0,n=Math.max(1,o.length);for(a=new Array(n-1);++t<n;)a[t-1]=N(r,t/n);return c}function c(t){return isNaN(t=+t)?e:o[i(a,t)]}return c.invertExtent=function(t){var n=o.indexOf(t);return n<0?[NaN,NaN]:[n>0?a[n-1]:r[0],n<a.length?a[n]:r[r.length-1]]},c.domain=function(t){if(!arguments.length)return r.slice();r=[];for(var e,i=0,o=t.length;i<o;++i)null==(e=t[i])||isNaN(e=+e)||r.push(e);return r.sort(n),u()},c.range=function(t){return arguments.length?(o=ph.call(t),u()):o.slice()},c.unknown=function(t){return arguments.length?(e=t,c):e},c.quantiles=function(){return a.slice()},c.copy=function(){return t().domain(r).range(o).unknown(e)},sh.apply(c,arguments)},t.scaleQuantize=function t(){var n,e=0,r=1,o=1,a=[.5],u=[0,1];function c(t){return t<=t?u[i(a,t,0,o)]:n}function f(){var t=-1;for(a=new Array(o);++t<o;)a[t]=((t+1)*r-(t-o)*e)/(o+1);return c}return c.domain=function(t){return arguments.length?(e=+t[0],r=+t[1],f()):[e,r]},c.range=function(t){return arguments.length?(o=(u=ph.call(t)).length-1,f()):u.slice()},c.invertExtent=function(t){var n=u.indexOf(t);return n<0?[NaN,NaN]:n<1?[e,a[0]]:n>=o?[a[o-1],r]:[a[n-1],a[n]]},c.unknown=function(t){return arguments.length?(n=t,c):c},c.thresholds=function(){return a.slice()},c.copy=function(){return t().domain([e,r]).range(u).unknown(n)},sh.apply(Eh(c),arguments)},t.scaleThreshold=function t(){var n,e=[.5],r=[0,1],o=1;function a(t){return t<=t?r[i(e,t,0,o)]:n}return a.domain=function(t){return arguments.length?(e=ph.call(t),o=Math.min(e.length,r.length-1),a):e.slice()},a.range=function(t){return arguments.length?(r=ph.call(t),o=Math.min(e.length,r.length-1),a):r.slice()},a.invertExtent=function(t){var n=r.indexOf(t);return[e[n-1],e[n]]},a.unknown=function(t){return arguments.length?(n=t,a):n},a.copy=function(){return t().domain(e).range(r).unknown(n)},sh.apply(a,arguments)},t.scaleTime=function(){return sh.apply(Nv(Nd,wd,cd,od,rd,nd,Kh,Wh,t.timeFormat).domain([new Date(2e3,0,1),new Date(2e3,0,2)]),arguments)},t.scaleUtc=function(){return sh.apply(Nv(Wd,Vd,Rd,Cd,kd,Td,Kh,Wh,t.utcFormat).domain([Date.UTC(2e3,0,1),Date.UTC(2e3,0,2)]),arguments)},t.scaleSequential=function t(){var n=Eh(Av()(mh));return n.copy=function(){return Tv(n,t())},lh.apply(n,arguments)},t.scaleSequentialLog=function t(){var n=Uh(Av()).domain([1,10]);return n.copy=function(){return Tv(n,t()).base(n.base())},lh.apply(n,arguments)},t.scaleSequentialPow=Sv,t.scaleSequentialSqrt=function(){return Sv.apply(null,arguments).exponent(.5)},t.scaleSequentialSymlog=function t(){var n=Yh(Av());return n.copy=function(){return Tv(n,t()).constant(n.constant())},lh.apply(n,arguments)},t.scaleSequentialQuantile=function t(){var e=[],r=mh;function o(t){if(!isNaN(t=+t))return r((i(e,t)-1)/(e.length-1))}return o.domain=function(t){if(!arguments.length)return e.slice();e=[];for(var r,i=0,a=t.length;i<a;++i)null==(r=t[i])||isNaN(r=+r)||e.push(r);return e.sort(n),o},o.interpolator=function(t){return arguments.length?(r=t,o):r},o.copy=function(){return t(r).domain(e)},lh.apply(o,arguments)},t.scaleDiverging=function t(){var n=Eh(kv()(mh));return n.copy=function(){return Tv(n,t())},lh.apply(n,arguments)},t.scaleDivergingLog=function t(){var n=Uh(kv()).domain([.1,1,10]);return n.copy=function(){return Tv(n,t()).base(n.base())},lh.apply(n,arguments)},t.scaleDivergingPow=Ev,t.scaleDivergingSqrt=function(){return Ev.apply(null,arguments).exponent(.5)},t.scaleDivergingSymlog=function t(){var n=Yh(kv());return n.copy=function(){return Tv(n,t()).constant(n.constant())},lh.apply(n,arguments)},t.tickFormat=kh,t.schemeCategory10=Pv,t.schemeAccent=zv,t.schemeDark2=Rv,t.schemePaired=Dv,t.schemePastel1=qv,t.schemePastel2=Lv,t.schemeSet1=Uv,t.schemeSet2=Ov,t.schemeSet3=Bv,t.interpolateBrBG=Iv,t.schemeBrBG=Fv,t.interpolatePRGn=Hv,t.schemePRGn=jv,t.interpolatePiYG=Gv,t.schemePiYG=Xv,t.interpolatePuOr=$v,t.schemePuOr=Vv,t.interpolateRdBu=Zv,t.schemeRdBu=Wv,t.interpolateRdGy=Jv,t.schemeRdGy=Qv,t.interpolateRdYlBu=tg,t.schemeRdYlBu=Kv,t.interpolateRdYlGn=eg,t.schemeRdYlGn=ng,t.interpolateSpectral=ig,t.schemeSpectral=rg,t.interpolateBuGn=ag,t.schemeBuGn=og,t.interpolateBuPu=cg,t.schemeBuPu=ug,t.interpolateGnBu=sg,t.schemeGnBu=fg,t.interpolateOrRd=hg,t.schemeOrRd=lg,t.interpolatePuBuGn=pg,t.schemePuBuGn=dg,t.interpolatePuBu=gg,t.schemePuBu=vg,t.interpolatePuRd=_g,t.schemePuRd=yg,t.interpolateRdPu=mg,t.schemeRdPu=bg,t.interpolateYlGnBu=wg,t.schemeYlGnBu=xg,t.interpolateYlGn=Ng,t.schemeYlGn=Mg,t.interpolateYlOrBr=Tg,t.schemeYlOrBr=Ag,t.interpolateYlOrRd=kg,t.schemeYlOrRd=Sg,t.interpolateBlues=Cg,t.schemeBlues=Eg,t.interpolateGreens=zg,t.schemeGreens=Pg,t.interpolateGreys=Dg,t.schemeGreys=Rg,t.interpolatePurples=Lg,t.schemePurples=qg,t.interpolateReds=Og,t.schemeReds=Ug,t.interpolateOranges=Yg,t.schemeOranges=Bg,t.interpolateCubehelixDefault=Fg,t.interpolateRainbow=function(t){(t<0||t>1)&&(t-=Math.floor(t));var n=Math.abs(t-.5);return Hg.h=360*t-100,Hg.s=1.5-1.5*n,Hg.l=.8-.9*n,Hg+""},t.interpolateWarm=Ig,t.interpolateCool=jg,t.interpolateSinebow=function(t){var n;return t=(.5-t)*Math.PI,Xg.r=255*(n=Math.sin(t))*n,Xg.g=255*(n=Math.sin(t+Gg))*n,Xg.b=255*(n=Math.sin(t+Vg))*n,Xg+""},t.interpolateViridis=Wg,t.interpolateMagma=Zg,t.interpolateInferno=Qg,t.interpolatePlasma=Jg,t.create=function(t){return zt(W(t).call(document.documentElement))},t.creator=W,t.local=Dt,t.matcher=tt,t.mouse=Ot,t.namespace=$,t.namespaces=V,t.clientPoint=Ut,t.select=zt,t.selectAll=function(t){return"string"==typeof t?new Ct([document.querySelectorAll(t)],[document.documentElement]):new Ct([null==t?[]:t],Et)},t.selection=Pt,t.selector=Q,t.selectorAll=K,t.style=ct,t.touch=Bt,t.touches=function(t,n){null==n&&(n=Lt().touches);for(var e=0,r=n?n.length:0,i=new Array(r);e<r;++e)i[e]=Ut(t,n[e]);return i},t.window=ut,t.customEvent=St,t.arc=function(){var t=hy,n=dy,e=Kg(0),r=null,i=py,o=vy,a=gy,u=null;function c(){var c,f,s,l=+t.apply(this,arguments),h=+n.apply(this,arguments),d=i.apply(this,arguments)-fy,p=o.apply(this,arguments)-fy,v=ty(p-d),g=p>d;if(u||(u=c=Hi()),h<l&&(f=h,h=l,l=f),h>uy)if(v>sy-uy)u.moveTo(h*ey(d),h*oy(d)),u.arc(0,0,h,d,p,!g),l>uy&&(u.moveTo(l*ey(p),l*oy(p)),u.arc(0,0,l,p,d,g));else{var y,_,b=d,m=p,x=d,w=p,M=v,N=v,A=a.apply(this,arguments)/2,T=A>uy&&(r?+r.apply(this,arguments):ay(l*l+h*h)),S=iy(ty(h-l)/2,+e.apply(this,arguments)),k=S,E=S;if(T>uy){var C=ly(T/l*oy(A)),P=ly(T/h*oy(A));(M-=2*C)>uy?(x+=C*=g?1:-1,w-=C):(M=0,x=w=(d+p)/2),(N-=2*P)>uy?(b+=P*=g?1:-1,m-=P):(N=0,b=m=(d+p)/2)}var z=h*ey(b),R=h*oy(b),D=l*ey(w),q=l*oy(w);if(S>uy){var L,U=h*ey(m),O=h*oy(m),B=l*ey(x),Y=l*oy(x);if(v<cy&&(L=function(t,n,e,r,i,o,a,u){var c=e-t,f=r-n,s=a-i,l=u-o,h=l*c-s*f;if(!(h*h<uy))return[t+(h=(s*(n-o)-l*(t-i))/h)*c,n+h*f]}(z,R,B,Y,U,O,D,q))){var F=z-L[0],I=R-L[1],j=U-L[0],H=O-L[1],X=1/oy(((s=(F*j+I*H)/(ay(F*F+I*I)*ay(j*j+H*H)))>1?0:s<-1?cy:Math.acos(s))/2),G=ay(L[0]*L[0]+L[1]*L[1]);k=iy(S,(l-G)/(X-1)),E=iy(S,(h-G)/(X+1))}}N>uy?E>uy?(y=yy(B,Y,z,R,h,E,g),_=yy(U,O,D,q,h,E,g),u.moveTo(y.cx+y.x01,y.cy+y.y01),E<S?u.arc(y.cx,y.cy,E,ny(y.y01,y.x01),ny(_.y01,_.x01),!g):(u.arc(y.cx,y.cy,E,ny(y.y01,y.x01),ny(y.y11,y.x11),!g),u.arc(0,0,h,ny(y.cy+y.y11,y.cx+y.x11),ny(_.cy+_.y11,_.cx+_.x11),!g),u.arc(_.cx,_.cy,E,ny(_.y11,_.x11),ny(_.y01,_.x01),!g))):(u.moveTo(z,R),u.arc(0,0,h,b,m,!g)):u.moveTo(z,R),l>uy&&M>uy?k>uy?(y=yy(D,q,U,O,l,-k,g),_=yy(z,R,B,Y,l,-k,g),u.lineTo(y.cx+y.x01,y.cy+y.y01),k<S?u.arc(y.cx,y.cy,k,ny(y.y01,y.x01),ny(_.y01,_.x01),!g):(u.arc(y.cx,y.cy,k,ny(y.y01,y.x01),ny(y.y11,y.x11),!g),u.arc(0,0,l,ny(y.cy+y.y11,y.cx+y.x11),ny(_.cy+_.y11,_.cx+_.x11),g),u.arc(_.cx,_.cy,k,ny(_.y11,_.x11),ny(_.y01,_.x01),!g))):u.arc(0,0,l,w,x,g):u.lineTo(D,q)}else u.moveTo(0,0);if(u.closePath(),c)return u=null,c+""||null}return c.centroid=function(){var e=(+t.apply(this,arguments)+ +n.apply(this,arguments))/2,r=(+i.apply(this,arguments)+ +o.apply(this,arguments))/2-cy/2;return[ey(r)*e,oy(r)*e]},c.innerRadius=function(n){return arguments.length?(t="function"==typeof n?n:Kg(+n),c):t},c.outerRadius=function(t){return arguments.length?(n="function"==typeof t?t:Kg(+t),c):n},c.cornerRadius=function(t){return arguments.length?(e="function"==typeof t?t:Kg(+t),c):e},c.padRadius=function(t){return arguments.length?(r=null==t?null:"function"==typeof t?t:Kg(+t),c):r},c.startAngle=function(t){return arguments.length?(i="function"==typeof t?t:Kg(+t),c):i},c.endAngle=function(t){return arguments.length?(o="function"==typeof t?t:Kg(+t),c):o},c.padAngle=function(t){return arguments.length?(a="function"==typeof t?t:Kg(+t),c):a},c.context=function(t){return arguments.length?(u=null==t?null:t,c):u},c},t.area=My,t.line=wy,t.pie=function(){var t=Ay,n=Ny,e=null,r=Kg(0),i=Kg(sy),o=Kg(0);function a(a){var u,c,f,s,l,h=a.length,d=0,p=new Array(h),v=new Array(h),g=+r.apply(this,arguments),y=Math.min(sy,Math.max(-sy,i.apply(this,arguments)-g)),_=Math.min(Math.abs(y)/h,o.apply(this,arguments)),b=_*(y<0?-1:1);for(u=0;u<h;++u)(l=v[p[u]=u]=+t(a[u],u,a))>0&&(d+=l);for(null!=n?p.sort(function(t,e){return n(v[t],v[e])}):null!=e&&p.sort(function(t,n){return e(a[t],a[n])}),u=0,f=d?(y-h*b)/d:0;u<h;++u,g=s)c=p[u],s=g+((l=v[c])>0?l*f:0)+b,v[c]={data:a[c],index:u,value:l,startAngle:g,endAngle:s,padAngle:_};return v}return a.value=function(n){return arguments.length?(t="function"==typeof n?n:Kg(+n),a):t},a.sortValues=function(t){return arguments.length?(n=t,e=null,a):n},a.sort=function(t){return arguments.length?(e=t,n=null,a):e},a.startAngle=function(t){return arguments.length?(r="function"==typeof t?t:Kg(+t),a):r},a.endAngle=function(t){return arguments.length?(i="function"==typeof t?t:Kg(+t),a):i},a.padAngle=function(t){return arguments.length?(o="function"==typeof t?t:Kg(+t),a):o},a},t.areaRadial=Py,t.radialArea=Py,t.lineRadial=Cy,t.radialLine=Cy,t.pointRadial=zy,t.linkHorizontal=function(){return Ly(Uy)},t.linkVertical=function(){return Ly(Oy)},t.linkRadial=function(){var t=Ly(By);return t.angle=t.x,delete t.x,t.radius=t.y,delete t.y,t},t.symbol=function(){var t=Kg(Yy),n=Kg(64),e=null;function r(){var r;if(e||(e=r=Hi()),t.apply(this,arguments).draw(e,+n.apply(this,arguments)),r)return e=null,r+""||null}return r.type=function(n){return arguments.length?(t="function"==typeof n?n:Kg(n),r):t},r.size=function(t){return arguments.length?(n="function"==typeof t?t:Kg(+t),r):n},r.context=function(t){return arguments.length?(e=null==t?null:t,r):e},r},t.symbols=e_,t.symbolCircle=Yy,t.symbolCross=Fy,t.symbolDiamond=Hy,t.symbolSquare=Wy,t.symbolStar=$y,t.symbolTriangle=Qy,t.symbolWye=n_,t.curveBasisClosed=function(t){return new a_(t)},t.curveBasisOpen=function(t){return new u_(t)},t.curveBasis=function(t){return new o_(t)},t.curveBundle=f_,t.curveCardinalClosed=p_,t.curveCardinalOpen=g_,t.curveCardinal=h_,t.curveCatmullRomClosed=x_,t.curveCatmullRomOpen=M_,t.curveCatmullRom=b_,t.curveLinearClosed=function(t){return new N_(t)},t.curveLinear=by,t.curveMonotoneX=function(t){return new E_(t)},t.curveMonotoneY=function(t){return new C_(t)},t.curveNatural=function(t){return new z_(t)},t.curveStep=function(t){return new D_(t,.5)},t.curveStepAfter=function(t){return new D_(t,1)},t.curveStepBefore=function(t){return new D_(t,0)},t.stack=function(){var t=Kg([]),n=L_,e=q_,r=U_;function i(i){var o,a,u=t.apply(this,arguments),c=i.length,f=u.length,s=new Array(f);for(o=0;o<f;++o){for(var l,h=u[o],d=s[o]=new Array(c),p=0;p<c;++p)d[p]=l=[0,+r(i[p],h,p,i)],l.data=i[p];d.key=h}for(o=0,a=n(s);o<f;++o)s[a[o]].index=o;return e(s,a),s}return i.keys=function(n){return arguments.length?(t="function"==typeof n?n:Kg(Ry.call(n)),i):t},i.value=function(t){return arguments.length?(r="function"==typeof t?t:Kg(+t),i):r},i.order=function(t){return arguments.length?(n=null==t?L_:"function"==typeof t?t:Kg(Ry.call(t)),i):n},i.offset=function(t){return arguments.length?(e=null==t?q_:t,i):e},i},t.stackOffsetExpand=function(t,n){if((r=t.length)>0){for(var e,r,i,o=0,a=t[0].length;o<a;++o){for(i=e=0;e<r;++e)i+=t[e][o][1]||0;if(i)for(e=0;e<r;++e)t[e][o][1]/=i}q_(t,n)}},t.stackOffsetDiverging=function(t,n){if((u=t.length)>1)for(var e,r,i,o,a,u,c=0,f=t[n[0]].length;c<f;++c)for(o=a=0,e=0;e<u;++e)(i=(r=t[n[e]][c])[1]-r[0])>=0?(r[0]=o,r[1]=o+=i):i<0?(r[1]=a,r[0]=a+=i):r[0]=o},t.stackOffsetNone=q_,t.stackOffsetSilhouette=function(t,n){if((e=t.length)>0){for(var e,r=0,i=t[n[0]],o=i.length;r<o;++r){for(var a=0,u=0;a<e;++a)u+=t[a][r][1]||0;i[r][1]+=i[r][0]=-u/2}q_(t,n)}},t.stackOffsetWiggle=function(t,n){if((i=t.length)>0&&(r=(e=t[n[0]]).length)>0){for(var e,r,i,o=0,a=1;a<r;++a){for(var u=0,c=0,f=0;u<i;++u){for(var s=t[n[u]],l=s[a][1]||0,h=(l-(s[a-1][1]||0))/2,d=0;d<u;++d){var p=t[n[d]];h+=(p[a][1]||0)-(p[a-1][1]||0)}c+=l,f+=h*l}e[a-1][1]+=e[a-1][0]=o,c&&(o-=f/c)}e[a-1][1]+=e[a-1][0]=o,q_(t,n)}},t.stackOrderAppearance=O_,t.stackOrderAscending=Y_,t.stackOrderDescending=function(t){return Y_(t).reverse()},t.stackOrderInsideOut=function(t){var n,e,r=t.length,i=t.map(F_),o=O_(t),a=0,u=0,c=[],f=[];for(n=0;n<r;++n)e=o[n],a<u?(a+=i[e],c.push(e)):(u+=i[e],f.push(e));return f.reverse().concat(c)},t.stackOrderNone=L_,t.stackOrderReverse=function(t){return L_(t).reverse()},t.timeInterval=$h,t.timeMillisecond=Wh,t.timeMilliseconds=Zh,t.utcMillisecond=Wh,t.utcMilliseconds=Zh,t.timeSecond=Kh,t.timeSeconds=td,t.utcSecond=Kh,t.utcSeconds=td,t.timeMinute=nd,t.timeMinutes=ed,t.timeHour=rd,t.timeHours=id,t.timeDay=od,t.timeDays=ad,t.timeWeek=cd,t.timeWeeks=vd,t.timeSunday=cd,t.timeSundays=vd,t.timeMonday=fd,t.timeMondays=gd,t.timeTuesday=sd,t.timeTuesdays=yd,t.timeWednesday=ld,t.timeWednesdays=_d,t.timeThursday=hd,t.timeThursdays=bd,t.timeFriday=dd,t.timeFridays=md,t.timeSaturday=pd,t.timeSaturdays=xd,t.timeMonth=wd,t.timeMonths=Md,t.timeYear=Nd,t.timeYears=Ad,t.utcMinute=Td,t.utcMinutes=Sd,t.utcHour=kd,t.utcHours=Ed,t.utcDay=Cd,t.utcDays=Pd,t.utcWeek=Rd,t.utcWeeks=Yd,t.utcSunday=Rd,t.utcSundays=Yd,t.utcMonday=Dd,t.utcMondays=Fd,t.utcTuesday=qd,t.utcTuesdays=Id,t.utcWednesday=Ld,t.utcWednesdays=jd,t.utcThursday=Ud,t.utcThursdays=Hd,t.utcFriday=Od,t.utcFridays=Xd,t.utcSaturday=Bd,t.utcSaturdays=Gd,t.utcMonth=Vd,t.utcMonths=$d,t.utcYear=Wd,t.utcYears=Zd,t.timeFormatDefaultLocale=hv,t.timeFormatLocale=tp,t.isoFormat=dv,t.isoParse=pv,t.now=nr,t.timer=ir,t.timerFlush=or,t.timeout=fr,t.interval=function(t,n,e){var r=new rr,i=n;return null==n?(r.restart(t,n,e),r):(n=+n,e=null==e?nr():+e,r.restart(function o(a){a+=i,r.restart(o,i+=n,e),t(a)},n,e),r)},t.transition=Cr,t.active=function(t,n){var e,r,i=t.__transition;if(i)for(r in n=null==n?null:n+"",i)if((e=i[r]).state>dr&&e.name===n)return new Er([[t]],fi,n,+r);return null},t.interrupt=Mr,t.voronoi=function(){var t=j_,n=H_,e=null;function r(r){return new Nb(r.map(function(e,i){var o=[Math.round(t(e,i,r)/xb)*xb,Math.round(n(e,i,r)/xb)*xb];return o.index=i,o.data=e,o}),e)}return r.polygons=function(t){return r(t).polygons()},r.links=function(t){return r(t).links()},r.triangles=function(t){return r(t).triangles()},r.x=function(n){return arguments.length?(t="function"==typeof n?n:I_(+n),r):t},r.y=function(t){return arguments.length?(n="function"==typeof t?t:I_(+t),r):n},r.extent=function(t){return arguments.length?(e=null==t?null:[[+t[0][0],+t[0][1]],[+t[1][0],+t[1][1]]],r):e&&[[e[0][0],e[0][1]],[e[1][0],e[1][1]]]},r.size=function(t){return arguments.length?(e=null==t?null:[[0,0],[+t[0],+t[1]]],r):e&&[e[1][0]-e[0][0],e[1][1]-e[0][1]]},r},t.zoom=function(){var n,e,r=zb,i=Rb,o=Ub,a=qb,u=Lb,c=[0,1/0],f=[[-1/0,-1/0],[1/0,1/0]],s=250,l=De,h=[],d=I("start","zoom","end"),p=500,v=150,g=0;function y(t){t.property("__zoom",Db).on("wheel.zoom",N).on("mousedown.zoom",A).on("dblclick.zoom",T).filter(u).on("touchstart.zoom",S).on("touchmove.zoom",k).on("touchend.zoom touchcancel.zoom",E).style("touch-action","none").style("-webkit-tap-highlight-color","rgba(0,0,0,0)")}function _(t,n){return(n=Math.max(c[0],Math.min(c[1],n)))===t.k?t:new Sb(n,t.x,t.y)}function b(t,n,e){var r=n[0]-e[0]*t.k,i=n[1]-e[1]*t.k;return r===t.x&&i===t.y?t:new Sb(t.k,r,i)}function m(t){return[(+t[0][0]+ +t[1][0])/2,(+t[0][1]+ +t[1][1])/2]}function x(t,n,e){t.on("start.zoom",function(){w(this,arguments).start()}).on("interrupt.zoom end.zoom",function(){w(this,arguments).end()}).tween("zoom",function(){var t=arguments,r=w(this,t),o=i.apply(this,t),a=e||m(o),u=Math.max(o[1][0]-o[0][0],o[1][1]-o[0][1]),c=this.__zoom,f="function"==typeof n?n.apply(this,t):n,s=l(c.invert(a).concat(u/c.k),f.invert(a).concat(u/f.k));return function(t){if(1===t)t=f;else{var n=s(t),e=u/n[2];t=new Sb(e,a[0]-n[0]*e,a[1]-n[1]*e)}r.zoom(null,t)}})}function w(t,n){for(var e,r=0,i=h.length;r<i;++r)if((e=h[r]).that===t)return e;return new M(t,n)}function M(t,n){this.that=t,this.args=n,this.index=-1,this.active=0,this.extent=i.apply(t,n)}function N(){if(r.apply(this,arguments)){var t=w(this,arguments),n=this.__zoom,e=Math.max(c[0],Math.min(c[1],n.k*Math.pow(2,a.apply(this,arguments)))),i=Ot(this);if(t.wheel)t.mouse[0][0]===i[0]&&t.mouse[0][1]===i[1]||(t.mouse[1]=n.invert(t.mouse[0]=i)),clearTimeout(t.wheel);else{if(n.k===e)return;t.mouse=[i,n.invert(i)],Mr(this),t.start()}Pb(),t.wheel=setTimeout(function(){t.wheel=null,t.end()},v),t.zoom("mouse",o(b(_(n,e),t.mouse[0],t.mouse[1]),t.extent,f))}}function A(){if(!e&&r.apply(this,arguments)){var n=w(this,arguments),i=zt(t.event.view).on("mousemove.zoom",function(){if(Pb(),!n.moved){var e=t.event.clientX-u,r=t.event.clientY-c;n.moved=e*e+r*r>g}n.zoom("mouse",o(b(n.that.__zoom,n.mouse[0]=Ot(n.that),n.mouse[1]),n.extent,f))},!0).on("mouseup.zoom",function(){i.on("mousemove.zoom mouseup.zoom",null),jt(t.event.view,n.moved),Pb(),n.end()},!0),a=Ot(this),u=t.event.clientX,c=t.event.clientY;It(t.event.view),Cb(),n.mouse=[a,this.__zoom.invert(a)],Mr(this),n.start()}}function T(){if(r.apply(this,arguments)){var n=this.__zoom,e=Ot(this),a=n.invert(e),u=n.k*(t.event.shiftKey?.5:2),c=o(b(_(n,u),e,a),i.apply(this,arguments),f);Pb(),s>0?zt(this).transition().duration(s).call(x,c,e):zt(this).call(y.transform,c)}}function S(){if(r.apply(this,arguments)){var e,i,o,a,u=w(this,arguments),c=t.event.changedTouches,f=c.length;for(Cb(),i=0;i<f;++i)a=[a=Bt(this,c,(o=c[i]).identifier),this.__zoom.invert(a),o.identifier],u.touch0?u.touch1||(u.touch1=a):(u.touch0=a,e=!0);if(n&&(n=clearTimeout(n),!u.touch1))return u.end(),void((a=zt(this).on("dblclick.zoom"))&&a.apply(this,arguments));e&&(n=setTimeout(function(){n=null},p),Mr(this),u.start())}}function k(){var e,r,i,a,u=w(this,arguments),c=t.event.changedTouches,s=c.length;for(Pb(),n&&(n=clearTimeout(n)),e=0;e<s;++e)i=Bt(this,c,(r=c[e]).identifier),u.touch0&&u.touch0[2]===r.identifier?u.touch0[0]=i:u.touch1&&u.touch1[2]===r.identifier&&(u.touch1[0]=i);if(r=u.that.__zoom,u.touch1){var l=u.touch0[0],h=u.touch0[1],d=u.touch1[0],p=u.touch1[1],v=(v=d[0]-l[0])*v+(v=d[1]-l[1])*v,g=(g=p[0]-h[0])*g+(g=p[1]-h[1])*g;r=_(r,Math.sqrt(v/g)),i=[(l[0]+d[0])/2,(l[1]+d[1])/2],a=[(h[0]+p[0])/2,(h[1]+p[1])/2]}else{if(!u.touch0)return;i=u.touch0[0],a=u.touch0[1]}u.zoom("touch",o(b(r,i,a),u.extent,f))}function E(){var n,r,i=w(this,arguments),o=t.event.changedTouches,a=o.length;for(Cb(),e&&clearTimeout(e),e=setTimeout(function(){e=null},p),n=0;n<a;++n)r=o[n],i.touch0&&i.touch0[2]===r.identifier?delete i.touch0:i.touch1&&i.touch1[2]===r.identifier&&delete i.touch1;i.touch1&&!i.touch0&&(i.touch0=i.touch1,delete i.touch1),i.touch0?i.touch0[1]=this.__zoom.invert(i.touch0[0]):i.end()}return y.transform=function(t,n){var e=t.selection?t.selection():t;e.property("__zoom",Db),t!==e?x(t,n):e.interrupt().each(function(){w(this,arguments).start().zoom(null,"function"==typeof n?n.apply(this,arguments):n).end()})},y.scaleBy=function(t,n){y.scaleTo(t,function(){return this.__zoom.k*("function"==typeof n?n.apply(this,arguments):n)})},y.scaleTo=function(t,n){y.transform(t,function(){var t=i.apply(this,arguments),e=this.__zoom,r=m(t),a=e.invert(r),u="function"==typeof n?n.apply(this,arguments):n;return o(b(_(e,u),r,a),t,f)})},y.translateBy=function(t,n,e){y.transform(t,function(){return o(this.__zoom.translate("function"==typeof n?n.apply(this,arguments):n,"function"==typeof e?e.apply(this,arguments):e),i.apply(this,arguments),f)})},y.translateTo=function(t,n,e){y.transform(t,function(){var t=i.apply(this,arguments),r=this.__zoom,a=m(t);return o(kb.translate(a[0],a[1]).scale(r.k).translate("function"==typeof n?-n.apply(this,arguments):-n,"function"==typeof e?-e.apply(this,arguments):-e),t,f)})},M.prototype={start:function(){return 1==++this.active&&(this.index=h.push(this)-1,this.emit("start")),this},zoom:function(t,n){return this.mouse&&"mouse"!==t&&(this.mouse[1]=n.invert(this.mouse[0])),this.touch0&&"touch"!==t&&(this.touch0[1]=n.invert(this.touch0[0])),this.touch1&&"touch"!==t&&(this.touch1[1]=n.invert(this.touch1[0])),this.that.__zoom=n,this.emit("zoom"),this},end:function(){return 0==--this.active&&(h.splice(this.index,1),this.index=-1,this.emit("end")),this},emit:function(t){St(new Tb(y,t,this.that.__zoom),d.apply,d,[t,this.that,this.args])}},y.wheelDelta=function(t){return arguments.length?(a="function"==typeof t?t:Ab(+t),y):a},y.filter=function(t){return arguments.length?(r="function"==typeof t?t:Ab(!!t),y):r},y.touchable=function(t){return arguments.length?(u="function"==typeof t?t:Ab(!!t),y):u},y.extent=function(t){return arguments.length?(i="function"==typeof t?t:Ab([[+t[0][0],+t[0][1]],[+t[1][0],+t[1][1]]]),y):i},y.scaleExtent=function(t){return arguments.length?(c[0]=+t[0],c[1]=+t[1],y):[c[0],c[1]]},y.translateExtent=function(t){return arguments.length?(f[0][0]=+t[0][0],f[1][0]=+t[1][0],f[0][1]=+t[0][1],f[1][1]=+t[1][1],y):[[f[0][0],f[0][1]],[f[1][0],f[1][1]]]},y.constrain=function(t){return arguments.length?(o=t,y):o},y.duration=function(t){return arguments.length?(s=+t,y):s},y.interpolate=function(t){return arguments.length?(l=t,y):l},y.on=function(){var t=d.on.apply(d,arguments);return t===d?y:t},y.clickDistance=function(t){return arguments.length?(g=(t=+t)*t,y):Math.sqrt(g)},y},t.zoomTransform=Eb,t.zoomIdentity=kb,Object.defineProperty(t,"__esModule",{value:!0})});
```
