## Supplementary figures and images for "Exploring dimension-reduced embeddings with Sleepwalk"

### Fig_A.png

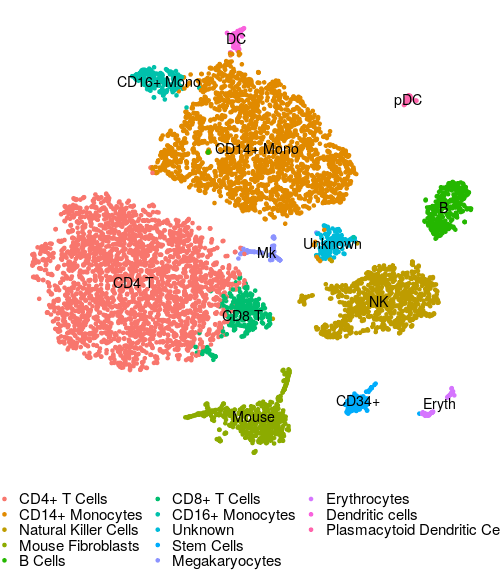
